# A chemical reaction network model of PURE

**DOI:** 10.1101/2023.08.14.553301

**Authors:** Zoila Jurado, Ayush Pandey, Richard M. Murray

**Affiliations:** Division of Engineering and Applied Science, California Institute of Technology, Pasadena, CA; Division of Biology and Biological Engineering, California Institute of Technology, Pasadena, CA

## Abstract

Cell-free expression systems provide a method for rapid DNA circuit prototyping and functional protein synthesis. While crude extracts remain a black box with many components carrying out unknown reactions, the PURE system contains only the required transcription and translation components for protein production. All proteins and small molecules are at known concentrations, opening up the possibility of detailed modeling for reliable computational predictions. However, there is little to no experimental data supporting the expression of target proteins for detailed protein models PURE models. In this work, we build a chemical reaction network transcription model for PURE protein synthesis. We compare the transcription models using DNA encoding for the malachite-green aptamer (MGapt) to measure mRNA production. Furthermore, we expand the PURE detailed translation model for an arbitrary set of amino acids and length. Lastly, we combine the transcription and the expanded translation models to create a PURE protein synthesis model built purely from mass-action reactions. We use the combined model to capture the translation of a plasmid encoding MGapt and deGFP under a T7-promoter and a strong RBS. The model accurately predicts the MGapt mRNA production for the first two hours, the dynamics of deGFP expression, and the total protein production with an accuracy within 10 %.

## Introduction

Cell-free protein synthesis (CFPS) systems are divided into two broad categories. The first widely used CFPS is based on cell lysate, first implemented in the 1960s to express synthetic RNAs to decipher the genetic code [1]. Cell lysate-based systems or TX-TL (transcription and translation) utilize cellular machinery harvested from the cell. After multiple stages of growth, lysis, and clarifying spins, an energy buffer with amino acids is added to the lysate to create a protein expression system [2, 3]. The widespread use of TX-TL, now commercially available, has its limitations. The research field is limited by batch-to-batch variability, affecting lifetime and total protein expression [4]. The batch-to-batch variability can result from multiple variables such as cell strain, optical density (OD) at the time of harvest, lysis method, energy mixture composition, and reagents batches. The ability to achieve ‘design–build–test’ cycles, similar to those found in other traditional engineering fields, is thus ultimately limited by the CFPS system’s predictability.

Over the last decade, multiple TX-TL protein expression models have been put forth and have shown to accurately model mRNA and protein production [5, 6, 7, 8]. However, these models lack the ability to predict expression without characterizing the models to their specific experimental data sets. Steps have been made towards more predictive models, modeling the behavior of whole circuits through the use of a software toolbox and characterizing components of the entire model [9]. Though these TX-TL models can help understand phenomena or estimate unknown parameters, they continue to be constrained by the unknown composition of the cellular lysate.

One advantage of using cellular lysate is the retention of biological pathways of the cell strain, such as glycolysis, allowing for energy regeneration. The extent to which cellular processes remain functional is still undetermined. Characterization of lysate as the next step for TX-TL modeling relies on LCMS to measure small molecules, proteins, and lipids to understand how much of the core metabolism is active, potential side reactions, and waste generation effects [10]. However, measuring all proteins, small molecules, and lipids, and mapping the chemical reactions associated with each in lysate, is difficult. Thus, having a universal, batch-independent, and detailed TX-TL model is unlikely.

The second category of CFPS utilizes purified components of transcription and translation machinery to reconstruct the “central dogma.” The first reported attempt of using purified proteins to express functional proteins was in 1977 [11], having limited success. Weissbach’s group provided a starting point to which Ganoza *et al.* [12], and Pavlov *et al.* [13, 14] attempted to use precharged aminoacyl-tRNAs or partially purified aminoacyl-tRNA synthetase alongside purified proteins. Shortly after, in 2001, Shimizu *et al.* [15] achieved successful protein production using PURE — Protein synthesis Using purified Recombinant Elements. The PURE system contains all transcription and translation proteins required for protein production at known concentrations. As a direct contrast to cellular lysate-based CFPS, this allows for batch-to-batch and inter-laboratory repeatability. Consequently, the PURE system presents an opportunity for detailed modeling, allowing for reliable computational predictions. Without requiring recharacterization for every experimental run, these models could be integrated into pipelines to prototype larger circuits using the PURE system. Nonetheless, even with complete control and knowledge of the composition, existing PURE models fall back to the phenomenological modeling of transcription and translation [5, 16, 17, 18, 19] by grouping all NTPs as one variable, not modeling each step of protein production, and employing Hill functions instead of chemical reaction equations.

In 2017, Shimizu’s group introduced a MATLAB model for the translation mechanisms in PURE [20, 21]. It is a detailed model that distinguishes between NTPs and incorporates each stage in the translation of an fMGG peptide. This computational model consists of 968 mass-action reactions and 241 species, including the 27 components that initialize the PURE system. Time courses of all components can be tracked in this model, which provides a valuable method to explore and systematically model the protein synthesis in PURE. But, fMGG is a small peptide, and extending this detailed model to the commonly used proteins in PURE is not straightforward. Explicitly writing each reaction and all possible species would be increasingly tedious as protein length increases. Moreover, since transcription is not modeled, this model is an incomplete description of the PURE system and cannot be experimentally validated. To address these limitations, in this paper, we demonstrate (1) a detailed model of PURE for the transcription of arbitrary DNA sequences with mechanistic details for each step in the transcription; (2) a generalization of Shimizu’s group translation model for arbitrary proteins; and (3) the experimental validation of a complete transcription and translation model of the PURE system.

## Results and Discussion

### Creating a chemical reaction network of transcription for any DNA sequence

To model the detailed step-by-step mechanistic process of transcription, we built a chemical reaction network (CRN). The chemical reactions of the transcription model were initially based on the reactions and rates proposed in the TX-TL model by Tuza *et al.* [22]. The user is required to input the DNA sequence of the desired protein, not including the promoter or terminator portions. To adapt and expand this model efficiently for any arbitrary mRNA sequence, we used a CRN compiler tool called BioCRNpyler [23].

BioCRNpyler is a Python-based software package that can easily compile CRN models from simple descriptions of the parts of the system. For the transcription model, we include a detailed description of the process in such a way that the transcription mechanisms can be adaptable according to the mRNA sequence being transcribed. BioCRNpyler also contains a library of parts and parameters that can be used to share parts of the model in other larger system models. We use the features in BioCRNpyler to generate species and reactions depending on user input and to store these mechanisms in a way that can be used in larger models.

We split transcription into three groups: initiation, elongation, and termination, as illustrated in Figure 1 for mRNA_length=*n*_. We model the detailed mechanisms for each group to include all PURE components’ interactions. Initiation steps include NTP degradation and a one-step GTP-dependent activation of T7 RNAP [24]:

**Figure 1:**
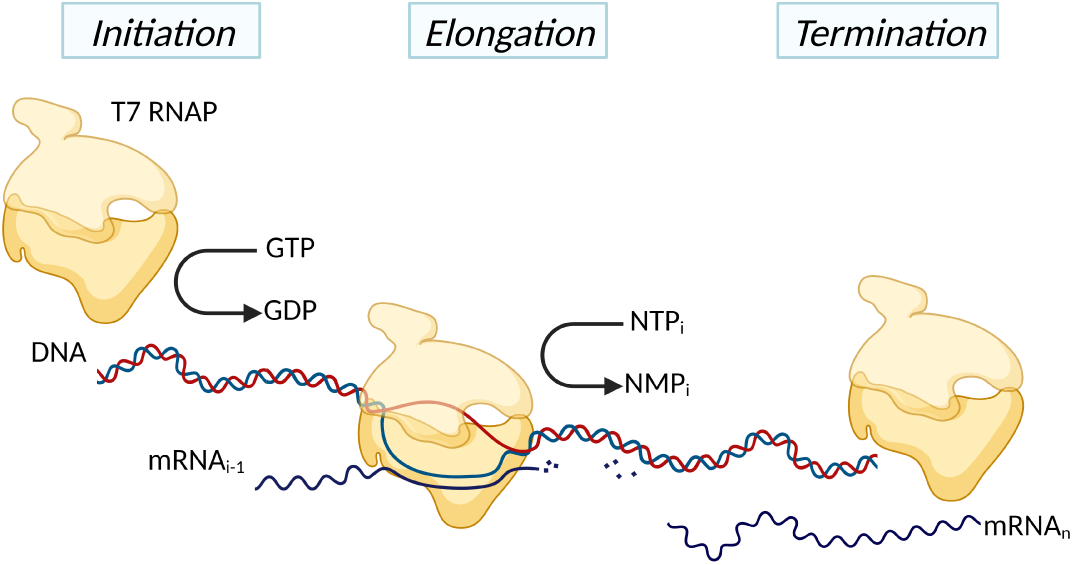
Schematic of mRNA synthesis with a reconstituted *E. coli* transcription system. We split the transcription reactions into three groups: initiation, elongation, and termination. Auxiliary reactions such as energy recycling or those explicitly related to translation are not included. Created with BioRender.com

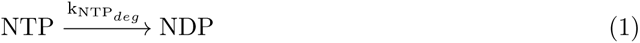

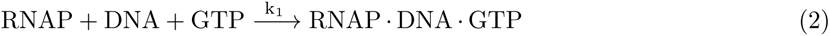

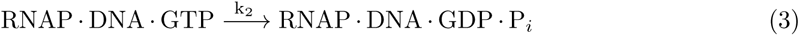

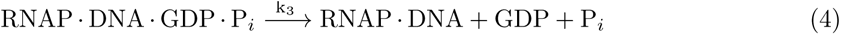

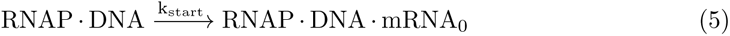

Elongation steps model each binding state separately for the addition of an NTP. The elongations steps are repeated for each nucleic acid addition along the growing mRNA chain of length *n*:

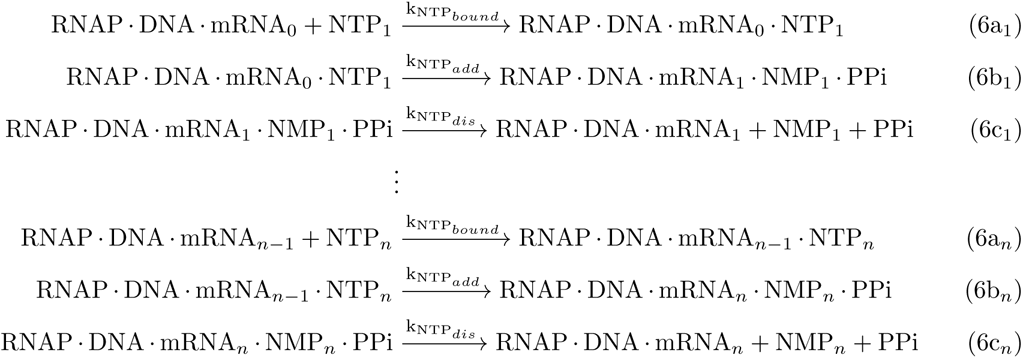

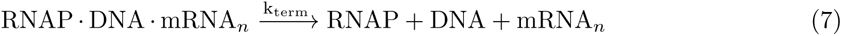

In our transcription reactions, we do not explicitly account for multiple polymerases simultaneously bound on a singular DNA strand. However, the effects of simultaneous transcription can be incorporated into the reaction rates. Additionally, no auxiliary reactions, such as NTP recycling, are incorporated into the transcription model.

### Validation of transcription model using pT7-MGapt-tT7 plasmid

To validate the transcription models, we measured the T7 RNAP-driven transcription of malachite-green aptamer (MGapt) without a ribosome binding site (RBS). The lack of RBS limits the reactions associated with translation, allowing the focus to be on transcription. A 10 µL reaction was done in triplicate using New England Biolabs (NEB) PURExpress In Vitro Protein Synthesis Kit. The reaction contained 5 nm of plasmid DNA with construct pT7-MGapt-tT7, 10 µm malachite green oxalate, and 8 units of RNase inhibitor. The samples were mixed in PCR tubes with 5% excess, then 10 µL was added to a 384-well plate and read for 3 hours at 37 °C in a BioTek H1MF plate reader. The total amount of mRNA was calculated using calibration curves (Figure S3A).

To parameterize the CRN model constructed using BioCRNpyler, we initialized the parameters using the values from Tuza *et al.* [22]. The initial conditions were taken from PURE components [25] for NTPs and T7 RNAP concentrations. To get reliable predictions of transcription in PURE, we trained the model using experimental data for the MGapt fluorescence. However, this model training using parameter identification is not straightforward as the CRN model we have constructed has many species and parameters. Since data is only available for one species in the model, it is crucial to assess the empirical identifiability of the model parameters. Finally, to account for the intrinsic noise observed in the experimental data, we use Bayesian inference to obtain a distribution of possible parameter values given the experimental data. To achieve both of these tasks (assessing identifiability and finding posterior parameter distributions), we use a biological data analysis pipeline [26] using the Python package Bioscrape [27]. This pipeline provides an effective interface to the BioCRNpyler model to run these analyses.

To choose the parameters to identify, we use the local sensitivity analysis of all species in the model against all parameters and for all time. The sensitivity analysis heatmap for the pT7-MGapt-tT7 model is shown in Figure 2A. We choose the parameters that are most sensitive in the fluorescence of the MGapt output. Out of the nine reaction rates highlighted in equations (1)-(7), we find that the most sensitive parameters are *k*_start_ and *k*_2_. These parameters correspond to the start of the transcription initiation and the formation rate of the RNAP-bound GDP and phosphate complex, respectively.

**Figure 2:**
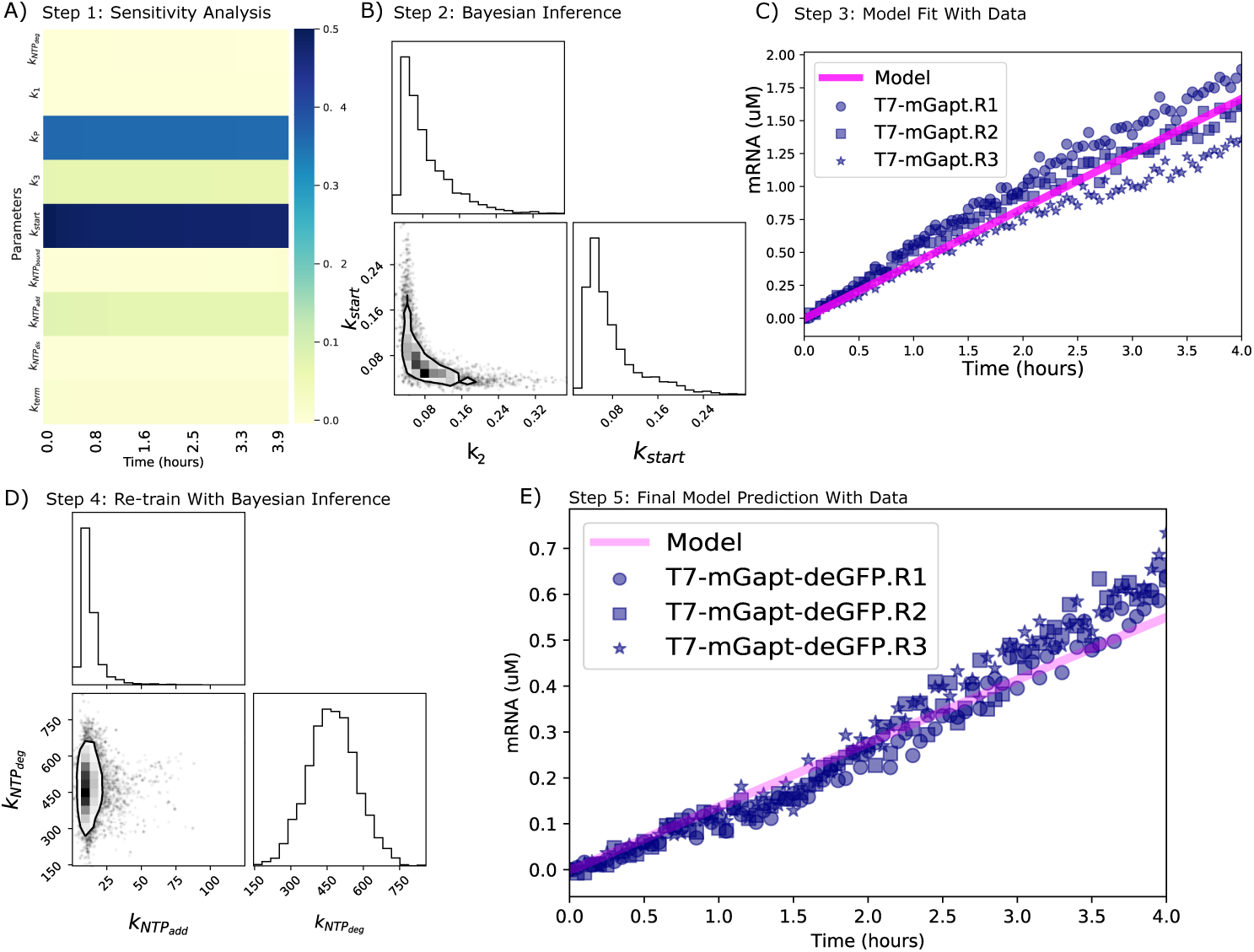
Modeling, analysis, and parameter learning for the PURE transcription model. A) Step 1: The sensitivity of the MGapt fluorescence to the CRN model parameters for all time. B) Step 2: The posterior distributions of parameters obtained after running Bayesian inference on *k*_2_ and *k*_start_. The corner plot depicts the covariance of the two parameters with the contour showing the 75% probability region for the parameter values. C) Step 3: With parameter values for *k*_2_ and *k*_start_ sampled from the posterior distributions, the model simulations (in magenta) are shown alongside the experimental data for three biological replicates (scattered points). D) Step 4: For accurate predictions, we re-train the model parameters with the data for the pT7-MGapt-UTR1-deGFP-tT7 construct. We infer the posterior parameter distributions for *k*_NTPadd_ *k*_NTPdeg_ using the MGapt fluorescence. E) Step 5: With parameter values drawn from the posterior distributions shown in (D), the model predictions (magenta) are shown alongside the experimental data.

We use the Bayesian inference tools in Bioscrape to identify the posterior parameter distributions for these two parameters. The corner plot in Figure 2B shows the posterior parameter distributions and their covariance. This chart provides us with a distribution to sample from when predicting the output using the fitted model. The model simulations with parameter values drawn from the posterior along with the experimental data are shown in Figure 2C. The final parameter values used are given in Table S1.

### Analysis of transcription model using pT7-MGapt-UTR1-deGFP-tT7

Transcription does not occur independent of translation in PURE. To further validate our transcription model, we proceeded to a coupled transcription-translation system. Significantly, to minimize interactions of the MGapt with the physical mechanism of translation, the MGapt preceded the RBS. Following the same protocol for the transcription-only system, we replaced pT7-MGapt-tT7 with pT7-MGapt-UTR1-deGFP-tT7 to measure the total mRNA production and deGFP expression. The total amount of deGFP was calculated using calibration curves (Figure S3B).

With the identified parameters from the pT7-MGapt-tT7 PURE transcription model, we found that the model did not accurately predict the MGapt fluorescence observed experimentally for the transcription of the longer mRNA sequence in pT7-MGapt-UTR1-deGFP-tT7. The model for pT7-MGapt-UTR1-deGFP-tT7 is around ten times larger than the model for the transcription of MGapt only. The model for pT7-MGapt-tT7 has 285 species and 276 parameters, while the model for the longer mRNA sequence of pT7-MGapt-UTR1-deGFP-tT7 has 2429 species and 2424 parameters. We found that the predictions from the larger model do not match the experimental data. This is expected because we did not identify all parameters in the transcription model in the previous step. From the sensitivity analysis of the larger model, we find that *k*_NTP_*_add_* and *k*_NTPdeg_ are two suitable candidate parameters to infer from the data. Following a similar pipeline as before, we find the posterior parameter distributions for the two parameters (shown in 2D). Finally, we simulate the model predictions by sampling the values of the parameters from the posterior probability distributions and plot these predictions alongside the experimental data shown in Figure 2E.

Summarizing the computational analysis, we utilized transcription parameters based on results from the literature. Then we inferred *k*_start_, *k*_2_, *k*_NTP_*_add_*, and *k*_NTPdeg_ in the PURE TX-models using the experimental data. The final trained model accurately predicts the transcription activity of the DNA sequence for which the transcription reactions were generated.

### Expansion of PURE translation model for an arbitrary set of amino acids

The MATLAB model for the PURE translation [20, 21] is limited to the case of the fMGG peptide. The model consists of 968 reactions and 241 species, all explicitly written out. As a result, the ability to change or extend the peptide is labor-some and futile. To make peptide variation more tractable, we first converted the MATLAB fMGG translation model to Python using BioCRNpyler [23]. The comparison between the BioCRNpyler model (magenta line) and MATLAB model (blue circles) can be seen in Figure 3A, with the error between the two in orange. While retaining all the reactions and parameters from the spreadsheet-based MATLAB model, the translation to Python enables user-friendly scripting and loops to iterate over an arbitrary length of peptide.

**Figure 3:**
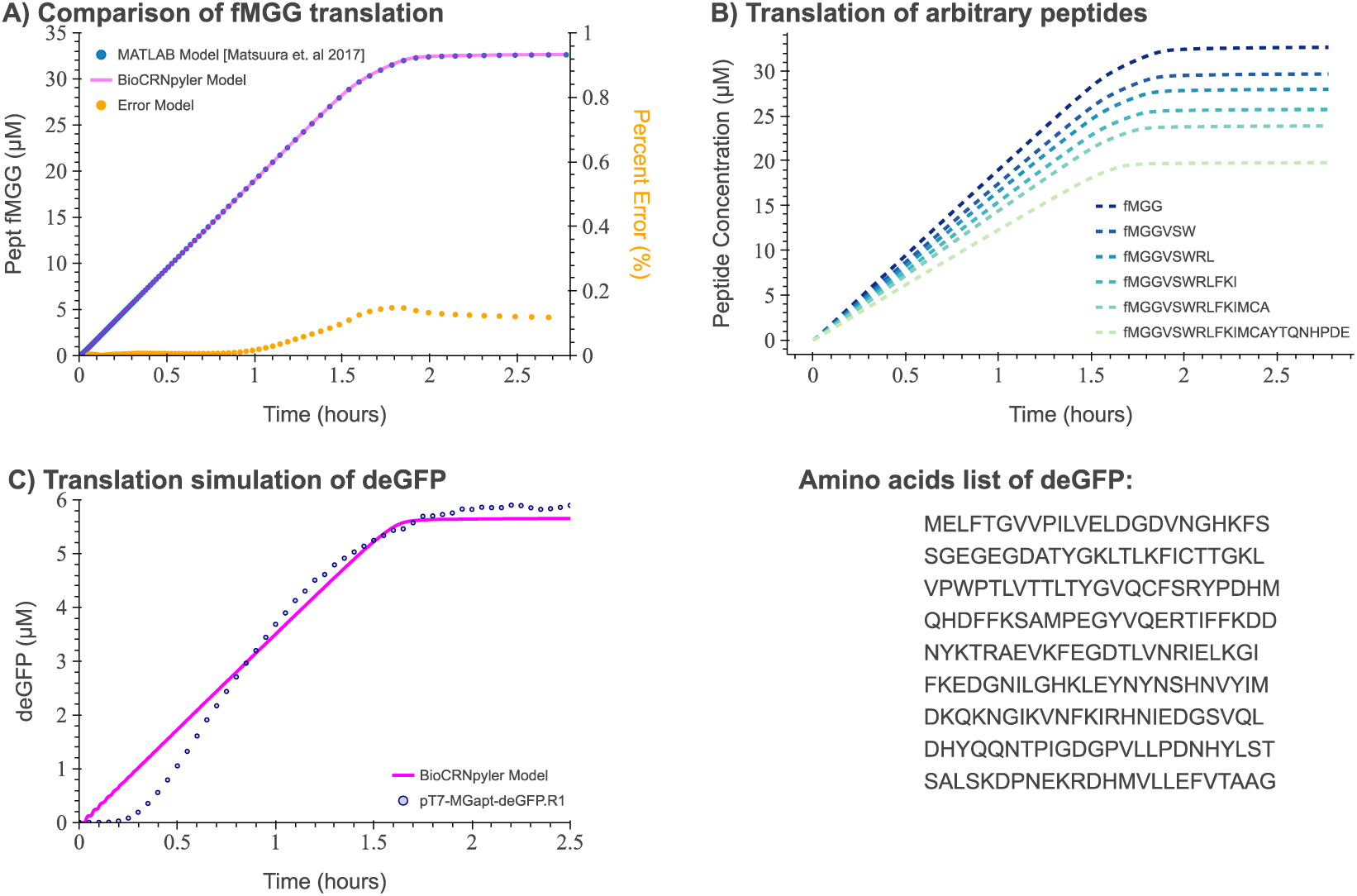
A) Comparison of original MATLAB model to BioCRNpyler model. BioCRNpyler model (magenta line) and MATLAB model (blue circles) overlap, and the difference between models in (orange circles) on a secondary axis. B) Expansion of BioCRNpyler with different amino acids to the original fMGG peptide. C) Left: Translation model prediction of deGFP (magenta line) overlaying experimental result (blue circles). Right: Amino acid list used as input for the BioCRNpyler translation model.

The expanded PURE translation model in BioCRNpyler was built by adding different amino acids one by one until all 20 amino acids were incorporated (see Figure 3B). As expected, the amount of the final peptide decreases as the amino acid chain lengthens. Finally, the model was expanded for repeated amino acids and arbitrary amino acid sequences of length greater than 21. The translation of green fluorescent protein (deGFP) (mRNA*_n_*=805, Pept*_n_*=226) was modeled using the BioCRNpyler translation only-model with the initial conditions adopted from the MATLAB model.

Initial conditions for tRNAs and amino acids not associated with Met or Gly were absent in the MATLAB model. So, we set the initial conditions of the Gly-amino acids and tRNAs as given in Table S2 based on PURE concentration published by Kazuta *et al.* [28]. Modeling the translation of an experimentally relevant protein such as deGFP enabled us to compare the PURE models to experimental results (see Figure 3C, model in magenta overlays three experimental repeats in blue). Using the extended translation model, we set mRNA*_n_* to 0.126 µm, such that the total deGFP expression is comparable to experimental results of 6 µm, as seen in Figure 3C. The translation-only model does not accurately predict the first hour of protein expression. This discrepancy is expected since we start with a nonzero mRNA. In the combined transcription and translation model, mRNA would be produced, resulting in the delay of protein expression. Therefore, the immediate production of deGFP was expected, but further supports the need for a coupled transcription and translation model.

### Combination of PURE models

Finally, with the successful construction of separate transcription and translation models for arbitrary sequences, BioCRNpyler easily allows for the combination of the two models. Since we were unable to obtain the initial conditions of PURExpress from NEB and found contradicting concentrations across literature, the initial conditions were set to the Version 7 PURE concentration published in Table S1 [28]. The one exception is the small molecule creatine phosphate’s (CP) initial concentration is 10 mm; see Supplementary Information Table S2 for the initial conditions of all of the proteins and amino acids.

We found that combining the two models did not produce an accurate production of deGFP. We propose this discrepancy because of the translation model’s lack of multiple ribosome loading on an mRNA or mean ribosome load. Mean ribosome load (MRL) is a metric of the average ribosome count associated with an mRNA [29]. To incorporate MRL, we modified the termination reaction in equation (7) to 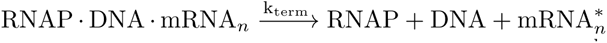, where species mRNA*^∗^* is an intermediate species. Additionally, we added a reaction 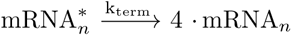 serving as a linker between the transcription model and translation model and accounting for MRL. The adjusted termination step resulted in the following:

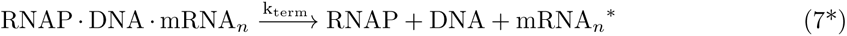

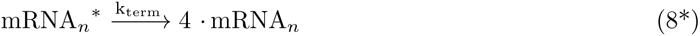

The scaling factor of 4 was determined empirically by simulating the combined system with scaling factors spanning from 1-10, as shown in Figure S1. The scaling factor is best interpreted as the translation of mRNA by four ribosomes simultaneously, on average. The average number of ribosomes associated with the given mRNA is dependent on the RBS and the mRNA 5’ end. Without quantifying the ribosome load of a particular DNA sequence, this feature enables other scaling factors to adjust the number of ribosomes simultaneously translating. This can be achieved by simply modifying the empirical factor of 4.

Finally, to account for protein folding the following reaction was added:

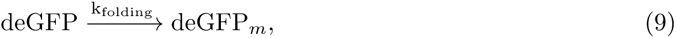

where *k_folding_*= 600*s^−^*^1^ [30], completing the combined PURE model. The total number of reactions of the combined transcription and translation model is 6988 with 6280 species. Similar to the translation model by Matsuura *et al.*, our model tracks all species. The combined model predictions are shown in Figure 4 overlaid with the experimental data (in blue shapes).

**Figure 4:**
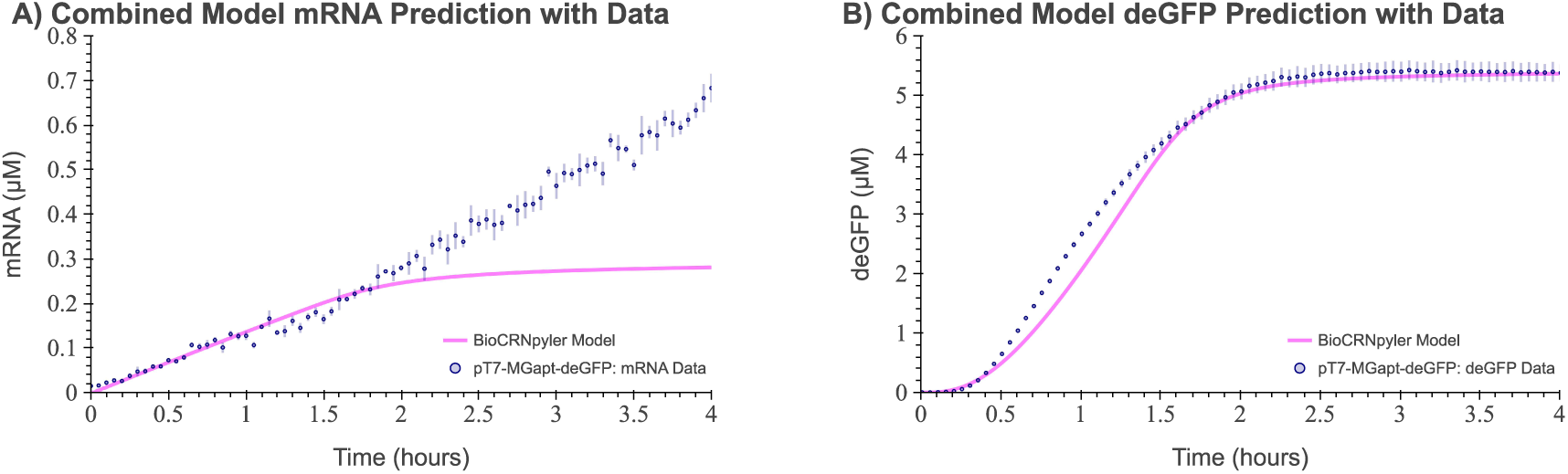
The combined transcription and translation model for pT7-MGapt-UTR1-deGFP-tT7, DNA=5 nm, with experimental result in PURExpress. A) Modeled mRNA production in the combined model (magenta line) overlaying with experimental data, three replicates (blue circles and blue error bars). B) Modeled deGFP expression in the combined model (magenta line) overlaying with experimental data, three replicates (blue circles and blue error bars).

As seen in Figure 4A, the model of mRNA production is consistent with experiments until approximately 2 hours. In the model, mRNA stabilizes around 2 hours, indicating that the mRNA production stops. The cessation of transcription at approximately 2 hours aligns with the expected lifespan of transcription in PURE. However, experimental results with NEB PURE show that MGapt fluorescence continues to increase past 2 hours. This can either indicate that mRNA production continues past 2 hours and NTPs become limited earlier in the model than in the experiments or that undetermined chemical reactions increase MGapt fluorescence over time.

The expression pattern of deGFP more closely resembles actual experimental data in terms of when its expression ceases, compared to the translation-only model presented earlier. This is clear in comparing the “kink” in Figure 3C when the deGFP stops expressing with a much smoother transition in Figure 4B. Additionally, the slopes of deGFP expression do not precisely match the experimental data. This suggests that even with the level of detail in the model of transcription and translation, there still are unidentified reactions. Despite these minor differences, we observe that both mRNA and deGFP are accurately modeled according to the properties of our interest in building cell-free biological circuits. The results are comparable for the specific protein, deGFP, within the 2 h reaction window, and the expression levels match the absolute values obtained in the experiments (see Figure 4B).

Finally, from the heatmaps in Figure 5A, we can infer that ATP and GTP are the likely limiting energy carriers in PURE. The consumption of GTP and ATP results in the discontinuation of mRNA and protein production in the model, supporting the need for further development of energy generation and recycling in CFPS. Additionally, Figure 5B shows that amino acids are not limiting in PURE. Figure 5B represents only free amino acids, those not in a complex with their respective tRNA. The excess of amino acids is apparent, with none of the 20 amino acid concentrations falling below 20 µm.

**Figure 5:**
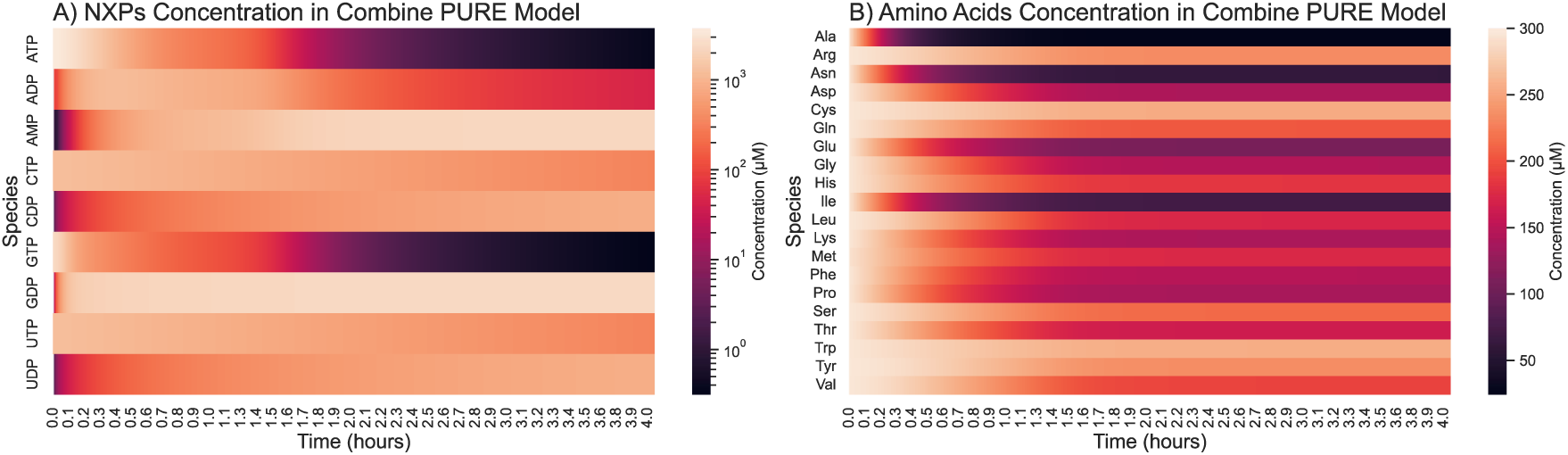
The concentrations of NXPs and amino acids over time in the combined transcription and translation model for pT7-MGapt-UTR1-deGFP-tT7, DNA= 5 nm. A) Concentrations (µm) of ATP, ADP, AMP, CTP, CDP, GTP, GDP, UTP, and UDP over simulation time using log-normal scale. B) Concentrations (µm) of all amino acids over simulation time.

## Conclusion

We have proposed a complete transcription and translation model of cell-free protein expression for arbitrary proteins in the PURE system. The existing model of translation in PURE from Matsuura *et al.* [20] only modeled the translation of the fMGG peptide. In this paper, we have developed a transcription model that incorporates each step of the growing mRNA strand and the specific NTP required to transcribe the mRNA from any given DNA sequence. Further, we have expanded on the PURE translation model so that proteins with any given amino acid sequence can be modeled. By combining the transcription and translation models that we developed, we present a complete model of protein expression in PURE cell-free systems. Using our approach, it is possible to create mathematical models of the expression of experimentally relevant proteins in PURE.

We validated our models using experimental data. We identified a distribution of possible parameters in the model with the experimental data for MGapt fluorescence. We showed that the validated model accurately predicts the MGapt fluorescence for two different plasmids — one where MGapt is expressed in an isolated manner and another where it is expressed together with deGFP. The combined model of the PURE cell-free system with the validated transcription model and the extended translation model was then used to predict deGFP expression. The model predictions agree with the experimental results.

This detailed model of the PUREexpress system is a step towards cell-free protein synthesis CFPS predictability. It can be used as a platform to guide the design of future probing experiments with biological circuits in PURE. Utilizing this detailed model with OnePot PURE [31], a version of PURE where all 36 proteins are co-cultured and purified together, can help circumvent batch-to-batch and inter-laboratory variability problems seen in extract-based systems. Using quantitative proteomics [32] initial conditions of each OnePot batch can be measured and simulated to predict batch yield and improve the reproducibility of OnePot PURE. Beyond this, our model can be instrumental in identifying potential directions for further research, particularly regarding the coupling between the transcription and the translation mechanisms.

The model can be improved by incorporating reactions that model potential inhibitory interactions, such as inorganic phosphates, pyrophosphates, and pH. Additional reactions could include DNA polymerases for DNA replication and ribosomal loading reactions beyond the linker reactions proposed in equations 7* and 8*. Modeling of incomplete transcription and translation products[33] could also be beneficial in improving the accuracy of the predictions. We hope this model will bring multiple research groups together by compiling data from fluorescent or LCMS measurements of different experimental conditions. Together we can begin to build a library of characterized parts for PURE or OnePot PURE to achieve a robust ‘design–build–test’ cycle.

## Materials and Methods

### Computational Modeling and Simulations

In this paper, we build on a previously published deterministic model of protein synthesis of a formyl-Met-Gly-Gly tripeptide (fMGG) [20]. This model is based on components of an *Escherichia coli*-based reconstituted *in vitro* translation system [15]. We use the chemical reaction network (CRN) formalism to create the detailed mechanistic model using a CRN compiler called BioCRNpyler [23]. The computational model has been developed such that it can synthesize peptides of arbitrary lengths and sequences. Our CRN model initially begins with a minimum of 86 components while creating other intermediate components and reactions depending on the sequence of the desired peptide.

BioCRNpyler outputs the model in the standard biological modeling language called the Systems Biology Markup Language (SBML) [34]. The exported SBML files can be simulated with any compatible SBML simulator. We choose to use the Bioscrape [27] Python package to simulate the SBML models. The CRN model is converted to an ordinary differential equation by Bioscrape and solved using Python odeint for desired initial conditions. To convert the CRN to an ODE, each reaction rate is written using the mass-action propensity [35]. We use Bioscrape because it supports sensitivity analysis and Bayesian inference tools for SBML models in addition to the model simulations. For each SBML model, we run the local sensitivity analysis to obtain the sensitivity of the measured species with all parameters and at all times. Then, we aim to identify the most sensitive parameters for the model using the experimental data. We perform the parameter identification using a Bayesian inference algorithm implemented in Bioscrape with emcee Python package [36]. Given the experimental data, we obtain a probability distribution for each identified parameter with Bayesian inference. Model simulations with parameter values sampled from these posterior probability distributions are then plotted against the experimental data to evaluate the quality of the model predictions. These posterior probability distributions also quantify the uncertainty in the data, which is an important advantage of Bayesian inference methods.

Summarizing the computational analysis for the PURE transcription model, we identified four out of nine parameters using the experimental data. The final trained model predicts the transcription from the plasmid’s DNA sequence. The translation kinetic parameters for simulation were taken from the computational model simulating the synthesis of a fMGG [20].

All parameters and initial conditions for the translation model were obtained from the PURE simulator [20] website (https://sites.google.com/view/puresimulator). The parameter values, initial conditions, and Bayesian inference chains are available in the Supplementary Information. All calculations and plotting were performed using the standard stack of Python packages – NumPy [37], SciPy [38], Pandas [39], Matplotlib [40], Bokeh [41], and Seaborn [42]. The run time for each simulation depends on the protein length for which the model is created. The simulation time varies from less than a second (for a model with around 200 species and parameters) to a few minutes (for a model with about 3000 species and parameters) on a personal computer running Intel i7-6700 2.6 GHz with 16GB of RAM. The simulation times and Bayesian inference routines can be sped up by around 10x by running the model simulations on a high-performance computing cluster.

### Plasmids

The original deGFP DNA plasmid, pTXTL-T7p14-mGapt, and pTXTL-T7p14-deGFP were purchased from Arbor Biosciences myTXTL Toolbox 2.0 plasmid collection [43]. The deGFP protein is on a T7 promoter with a UTR1 ribosome binding site (RBS). A malachite-green aptamer (MGapt) was cloned between the promoter and the RBS site using primers from Integrated DNA Technologies. Forward and reverse primers used can be found in Supplementary Information Table S3.

Both plasmids were first transformed into JM109 cells and grown overnight on plates with carbenicillin resistance at 37 °C. Colonies of each plate were picked and grown in 4.5 mL LB with 4.5 µL of carbenicillin (100 mg/µL) overnight. A glycerol stock of each plasmid was made with 500 µL of liquid culture and 500 µL of 50 % glycerol, and the rest was miniprepped using Qiagen Miniprep Kit. Before use in the cell-free reaction, all plasmids underwent an additional PCR purification step using a QiaQuick column (Qiagen), which removed excess salt and were eluted and stored in nuclease-free water at 4 °C for short-term storage and -20 °C for long-term storage.

### PURE reactions and fluorescence measurements

PURE reactions were mixed by following the protocol for NEB PURExpress (E6800), adjusted for a 10 µL reaction, and allowed to run in a 384-well plate (Nunc) at 37 °C. 5 nm of DNA was used, unless otherwise stated, 8 units of RNAse inhibitor (NEB), and 10 µm of malachite-green dye was added to each reaction.

Fluorescence measurements were read in a Synergy H1 plate reader (Biotek) at 3 min intervals using excitation/emission wavelengths set at 610/650 nm (MGapt) at gain 150 and 485/515 nm (deGFP) at gain 61. All samples were read in the same plate reader, and for deGFP relative fluorescence units (RFUs) were converted to nm of protein using a purified eGFP standard by following the protocol in paper [43]. Calibration curves for MGapt and deGFP are shown in Figure S3.

## Supporting information

Supporting scripts, data, and simulation results.

## Acknowledgments

We thank Anthony Chiang (Caltech SURF 2022 student) for motivating the follow-through of this project and Yoshihiro Shimizu (RIKEN Center for Biosystems Dynamics Research) for directing us to the initial concentrations for PURE. Research supported by the National Science Foundation award number 2152267 and the Air Force Office of Scientific Research (AFOSR) under MURI grant FA9550-22-1-0316. The computations presented here were conducted in the Resnick High Performance Computing Center, a facility supported by the Resnick Sustainability Institute at the California Institute of Technology, Pasadena, CA, USA.

## Notes

### Competing Interest Statement

The authors have declared no competing interest.

## References

[1] Marshall W. Nirenberg and J. Heinrich Matthaei. The dependence of cell-free protein synthesis in E. coli upon naturally occurring or synthetic polyribonucleotides. PNAS, 47(10):1588–1602, 1961.

[2] Zachary Z. Sun, Clarmyra A. Hayes, Jonghyeon Shin, Filippo Caschera, Richard M. Murray, and Vincent Noireaux. Protocols for implementing an Escherichia coli based TX-TL cell-free expression system for synthetic biology. JoVE, page e50762, 2013.

[3] Max Z. Levine, Nicole E. Gregorio, Michael C. Jewett, Katharine R. Watts, and Javin P. Oza. Escherichia coli-based cell-free protein synthesis: Protocols for a robust, flexible, and accessible platform technology. JoVE, page e58882, 2019.

[4] Stephanie D. Cole, Kathryn Beabout, Kendrick B. Turner, Zachary K. Smith, Vanessa L. Funk, Svetlana V. Harbaugh, Alvin T. Liem, Pierce A. Roth, Brian A. Geier, Peter A. Emanuel, et al. Quantification of interlaboratory cell-free protein synthesis variability. ACS Synth. Biol., 8(9):2080–2091, 2019.

[5] Ryan Marshall and Vincent Noireaux. Quantitative modeling of transcription and translation of an all-*E. coli* cell-free system. Sci Rep, 9:11980, 2019.

[6] Dan Siegal-Gaskins, Zoltán A. Tuza, Jongmin Kim, Vincent Noireaux, and Richard M. Murray. Gene circuit performance characterization and resource usage in a cell-free “breadboard”. ACS Synth. Biol., 3(6):416–425, 2014.

[7] Eyal Karzbrun, Jonghyeon Shin, Roy H. Bar-Ziv, and Vincent Noireaux. Coarse-grained dynamics of protein synthesis in a cell-free system. Physical Review Letters, 106(4):048104, 2011.

[8] Fabio Chizzolini, Michele Forlin, Noël Yeh Martín, Giuliano Berloffa, Dario Cecchi, and Sheref S. Mansy. Cell-free translation is more variable than transcription. ACS Synth. Biol., 6(4):638–647, 2017.

[9] Vipul Singhal, Zoltán A. Tuza, Zachary Z. Sun, and Richard M. Murray. A MATLAB toolbox for modeling genetic circuits in cell-free systems. Synthetic Biology, 6(1):ysab007, 2021.

[10] William Poole. Compilation and Inference with Chemical Reaction Networks. PhD thesis, California Institute of Technology, Pasadena, CA USA, 2022.

[11] Hsiang-Fu Kung, Betty Redfield, Benjamin V. Treadwell, Barnet Eskin, Carlos Spears, and Herbert Weissbach. DNA-directed in vitro synthesis of beta-galactosidase. Studies with purified factors. Journal of Biological Chemistry, 252(19):6889–6894, 1977.

[12] M. Clelia Ganoza, Christina Cunningham, and Robert M. Green. Isolation and point of action of a factor from Escherichia coli required to reconstruct translation. Proceedings of the National Academy of Sciences, 82(6):1648–1652, 1985.

[13] Michael Yu. Pavlov and Måns Ehrenberg. Rate of translation of natural mRNAs in an optimizedin VitroSystem. Archives of Biochemistry and Biophysics, 328(1):9–16, 1996.

[14] Michael Yu. Pavlov, David V. Freistroffer, Valérie Heurgué-Hamard, Richard H. Buckingham, and Måns Ehrenberg. Release factor RF3 abolishes competition between release factor RF1 and ribosome recycling factor (RRF) for a ribosome binding site. Journal of Molecular Biology, 273(2):389–401, 1997.

[15] Yoshihiro Shimizu, Akio Inoue, Yukihide Tomari, Tsutomu Suzuki, Takashi Yokogawa, Kazuya Nishikawa, and Takuya Ueda. Cell-free translation reconstituted with purified components. Nature Biotechnology, 19(8):751–755, 2001.

[16] Nadanai Laohakunakorn, Laura Grasemann, Barbora Lavickova, Grégoire Michielin, Amir Shahein, Zoe Swank, and Sebastian J. Maerkl. Bottom-up construction of complex biomolecular systems with cell-free synthetic biology. Frontiers in Bioengineering and Biotechnology, 8, 2020.

[17] Anne Doerr, Elise De Reus, Pauline Van Nies, Mischa Van der Haar, Katy Wei, Johannes Kattan, Aljoscha Wahl, and Christophe Danelon. Modelling cell-free rna and protein synthesis with minimal systems. Physical Biology, 16(2):025001, 2019.

[18] Tobias Stögbauer, Lukas Windhager, Ralf Zimmer, and Joachim O. Rädler. Experiment and mathematical modeling of gene expression dynamics in a cell-free system. Integrative Biology, 4(5):494–501, 2012.

[19] Fabio Mavelli, Roberto Marangoni, and Pasquale Stano. A simple protein synthesis model for the PURE system operation. Bulletin of Mathematical Biology, 77:1185–1212, 2015.

[20] Tomoaki Matsuura, Naoki Tanimura, Kazufumi Hosoda, Tetsuya Yomo, and Yoshihiro Shimizu. Reaction dynamics analysis of a reconstituted Escherichia coli protein translation system by computational modeling. Proceedings of the National Academy of Sciences, 114(8):E1336–E1344, 2017.

[21] Tomoaki Matsuura, Kazufumi Hosoda, and Yoshihiro Shimizu. Robustness of a reconstituted Escherichia coli protein translation system analyzed by computational modeling. ACS Synth. Biol., 7(8):1964–1972, 2018.

[22] Zoltán A. Tuza, Dan Siegal-Gaskins, Jongmin Kim, and Gábor Szederkényi. Analysis-based parameter estimation of an *in vitro* transcription-translation system. In Eur. Control Conf. 2015, page 1554–1560, 2015.

[23] William Poole, Ayush Pandey, Andrey Shur, Zoltán A. Tuza, and Richard M. Murray. BioCRNpyler: Compiling chemical reaction networks from biomolecular parts in diverse contexts. PLOS Computational Biology, 18(4):e1009987, 2022.

[24] Yiping Jia and Smita S. Patel. Kinetic mechanism of GTP binding and RNA synthesis during transcription initiation by bacteriophage T7 RNA polymerase. Journal of Biological Chemistry, 272(48):30147–30153, 1997.

[25] Yoshihiro Shimizu, Yutetsu Kuruma, Bei-Wen Ying, So Umekage, and Takuya Ueda. Cell-free translation systems for protein engineering. The FEBS Journal, 273(18):4133–4140, 2006.

[26] Ayush Pandey, Makena L. Rodriguez, William Poole, and Richard M. Murray. Characterization of integrase and excisionase activity in a cell-free protein expression system using a modeling and analysis pipeline. ACS Synth. Biol., 12(2):511–523, 2023.

[27] Ayush Pandey, William Poole, Anandh Swaminathan, Victoria Hsiao, and Richard M. Murray. Fast and flexible simulation and parameter estimation for synthetic biology using bioscrape. Journal of Open Source Software, 8(83):5057, 2023.

[28] Yasuaki Kazuta, Tomoaki Matsuura, Norikazu Ichihashi, and Tetsuya Yomo. Synthesis of milligram quantities of proteins using a reconstituted in vitro protein synthesis system. Journal of Bioscience and Bioengineering, 118(5):554–557, 2014.

[29] Alexander Karollus, Zĭga Avsec, and Julien Gagneur. Predicting mean ribosome load for 5’UTR of any length using deep learning. PLoS Comput. Biol., 17(5):e1008982, 2021.

[30] Hendrik Dietz and Matthias Rief. Exploring the energy landscape of GFP by single-molecule mechanical experiments. PNAS, 101(46):16192–16197, 2004.

[31] Barbora Lavickova and Sebastian J. Maerkl. A simple, robust, and low-cost method to produce the PURE cell-free system. ACS Synth. Biol., 8(2):455–462, 2019.

[32] Svitlana Rozanova, Katalin Barkovits, Miroslav Nikolov, Carla Schmidt, Henning Urlaub, and Katrin Marcus. *Quantitative Mass Spectrometry-Based Proteomics: An Overview*, pages 85–116. Springer US, New York, NY, 2021.

[33] Jun Li, Chi Zhang, Poyi Huang, Erkin Kuru, Eliot T. C. Forster-Benson, Taibo Li, and George M. Churcha. Dissecting limiting factors of the protein synthesis using recombinant elements (PURE) system. Translation, 5(1):e1327006, 2017.

[34] Michael Hucka, Andrew Finney, Herbert M. Sauro, Hamid Bolouri, John C. Doyle, Hiroaki Kitano, Adam P. Arkin, Benjamin J. Bornstein, Dennis Bray, Athel Cornish-Bowden, et al. The systems biology markup language (sbml): a medium for representation and exchange of biochemical network models. Bioinformatics, 19(4):524–531, 2003.

[35] Domitilla Del Vecchio and Richard M. Murray. Biomolecular Feedback Systems. Princeton University Press Princeton, NJ, 2015.

[36] Daniel Foreman-Mackey, David W. Hogg, Dustin Lang, and Jonathan Goodman. emcee: the MCMC hammer. Publications of the Astronomical Society of the Pacific, 125(925):306, 2013.

[37] Charles R. Harris, K. Jarrod Millman, Stéfan J. Van Der Walt, Ralf Gommers, Pauli Virtanen, David Cournapeau, Eric Wieser, Julian Taylor, Sebastian Berg, Nathaniel J. Smith, et al. Array programming with NumPy. Nature, 585(7825):357–362, 2020.

[38] Pauli Virtanen, Ralf Gommers, Travis E Oliphant, Matt Haberland, Tyler Reddy, David Cournapeau, Evgeni Burovski, Pearu Peterson, Warren Weckesser, Jonathan Bright, et al. Scipy 1.0: fundamental algorithms for scientific computing in Python. Nature Methods, 17(3):261–272, 2020.

[39] Wes McKinney et al. pandas: a foundational Python library for data analysis and statistics. Python for High Performance and Scientific Computing, 14(9):1–9, 2011.

[40] J. D. Hunter. Matplotlib: A 2D graphics environment. Computing in Science & Engineering, 9(3):90–95, 2007.

[41] Bokeh Development Team. Bokeh: Python visualization library, 2023.

[42] Michael L. Waskom. Seaborn: statistical data visualization. Journal of Open Source Software, 6(60):3021, 2021.

[43] Arbor Biosciences. myTXTL T7 Expression Kit. Michigan, United States of America, 2019.

