## Supporting Figures S1-S3; Supporting Tables S1-S3 for "A chemical reaction network model of PURE"

August 14, 2023

### 1 Parameter Values

The parameter values for the transcription model are given in Table S1. The initial chemical reactions rates of the transcription model were based on the TX-TL model by Tuza *et al.* [1].

**Table S1:** Transcription model parameters for PURE cell-free extract

| Parameter | Description | Value | Unit |
|---|---|---|---|
| $k_{\mathrm{NTP_{deg}}}$ | Degradation of nucleotides (ATP, GTP, UTP, and CTP) | $1.75 \times 10^{-4}$ | $\mathrm{hour}^{-1}$ |
| $k_1$ | Binding of RNAP and GTP to the DNA | 0.29 | $\mathrm{nM}^{-2}\mathrm{hour}^{-1}$ |
| $k_2$ | Rate of formation of the RNAP bound GDP and Phosphate complex on the DNA from RNAP bound GTP complex | 0.08 | $\mathrm{hour}^{-1}$ |
| $k_3$ | Unbinding of GDP and Phosphate from the RNAP and DNA complex | 0.42 | $\mathrm{hour}^{-1}$ |
| $k_{\mathrm{start}}$ | Start of the initiation of the mRNA transcript, ($\mathrm{mRNA_0}$) from the RNAP and DNA complex | 0.06 | $\mathrm{hour}^{-1}$ |
| $k_{\mathrm{NTP_{bound}}}$ | Binding rate of NTP to the RNAP bound DNA, complex with initiated mRNA transcript | 0.73 | $\mathrm{nM}^{-1}\mathrm{hour}^{-1}$ |
| $k_{\mathrm{NTP_{add}}}$ | Rate of elongation of the transcript | 35 | $\mathrm{hour}^{-1}$ |
| $k_{\mathrm{NTP_{dis}}}$ | Unbinding rate of NMP and PPi from the open complex | 948.25 | $\mathrm{hour}^{-1}$ |
| $k_{\mathrm{term}}$ | Termination rate | 2.6 | $\mathrm{hour}^{-1}$ |

The model's number of parameters depends on the transcript sequence. For example, the transcription model for Malachite Green aptamer has 276 parameters.

#### 2 Initial Conditions

The initial conditions for the combined transcription and translation model are given in Table S2; based on Version 7 PURE concentration published in Table S1 by Kazuta *et al.* [2]

**Table S2:** Translation initial condition for PURE cell-free reaction model.

| Species | Value | Unit |
|---|---|---|
| ATP | 3750 | µM |
| GTP | 2500 | µM |
| CTP | 1250 | µM |
| UTP | 1250 | µM |
| DNA | 0.005 | µM |
| T7 RNAP | 1 | µM |

| Species | Value | Unit |
|---|---|---|
| mRNA | 0 | µM |
| CK | 10 | µM |
| CP | 10000 | µM |
| EFG | 4.3 | µM |
| EFTs | 13 | µM |
| EFTu | 80 | µM |
| FD | 126.8498943 | µM |
| IF1 | 99 | µM |
| IF2 | 4.1 | µM |
| IF3 | 4.9 | µM |
| MK | 5.6 | µM |
| MTF | 2.4 | µM |
| NDK | 1.8 | µM |
| PPiase | 0.16 | µM |
| RF1 | 0.2 | µM |
| RF2 | 0.2 | µM |
| RF3 | 0.7 | µM |
| RRF | 16 | µM |
| RS70S | 3 | µM |
| AAs | 300 | µM |
| tRNAs | 3.54767184 | µM |

| Species | Value | Unit |
|---|---|---|
| AlaRS | 3 | µM |
| ArgRS | 0.12 | µM |
| AsnRS | 1.7 | µM |
| AspRS | 0.49 | µM |
| CysRS | 0.1 | µM |
| GlnRS | 0.24 | µM |
| GlyRS | 0.35 | µM |
| GluRS | 0.9 | µM |
| HisRS | 0.34 | µM |
| IleRS | 1.5 | µM |
| LeuRS | 0.16 | µM |
| LysRS | 0.46 | µM |
| MetRS | 0.44 | µM |
| PheRS | 0.54 | µM |
| ProRS | 0.67 | µM |
| SerRS | 0.16 | µM |
| ThrRS | 0.34 | µM |
| TrpRS | 0.11 | µM |
| TyrRS | 0.03 | µM |
| ValRS | 0.07 | µM |

### 3 Primers

**Table S3:** Primers used to clone in MGapt, GGGATCCCGACTGGCGAGAGCCAGGTAACGAATGGATC, to pTXTL-T7p14-deGFP DNA originally obtained from myTXTL [3]. The bold text identifies the binding region of the plasmid.

| Name | Seq |
|---|---|
| pT7_MGapt_FOR | GAGCCAGGTAACGAATGGATCCAAT**AATTTTGTTTAACTTTAAGAAGGAGATATACCATG** |
| pT7_MGapt_REV | ATTGGATCCATTCGTTACCTGGCTCTCGCCAGTCGGGATCCC**TCTAGAGGGAAACCGTTG** |

### 4 Additional graphs

#### 4.1 Incorporation of Mean Ribosome Load (MRL)

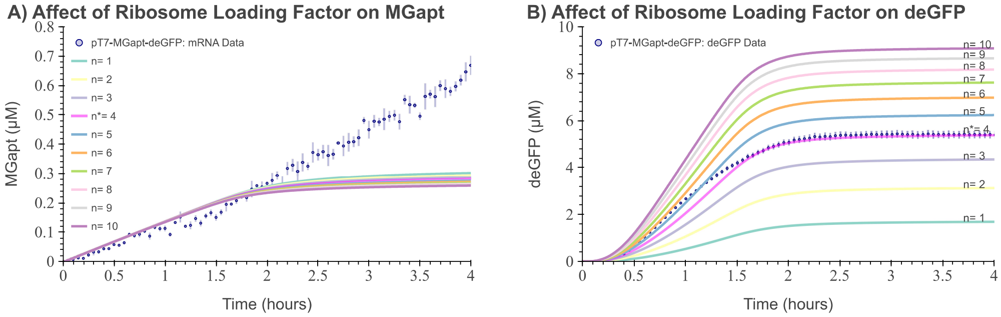


**Figure S1:** Empirical determination of scaling factor, $n$, for the reaction linking the transcription and translation models. The scaling factor allows for the incorporation of ribosome loading in the translation model. A) Simulation results of scaling factor tests, $n = 1$ through 10, on MGapt production(solid lines) overlaying MGapt data (blue circles). B) Simulation results of scaling factor tests, $n = 1$ through 10, of deGFP expression (solid lines) overlaying deGFP data (blue circles).

#### 4.2 Absolute Error of Combined BioCRNpyler Model

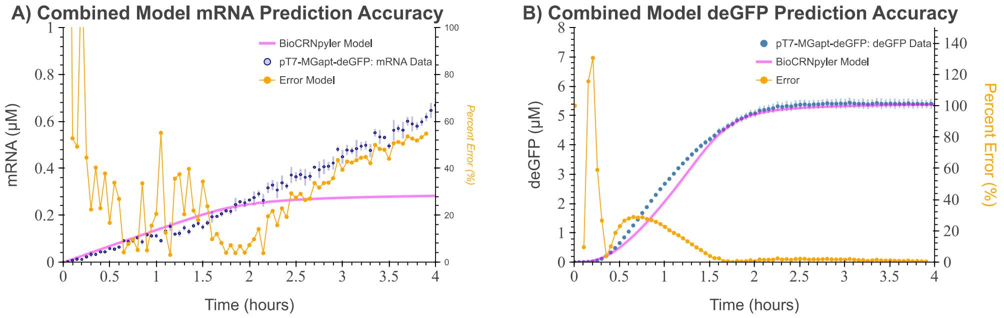


**Figure S2:** Absolute error of combined BioCRNpyler model compared to experimental results. A) Modeled mRNA production in the combined model (magenta line) overlayed with experimental data, three replicates (blue circles and blue error bars), and the absolute error (orange circles) on a secondary axis. B) Modeled deGFP expression in the combined model (magenta line) overlayed with experimental data, three replicates (blue circles and blue error bars), and the absolute error (orange circles) on a secondary axis.

### 5 deGFP protein and MGaptamer mRNA Calibration

The fluorescence calibration curve for MGaptamer was generated using single-stranded RNA purchased from Integrated DNA Technologies (IDT), rArCrUrGrGrArUrCrCrCrGrArCrUrGrGrCrGrArGrArGrCrCrArGrGrUrArArCrGrArArUrGrGrArUrCrCrArArU. The sample arrived lyophilized in tube and weighed 64.5 nmol (0.92 mg). To achieve the concentration of 100 µM 645 µL of nuclease-free water (NFW) was added. The sample was vortexed for several minutes and heated to 55°C for 5 min before vortexing again. The stock concentration was measured by a Nanodrop 2000c before serial dilutions in 1X PBS. Next, respective dilutions were deposited onto the bottom of a Nunc 384 well plate using an Echo 525 Acoustic Liquid Handler. Each well contained a total volume of 10 µL with four technical replicates containing 10 µM of Malachite Green dye. The Nunc 384 well plate was read using a BioTek H1MF plate reader at 37 °C and at 610/650 (Ex/Em) and gain 150. Each point on each calibration curve represents the average of 20 points, four replicates read over 10 minutes at 2.5-minute intervals to generate 5 points per replicate. The points were all background-subtracted from the negative control such that the 0 µM samples had zero fluorescence. Points were fit using linear regression and were not forced to go through the origin. Fits for each calibration curve are indicated in Fig. S3A.

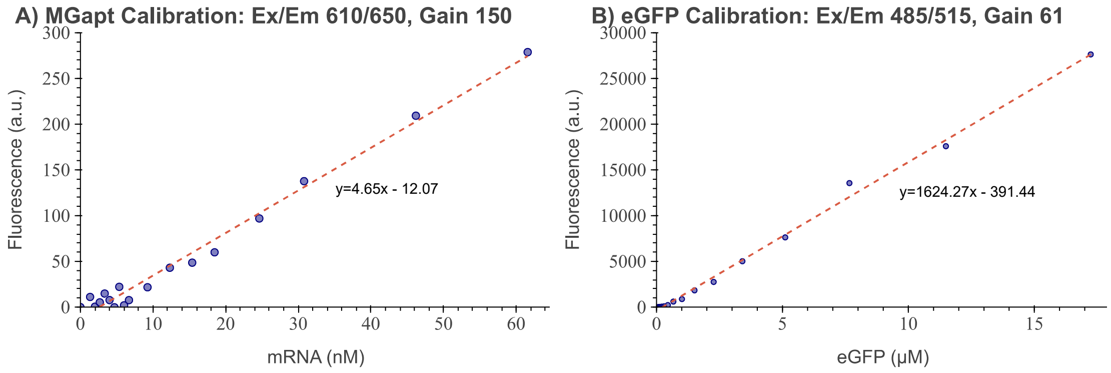


**Figure S3:** The fluorescence calibration curve for deGFP and MGaptamer used to convert RFU to µM.

The fluorescence calibration curve for deGFP was generated using purchased purified eGFP from Cell Biolabs (STA-201). Samples were prepared as described in the myTXTL manual [3]. The 1 mg mL$^{-1}$ eGFP (29.0 kDa was estimated to have a concentration of 34.483 µM). The eGFP stock was diluted in series in 1X PBS and 10 µL of each dilution was pipetted onto the wall of a Nunc 384 well plate, spun down, and then sealed with a plastic film. The plate was allowed to sit for 45 min at room temperature before being read in a BioTek H1MF plate reader at 30 °C and at 485/515 (Ex/Em) and gain of 61. Each point on the calibration curve represents the average of 12 points; three replicates read over 3 minutes at 1-minute intervals to generate 4 points per replicate. The points were all background-subtracted such that the PBS-only samples had zero fluorescence. Points were fit using linear regression and were not forced to go through the origin. Fits for each calibration curve are indicated in the S3B.

#### References


[1] Zoltán A. Tuza, Dan Siegal-Gaskins, Jongmin Kim, and Gábor Szederkényi. Analysis-based parameter estimation of an *in vitro* transcription-translation system. In *Eur. Control Conf. 2015*, page 1554–1560, 2015.

[2] Yasuaki Kazuta, Tomoaki Matsuura, Norikazu Ichihashi, and Tetsuya Yomo. Synthesis of milligram quantities of proteins using a reconstituted in vitro protein synthesis system. *Journal of Bioscience and Bioengineering*, 118(5):554–557, 2014.

[3] Arbor Biosciences. *myTXTL T7 Expression Kit.* Michigan, United States of America, 2019.
