## Supporting scripts, data, and simulation results. for "A chemical reaction network model of PURE": CRN_PURE_Model_Figures.html


### A chemical reaction network model of PURE: Plotting notebook¶

August 8, 2023

[1] Division of Engineering and Applied Science, California Institute of Technology, Pasadena, CA \
[2] Division of Biology and Biological Engineering, California Institute of Technology, Pasadena, CA \

This notebook was used to plot all figures depicted in the *A chemical reaction network model of PURE paper* and supplementary information. All experimental data with plasmid containing the construct pT7-MGapt-UTR1-deGFP-tT7 at 5 μM was collected and compiled into one file. All simulations were run and saved on a previous notebook, then uploaded to be plotted.

### Importing required packages and definitions¶

In [1]:

```
#workhorses
import numpy as np
import pandas as pd
import math
from scipy import stats
import seaborn as sns; sns.set()
import matplotlib.pyplot as plt
from matplotlib.colors import LogNorm, Normalize
from matplotlib.ticker import MaxNLocator
import random 

import bokeh

from bokeh.models import LinearAxis, Range1d
from bokeh.plotting import figure, show
import matplotlib.pyplot as plt

import holoviews as hv
hv.extension('bokeh')
from sklearn.metrics import r2_score
from bokeh.io import export_svgs
from bokeh.models import Title
from bokeh.plotting import gridplot,figure, output_file, show
from bokeh.models.glyphs import Text

#for custom colormaps
from matplotlib.colors import LinearSegmentedColormap
import seaborn as sns; sns.set_theme()

%matplotlib inline
import bokeh.io
import bokeh.plotting
bokeh.io.output_notebook()
from bokeh.themes import Theme

# Modules needed from Bokeh.
from bokeh.io import output_file, show
from bokeh.models import LinearAxis, Range1d

colors = bokeh.palettes.Colorblind[8]
colors2= bokeh.palettes.Set2[8]
colors3= bokeh.palettes.Set3[10]

try:
    import dnaplotlib as dpl
    dpl_enabled = True
except (ModuleNotFoundError,ImportError) as e:
    dpl_enabled = False
    
from bokeh.io import export_png
from bokeh.io import export_svgs
import svglib.svglib as svglib
from bokeh.models import Label

#Get directory
import os
directory = os.getcwd()
```

Loading BokehJS ...

### Getting experimental and simulated data¶

In [2]:

```
# Getting experimental data
folder_exp='/MGapt_deGFP_Exp_PURE_Data'
filename_exp = '/PURE_Mgapt_deGFP_CFP_complied_exp.xlsx'

GFP= pd.read_excel(directory + folder_exp + filename_exp, sheet_name='degfp_tidy', engine='openpyxl')
MGapt= pd.read_excel(directory + folder_exp + filename_exp, sheet_name='mgapt_tidy', engine='openpyxl')

#Normaliz to data
GFP['uM']= (GFP['value']+391.44)/1624.27
MGapt['uM']= ((MGapt['value']+12.07)/4.65)/1000
```

In [3]:

```
# Getting simulation results
filename= '/combined_mGapt-deGFP_5nM.csv'
PURE_Model_Final=pd.read_csv(directory + filename)
```

#### Calculating mean and error bars¶

In [4]:

```
#Mean and error for GFP
all_data= GFP

df0=all_data['dna']=='mgapt-degfp_nM' 
dfn=all_data['dna']=='Neg' 
df2 = all_data[df0 | dfn]

for conc in [5]: #conc=5
    dfconc=df2['mgapt-degfp_nM']== conc
    df = df2[dfconc]

    arr = df['uM'].values.copy()
    arr.resize(3,121)
    DF=pd.DataFrame(arr).T

    DF['uM_ave']=DF.iloc[:, [0,1,2]].mean(axis=1)
    DF['sem']=stats.sem(DF.iloc[:, [0,1,2]].T)
    DF['Time']=df['Time'].reset_index(drop=True)[0:121]

    # Add error bars to the DataFrame
    DF['error_low'] = DF['uM_ave'] - DF['sem']
    DF['error_high'] = DF['uM_ave'] + DF['sem']

    # For convenience
    x = 'Time'
    y = 'uM_ave'

    GFP_Data=DF
```

In [5]:

```
#Mean and error for MGapt
all_data=MGapt

df0=all_data['dna']=='mgapt-degfp_nM' 
dfn=all_data['dna']=='Neg' 
df2 = all_data[df0 | dfn]

for conc in [5]: #conc=5
    dfconc=df2['mgapt-degfp_nM']== conc
    df = df2[dfconc]
    
    arr = df['uM'].values.copy()
    arr.resize(3,121)
    DF=pd.DataFrame(arr).T

    DF['uM_ave']=DF.iloc[:, [0,1,2]].mean(axis=1)
    DF['sem']=stats.sem(DF.iloc[:, [0,1,2]].T)
    DF['Time']=df['Time'].reset_index(drop=True)[0:121]

    # Add error bars to the DataFrame
    DF['error_low'] = DF['uM_ave'] - DF['sem']
    DF['error_high'] = DF['uM_ave'] + DF['sem']
    
    # For convenience
    x = 'Time'
    y = 'uM_ave'
    
    mRNA_Data=DF
```

### Figure 3: Comparing MATLAB vs bioCRNpyler¶

In [6]:

```
#Upload results for peptide fMGG in Matsurra et al. MATLAB model
filename_fmgg = '/fMGG_MATLAB_Matsuura.xlsx'

data_matlab = pd.read_excel(directory + filename_fmgg, sheet_name='Data', engine='openpyxl')
data_matlab.shape

for x in data_matlab.columns:
    count = data_matlab[x].sum()
    if count ==0:
        data_matlab=data_matlab.drop(columns=[x])
```

In [7]:

```
#Upload results for TL only BioCRNpyler model
folder_exTL='/TL_extension_arbAA/'
filename_exTL = 'fMGG_TLonly_BioCRNpyler.csv'

data_crn =pd.read_csv(directory + folder_exTL + filename_exTL,)
data_crn.shape

for x in data_crn.columns:
    count = data_crn[x].sum()
    if count ==0:
        data_crn=data_crn.drop(columns=[x])
```

#### Error for translation of peptide fMGG in BioCRNpyler to MATLAB model¶

In [8]:

```
timepoints = np.linspace(10**-5, 10**4, 500)
```

In [9]:

```
df_Diff=pd.DataFrame(columns=[0,1,2])

for k in range(len(data_matlab['Time'])-1):
    time = data_matlab['Time'][k]

    i=0
    while timepoints[i] <= time >= timepoints[i+1]:
        i += 1

    value=(data_crn['Pept0003'][i+1]-data_crn['Pept0003'][i])/(timepoints[i+1]-timepoints[i])*(time-timepoints[i])+data_crn['Pept0003'][i]
    diff= value-data_matlab['Pept0003'][k]
    error= diff/data_matlab['Pept0003'][k]*100
    er=np.absolute(error)
    
    df_add=pd.DataFrame(np.array([[value, diff, er]]),columns=[0,1,2])
    df_Diff=pd.concat([df_Diff, df_add], ignore_index=True)
```

```
C:\Users\zoila\AppData\Local\Temp\ipykernel_2688\4206078010.py:12: RuntimeWarning: divide by zero encountered in scalar divide
  error= diff/data_matlab['Pept0003'][k]*100
```

In [10]:

```
p1 = bokeh.plotting.figure(toolbar_location='right',
    outline_line_color= None,
    min_border_right=10,
    height=400,
    width=600,)

p1.title.text = "A) Comparison of fMGG translation"
p1.xaxis.axis_label = 'Time (hours)'
p1.yaxis.axis_label = 'Pept fMGG (μM)'
p1.y_range=Range1d(0, 35)
p1.x_range=Range1d(0, 2.8)
p1.outline_line_color=None

# p1.yaxis
p1.ygrid.visible = False
p1.yaxis.axis_label_text_font_size='15pt' 
p1.yaxis.major_label_text_font_size = '15pt'
p1.yaxis.major_label_text_font='Work Sans'
p1.yaxis.axis_label_standoff=15
p1.yaxis.axis_label_text_font_style='normal' 

# p1.xaxis
p1.xgrid.visible = False
p1.xaxis.axis_label_text_font_size='15pt' 
p1.xaxis.major_label_text_font_size = '15pt'
p1.xaxis.major_label_text_font='Work Sans'
p1.xaxis.axis_label_standoff=15
p1.xaxis.axis_label_text_font_style='normal' 

# p1.title
p1.title.text_font_size= '18pt' 
p1.title.align= 'left'
p1.title.offset=-70.0

p1.scatter(data_matlab['Time']/3600,  data_matlab['Pept0003'], legend_label = r"MATLAB Model" "\n" r"[Matsuura et al. 2017]", color = colors[0], fill_alpha=.9)
p1.line(timepoints/3600,  data_crn['Pept0003'], line_width = 3, legend_label = "BioCRNpyler Model", color = "magenta", line_alpha=.5)

p1.extra_y_ranges['foo'] = Range1d(0, 1)
p1.scatter(data_matlab['Time'][390:-1]/3600,  df_Diff[2][390:], legend_label = "Error Model", color = 'orange',y_range_name="foo")

ax2 = LinearAxis(y_range_name="foo", axis_label="Percent Error (%)")
ax2.axis_label_text_color ="orange"
p1.add_layout(ax2, 'right')

# p1.legend
p1.legend.location='top_left'
p1.legend.background_fill_alpha= 0
p1.legend.border_line_color= None
p1.legend.label_text_font_size= '10pt'

bokeh.io.show(p1)
```

#### Adding amino acids to 'fMGG'¶

In [11]:

```
#List of peptides translated to plot
ls_fMGG= ['fMGG','fMGGVSW','fMGGVSWRL',
          'fMGGVSWRLFKI','fMGGVSWRLFKIMCA',
         'fMGGVSWRLFKIMCAYTQNHPDE']
```

In [12]:

```
p2 = bokeh.plotting.figure(toolbar_location='right',
    outline_line_color= None,
    min_border_right=10,
    height=400,
    width=600,)

p2.title.text = "B) Translation of arbitrary peptides"
p2.xaxis.axis_label = 'Time (hours)'
p2.yaxis.axis_label = 'Peptide Concentration (μM)'
p2.outline_line_color=None

# p2.yaxis
p2.ygrid.visible = False
p2.yaxis.axis_label_text_font_size='15pt' 
p2.yaxis.major_label_text_font_size = '15pt'
p2.yaxis.major_label_text_font='Work Sans'
p2.yaxis.axis_label_standoff=15
p2.yaxis.axis_label_text_font_style='normal' 

# p2.xaxis
p2.xgrid.visible = False
p2.xaxis.axis_label_text_font_size='15pt' 
p2.xaxis.major_label_text_font_size = '15pt'
p2.xaxis.major_label_text_font='Work Sans'
p2.xaxis.axis_label_standoff=15
p2.xaxis.axis_label_text_font_style='normal' 

# p2.title
p2.title.text_font_size= '18pt' 
p2.title.align= 'left'
p2.title.offset=-70.0

#colors
colors2=bokeh.palettes.YlGnBu[8]

i=0
for file in ls_fMGG:
    data_mgg = pd.read_csv(directory + folder_exTL + file+'_TLonly_BioCRNpyler.csv')
    peptl=len(file)-1
    if peptl<10:
        p2.line(data_mgg['time']/3600,  data_mgg['Pept000'+str(peptl)], line_width = 3, line_dash='dashed', legend_label = file, color = colors2[i])
        i+=1
    else:
        p2.line(data_mgg['time']/3600,  data_mgg['Pept00'+str(peptl)], line_width = 3, line_dash='dashed', legend_label = file, color = colors2[i])
        i+=1
        
# p2.legend
p2.legend.location='bottom_right'
p2.legend.background_fill_alpha= 0
p2.legend.border_line_color= None
p2.legend.label_text_font_size= '10pt'

bokeh.io.show(p2)
```

#### Moving to deGFP, with mRNA set at 0.126 uM¶

##### Plot results along with experiemntal data¶

In [13]:

```
#Experimental data set of one repeat
filename='/2022.08.18_PURE_deGFP.csv'
deGFP_Data = pd.read_csv(directory + folder_exp+filename)
deGFP_Data['deGFP_uM']=(deGFP_Data['deGFP']+391.44)/1624.27-0.24
```

In [14]:

```
#TL model of MGapt-deGFP construct
filename='/CRN_PURE_TLonly_MGaptdeGFP_results.csv'
MGapt_deGFP_TLonly = pd.read_csv(directory + filename)
```

In [15]:

```
p3 = bokeh.plotting.figure(toolbar_location='right',
    outline_line_color= None,
    min_border_right=10,
    height=400,
    width=600,)

p3.title.text = "C) Translation simulation of deGFP"
p3.xaxis.axis_label = 'Time (hours)'
p3.yaxis.axis_label = 'deGFP (μM)'
p3.outline_line_color=None

# p3.yaxis
p3.ygrid.visible = False
p3.yaxis.axis_label_text_font_size='15pt' 
p3.yaxis.major_label_text_font_size = '15pt'
p3.yaxis.major_label_text_font='Work Sans'
p3.yaxis.axis_label_standoff=15
p3.yaxis.axis_label_text_font_style='normal' 

# p3.xaxis
p3.xgrid.visible = False
p3.xaxis.axis_label_text_font_size='15pt' 
p3.xaxis.major_label_text_font_size = '15pt'
p3.xaxis.major_label_text_font='Work Sans'
p3.xaxis.axis_label_standoff=15
p3.xaxis.axis_label_text_font_style='normal' 

# p3.title
p3.title.text_font_size= '18pt' 
p3.title.align= 'left'
p3.title.offset=-65.0

p3.line(MGapt_deGFP_TLonly['time']/3600, MGapt_deGFP_TLonly['Pept0225'],legend_label = "BioCRNpyler Model", line_color="magenta", line_width=3) 
#Experimental data set 1
p3.scatter(deGFP_Data['timepoints']/3600, deGFP_Data['deGFP_uM'], legend_label = "pT7-MGapt-deGFP.R1", radius=0.01, fill_alpha=0.2, color="navy") 
p3.legend.location='bottom_right'
p3.x_range=Range1d(0, 2.5)
p3.y_range=Range1d(0, 6)

# p2.legend
p3.legend.location='bottom_right'
p3.legend.background_fill_alpha= 0
p3.legend.border_line_color= None
p3.legend.label_text_font_size= '10pt'

bokeh.io.show(p3)
```

#### Adding empty space text¶

In [16]:

```
p4 = bokeh.plotting.figure(toolbar_location='right',
    outline_line_color= None,
    min_border_right=10,
    height=400,
    width=600,)

p4.y_range=Range1d(-5, 10)
p4.x_range=Range1d(-3, 3)
p4.title.text = "Amino acids list of deGFP:"
# p4.title.offset=-10.0
p4.outline_line_color=None

#Remove ticks
p4.xaxis.major_tick_line_color = None  # turn off x-axis major ticks
p4.xaxis.minor_tick_line_color = None  # turn off x-axis minor ticks

p4.yaxis.major_tick_line_color = None  # turn off y-axis major ticks
p4.yaxis.minor_tick_line_color = None  # turn off y-axis minor ticks

p4.yaxis.major_label_text_font_size = '0pt'
p4.xaxis.major_label_text_font_size = '0pt'

p4.ygrid.visible = False 
p4.xgrid.visible = False

p4.xaxis.axis_line_color=None
p4.yaxis.axis_line_color=None

# p4.title
p4.title.text_font_size= '18pt' 
p4.title.align= 'left'

#Amino acid list
aa_gfp='MELFTGVVPILVELDGDVNGHKFSVSGEGEGDATYGKLTLKFICTTGKLPVPWPTLVTTLTYGVQCFSRYPDHMKQHDFFKSAMPEGYVQERTIFFKDDGNYKTRAEVKFEGDTLVNRIELKGIDFKEDGNILGHKLEYNYNSHNVYIMADKQKNGIKVNFKIRHNIEDGSVQLADHYQQNTPIGDGPVLLPDNHYLSTQSALSKDPNEKRDHMVLLEFVTAAGI'

for x in range(9):
    start=x*25
    end=(x+1)*25-1
    mytext0 = Label(x=-1.5, y=9-x*1.5, text=aa_gfp[start:end],
                   text_font_size='15pt',
                  level='glyph',
                  x_offset=-15,
                  y_offset=-13.5,)
    p4.add_layout(mytext0)

bokeh.io.show(p4)
```

#### Plot as grid and save¶

In [17]:

```
# using grid layout on f1,f2,f3 plots
grid1=gridplot([[p1,p2,],[p3,p4]], width=600, height=400)
show(grid1)
```

#### Plot as Grid and save as .svg¶

In [18]:

```
count=1
for pic in [p1,p2,p3,p4]:
    pic.output_backend = "svg"
    export_svgs(pic, filename = 'Figures_PURE_Model/Fig3_TL_Expansion_model' + str(count) + '.svg',width=800, height=600)
    count+=1
    
export_svgs(grid1, filename = 'Figures_PURE_Model/Fig3_TL_Expansion_model.svg', width=800, height=600)
```

```
C:\Users\zoila\anaconda3\envs\python38\lib\site-packages\bokeh\io\export.py:308: UserWarning: Export method called with width or height argument on a non-Plot model. The size values will be ignored.
  warnings.warn("Export method called with width or height argument on a non-Plot model. The size values will be ignored.")
```

Out[18]:

```
['Figures_PURE_Model/Fig3_TL_Expansion_model.svg',
 'Figures_PURE_Model/Fig3_TL_Expansion_model_1.svg',
 'Figures_PURE_Model/Fig3_TL_Expansion_model_2.svg',
 'Figures_PURE_Model/Fig3_TL_Expansion_model_3.svg']
```

### Figure 5: Molecules: Plot consumables¶

In [19]:

```
Time=np.round(PURE_Model_Final['time']/3600,decimals=1)
```

In [20]:

```
list_ntps=['ATP','ADP','AMP',
       'CTP','CDP',
       'GTP','GDP',
       'UTP','UDP',]

list_aa=['Ala','Arg','Asn','Asp','Cys', 
          'Gln','Glu','Gly','His','Ile',
          'Leu','Lys','Met','Phe','Pro',
          'Ser','Thr','Trp','Tyr','Val',]

list_etc=['CP', 'PO4','PPi',]
```

#### Plots NXPs¶

In [21]:

```
data_ntps=PURE_Model_Final[list_ntps]
data_ntps['Time']=Time
data_ntps=data_ntps.set_index('Time')
```

```
C:\Users\zoila\AppData\Local\Temp\ipykernel_2688\1162131899.py:2: SettingWithCopyWarning: 
A value is trying to be set on a copy of a slice from a DataFrame.
Try using .loc[row_indexer,col_indexer] = value instead

See the caveats in the documentation: https://pandas.pydata.org/pandas-docs/stable/user_guide/indexing.html#returning-a-view-versus-a-copy
  data_ntps['Time']=Time
```

In [22]:

```
fig1, ax1 = plt.subplots(figsize=(10,5)) 
ax1 = sns.heatmap(data_ntps.transpose(),fmt=".1f",norm=LogNorm(),
                 cbar_kws={'label': 'Concentration (μM)'})

ax1.set_xlabel("Time (hours)", fontsize=15)
ax1.set_ylabel('Species', fontsize=15)

plt.title("A) NXPs Concentration in Combine PURE Model", loc='left',fontsize=18)
plt.show()
```

In [23]:

```
image_format = 'svg' # e.g .png, .svg, etc.
image_name = 'Figures_PURE_Model/Fig5_nxps.svg'

fig1.savefig(image_name, format=image_format, dpi=1200)
```

#### Plots AAs¶

In [24]:

```
data_aa=PURE_Model_Final[list_aa]
data_aa['Time']=Time
data_aa=data_aa.set_index('Time')
```

```
C:\Users\zoila\AppData\Local\Temp\ipykernel_2688\4075196112.py:2: SettingWithCopyWarning: 
A value is trying to be set on a copy of a slice from a DataFrame.
Try using .loc[row_indexer,col_indexer] = value instead

See the caveats in the documentation: https://pandas.pydata.org/pandas-docs/stable/user_guide/indexing.html#returning-a-view-versus-a-copy
  data_aa['Time']=Time
```

In [25]:

```
fig2, ax2 = plt.subplots(figsize=(10,5)) 
ax2 = sns.heatmap(data_aa.transpose(),fmt=".1f",
                 cbar_kws={'label': 'Concentration (μM)'})

ax2.set_xlabel("Time (hours)", fontsize=15)
ax2.set_ylabel('Species', fontsize=15)

plt.title("B) Amino Acids Concentration in Combine PURE Model", loc='left',fontsize=18)
plt.show()
```

In [26]:

```
image_format = 'svg' # e.g .png, .svg, etc.
image_name = 'Figures_PURE_Model/Fig5_aa.svg'

fig2.savefig(image_name, format=image_format, dpi=1200)
```

### Figure of Mean Ribosome Load (MRL) Simulations¶

Effects of multiplier (n) on deGFP expression

In [27]:

```
#Upload and compile mean ribosome load simulations
gfp_all=pd.DataFrame()
mrna_all=pd.DataFrame()
filename_mrl = '/MGapt-deGFP_Model_MRL_runs/mGapt-deGFP_5nM_'

for x in ['n1','n2','n3','n4','n5','n6','n7','n8','n9','n10']:
    full_data=pd.read_csv(directory + filename_mrl+x+'.csv')
    gfp_all[x]=full_data['deGFP_m']
    mrna_all[x]=full_data['mRNA_t']
```

#### deGFP¶

In [28]:

```
pRBSb = bokeh.plotting.figure(toolbar_location='right',
    outline_line_color= None,
    min_border_right=10,
    height=400,
    width=600,)

pRBSb.title.text = "B) Affect of Ribosome Loading Factor on deGFP"
pRBSb.xaxis.axis_label = 'Time (hours)'
pRBSb.yaxis.axis_label = 'deGFP (μM)'
pRBSb.y_range=Range1d(0, 10)
pRBSb.x_range=Range1d(0, 4)
pRBSb.outline_line_color=None

# pRBSbyaxis
pRBSb.ygrid.visible = False
pRBSb.yaxis.axis_label_text_font_size='15pt' 
pRBSb.yaxis.major_label_text_font_size = '15pt'
pRBSb.yaxis.major_label_text_font='Work Sans'
pRBSb.yaxis.axis_label_standoff=15
pRBSb.yaxis.axis_label_text_font_style='normal' 

# pRBSbxaxis
pRBSb.xgrid.visible = False
pRBSb.xaxis.axis_label_text_font_size='15pt' 
pRBSb.xaxis.major_label_text_font_size = '15pt'
pRBSb.xaxis.major_label_text_font='Work Sans'
pRBSb.xaxis.axis_label_standoff=15
pRBSb.xaxis.axis_label_text_font_style='normal' 

# pRBSbtitle
pRBSb.title.text_font_size= '18pt' 
pRBSb.title.align= 'left'
pRBSb.title.offset=-70.0


#Data from experiments
pRBSb.circle(
    # source=DF,
    x=GFP_Data['Time'], y=GFP_Data['uM_ave']-GFP_Data['uM_ave'][0],
    legend_label= ("pT7-MGapt-deGFP: deGFP Data"),color= 'navy',radius=0.01, fill_alpha=0.2,)

# Add error bars
pRBSb.segment(
    # source=DF,
    x0=GFP_Data['Time'], y0=GFP_Data['error_low']-GFP_Data['uM_ave'][0], 
    x1=GFP_Data['Time'], y1=GFP_Data['error_high']-GFP_Data['uM_ave'][0], 
    line_width=2, color= 'navy', line_alpha=.25)

#plot over all n span
i=0
for x in gfp_all.columns:
    y=gfp_all[x]
    if x!='n4':
        pRBSb.line(PURE_Model_Final['time']/3600,  y, line_width = 3,  color = colors3[i])
        n_label = Label(x=3.7, y=y.max()-.1, text='n= '+str(i+1),text_baseline = 'bottom',text_font_size = '14px')
        pRBSb.add_layout(n_label)
        i=i+1
    else:
        pRBSb.line(PURE_Model_Final['time']/3600,  PURE_Model_Final['deGFP_m'], line_width = 3, color = 'magenta', line_alpha=.5)
        n_label = Label(x=3.7, y=PURE_Model_Final['deGFP_m'].max(), text='n*= '+str(i+1),text_baseline = 'bottom',text_font_size = '14px')
        pRBSb.add_layout(n_label)
        i=i+1

# pRBSb.legend
pRBSb.legend.location='top_left'
pRBSb.legend.background_fill_alpha= 0
pRBSb.legend.border_line_color= None
pRBSb.legend.label_text_font_size= '10pt'

bokeh.io.show(pRBSb)
```

#### mRNA¶

In [29]:

```
pRBSa = bokeh.plotting.figure(toolbar_location='right',
    outline_line_color= None,
    min_border_right=10,
    height=400,
    width=600,)

pRBSa.title.text = "A) Affect of Ribosome Loading Factor on MGapt"
pRBSa.xaxis.axis_label = 'Time (hours)'
pRBSa.yaxis.axis_label = 'MGapt (μM)'
pRBSa.y_range=Range1d(0, 0.8)
pRBSa.x_range=Range1d(0, 4)
pRBSa.outline_line_color=None

# pRBSa.yaxis
pRBSa.ygrid.visible = False
pRBSa.yaxis.axis_label_text_font_size='15pt' 
pRBSa.yaxis.major_label_text_font_size = '15pt'
pRBSa.yaxis.major_label_text_font='Work Sans'
pRBSa.yaxis.axis_label_standoff=15
pRBSa.yaxis.axis_label_text_font_style='normal' 

# pRBSa.xaxis
pRBSa.xgrid.visible = False
pRBSa.xaxis.axis_label_text_font_size='15pt' 
pRBSa.xaxis.major_label_text_font_size = '15pt'
pRBSa.xaxis.major_label_text_font='Work Sans'
pRBSa.xaxis.axis_label_standoff=15
pRBSa.xaxis.axis_label_text_font_style='normal' 

# pRBSa.title
pRBSa.title.text_font_size= '18pt' 
pRBSa.title.align= 'left'
pRBSa.title.offset=-70.0

#Data from experiments
pRBSa.circle(
    x=mRNA_Data['Time'], y=mRNA_Data['uM_ave']-mRNA_Data['uM_ave'][0],  legend_label ="pT7-MGapt-deGFP: mRNA Data",color= "navy", radius=0.01, fill_alpha=0.2,)

# Add error bars
pRBSa.segment(
    x0=mRNA_Data['Time'], y0=mRNA_Data['error_low']-mRNA_Data['uM_ave'][0], 
    x1=mRNA_Data['Time'], y1=mRNA_Data['error_high']-mRNA_Data['uM_ave'][0],
    line_width=2, color= "navy", line_alpha=.25)

#plot over all n span
i=0
for x in mrna_all.columns:
    y=mrna_all[x]
    if x!='n4':
        pRBSa.line(PURE_Model_Final['time']/3600,  y, line_width = 3, legend_label = 'n= '+str(i+1), color = colors3[i])
        i=i+1
    else:
        pRBSa.line(PURE_Model_Final['time']/3600,  PURE_Model_Final['mRNA_t'], line_width = 3, legend_label = 'n*= 4', color = 'magenta', line_alpha=.5)
        i=i+1

# pRBSa.legend
pRBSa.legend.location='top_left'
pRBSa.legend.background_fill_alpha= 0
pRBSa.legend.border_line_color= None
pRBSa.legend.label_text_font_size= '10pt'

bokeh.io.show(pRBSa)
```

#### Plot as Grid and save as .svg¶

In [30]:

```
# using grid layout on plots
pRBS=gridplot([[pRBSa, pRBSb]], width=600, height=400)
show(pRBS)
```

In [31]:

```
count=1
for pic in [pRBSa, pRBSb]:
    pic.output_backend = "svg"
    export_svgs(pic, filename = 'Figures_PURE_Model/Fig_pRBS' + str(count) + '.svg',width=800, height=600)
    count+=1
    
export_svgs(grid1, filename = 'Figures_PURE_Model/Fig_pRBS.svg', width=800, height=600)
```

```
C:\Users\zoila\anaconda3\envs\python38\lib\site-packages\bokeh\io\export.py:308: UserWarning: Export method called with width or height argument on a non-Plot model. The size values will be ignored.
  warnings.warn("Export method called with width or height argument on a non-Plot model. The size values will be ignored.")
```

Out[31]:

```
['Figures_PURE_Model/Fig_pRBS.svg',
 'Figures_PURE_Model/Fig_pRBS_1.svg',
 'Figures_PURE_Model/Fig_pRBS_2.svg',
 'Figures_PURE_Model/Fig_pRBS_3.svg']
```

### Figure\_Absolute Error: Plot data vs model error¶

#### deGFP¶

##### Error calculations¶

In [32]:

```
#Trimming experimental data for first 4 hours to be consistent with simulation
DF_data_degfp=GFP_Data[:80]
timepoints = np.linspace(10**-5, 4*60*60, 500)
DF_data_degfp['uM']=DF_data_degfp['uM_ave']-DF_data_degfp['uM_ave'].min()
timepoints=timepoints/3600
```

```
C:\Users\zoila\AppData\Local\Temp\ipykernel_2688\4253514129.py:4: SettingWithCopyWarning: 
A value is trying to be set on a copy of a slice from a DataFrame.
Try using .loc[row_indexer,col_indexer] = value instead

See the caveats in the documentation: https://pandas.pydata.org/pandas-docs/stable/user_guide/indexing.html#returning-a-view-versus-a-copy
  DF_data_degfp['uM']=DF_data_degfp['uM_ave']-DF_data_degfp['uM_ave'].min()
```

In [33]:

```
#Calculate absolute error between simulation and experimental data
df_Diff_degfp = pd.DataFrame()
df_Diff_degfp

for k in range(len(DF_data_degfp['Time'])-1):
    time = DF_data_degfp['Time'][k]
    i=0
    while timepoints[i] <= time >= timepoints[i+1]:
        i += 1

    value=(PURE_Model_Final['deGFP_m'][i+1]-PURE_Model_Final['deGFP_m'][i])/(timepoints[i+1]-timepoints[i])*(time-timepoints[i])+PURE_Model_Final['deGFP_m'][i]
    diff= value-DF_data_degfp['uM'][k]
    error= diff/DF_data_degfp['uM'][k]*100
    er=np.absolute(error)
    
    df_Diff_degfp[k]=[value,diff,er]
    
df_Diff_degfp=df_Diff_degfp.transpose()
```

```
C:\Users\zoila\AppData\Local\Temp\ipykernel_2688\3971434117.py:13: RuntimeWarning: divide by zero encountered in scalar divide
  error= diff/DF_data_degfp['uM'][k]*100
```

##### Plot¶

In [34]:

```
pErrorb = bokeh.plotting.figure(toolbar_location='right',
    outline_line_color= None,
    min_border_right=10,
    height=400,
    width=600,)

pErrorb.title.text = "B) Combined Model deGFP Prediction Accuracy"
pErrorb.xaxis.axis_label = 'Time (hours)'
pErrorb.yaxis.axis_label = 'deGFP (μM)'
pErrorb.y_range=Range1d(0, 8)
pErrorb.x_range=Range1d(0, 4)
pErrorb.outline_line_color=None

# pErrorb.yaxis
pErrorb.ygrid.visible = False
pErrorb.yaxis.axis_label_text_font_size='15pt' 
pErrorb.yaxis.major_label_text_font_size = '15pt'
pErrorb.yaxis.major_label_text_font='Work Sans'
pErrorb.yaxis.axis_label_standoff=15
pErrorb.yaxis.axis_label_text_font_style='normal' 

# pErrorb.xaxis
pErrorb.xgrid.visible = False
pErrorb.xaxis.axis_label_text_font_size='15pt' 
pErrorb.xaxis.major_label_text_font_size = '15pt'
pErrorb.xaxis.major_label_text_font='Work Sans'
pErrorb.xaxis.axis_label_standoff=15
pErrorb.xaxis.axis_label_text_font_style='normal' 

# pErrorb.title
pErrorb.title.text_font_size= '18pt' 
pErrorb.title.align= 'left'

pErrorb.title.offset=-65.0

pErrorb.circle(
    # source=DF,
    x=GFP_Data['Time'],
    y=GFP_Data['uM_ave']-GFP_Data['uM_ave'][0],
    legend_label= ("pT7-MGapt-deGFP: deGFP Data"), 
    color= 'navy',radius=0.01, fill_alpha=0.2,)

# Add error bars
pErrorb.segment(
    # source=DF,
    x0=GFP_Data['Time'],
    y0=GFP_Data['error_low']-GFP_Data['uM_ave'][0],
    x1=GFP_Data['Time'],
    y1=GFP_Data['error_high']-GFP_Data['uM_ave'][0],
    line_width=2,
    color= 'navy',
    line_alpha=.25
)

    
df_Diff = pd.DataFrame()
df_Diff
DF_data=GFP_Data[:80]
DF_data['uM']=DF['uM_ave']-DF['uM_ave'][0]
timepoints = np.linspace(10**-5, 4*60*60, 500)
timepoints=timepoints/3600
# DF_data
for k in range(len(DF_data['Time'])-1):
    time = DF_data['Time'][k]
    i=0
    while timepoints[i] <= time >= timepoints[i+1]:
        i += 1
    value=(PURE_Model_Final['deGFP_m'][i+1]-PURE_Model_Final['deGFP_m'][i])/(timepoints[i+1]-timepoints[i])*(time-timepoints[i])+PURE_Model_Final['deGFP_m'][i]
    diff= value-DF_data['uM'][k]
    error= diff/DF_data['uM'][k]*100
    er=np.absolute(error)
    
    df_Diff[k]=[value,diff,er]
    
df_Diff=df_Diff.transpose()

# Model data
pErrorb.line(PURE_Model_Final['time']/3600, PURE_Model_Final['deGFP_m'],legend_label = "BioCRNpyler Model", line_color='magenta', 
        line_dash= "solid", line_width=3,line_alpha=.5) 

pErrorb.extra_y_ranges['foo'] = Range1d(0, 150)
pErrorb.scatter(DF_data['Time'][:79],  df_Diff_degfp[2], legend_label = "Error", color = 'orange',y_range_name="foo")
pErrorb.line(DF_data['Time'][:79],  df_Diff_degfp[2], legend_label = "Error", color = 'orange',y_range_name="foo")

ax2 = LinearAxis(y_range_name="foo", axis_label="Percent Error (%)")
ax2.axis_label_text_color ="orange"
pErrorb.add_layout(ax2, 'right')

# p1.legend
pErrorb.legend.location='top_right'
pErrorb.legend.background_fill_alpha= 0
pErrorb.legend.border_line_color= None
pErrorb.legend.label_text_font_size= '10pt'

bokeh.io.show(pErrorb)
```

```
C:\Users\zoila\AppData\Local\Temp\ipykernel_2688\1763936343.py:59: SettingWithCopyWarning: 
A value is trying to be set on a copy of a slice from a DataFrame.
Try using .loc[row_indexer,col_indexer] = value instead

See the caveats in the documentation: https://pandas.pydata.org/pandas-docs/stable/user_guide/indexing.html#returning-a-view-versus-a-copy
  DF_data['uM']=DF['uM_ave']-DF['uM_ave'][0]
C:\Users\zoila\AppData\Local\Temp\ipykernel_2688\1763936343.py:70: RuntimeWarning: divide by zero encountered in scalar divide
  error= diff/DF_data['uM'][k]*100
```

#### mRNA¶

##### Error calculations¶

In [35]:

```
DF_data_mRNA=mRNA_Data[:80]
timepoints = np.linspace(10**-5, 4*60*60, 500)
DF_data_mRNA['uM']=DF_data_mRNA['uM_ave']-DF_data_mRNA['uM_ave'].min()
timepoints=timepoints/3600
```

```
C:\Users\zoila\AppData\Local\Temp\ipykernel_2688\3879975904.py:3: SettingWithCopyWarning: 
A value is trying to be set on a copy of a slice from a DataFrame.
Try using .loc[row_indexer,col_indexer] = value instead

See the caveats in the documentation: https://pandas.pydata.org/pandas-docs/stable/user_guide/indexing.html#returning-a-view-versus-a-copy
  DF_data_mRNA['uM']=DF_data_mRNA['uM_ave']-DF_data_mRNA['uM_ave'].min()
```

In [36]:

```
df_Diff_mrna = pd.DataFrame()
df_Diff_mrna

for k in range(len(DF_data_mRNA['Time'])-1):
    time = DF_data_mRNA['Time'][k]
    i=0
    while timepoints[i] <= time >= timepoints[i+1]:
        i += 1
    value=(PURE_Model_Final['mRNA_t'][i+1]-PURE_Model_Final['mRNA_t'][i])/(timepoints[i+1]-timepoints[i])*(time-timepoints[i])+PURE_Model_Final['mRNA_t'][i]
    diff= value-DF_data_mRNA['uM'][k]
    error= diff/DF_data_mRNA['uM'][k]*100
    er=np.absolute(error)
    
    df_Diff_mrna[k]=[value,diff,er]
    
df_Diff_mrna=df_Diff_mrna.transpose()
```

```
C:\Users\zoila\AppData\Local\Temp\ipykernel_2688\1624804730.py:11: RuntimeWarning: divide by zero encountered in scalar divide
  error= diff/DF_data_mRNA['uM'][k]*100
```

##### Plot¶

In [37]:

```
pErrora = bokeh.plotting.figure(toolbar_location='right',
    outline_line_color= None,
    min_border_right=10,
    height=400,
    width=600,)


pErrora.title.text = "A) Combined Model mRNA Prediction Accuracy"
pErrora.xaxis.axis_label = 'Time (hours)'
pErrora.yaxis.axis_label = 'mRNA (μM)'
pErrora.outline_line_color=None

# pErrora.yaxis
pErrora.ygrid.visible = False
pErrora.yaxis.axis_label_text_font_size='15pt' 
pErrora.yaxis.major_label_text_font_size = '15pt'
pErrora.yaxis.major_label_text_font='Work Sans'
pErrora.yaxis.axis_label_standoff=15
pErrora.yaxis.axis_label_text_font_style='normal' 

# pErrora.xaxis
pErrora.xgrid.visible = False
pErrora.xaxis.axis_label_text_font_size='15pt' 
pErrora.xaxis.major_label_text_font_size = '15pt'
pErrora.xaxis.major_label_text_font='Work Sans'
pErrora.xaxis.axis_label_standoff=15
pErrora.xaxis.axis_label_text_font_style='normal' 

# pErrora.title
pErrora.title.text_font_size= '18pt' 
pErrora.title.align= 'left'
pErrora.title.offset=-65.0

mult=.0645
# Model results
pErrora.line(PURE_Model_Final['time']/3600, PURE_Model_Final['mRNA_t'],legend_label = "BioCRNpyler Model", line_color="magenta", line_width=3, line_alpha=.5) #mRNA

# Experimental data set 1
pErrora.circle(
    x=mRNA_Data['Time'], y=mRNA_Data['uM_ave']-mRNA_Data['uM_ave'][0], legend_label = "pT7-MGapt-deGFP: mRNA Data", color= "navy", radius=0.01, fill_alpha=0.2,)

# Add error bars
pErrora.segment(
    x0=mRNA_Data['Time'], y0=mRNA_Data['error_low']-mRNA_Data['uM_ave'][0], 
    x1=mRNA_Data['Time'], y1=mRNA_Data['error_high']-mRNA_Data['uM_ave'][0],
    line_width=2, color= "navy", line_alpha=.25)

# Add error between measured and simulated
pErrora.extra_y_ranges['foo'] = Range1d(0, 100)
pErrora.line(DF_data['Time'][:79],  df_Diff_mrna[2], legend_label = "Error Model", color = 'orange',y_range_name="foo")
pErrora.scatter(DF_data['Time'][:79],  df_Diff_mrna[2], legend_label = "Error Model", color = 'orange',y_range_name="foo")

ax2 = LinearAxis(y_range_name="foo", axis_label="Percent Error (%)")
ax2.axis_label_text_color ="orange"
pErrora.add_layout(ax2, 'right')

pErrora.x_range=Range1d(0, 4)
pErrora.y_range=Range1d(0, 1)

# pErrora.legend
pErrora.legend.location='top_right'
pErrora.legend.background_fill_alpha= 0
pErrora.legend.border_line_color= None
pErrora.legend.label_text_font_size= '10pt'

bokeh.io.show(pErrora)
```

#### Plot as grid and save¶

In [38]:

```
# using grid layout on plots
pError=gridplot([[pErrora, pErrorb]], width=600, height=400)
show(pError)
```

In [39]:

```
count=1
for pic in [pErrora,pErrorb]:
    pic.output_backend = "svg"
    export_svgs(pic, filename = 'Figures_PURE_Model/Fig_pError' + str(count) + '.svg',width=800, height=600)
    count+=1
    
export_svgs(grid1, filename = 'Figures_PURE_Model/Fig_pError.svg', width=800, height=600)
```

```
C:\Users\zoila\anaconda3\envs\python38\lib\site-packages\bokeh\io\export.py:308: UserWarning: Export method called with width or height argument on a non-Plot model. The size values will be ignored.
  warnings.warn("Export method called with width or height argument on a non-Plot model. The size values will be ignored.")
```

Out[39]:

```
['Figures_PURE_Model/Fig_pError.svg',
 'Figures_PURE_Model/Fig_pError_1.svg',
 'Figures_PURE_Model/Fig_pError_2.svg',
 'Figures_PURE_Model/Fig_pError_3.svg']
```

### Calibrations Plots¶

#### mRNA Calibrations¶

In [40]:

```
#MGapt Calibration file
filename = '/Calibrations_Biotek/2022.10.04_mRNA_mGapt_data_37C.csv'
mGapt_Data =  pd.read_csv(directory+filename, delimiter = '\,', names = ['mRNA (nM)','Biotek 1', 'Biotek 2','Biotek 3', 'Biotek 4'], skiprows = 1)
```

```
C:\Users\zoila\AppData\Local\Temp\ipykernel_2688\1035132161.py:3: ParserWarning: Falling back to the 'python' engine because the 'c' engine does not support regex separators (separators > 1 char and different from '\s+' are interpreted as regex); you can avoid this warning by specifying engine='python'.
  mGapt_Data =  pd.read_csv(directory+filename, delimiter = '\,', names = ['mRNA (nM)','Biotek 1', 'Biotek 2','Biotek 3', 'Biotek 4'], skiprows = 1)
```

In [41]:

```
pCal_mRNA = bokeh.plotting.figure(toolbar_location='right',
    outline_line_color= None,
    min_border_right=10,
    height=400,
    width=600,)

pCal_mRNA.title.text = "A) MGapt Calibration: Ex/Em 610/650, Gain 150 "
pCal_mRNA.xaxis.axis_label = 'mRNA (nM)'
pCal_mRNA.yaxis.axis_label = 'Fluorescence (a.u.)'
pCal_mRNA.y_range=Range1d(0, 300)
pCal_mRNA.x_range=Range1d(0, 65)
pCal_mRNA.outline_line_color=None

# pCal_mRNAyaxis
pCal_mRNA.ygrid.visible = False
pCal_mRNA.yaxis.axis_label_text_font_size='15pt' 
pCal_mRNA.yaxis.major_label_text_font_size = '15pt'
pCal_mRNA.yaxis.major_label_text_font='Work Sans'
pCal_mRNA.yaxis.axis_label_standoff=15
pCal_mRNA.yaxis.axis_label_text_font_style='normal' 

# pCal_mRNAxaxis
pCal_mRNA.xgrid.visible = False
pCal_mRNA.xaxis.axis_label_text_font_size='15pt' 
pCal_mRNA.xaxis.major_label_text_font_size = '15pt'
pCal_mRNA.xaxis.major_label_text_font='Work Sans'
pCal_mRNA.xaxis.axis_label_standoff=15
pCal_mRNA.xaxis.axis_label_text_font_style='normal' 

# pCal_mRNAtitle
pCal_mRNA.title.text_font_size= '18pt' 
pCal_mRNA.title.align= 'left'
pCal_mRNA.title.offset=-70.0


#Data from experiments
pCal_mRNA.scatter(mGapt_Data['mRNA (nM)'], mGapt_Data['Biotek 1'], radius=0.5, fill_alpha=0.5, color="navy") #mRNA
m4, b4 = np.polyfit(mGapt_Data['mRNA (nM)'], mGapt_Data['Biotek 1'], 1)
pCal_mRNA.line(mGapt_Data['mRNA (nM)'],mGapt_Data['mRNA (nM)']*m4+b4 ,  color="#d95b43", line_dash= 'dashed', line_width=2) #trendline

citation4 = Label(x=200, y=100, x_units='screen', y_units='screen',
                 text='y= '+str(np.around(m4, decimals=2, out=None))+'x + ' + str(np.around(b4, decimals=2, out=None)),# render_mode='css',
                 border_line_color=None, border_line_alpha=1.0,
                 background_fill_color='white', background_fill_alpha=1.0, text_font_size='14pt')
pCal_mRNA.add_layout(citation4)

bokeh.io.show(pCal_mRNA)
```

#### deGFP Calibrations¶

In [42]:

```
#eGFP Calibration file
filename = '/Calibrations_Biotek/2022.10.04_eGFP_data_30C.csv'
eGFP_Data =  pd.read_csv(directory+filename, delimiter = '\,', names = ['eGFP (uM)','Biotek 1','Biotek 4',], skiprows = 1)
```

```
C:\Users\zoila\AppData\Local\Temp\ipykernel_2688\2386572246.py:3: ParserWarning: Falling back to the 'python' engine because the 'c' engine does not support regex separators (separators > 1 char and different from '\s+' are interpreted as regex); you can avoid this warning by specifying engine='python'.
  eGFP_Data =  pd.read_csv(directory+filename, delimiter = '\,', names = ['eGFP (uM)','Biotek 1','Biotek 4',], skiprows = 1)
```

In [43]:

```
pCal_eGFP = bokeh.plotting.figure(toolbar_location='right',
    outline_line_color= None,
    min_border_right=10,
    height=400,
    width=600,)

pCal_eGFP.title.text = "B) eGFP Calibration: Ex/Em 485/515, Gain 61 "
pCal_eGFP.xaxis.axis_label = 'eGFP (μM)'
pCal_eGFP.yaxis.axis_label = 'Fluorescence (a.u.)'
pCal_eGFP.y_range=Range1d(0, 30000)
pCal_eGFP.x_range=Range1d(0, 18)
pCal_eGFP.outline_line_color=None

# pCal_eGFPyaxis
pCal_eGFP.ygrid.visible = False
pCal_eGFP.yaxis.axis_label_text_font_size='15pt' 
pCal_eGFP.yaxis.major_label_text_font_size = '15pt'
pCal_eGFP.yaxis.major_label_text_font='Work Sans'
pCal_eGFP.yaxis.axis_label_standoff=15
pCal_eGFP.yaxis.axis_label_text_font_style='normal' 

# pCal_eGFPxaxis
pCal_eGFP.xgrid.visible = False
pCal_eGFP.xaxis.axis_label_text_font_size='15pt' 
pCal_eGFP.xaxis.major_label_text_font_size = '15pt'
pCal_eGFP.xaxis.major_label_text_font='Work Sans'
pCal_eGFP.xaxis.axis_label_standoff=15
pCal_eGFP.xaxis.axis_label_text_font_style='normal' 

# pCal_eGFPtitle
pCal_eGFP.title.text_font_size= '18pt' 
pCal_eGFP.title.align= 'left'
pCal_eGFP.title.offset=-70.0


#Data from experiments
pCal_eGFP.scatter(eGFP_Data['eGFP (uM)'], eGFP_Data['Biotek 4'], radius=0.1, fill_alpha=0.5, color="navy")
m4, b4 = np.polyfit(eGFP_Data['eGFP (uM)'], eGFP_Data['Biotek 4'], 1)
pCal_eGFP.line(eGFP_Data['eGFP (uM)'],eGFP_Data['eGFP (uM)']*m4+b4 ,  color="#d95b43", line_dash= 'dashed', line_width=2) #trendline

citation4 = Label(x=200, y=100, x_units='screen', y_units='screen',
                 text='y= '+str(np.around(m4, decimals=2, out=None))+'x' + str(np.around(b4, decimals=2, out=None)), #render_mode='css',
                 border_line_color=None, border_line_alpha=1.0,
                 background_fill_color='white', background_fill_alpha=1.0, text_font_size='14pt')
pCal_eGFP.add_layout(citation4)

bokeh.io.show(pCal_eGFP)
```

#### Plot as Grid and save as .svg¶

In [44]:

```
# using grid layout on plots
pCal=gridplot([[pCal_mRNA, pCal_eGFP]], width=600, height=400)
show(pCal)
```

In [45]:

```
count=1
for pic in [pCal_mRNA, pCal_eGFP]:
    pic.output_backend = "svg"
    export_svgs(pic, filename = 'Figures_PURE_Model/Fig_pCal' + str(count) + '.svg',width=800, height=600)
    count+=1

export_svgs(pCal, filename = 'Figures_PURE_Model/Fig_pCal.svg', width=800, height=600)
```

```
C:\Users\zoila\anaconda3\envs\python38\lib\site-packages\bokeh\io\export.py:308: UserWarning: Export method called with width or height argument on a non-Plot model. The size values will be ignored.
  warnings.warn("Export method called with width or height argument on a non-Plot model. The size values will be ignored.")
```

Out[45]:

```
['Figures_PURE_Model/Fig_pCal.svg', 'Figures_PURE_Model/Fig_pCal_1.svg']
```

#### Computing environment¶

In [46]:

```
%load_ext watermark
%watermark -v -p bioscrape,bokeh,panel,jupyterlab,biocrnpyler
```

```
Python implementation: CPython
Python version       : 3.8.17
IPython version      : 8.12.2

bioscrape  : 1.2.1
bokeh      : 2.4.0
panel      : 0.13.1
jupyterlab : 3.6.5
biocrnpyler: 1.1.1
```
