## Supporting scripts, data, and simulation results. for "A chemical reaction network model of PURE": CRN_PURE_TLonly_Final.html


### PURE TL-only Model in BioCRNpyler¶

August 8, 2023

[1] Division of Engineering and Applied Science, California Institute of Technology, Pasadena, CA \
[2] Division of Biology and Biological Engineering, California Institute of Technology, Pasadena, CA \

The PURE TL model in BioCRNpyler is based on the PURE translation models from Matsuura *et al.* [2,3] originally modeled in MATLAB. This script extends the reactions explicitly listed in Matsuura’s model by allowing the user to input amino acids of arbitrary length. The PURE TL does not account for multiple tRNAs coding for the same amino acid. Additionally, all reactions where k is zero were commented out since our modeling engineering, bioscrape, does not allow reaction constants of value zero.
The reactions were broken up into 6 subsections: species and reactions based on amino acids needed (1-2) and species and reactions of growing peptide (3-6).

1. Non-nucleic acid Species and Reactions
2. Addition of AA individual reactions
3. Met Species and Reaction (Start codon)
4. Elongation
5. Termination (End codon)
6. Removal of Ribosome

##### Citation:¶

[2] Matsuura, T., Tanimura, N., Hosoda, K., and Shimizu, Y. Reaction dynamics analysis of a reconstituted *Escherichia coli* protein translation system by computational modeling. PNAS. 2017, 114, 8. (PNAS, 2017) \
[3] Matsuura, T., Hosoda, K., and Shimizu, Y. Robustness of a reconstituted *Escherichia coli* protein translation system analyzed by computational modeling. ACS Synth. Biol. 2018, 7, 8, 1964–1972. (ACS Synth. Biol., 2018)

### Importing required packages and definitions¶

#### Packages and plotting basics¶

In [1]:

```
from typing import List, Union
import numpy as np
import pandas as pd
import pylab as plt
import csv
import numpy as np
import pandas as pd
import math
import seaborn as sns; sns.set()
import matplotlib.pyplot as plt
import random 
import time
import datetime
from itertools import groupby

from biocrnpyler import *
import itertools as it
from biocrnpyler.component import Component
from biocrnpyler.chemical_reaction_network import Species, Reaction, ChemicalReactionNetwork
from biocrnpyler.mechanism import Mechanism
from biocrnpyler.reaction import Reaction
from biocrnpyler.species import Complex, Species, WeightedSpecies

#Get directory
import os
directory = os.getcwd()

def flatten(t):
    return [item for sublist in t for item in sublist]
```

##### Bokeh¶

In [2]:

```
%matplotlib inline
import bokeh.io
import bokeh.plotting
bokeh.io.output_notebook()
from bokeh.themes import Theme

# Modules needed from Bokeh.
from bokeh.io import output_file, show
from bokeh.plotting import figure
from bokeh.models import LinearAxis, Range1d

colors = bokeh.palettes.Colorblind[8]

try:
    import dnaplotlib as dpl
    dpl_enabled = True
except (ModuleNotFoundError,ImportError) as e:
    dpl_enabled = False
    
theme = Theme(json={'attrs': {
# apply defaults to Figure properties
'Figure': {
    'toolbar_location': 'right',
    'outline_line_color': None,
    'min_border_right': 10,
#     'sizing_mode': 'stretch_width',
    'height':600,
    'width':800,
},'Grid': {
    'grid_line_color': None,
},
'Title': {
    'text_font_size': '20pt',
    'align': 'center'
},
    
    
# apply defaults to Axis properties
'Axis': {
#     'minor_tick_out': None,
#     'minor_tick_in': None,
    'major_label_text_font_size': '15pt',
    'axis_label_text_font_size': '15pt',
#     'axis_label_text_font': 'Work Sans',
    'axis_label_text_font_style': 'normal',
    'axis_label_standoff':15
},


# apply defaults to Legend properties
'Legend': {
    'background_fill_alpha': 0.8,
    'location': 'top_right',
    "label_text_font_size": '15pt'
}}})

bokeh.io.curdoc().theme = theme
from bokeh.io import export_png
```

Loading BokehJS ...

### Define the directory where parameters and initial conditions are saved as a CSV.¶

In [3]:

```
filename_parameters = '/fMGG_synthesis_parameters_CRN.csv'

with open(directory + filename_parameters, mode='r') as infile:
    reader = csv.reader(infile)
    rxn_k= {rows[0]:float(rows[1]) for rows in reader}
```

### User defined transcribed unit¶

In [4]:

```
def translate(seq):
       
    table = {
        'ATA':'Ile', 'ATC':'Ile', 'ATT':'Ile', 'ATG':'Met',
        'ACA':'Thr', 'ACC':'Thr', 'ACG':'Thr', 'ACT':'Thr',
        'AAC':'Asn', 'AAT':'Asn', 'AAA':'Lys', 'AAG':'Lys',
        'AGC':'Ser', 'AGT':'Ser', 'AGA':'Arg', 'AGG':'Arg',                 
        'CTA':'Leu', 'CTC':'Leu', 'CTG':'Leu', 'CTT':'Leu',
        'CCA':'Pro', 'CCC':'Pro', 'CCG':'Pro', 'CCT':'Pro',
        'CAC':'His', 'CAT':'His', 'CAA':'Gln', 'CAG':'Gln',
        'CGA':'Arg', 'CGC':'Arg', 'CGG':'Arg', 'CGT':'Arg',
        'GTA':'Val', 'GTC':'Val', 'GTG':'Val', 'GTT':'Val',
        'GCA':'Ala', 'GCC':'Ala', 'GCG':'Ala', 'GCT':'Ala',
        'GAC':'Asp', 'GAT':'Asp', 'GAA':'Glu', 'GAG':'Glu',
        'GGA':'Gly', 'GGC':'Gly', 'GGG':'Gly', 'GGT':'Gly',
        'TCA':'Ser', 'TCC':'Ser', 'TCG':'Ser', 'TCT':'Ser',
        'TTC':'Phe', 'TTT':'Phe', 'TTA':'Leu', 'TTG':'Leu',
        'TAC':'Tyr', 'TAT':'Tyr', 'TAA':'_', 'TAG':'_',
        'TGC':'Cys', 'TGT':'Cys', 'TGA':'_', 'TGG':'Trp',
    }
    protein =[]
    if len(seq)%3 == 0:
        for i in range(0, len(seq), 3):
            codon = seq[i:i + 3]
            if table[codon] != '_':
                protein+= [table[codon]]
            else:
                break
    return protein
```

### Input the desired peptide or protein amino acid chain¶

In [5]:

```
#User will copy the sequence bwteen end of promoter and start of terminator of DNA strain
#deGFP
dna_seq='ATGGAGCTTTTCACTGGCGTTGTTCCCATCCTGGTCGAGCTGGACGGCGACGTAAACGGCCACAAGTTCAGCGTGTCCGGCGAGGGCGAGGGCGATGCCACCTACGGCAAGCTGACCCTGAAGTTCATCTGCACCACCGGCAAGCTGCCCGTGCCCTGGCCCACCCTCGTGACCACCCTGACCTACGGCGTGCAGTGCTTCAGCCGCTACCCCGACCACATGAAGCAGCACGACTTCTTCAAGTCCGCCATGCCCGAAGGCTACGTCCAGGAGCGCACCATCTTCTTCAAGGACGACGGCAACTACAAGACCCGCGCCGAGGTGAAGTTCGAGGGCGACACCCTGGTGAACCGCATCGAGCTGAAGGGCATCGACTTCAAGGAGGACGGCAACATCCTGGGGCACAAGCTGGAGTACAACTACAACAGCCACAACGTCTATATCATGGCCGACAAGCAGAAGAACGGCATCAAGGTGAACTTCAAGATCCGCCACAACATCGAGGACGGCAGCGTGCAGCTCGCCGACCACTACCAGCAGAACACCCCCATCGGCGACGGCCCCGTGCTGCTGCCCGACAACCACTACCTGAGCACCCAGTCCGCCCTGAGCAAAGACCCCAACGAGAAGCGCGATCACATGGTCCTGCTGGAGTTCGTGACCGCCGCCGGGATCTAA'

#amino acid seq used to test expanded TL-only model
# protein=['Met','Gly','Gly','Val','Ser','Trp','Arg','Leu','Phe','Lys','Ile','Met','Cys','Ala','Tyr','Thr','Gln','Asn','His','Pro','Asp','Glu']
```

#### Converting DNA seq to AA for TL¶

In [6]:

```
for bp in range(0,len(dna_seq)-2):
    bp_aa = dna_seq[bp:bp+3]
    if bp_aa=='ATG':
        print('start codon found')
        start = bp
        remainder= len(dna_seq[bp:]) % 3

        if remainder ==0:
            CDS= dna_seq
        else:
            CDS= dna_seq[bp:-remainder]
        # print(CDS)
        protein= translate(CDS)
        break
    if bp == len(dna_seq)-2:
        print('No start codon found')

x=protein
```

```
start codon found
```

### Translation (TL)¶

#### Non-nucleic acid Species and Reactions¶

Creates all the species and reaction around the amino acids in the given chain, common protein and small molecules.

In [7]:

```
# Makes a list of the aa needed in the given amino acid chain, removing any duplicates.
AA= list(set(x[1:],)) #removes any duplicated amino acids

#Makes empty array for species and reactions
list_of_reactions = []
list_species_aa=[]

#Reference lists for aa and respective codon.
list_AA =['Ala', 'Arg', 'Asn', 'Asp', 'Cys', 
          'Gln', 'Glu', 'Gly', 'His', 'Ile', 
          'Leu', 'Lys', 'Met', 'Phe', 'Pro',
          'Ser', 'Thr', 'Trp', 'Tyr', 'Val']
list_condon =['AGU', 'CCG', 'AUU','AUC','ACA',
             'UUG','UUC','GCC','AUG','AAU',
             'CAG','CUU','CAU','GAA','AGG',
             'GCU','AGU','CCA','AUA','CAC',]
```

#### General proteins and small molecules¶

In [8]:

```
#Make species for general proteins and small molecules
ADP = Species('ADP')
AMP = Species('AMP')
ATP = Species('ATP') 
CK = Species('CK')
CK_ADP = Species('CK_ADP')
CK_ATP = Species('CK_ATP')
CK_CP = Species('CK_CP')
CK_CP_ADP = Species('CK_CP_ADP')
CK_Cr = Species('CK_Cr')
CK_Cr_ATP = Species('CK_Cr_ATP')

CK_degraded = Species('CK_degraded')
CP = Species('CP')
Cr = Species('Cr')
EFG = Species('EFG')
EFG_degraded = Species('EFG_degraded')
EFG_GDP = Species('EFG_GDP')
EFG_GTP = Species('EFG_GTP')
EFTs = Species('EFTs')
EFTs_degraded = Species('EFTs_degraded')
EFTu = Species('EFTu')

EFTu_degraded = Species('EFTu_degraded')
EFTu_EFTs = Species('EFTu_EFTs')
EFTu_GDP = Species('EFTu_GDP')
EFTu_GDP_EFTs = Species('EFTu_GDP_EFTs')
EFTu_GTP = Species('EFTu_GTP')
EFTu_GTP_EFTs = Species('EFTu_GTP_EFTs')
FD = Species('FD')
GDP = Species('GDP')
GMP = Species('GMP')
GTP = Species('GTP')

IF1 = Species('IF1')
IF1_degraded = Species('IF1_degraded')
IF2 = Species('IF2')
IF2_degraded = Species('IF2_degraded')
IF2_GDP = Species('IF2_GDP')
IF2_GTP = Species('IF2_GTP')
IF2_GTP_fMettRNAfMetCAU = Species('IF2_GTP_fMettRNAfMetCAU')
IF3 = Species('IF3')
IF3_degraded = Species('IF3_degraded')
MK = Species('MK')

MK_ADP_1 = Species('MK_ADP_1')
MK_ADP_2 = Species('MK_ADP_2')
MK_ADP_ADP = Species('MK_ADP_ADP')
MK_AMP = Species('MK_AMP')
MK_ATP = Species('MK_ATP')
MK_ATP_AMP = Species('MK_ATP_AMP')
MK_degraded = Species('MK_degraded')
mRNA = Species('mRNA')
mRNA_degraded = Species('mRNA_degraded')
MTF = Species('MTF')

MTF_degraded = Species('MTF_degraded')
MTF_FD = Species('MTF_FD')
MTF_THF = Species('MTF_THF')
NDK = Species('NDK')
NDK_ADP = Species('NDK_ADP')
NDK_ATP = Species('NDK_ATP')
NDK_degraded = Species('NDK_degraded')
NDK_GDP = Species('NDK_GDP')
NDK_GDP_ATP = Species('NDK_GDP_ATP')
NDK_GTP = Species('NDK_GTP')

NDK_GTP_ADP = Species('NDK_GTP_ADP')
PO4 = Species('PO4')
PPi = Species('PPi')
PPiase = Species('PPiase')
PPiase_degraded = Species('PPiase_degraded')
PPiase_PO4 = Species('PPiase_PO4')
PPiase_PO4_PO4 = Species('PPiase_PO4_PO4')
PPiase_PPi = Species('PPiase_PPi')
RF1 = Species('RF1')
RF1_degraded = Species('RF1_degraded')

RF2 = Species('RF2')
RF2_degraded = Species('RF2_degraded')
RF3 = Species('RF3')
RF3_degraded = Species('RF3_degraded')
RF3_GDP = Species('RF3_GDP')
RF3_GTP = Species('RF3_GTP')
RRF = Species('RRF')
RRF_degraded = Species('RRF_degraded')

RS30S = Species('RS30S')
RS30S_degraded = Species('RS30S_degraded')
RS30S_IF1_IF3_IF2_GTP_mRNA = Species('RS30S_IF1_IF3_IF2_GTP_mRNA')
RS30S_IF1_IF3_mRNA = Species('RS30S_IF1_IF3_mRNA')
RS30S_IF1_mRNA = Species('RS30S_IF1_mRNA')
RS30S_IF1 = Species('RS30S_IF1')
RS30S_IF1_IF2_GTP = Species('RS30S_IF1_IF2_GTP')
RS30S_IF1_IF2_GTP_mRNA = Species('RS30S_IF1_IF2_GTP_mRNA')
RS30S_IF1_IF3 = Species('RS30S_IF1_IF3')
RS30S_IF1_IF3_IF2_GTP = Species('RS30S_IF1_IF3_IF2_GTP')
RS30S_IF3_IF2_GTP = Species('RS30S_IF3_IF2_GTP')

RS30S_IF3_IF2_GTP_mRNA = Species('RS30S_IF3_IF2_GTP_mRNA')      
RS30S_IF2_GTP = Species('RS30S_IF2_GTP')
RS30S_IF2_GTP_mRNA = Species('RS30S_IF2_GTP_mRNA')
RS30S_IF3 = Species('RS30S_IF3')
RS30S_IF3_mRNA = Species('RS30S_IF3_mRNA')
RS30S_mRNA = Species('RS30S_mRNA')
RS50S = Species('RS50S')
RS50S_degraded = Species('RS50S_degraded')
RS50S_EFG_GDP = Species('RS50S_EFG_GDP')
RS50S_EFG_GDP_PO4 = Species('RS50S_EFG_GDP_PO4')

RS50S_EFG_GTP = Species('RS50S_EFG_GTP')
RS50S_RRF = Species('RS50S_RRF')
RS50S_RRF_EFG_GDP = Species('RS50S_RRF_EFG_GDP')
RS70S = Species('RS70S')
RS70S_EFG_GDP = Species('RS70S_EFG_GDP')
RS70S_EFG_GDP_PO4 = Species('RS70S_EFG_GDP_PO4')
RS70S_EFG_GTP = Species('RS70S_EFG_GTP')
RS70S_IF1 = Species('RS70S_IF1')
RS70S_IF3 = Species('RS70S_IF3')
RS70S_IF1_IF3 = Species('RS70S_IF1_IF3')         

THF = Species('THF')
fMet = Species('fMet')
fMet_degraded = Species('fMet_degraded')

#######################################################################################################################

#Creates a list of all the general species
list_species_gen=[ADP, AMP, ATP, CK, CK_ADP, CK_ATP, 
                  CK_CP, CK_CP_ADP, CK_Cr, CK_Cr_ATP, CK_degraded,
                  CP, Cr, EFG, EFG_degraded, EFG_GDP,
                  EFG_GTP, EFTs, EFTs_degraded, EFTu, EFTu_degraded,
                  EFTu_EFTs, EFTu_GDP, EFTu_GDP_EFTs, EFTu_GTP, EFTu_GTP_EFTs,
                  FD, GDP, GMP, GTP, IF1,
                  IF1_degraded, IF2, IF2_degraded, IF2_GDP, IF2_GTP,
                  IF2_GTP_fMettRNAfMetCAU, IF3, IF3_degraded, MK, MK_ADP_1, MK_ADP_2,
                  MK_ADP_ADP, MK_AMP, MK_ATP, MK_ATP_AMP, MK_degraded,
                  mRNA, mRNA_degraded, MTF, MTF_degraded, MTF_FD, MTF_THF,
                  NDK, NDK_ADP, NDK_ATP, NDK_degraded, NDK_GDP, NDK_GDP_ATP,
                  NDK_GTP, NDK_GTP_ADP, PO4, PPi, PPiase, PPiase_degraded,
                  PPiase_PO4, PPiase_PO4_PO4, PPiase_PPi, RF1, RF1_degraded,
                  RF2, RF2_degraded, RF3, RF3_degraded, RF3_GDP,
                  RF3_GTP, RRF, RRF_degraded, RS30S, RS30S_degraded,
                  RS30S_IF1_IF3_IF2_GTP_mRNA, RS30S_IF1_IF3_mRNA, RS30S_IF1_mRNA, RS30S_IF1, RS30S_IF1_IF2_GTP,
                  RS30S_IF1_IF2_GTP_mRNA, RS30S_IF1_IF3, RS30S_IF1_IF3_IF2_GTP, RS30S_IF3_IF2_GTP_mRNA,   
                  RS30S_IF2_GTP, RS30S_IF2_GTP_mRNA, RS30S_IF3, RS30S_IF3_mRNA, RS30S_IF3_IF2_GTP,
                  RS30S_mRNA, RS50S, RS50S_degraded, RS50S_EFG_GDP, RS50S_EFG_GDP_PO4,
                  RS50S_EFG_GTP, RS50S_RRF, RS50S_RRF_EFG_GDP, RS70S, RS70S_EFG_GDP,
                  RS70S_EFG_GDP_PO4, RS70S_EFG_GTP, RS70S_IF1, RS70S_IF3, RS70S_IF1_IF3,        
                  THF, fMet, fMet_degraded,]

#######################################################################################################################

#Creates a list of all the reactions only involving the general species. The reaction rates are read from the rxn_k dictionary created initially.
list_of_reaction_gen= [
    # Reaction.from_massaction([RS50S],[RS50S_degraded], k_forward = rxn_k['re0000000003_k1']),
    # Reaction.from_massaction([RS30S],[RS30S_degraded], k_forward = rxn_k['re0000000004_k1']),
    # Reaction.from_massaction([fMet],[fMet_degraded], k_forward = rxn_k['re0000000007_k1']),
    # Reaction.from_massaction([mRNA],[mRNA_degraded], k_forward = rxn_k['re0000000008_k1']),
    # Reaction.from_massaction([EFTu],[EFTu_degraded], k_forward = rxn_k['re0000000029_k1']),
    # Reaction.from_massaction([EFG],[EFG_degraded], k_forward = rxn_k['re0000000030_k1']),
    
    Reaction.from_massaction([EFTu,EFTs],[EFTu_EFTs], k_forward = rxn_k['re0000000261_k1']),
    Reaction.from_massaction([EFTu_EFTs],[EFTu,EFTs], k_forward = rxn_k['re0000000262_k1']),
    Reaction.from_massaction([EFTu_EFTs,GDP],[EFTu_GDP_EFTs], k_forward = rxn_k['re0000000263_k1']),
    Reaction.from_massaction([EFTu_GDP_EFTs],[EFTu_EFTs,GDP], k_forward = rxn_k['re0000000264_k1']),
    Reaction.from_massaction([EFTu_GDP_EFTs],[EFTu_GDP,EFTs], k_forward = rxn_k['re0000000265_k1']),
    Reaction.from_massaction([EFTu_GDP,EFTs],[EFTu_GDP_EFTs], k_forward = rxn_k['re0000000266_k1']),
    Reaction.from_massaction([EFTu_GDP],[EFTu,GDP], k_forward = rxn_k['re0000000267_k1']),
    Reaction.from_massaction([EFTu,GDP],[EFTu_GDP], k_forward = rxn_k['re0000000268_k1']),
    Reaction.from_massaction([EFTu_EFTs,GTP],[EFTu_GTP_EFTs], k_forward = rxn_k['re0000000269_k1']),
    Reaction.from_massaction([EFTu_GTP_EFTs],[EFTu_EFTs,GTP], k_forward = rxn_k['re0000000270_k1']),
    Reaction.from_massaction([EFTu_GTP_EFTs],[EFTu_GTP,EFTs], k_forward = rxn_k['re0000000271_k1']),
    Reaction.from_massaction([EFTu_GTP,EFTs],[EFTu_GTP_EFTs], k_forward = rxn_k['re0000000272_k1']),
    Reaction.from_massaction([EFTu,GTP],[EFTu_GTP], k_forward = rxn_k['re0000000273_k1']),
    Reaction.from_massaction([EFTu_GTP],[EFTu,GTP], k_forward = rxn_k['re0000000274_k1']),
    
    # Reaction.from_massaction([EFTs],[EFTs_degraded], k_forward = rxn_k['re0000000277_k1']),
    # Reaction.from_massaction([EFTu_GTP],[EFTu_degraded,GTP], k_forward = rxn_k['re0000000278_k1']),
    # Reaction.from_massaction([EFTu_GDP],[EFTu_degraded,GDP], k_forward = rxn_k['re0000000279_k1']),
    # Reaction.from_massaction([EFTu_EFTs],[EFTu_degraded,EFTs], k_forward = rxn_k['re0000000280_k1']),
    # Reaction.from_massaction([EFTu_EFTs],[EFTs_degraded,EFTu], k_forward = rxn_k['re0000000281_k1']),
    # Reaction.from_massaction([EFTu_GDP_EFTs],[EFTu_degraded,EFTs,GDP], k_forward = rxn_k['re0000000282_k1']),
    # Reaction.from_massaction([EFTu_GDP_EFTs],[EFTs_degraded,EFTu,GDP], k_forward = rxn_k['re0000000283_k1']),
    # Reaction.from_massaction([EFTu_GTP_EFTs],[EFTu_degraded,EFTs,GTP], k_forward = rxn_k['re0000000284_k1']),
    # Reaction.from_massaction([EFTu_GTP_EFTs],[EFTs_degraded,EFTu,GTP], k_forward = rxn_k['re0000000285_k1']),

    Reaction.from_massaction([EFG_GDP],[EFG,GDP], k_forward = rxn_k['re0000000292_k1']),
    Reaction.from_massaction([EFG,GDP],[EFG_GDP], k_forward = rxn_k['re0000000293_k1']),
    Reaction.from_massaction([EFG,GTP],[EFG_GTP], k_forward = rxn_k['re0000000294_k1']),
    Reaction.from_massaction([EFG_GTP],[EFG,GTP], k_forward = rxn_k['re0000000295_k1']),
    # Reaction.from_massaction([EFG_GTP],[EFG_degraded,GTP], k_forward = rxn_k['re0000000296_k1']),
    # Reaction.from_massaction([EFG_GDP],[EFG_degraded,GDP], k_forward = rxn_k['re0000000297_k1']),
    Reaction.from_massaction([RS50S,EFG_GTP],[RS50S_EFG_GTP], k_forward = rxn_k['re0000000298_k1']),
    Reaction.from_massaction([RS50S_EFG_GTP],[RS50S,EFG_GTP], k_forward = rxn_k['re0000000299_k1']),
    Reaction.from_massaction([RS70S,EFG_GTP],[RS70S_EFG_GTP], k_forward = rxn_k['re0000000300_k1']),
    Reaction.from_massaction([RS70S_EFG_GTP],[RS70S,EFG_GTP], k_forward = rxn_k['re0000000301_k1']),

    Reaction.from_massaction([RS50S_EFG_GTP],[RS50S_EFG_GDP_PO4], k_forward = rxn_k['re0000000302_k1']),
    Reaction.from_massaction([RS50S_EFG_GDP_PO4],[RS50S_EFG_GTP], k_forward = rxn_k['re0000000303_k1']),
    Reaction.from_massaction([RS70S_EFG_GTP],[RS70S_EFG_GDP_PO4], k_forward = rxn_k['re0000000304_k1']),
    Reaction.from_massaction([RS70S_EFG_GDP_PO4],[RS70S_EFG_GTP], k_forward = rxn_k['re0000000305_k1']),
    Reaction.from_massaction([RS50S_EFG_GDP_PO4],[RS50S_EFG_GDP,PO4], k_forward = rxn_k['re0000000306_k1']),
    Reaction.from_massaction([RS70S_EFG_GDP_PO4],[RS70S_EFG_GDP,PO4], k_forward = rxn_k['re0000000307_k1']),
    Reaction.from_massaction([RS50S_EFG_GDP],[RS50S,EFG_GDP], k_forward = rxn_k['re0000000308_k1']),
    Reaction.from_massaction([RS70S_EFG_GDP],[RS70S,EFG_GDP], k_forward = rxn_k['re0000000309_k1']),
    # Reaction.from_massaction([RS50S_EFG_GTP],[EFG_degraded,GTP,RS50S], k_forward = rxn_k['re0000000310_k1']),
    # Reaction.from_massaction([RS50S_EFG_GTP],[RS50S_degraded,GTP,EFG], k_forward = rxn_k['re0000000311_k1']),

    # Reaction.from_massaction([RS50S_EFG_GDP_PO4],[EFG_degraded,GDP,PO4,RS50S], k_forward = rxn_k['re0000000312_k1']),
    # Reaction.from_massaction([RS50S_EFG_GDP_PO4],[RS50S_degraded,EFG,GDP,PO4], k_forward = rxn_k['re0000000313_k1']),
    # Reaction.from_massaction([RS50S_EFG_GDP],[EFG_degraded,GDP,RS50S], k_forward = rxn_k['re0000000314_k1']),
    # Reaction.from_massaction([RS50S_EFG_GDP],[RS50S_degraded,EFG,GDP], k_forward = rxn_k['re0000000315_k1']),
    # Reaction.from_massaction([RS70S_EFG_GTP],[EFG_degraded,RS30S,GTP,RS50S], k_forward = rxn_k['re0000000316_k1']),
    # Reaction.from_massaction([RS70S_EFG_GTP],[RS50S_degraded,EFG,GTP,RS30S], k_forward = rxn_k['re0000000317_k1']),
    # Reaction.from_massaction([RS70S_EFG_GDP_PO4],[EFG_degraded,RS30S,GDP,PO4,RS50S], k_forward = rxn_k['re0000000318_k1']),
    # Reaction.from_massaction([RS70S_EFG_GDP_PO4],[RS50S_degraded,EFG,PO4,GDP,RS30S], k_forward = rxn_k['re0000000319_k1']),
    # Reaction.from_massaction([RS70S_EFG_GTP],[RS30S_degraded,EFG,RS50S,GTP], k_forward = rxn_k['re0000000320_k1']),
    # Reaction.from_massaction([RS70S_EFG_GDP_PO4],[RS30S_degraded,EFG,PO4,RS50S,GDP], k_forward = rxn_k['re0000000321_k1']),

    # Reaction.from_massaction([RS70S_EFG_GDP],[RS50S_degraded,EFG,GDP,RS30S], k_forward = rxn_k['re0000000322_k1']),
    # Reaction.from_massaction([RS70S_EFG_GDP],[EFG_degraded,RS30S,GDP,RS50S], k_forward = rxn_k['re0000000323_k1']),
    # Reaction.from_massaction([RS70S_EFG_GDP],[RS30S_degraded,EFG,RS50S,GDP], k_forward = rxn_k['re0000000324_k1']),
    # Reaction.from_massaction([RS50S_EFG_GDP,PO4],[RS50S_EFG_GDP_PO4], k_forward = rxn_k['re0000000325_k1']),
    # Reaction.from_massaction([RS70S_EFG_GDP,PO4],[RS70S_EFG_GDP_PO4], k_forward = rxn_k['re0000000326_k1']),
    # Reaction.from_massaction([RS50S,EFG_GDP],[RS50S_EFG_GDP], k_forward = rxn_k['re0000000327_k1']),
    # Reaction.from_massaction([RS70S,EFG_GDP],[RS70S_EFG_GDP], k_forward = rxn_k['re0000000328_k1']),
    # Reaction.from_massaction([CK],[CK_degraded], k_forward = rxn_k['re0000000329_k1']),
    Reaction.from_massaction([CK,ADP],[CK_ADP], k_forward = rxn_k['re0000000330_k1']),
    Reaction.from_massaction([CK_ADP],[CK,ADP], k_forward = rxn_k['re0000000331_k1']),

    Reaction.from_massaction([CK,CP],[CK_CP], k_forward = rxn_k['re0000000332_k1']),
    Reaction.from_massaction([CK_CP],[CK,CP], k_forward = rxn_k['re0000000333_k1']),
    Reaction.from_massaction([CK_CP,ADP],[CK_CP_ADP], k_forward = rxn_k['re0000000334_k1']),
    Reaction.from_massaction([CK_CP_ADP],[CK_CP,ADP], k_forward = rxn_k['re0000000335_k1']),
    Reaction.from_massaction([CK_ADP,CP],[CK_CP_ADP], k_forward = rxn_k['re0000000336_k1']),
    Reaction.from_massaction([CK_CP_ADP],[CP,CK_ADP], k_forward = rxn_k['re0000000337_k1']),
    Reaction.from_massaction([CK_CP_ADP],[CK_Cr_ATP], k_forward = rxn_k['re0000000338_k1']),
    Reaction.from_massaction([CK_Cr_ATP],[CK_CP_ADP], k_forward = rxn_k['re0000000339_k1']),
    Reaction.from_massaction([CK_Cr_ATP],[CK_Cr,ATP], k_forward = rxn_k['re0000000340_k1']),
    Reaction.from_massaction([CK_Cr,ATP],[CK_Cr_ATP], k_forward = rxn_k['re0000000341_k1']),

    Reaction.from_massaction([CK_Cr_ATP],[CK_ATP,Cr], k_forward = rxn_k['re0000000342_k1']),
    Reaction.from_massaction([CK_ATP,Cr],[CK_Cr_ATP], k_forward = rxn_k['re0000000343_k1']),
    Reaction.from_massaction([CK_ATP],[CK,ATP], k_forward = rxn_k['re0000000344_k1']),
    Reaction.from_massaction([CK,ATP],[CK_ATP], k_forward = rxn_k['re0000000345_k1']),
    Reaction.from_massaction([CK_Cr],[CK,Cr], k_forward = rxn_k['re0000000346_k1']),
    Reaction.from_massaction([CK,Cr],[CK_Cr], k_forward = rxn_k['re0000000347_k1']),
    # Reaction.from_massaction([CK_CP],[CK_degraded,CP], k_forward = rxn_k['re0000000348_k1']),
    # Reaction.from_massaction([CK_ADP],[CK_degraded,ADP], k_forward = rxn_k['re0000000349_k1']),
    # Reaction.from_massaction([CK_CP_ADP],[CK_degraded,ADP,CP], k_forward = rxn_k['re0000000350_k1']),
    # Reaction.from_massaction([CK_Cr_ATP],[CK_degraded,ATP,Cr], k_forward = rxn_k['re0000000351_k1']),

    # Reaction.from_massaction([CK_ATP],[CK_degraded,ATP], k_forward = rxn_k['re0000000352_k1']),
    # Reaction.from_massaction([CK_Cr],[CK_degraded,Cr], k_forward = rxn_k['re0000000353_k1']),
    # Reaction.from_massaction([NDK],[NDK_degraded], k_forward = rxn_k['re0000000354_k1']),
    Reaction.from_massaction([NDK,ATP],[NDK_ATP], k_forward = rxn_k['re0000000355_k1']),
    Reaction.from_massaction([NDK_ATP],[NDK,ATP], k_forward = rxn_k['re0000000356_k1']),
    Reaction.from_massaction([NDK,GDP],[NDK_GDP], k_forward = rxn_k['re0000000357_k1']),
    Reaction.from_massaction([NDK_GDP],[NDK,GDP], k_forward = rxn_k['re0000000358_k1']),
    Reaction.from_massaction([NDK_GDP,ATP],[NDK_GDP_ATP], k_forward = rxn_k['re0000000359_k1']),
    Reaction.from_massaction([NDK_GDP_ATP],[NDK_GDP,ATP], k_forward = rxn_k['re0000000360_k1']),
    Reaction.from_massaction([NDK_ATP,GDP],[NDK_GDP_ATP], k_forward = rxn_k['re0000000361_k1']),

    Reaction.from_massaction([NDK_GDP_ATP],[NDK_ATP,GDP], k_forward = rxn_k['re0000000362_k1']),
    Reaction.from_massaction([NDK_GDP_ATP],[NDK_GTP_ADP], k_forward = rxn_k['re0000000363_k1']),
    # Reaction.from_massaction([NDK_GTP_ADP],[NDK_GDP_ATP], k_forward = rxn_k['re0000000364_k1']),
    Reaction.from_massaction([NDK_GTP_ADP],[NDK_ADP,GTP], k_forward = rxn_k['re0000000365_k1']),
    Reaction.from_massaction([NDK_GTP_ADP],[NDK_GTP,ADP], k_forward = rxn_k['re0000000366_k1']),
    Reaction.from_massaction([NDK_ADP],[NDK,ADP], k_forward = rxn_k['re0000000367_k1']),
    Reaction.from_massaction([NDK_GTP],[NDK,GTP], k_forward = rxn_k['re0000000368_k1']),
    # Reaction.from_massaction([NDK_GDP],[NDK_degraded,GDP], k_forward = rxn_k['re0000000369_k1']),
    # Reaction.from_massaction([NDK_ATP],[NDK_degraded,ATP], k_forward = rxn_k['re0000000370_k1']),
    # Reaction.from_massaction([NDK_GDP_ATP],[NDK_degraded,ATP,GDP], k_forward = rxn_k['re0000000371_k1']),

    # Reaction.from_massaction([NDK_GTP_ADP],[NDK_degraded,ADP,GTP], k_forward = rxn_k['re0000000372_k1']),
    # Reaction.from_massaction([NDK_ADP],[NDK_degraded,ADP], k_forward = rxn_k['re0000000373_k1']),
    # Reaction.from_massaction([NDK_GTP],[NDK_degraded,GTP], k_forward = rxn_k['re0000000374_k1']),
    Reaction.from_massaction([NDK_GTP,ADP],[NDK_GTP_ADP], k_forward = rxn_k['re0000000375_k1']),
    Reaction.from_massaction([NDK_ADP,GTP],[NDK_GTP_ADP], k_forward = rxn_k['re0000000376_k1']),
    Reaction.from_massaction([NDK,GTP],[NDK_GTP], k_forward = rxn_k['re0000000377_k1']),
    Reaction.from_massaction([NDK,ADP],[NDK_ADP], k_forward = rxn_k['re0000000378_k1']),
    # Reaction.from_massaction([MK],[MK_degraded], k_forward = rxn_k['re0000000379_k1']),
    Reaction.from_massaction([MK,ATP],[MK_ATP], k_forward = rxn_k['re0000000380_k1']),
    Reaction.from_massaction([MK_ATP],[ATP,MK], k_forward = rxn_k['re0000000381_k1']),

    Reaction.from_massaction([MK,AMP],[MK_AMP], k_forward = rxn_k['re0000000382_k1']),
    Reaction.from_massaction([MK_AMP],[MK,AMP], k_forward = rxn_k['re0000000383_k1']),
    Reaction.from_massaction([MK_AMP,ATP],[MK_ATP_AMP], k_forward = rxn_k['re0000000384_k1']),
    Reaction.from_massaction([MK_ATP_AMP],[ATP,MK_AMP], k_forward = rxn_k['re0000000385_k1']),
    Reaction.from_massaction([MK_ATP,AMP],[MK_ATP_AMP], k_forward = rxn_k['re0000000386_k1']),
    Reaction.from_massaction([MK_ATP_AMP],[MK_ATP,AMP], k_forward = rxn_k['re0000000387_k1']),
    Reaction.from_massaction([MK_ATP_AMP],[MK_ADP_ADP], k_forward = rxn_k['re0000000388_k1']),
    Reaction.from_massaction([MK_ADP_ADP],[MK_ATP_AMP], k_forward = rxn_k['re0000000389_k1']),
    Reaction.from_massaction([MK_ADP_ADP],[MK_ADP_1,ADP], k_forward = rxn_k['re0000000390_k1']),
    Reaction.from_massaction([MK_ADP_1,ADP],[MK_ADP_ADP], k_forward = rxn_k['re0000000391_k1']),

    Reaction.from_massaction([MK_ADP_ADP],[MK_ADP_2,ADP], k_forward = rxn_k['re0000000392_k1']),
    Reaction.from_massaction([MK_ADP_2,ADP],[MK_ADP_ADP], k_forward = rxn_k['re0000000393_k1']),
    Reaction.from_massaction([MK_ADP_1],[MK,ADP], k_forward = rxn_k['re0000000394_k1']),
    Reaction.from_massaction([MK,ADP],[MK_ADP_1], k_forward = rxn_k['re0000000395_k1']),
    Reaction.from_massaction([MK_ADP_2],[ADP,MK], k_forward = rxn_k['re0000000396_k1']),
    Reaction.from_massaction([MK,ADP],[MK_ADP_2], k_forward = rxn_k['re0000000397_k1']),
    # Reaction.from_massaction([MK_AMP],[MK_degraded,AMP], k_forward = rxn_k['re0000000398_k1']),
    # Reaction.from_massaction([MK_ATP],[MK_degraded,ATP], k_forward = rxn_k['re0000000399_k1']),
    # Reaction.from_massaction([MK_ATP_AMP],[MK_degraded,ATP], k_forward = rxn_k['re0000000400_k1']),
    # Reaction.from_massaction([MK_ADP_ADP],[MK_degraded,ADP], k_forward = rxn_k['re0000000401_k1']),

    # Reaction.from_massaction([MK_ADP_1],[MK_degraded,ADP], k_forward = rxn_k['re0000000402_k1']),
    # Reaction.from_massaction([MK_ADP_2],[MK_degraded,ADP], k_forward = rxn_k['re0000000403_k1']),
    # Reaction.from_massaction([PPiase],[PPiase_degraded], k_forward = rxn_k['re0000000404_k1']),
    Reaction.from_massaction([PPiase,PPi],[PPiase_PPi], k_forward = rxn_k['re0000000405_k1']),
    Reaction.from_massaction([PPiase_PPi],[PPiase,PPi], k_forward = rxn_k['re0000000406_k1']),
    Reaction.from_massaction([PPiase_PPi],[PPiase_PO4_PO4], k_forward = rxn_k['re0000000407_k1']),
    Reaction.from_massaction([PPiase_PO4_PO4],[PPiase_PPi], k_forward = rxn_k['re0000000408_k1']),
    Reaction.from_massaction([PPiase_PO4_PO4],[PPiase_PO4,PO4], k_forward = rxn_k['re0000000409_k1']),
    Reaction.from_massaction([PPiase_PO4,PO4],[PPiase_PO4_PO4], k_forward = rxn_k['re0000000410_k1']),
    Reaction.from_massaction([PPiase_PO4],[PPiase,PO4], k_forward = rxn_k['re0000000411_k1']),

    Reaction.from_massaction([PPiase,PO4],[PPiase_PO4], k_forward = rxn_k['re0000000412_k1']),
    # Reaction.from_massaction([PPiase_PPi],[PPiase_degraded,PPi], k_forward = rxn_k['re0000000413_k1']),
    # Reaction.from_massaction([PPiase_PO4_PO4],[PPiase_degraded,PO4], k_forward = rxn_k['re0000000414_k1']),
    # Reaction.from_massaction([PPiase_PO4],[PPiase_degraded,PO4], k_forward = rxn_k['re0000000415_k1']),
    # Reaction.from_massaction([MTF],[MTF_degraded], k_forward = rxn_k['re0000000416_k1']),
    Reaction.from_massaction([MTF,FD],[MTF_FD], k_forward = rxn_k['re0000000418_k1']),
    Reaction.from_massaction([MTF_FD],[MTF,FD], k_forward = rxn_k['re0000000419_k1']),
    Reaction.from_massaction([MTF_THF],[MTF,THF], k_forward = rxn_k['re0000000434_k1']),
    # Reaction.from_massaction([MTF,THF],[MTF_THF], k_forward = rxn_k['re0000000435_k1']),
    # Reaction.from_massaction([MTF_FD],[MTF_degraded,FD], k_forward = rxn_k['re0000000436_k1']),

    # Reaction.from_massaction([MTF_THF],[MTF_degraded,THF], k_forward = rxn_k['re0000000441_k1']),
    # Reaction.from_massaction([FD],[THF], k_forward = rxn_k['re0000000442_k1']),
    # Reaction.from_massaction([THF],[FD], k_forward = rxn_k['re0000000444_k1']),
    Reaction.from_massaction([IF2,GTP],[IF2_GTP], k_forward = rxn_k['re0000000445_k1']),
    Reaction.from_massaction([IF2_GTP],[GTP,IF2], k_forward = rxn_k['re0000000446_k1']),
    Reaction.from_massaction([IF2,GDP],[IF2_GDP], k_forward = rxn_k['re0000000447_k1']),
    Reaction.from_massaction([IF2_GDP],[IF2,GDP], k_forward = rxn_k['re0000000448_k1']),
    # Reaction.from_massaction([IF2],[IF2_degraded], k_forward = rxn_k['re0000000451_k1']),
    # Reaction.from_massaction([IF2_GDP],[IF2_degraded,GDP], k_forward = rxn_k['re0000000452_k1']),
    # Reaction.from_massaction([IF2_GTP],[IF2_degraded,GTP], k_forward = rxn_k['re0000000453_k1']),

    Reaction.from_massaction([RS70S],[RS30S,RS50S], k_forward = rxn_k['re0000000455_k1']),
    Reaction.from_massaction([RS30S,RS50S],[RS70S], k_forward = rxn_k['re0000000456_k1']),
    Reaction.from_massaction([RS70S,IF3],[RS70S_IF3], k_forward = rxn_k['re0000000457_k1']),
    Reaction.from_massaction([RS70S_IF3],[RS70S,IF3], k_forward = rxn_k['re0000000458_k1']),
    Reaction.from_massaction([RS30S,IF3],[RS30S_IF3], k_forward = rxn_k['re0000000459_k1']),
    Reaction.from_massaction([RS30S_IF3],[RS30S,IF3], k_forward = rxn_k['re0000000460_k1']),
    Reaction.from_massaction([RS70S_IF3],[RS30S_IF3,RS50S], k_forward = rxn_k['re0000000461_k1']),
    Reaction.from_massaction([RS30S_IF3,RS50S],[RS70S_IF3], k_forward = rxn_k['re0000000462_k1']),
    Reaction.from_massaction([RS30S_IF3,IF2_GTP],[RS30S_IF3_IF2_GTP], k_forward = rxn_k['re0000000463_k1']),
    Reaction.from_massaction([RS30S_IF3_IF2_GTP],[RS30S_IF3,IF2_GTP], k_forward = rxn_k['re0000000464_k1']),

    Reaction.from_massaction([RS30S_IF3,mRNA],[RS30S_IF3_mRNA], k_forward= rxn_k['re0000000469_k1']),
    Reaction.from_massaction([RS30S_IF3_mRNA],[RS30S_IF3,mRNA], k_forward= rxn_k['re0000000470_k1']),
    Reaction.from_massaction([RS30S_IF3_mRNA,IF2_GTP],[RS30S_IF3_IF2_GTP_mRNA], k_forward= rxn_k['re0000000471_k1']),
    Reaction.from_massaction([RS30S_IF3_IF2_GTP_mRNA],[RS30S_IF3_mRNA,IF2_GTP], k_forward= rxn_k['re0000000472_k1']),
    Reaction.from_massaction([RS30S_IF3_IF2_GTP,mRNA],[RS30S_IF3_IF2_GTP_mRNA], k_forward= rxn_k['re0000000481_k1']),
    Reaction.from_massaction([RS30S_IF3_IF2_GTP_mRNA],[RS30S_IF3_IF2_GTP,mRNA], k_forward= rxn_k['re0000000482_k1']),
    Reaction.from_massaction([RS70S_IF1],[RS30S_IF1,RS50S], k_forward = rxn_k['re0000000487_k1']),
    Reaction.from_massaction([RS30S_IF1,RS50S],[RS70S_IF1], k_forward = rxn_k['re0000000488_k1']),
    Reaction.from_massaction([RS70S_IF1,IF3],[RS70S_IF1_IF3], k_forward = rxn_k['re0000000489_k1']),
    Reaction.from_massaction([RS70S_IF1_IF3],[RS70S_IF1,IF3], k_forward = rxn_k['re0000000490_k1']),

    Reaction.from_massaction([RS30S_IF1,IF3],[RS30S_IF1_IF3], k_forward = rxn_k['re0000000491_k1']),
    Reaction.from_massaction([RS30S_IF1_IF3],[RS30S_IF1,IF3], k_forward = rxn_k['re0000000492_k1']),
    Reaction.from_massaction([RS70S_IF1_IF3],[RS30S_IF1_IF3,RS50S], k_forward = rxn_k['re0000000493_k1']),
    Reaction.from_massaction([RS30S_IF1_IF3,RS50S],[RS70S_IF1_IF3], k_forward = rxn_k['re0000000494_k1']),
    Reaction.from_massaction([RS30S_IF1_IF3,IF2_GTP],[RS30S_IF1_IF3_IF2_GTP], k_forward = rxn_k['re0000000495_k1']),
    Reaction.from_massaction([RS30S_IF1_IF3_IF2_GTP],[RS30S_IF1_IF3,IF2_GTP], k_forward = rxn_k['re0000000496_k1']),
    Reaction.from_massaction([RS70S,IF1],[RS70S_IF1], k_forward= rxn_k['re0000000501_k1']),
    Reaction.from_massaction([RS70S_IF1],[RS70S,IF1], k_forward = rxn_k['re0000000502_k1']),
    Reaction.from_massaction([RS30S,IF1],[RS30S_IF1], k_forward = rxn_k['re0000000503_k1']),
    Reaction.from_massaction([RS30S_IF1],[RS30S,IF1], k_forward = rxn_k['re0000000504_k1']),

    Reaction.from_massaction([RS70S_IF3,IF1],[RS70S_IF1_IF3], k_forward = rxn_k['re0000000505_k1']),
    Reaction.from_massaction([RS70S_IF1_IF3],[RS70S_IF3,IF1], k_forward = rxn_k['re0000000506_k1']),
    Reaction.from_massaction([RS30S_IF3,IF1],[RS30S_IF1_IF3], k_forward= rxn_k['re0000000507_k1']),
    Reaction.from_massaction([RS30S_IF1_IF3],[RS30S_IF3,IF1], k_forward = rxn_k['re0000000508_k1']),
    Reaction.from_massaction([RS30S_IF3_IF2_GTP,IF1],[RS30S_IF1_IF3_IF2_GTP], k_forward= rxn_k['re0000000509_k1']),
    Reaction.from_massaction([RS30S_IF1_IF3_IF2_GTP],[RS30S_IF3_IF2_GTP,IF1], k_forward = rxn_k['re0000000510_k1']),
    Reaction.from_massaction([RS30S_IF1_IF3,mRNA],[RS30S_IF1_IF3_mRNA], k_forward = rxn_k['re0000000513_k1']),
    Reaction.from_massaction([RS30S_IF1_IF3_mRNA],[RS30S_IF1_IF3,mRNA], k_forward = rxn_k['re0000000514_k1']),
    Reaction.from_massaction([RS30S_IF1_IF3_mRNA,IF2_GTP],[RS30S_IF1_IF3_IF2_GTP_mRNA], k_forward = rxn_k['re0000000515_k1']),
    Reaction.from_massaction([RS30S_IF1_IF3_IF2_GTP_mRNA],[RS30S_IF1_IF3_mRNA,IF2_GTP], k_forward = rxn_k['re0000000516_k1']),
    Reaction.from_massaction([RS30S_IF1_IF3_IF2_GTP,mRNA],[RS30S_IF1_IF3_IF2_GTP_mRNA], k_forward = rxn_k['re0000000525_k1']),

    Reaction.from_massaction([RS30S_IF1_IF3_IF2_GTP_mRNA],[RS30S_IF1_IF3_IF2_GTP,mRNA], k_forward = rxn_k['re0000000526_k1']),
    Reaction.from_massaction([RS30S_IF3_mRNA,IF1],[RS30S_IF1_IF3_mRNA], k_forward = rxn_k['re0000000531_k1']),
    Reaction.from_massaction([RS30S_IF1_IF3_mRNA],[RS30S_IF3_mRNA,IF1], k_forward = rxn_k['re0000000532_k1']),
    Reaction.from_massaction([RS30S_IF3_IF2_GTP_mRNA,IF1],[RS30S_IF1_IF3_IF2_GTP_mRNA], k_forward= rxn_k['re0000000535_k1']),
    Reaction.from_massaction([RS30S_IF1_IF3_IF2_GTP_mRNA],[RS30S_IF3_IF2_GTP_mRNA,IF1], k_forward = rxn_k['re0000000536_k1']),
    # Reaction.from_massaction([IF1],[IF1_degraded], k_forward = rxn_k['re0000000541_k1']),
    # Reaction.from_massaction([IF3],[IF3_degraded], k_forward = rxn_k['re0000000542_k1']),
    # Reaction.from_massaction([RS70S],[RS30S_degraded,RS50S], k_forward= rxn_k['re0000000543_k1']),
    # Reaction.from_massaction([RS70S],[RS50S_degraded,RS30S], k_forward= rxn_k['re0000000544_k1']),
    # Reaction.from_massaction([RS30S_IF1],[IF1_degraded,RS30S], k_forward = rxn_k['re0000000545_k1']),
    # Reaction.from_massaction([RS30S_IF1],[RS30S_degraded,IF1], k_forward = rxn_k['re0000000546_k1']),

    # Reaction.from_massaction([RS30S_IF3],[IF3_degraded,RS30S], k_forward= rxn_k['re0000000547_k1']),
    # Reaction.from_massaction([RS30S_IF3],[RS30S_degraded,IF3], k_forward= rxn_k['re0000000548_k1']),
    # Reaction.from_massaction([RS30S_IF3_mRNA],[IF3_degraded,mRNA,RS30S], k_forward= rxn_k['re0000000549_k1']),
    # Reaction.from_massaction([RS30S_IF3_mRNA],[RS30S_degraded,mRNA,IF3], k_forward= rxn_k['re0000000550_k1']),
    # Reaction.from_massaction([RS70S_IF3],[RS30S_degraded,RS50S,IF3], k_forward = rxn_k['re0000000553_k1']),
    # Reaction.from_massaction([RS70S_IF3],[RS50S_degraded,RS30S,IF3], k_forward = rxn_k['re0000000554_k1']),
    # Reaction.from_massaction([RS70S_IF3],[IF3_degraded,RS50S,RS30S], k_forward = rxn_k['re0000000555_k1']),
    # Reaction.from_massaction([RS70S_IF1],[RS30S_degraded,RS50S,IF1], k_forward = rxn_k['re0000000556_k1']),
    # Reaction.from_massaction([RS70S_IF1],[RS50S_degraded,RS30S,IF1], k_forward = rxn_k['re0000000557_k1']),
    # Reaction.from_massaction([RS70S_IF1],[IF1_degraded,RS50S,RS30S], k_forward = rxn_k['re0000000558_k1']),

    # Reaction.from_massaction([RS30S_IF1_IF3],[IF3_degraded,RS30S,IF1], k_forward = rxn_k['re0000000559_k1']),
    # Reaction.from_massaction([RS30S_IF1_IF3],[RS30S_degraded,IF3,IF1], k_forward = rxn_k['re0000000560_k1']),
    # Reaction.from_massaction([RS30S_IF1_IF3],[IF1_degraded,RS30S,IF3], k_forward = rxn_k['re0000000561_k1']),
    # Reaction.from_massaction([RS30S_IF3_IF2_GTP],[RS30S_degraded,GTP,IF2,IF3], k_forward= rxn_k['re0000000562_k1']),
    # Reaction.from_massaction([RS30S_IF3_IF2_GTP],[IF2_degraded,RS30S,GTP,IF3], k_forward= rxn_k['re0000000563_k1']),
    # Reaction.from_massaction([RS30S_IF3_IF2_GTP],[IF3_degraded,GTP,IF2,RS30S], k_forward= rxn_k['re0000000564_k1']),
    # Reaction.from_massaction([RS30S_IF1_IF3_mRNA],[IF3_degraded,mRNA,IF1,RS30S], k_forward = rxn_k['re0000000568_k1']),
    # Reaction.from_massaction([RS30S_IF1_IF3_mRNA],[RS30S_degraded,mRNA,IF3,IF1], k_forward = rxn_k['re0000000569_k1']),
    # Reaction.from_massaction([RS30S_IF1_IF3_mRNA],[IF1_degraded,mRNA,RS30S,IF3], k_forward = rxn_k['re0000000570_k1']),
    # Reaction.from_massaction([RS30S_IF3_IF2_GTP_mRNA],[RS30S_degraded,mRNA,GTP,IF3,IF2], k_forward= rxn_k['re0000000571_k1']),

    # Reaction.from_massaction([RS30S_IF3_IF2_GTP_mRNA],[IF3_degraded,GTP,mRNA,IF2,RS30S], k_forward= rxn_k['re0000000572_k1']),
    # Reaction.from_massaction([RS30S_IF3_IF2_GTP_mRNA],[IF2_degraded,mRNA,GTP,RS30S,IF3], k_forward= rxn_k['re0000000573_k1']),
    # Reaction.from_massaction([RS70S_IF1_IF3],[IF3_degraded,RS50S,RS30S,IF1], k_forward = rxn_k['re0000000580_k1']),
    # Reaction.from_massaction([RS70S_IF1_IF3],[RS50S_degraded,RS30S,IF3,IF1], k_forward = rxn_k['re0000000581_k1']),
    # Reaction.from_massaction([RS70S_IF1_IF3],[IF1_degraded,RS50S,RS30S,IF3], k_forward = rxn_k['re0000000582_k1']),
    # Reaction.from_massaction([RS70S_IF1_IF3],[RS30S_degraded,RS50S,IF3,IF1], k_forward = rxn_k['re0000000583_k1']),
    # Reaction.from_massaction([RS30S_IF1_IF3_IF2_GTP],[IF3_degraded,GTP,IF2,RS30S,IF1], k_forward = rxn_k['re0000000584_k1']),
    # Reaction.from_massaction([RS30S_IF1_IF3_IF2_GTP],[IF2_degraded,RS30S,GTP,IF3,IF1], k_forward = rxn_k['re0000000585_k1']),
    # Reaction.from_massaction([RS30S_IF1_IF3_IF2_GTP],[IF1_degraded,GTP,IF2,IF3,RS30S], k_forward = rxn_k['re0000000586_k1']),
    # Reaction.from_massaction([RS30S_IF1_IF3_IF2_GTP],[RS30S_degraded,GTP,IF2,IF3,IF1], k_forward = rxn_k['re0000000587_k1']),

    # Reaction.from_massaction([RS30S_IF1_IF3_IF2_GTP_mRNA],[IF3_degraded,mRNA,IF2,GTP,IF1,RS30S], k_forward = rxn_k['re0000000592_k1']),
    # Reaction.from_massaction([RS30S_IF1_IF3_IF2_GTP_mRNA],[RS30S_degraded,IF3,mRNA,GTP,IF2,IF1], k_forward = rxn_k['re0000000593_k1']),
    # Reaction.from_massaction([RS30S_IF1_IF3_IF2_GTP_mRNA],[IF2_degraded,RS30S,mRNA,GTP,IF3,IF1], k_forward = rxn_k['re0000000594_k1']),
    # Reaction.from_massaction([RS30S_IF1_IF3_IF2_GTP_mRNA],[IF1_degraded,mRNA,IF2,GTP,RS30S,IF3], k_forward = rxn_k['re0000000595_k1']),
    # Reaction.from_massaction([RS30S,IF2_GTP],[RS30S_IF2_GTP], k_forward = rxn_k['re0000000609_k1']),
    Reaction.from_massaction([RS30S_IF2_GTP],[IF2_GTP,RS30S], k_forward= rxn_k['re0000000610_k1']),
    Reaction.from_massaction([RS30S_IF2_GTP,IF3],[RS30S_IF3_IF2_GTP], k_forward= rxn_k['re0000000615_k1']),
    Reaction.from_massaction([RS30S_IF3_IF2_GTP],[IF3,RS30S_IF2_GTP], k_forward= rxn_k['re0000000616_k1']),
    Reaction.from_massaction([mRNA,RS30S],[RS30S_mRNA], k_forward = rxn_k['re0000000619_k1']),
    Reaction.from_massaction([RS30S_mRNA],[RS30S,mRNA], k_forward= rxn_k['re0000000620_k1']),

    Reaction.from_massaction([RS30S_mRNA,IF2_GTP],[RS30S_IF2_GTP_mRNA], k_forward= rxn_k['re0000000625_k1']),
    Reaction.from_massaction([RS30S_IF2_GTP_mRNA],[IF2_GTP,RS30S_mRNA], k_forward= rxn_k['re0000000626_k1']),
    Reaction.from_massaction([mRNA,RS30S_IF2_GTP],[RS30S_IF2_GTP_mRNA], k_forward = rxn_k['re0000000631_k1']),
    Reaction.from_massaction([RS30S_IF2_GTP_mRNA],[RS30S_IF2_GTP,mRNA], k_forward= rxn_k['re0000000632_k1']),
    Reaction.from_massaction([RS30S_mRNA,IF3],[RS30S_IF3_mRNA], k_forward= rxn_k['re0000000635_k1']),
    Reaction.from_massaction([RS30S_IF3_mRNA],[IF3,RS30S_mRNA], k_forward= rxn_k['re0000000636_k1']),
    Reaction.from_massaction([RS30S_IF2_GTP_mRNA,IF3],[RS30S_IF3_IF2_GTP_mRNA], k_forward= rxn_k['re0000000639_k1']),
    Reaction.from_massaction([RS30S_IF3_IF2_GTP_mRNA],[IF3,RS30S_IF2_GTP_mRNA], k_forward= rxn_k['re0000000640_k1']),
    Reaction.from_massaction([RS30S_IF1,IF2_GTP],[RS30S_IF1_IF2_GTP], k_forward = rxn_k['re0000000643_k1']),
    Reaction.from_massaction([RS30S_IF1_IF2_GTP],[IF2_GTP,RS30S_IF1], k_forward = rxn_k['re0000000644_k1']),

    Reaction.from_massaction([IF1,RS30S_IF2_GTP],[RS30S_IF1_IF2_GTP], k_forward = rxn_k['re0000000649_k1']),
    Reaction.from_massaction([RS30S_IF1_IF2_GTP],[IF1,RS30S_IF2_GTP], k_forward = rxn_k['re0000000650_k1']),
    Reaction.from_massaction([RS30S_IF1_IF2_GTP,IF3],[RS30S_IF1_IF3_IF2_GTP], k_forward = rxn_k['re0000000653_k1']),
    Reaction.from_massaction([RS30S_IF1_IF3_IF2_GTP],[IF3,RS30S_IF1_IF2_GTP], k_forward = rxn_k['re0000000654_k1']),
    Reaction.from_massaction([RS30S_IF1_mRNA,IF2_GTP],[RS30S_IF1_IF2_GTP_mRNA], k_forward= rxn_k['re0000000657_k1']),
    Reaction.from_massaction([RS30S_IF1_IF2_GTP_mRNA],[RS30S_IF1_mRNA,IF2_GTP], k_forward = rxn_k['re0000000658_k1']),
    Reaction.from_massaction([RS30S_IF1_mRNA,IF3],[RS30S_IF1_IF3_mRNA], k_forward= rxn_k['re0000000667_k1']),
    Reaction.from_massaction([RS30S_IF1_IF3_mRNA],[IF3,RS30S_IF1_mRNA], k_forward = rxn_k['re0000000668_k1']),
    Reaction.from_massaction([RS30S_IF1_IF2_GTP_mRNA,IF3],[RS30S_IF1_IF3_IF2_GTP_mRNA], k_forward = rxn_k['re0000000671_k1']),
    Reaction.from_massaction([RS30S_IF1_IF3_IF2_GTP_mRNA],[IF3,RS30S_IF1_IF2_GTP_mRNA], k_forward = rxn_k['re0000000672_k1']),

    Reaction.from_massaction([RS30S_IF1,mRNA],[RS30S_IF1_mRNA], k_forward = rxn_k['re0000000675_k1']),
    Reaction.from_massaction([RS30S_IF1_mRNA],[RS30S_IF1,mRNA], k_forward= rxn_k['re0000000676_k1']),
    Reaction.from_massaction([RS30S_mRNA,IF1],[RS30S_IF1_mRNA], k_forward= rxn_k['re0000000677_k1']),
    Reaction.from_massaction([RS30S_IF1_mRNA],[RS30S_mRNA,IF1], k_forward= rxn_k['re0000000678_k1']),
    Reaction.from_massaction([RS30S_IF2_GTP_mRNA,IF1],[RS30S_IF1_IF2_GTP_mRNA], k_forward= rxn_k['re0000000679_k1']),
    Reaction.from_massaction([RS30S_IF1_IF2_GTP_mRNA],[RS30S_IF2_GTP_mRNA,IF1], k_forward = rxn_k['re0000000680_k1']),
    Reaction.from_massaction([RS30S_IF1_IF2_GTP,mRNA],[RS30S_IF1_IF2_GTP_mRNA], k_forward = rxn_k['re0000000681_k1']),
    Reaction.from_massaction([RS30S_IF1_IF2_GTP_mRNA],[RS30S_IF1_IF2_GTP,mRNA], k_forward = rxn_k['re0000000682_k1']),
    # Reaction.from_massaction([RS30S_IF2_GTP],[RS30S_degraded,IF2,GTP], k_forward= rxn_k['re0000000689_k1']),
    # Reaction.from_massaction([RS30S_IF2_GTP],[IF2_degraded,GTP,RS30S], k_forward= rxn_k['re0000000690_k1']),

    # Reaction.from_massaction([RS30S_mRNA],[RS30S_degraded,mRNA], k_forward= rxn_k['re0000000693_k1']),
    # Reaction.from_massaction([RS30S_IF2_GTP_mRNA],[RS30S_degraded,IF2,GTP,mRNA], k_forward= rxn_k['re0000000695_k1']),
    # Reaction.from_massaction([RS30S_IF2_GTP_mRNA],[IF2_degraded,mRNA,RS30S,GTP], k_forward= rxn_k['re0000000696_k1']),
    # Reaction.from_massaction([RS30S_IF1_IF2_GTP],[RS30S_degraded,IF2,GTP,IF1], k_forward = rxn_k['re0000000699_k1']),
    # Reaction.from_massaction([RS30S_IF1_IF2_GTP],[IF2_degraded,GTP,RS30S,IF1], k_forward = rxn_k['re0000000700_k1']),
    # Reaction.from_massaction([RS30S_IF1_IF2_GTP],[IF1_degraded,RS30S,IF2,GTP], k_forward = rxn_k['re0000000701_k1']),
    # Reaction.from_massaction([RS30S_IF1_mRNA],[RS30S_degraded,IF1,mRNA], k_forward= rxn_k['re0000000705_k1']),
    # Reaction.from_massaction([RS30S_IF1_mRNA],[IF1_degraded,mRNA,RS30S], k_forward= rxn_k['re0000000706_k1']),
    # Reaction.from_massaction([RS30S_IF1_IF2_GTP_mRNA],[RS30S_degraded,IF2,GTP,IF1,mRNA], k_forward = rxn_k['re0000000709_k1']),
    # Reaction.from_massaction([RS30S_IF1_IF2_GTP_mRNA],[IF2_degraded,mRNA,RS30S,IF1,GTP], k_forward = rxn_k['re0000000710_k1']),

    # Reaction.from_massaction([RS30S_IF1_IF2_GTP_mRNA],[IF1_degraded,RS30S,mRNA,IF2,GTP], k_forward = rxn_k['re0000000711_k1']),
    # Reaction.from_massaction([ATP],[ADP,PO4], k_forward = rxn_k['re0000000784_k1']),
    # Reaction.from_massaction([ADP],[AMP,PO4], k_forward = rxn_k['re0000000785_k1']),!!!!!!!!!!REALLY???
    # Reaction.from_massaction([ATP],[AMP,PPi], k_forward = rxn_k['re0000000786_k1']),
    # Reaction.from_massaction([GTP],[GDP,PO4], k_forward = rxn_k['re0000000787_k1']),
    # Reaction.from_massaction([GDP],[GMP,PO4], k_forward = rxn_k['re0000000788_k1']),
    # Reaction.from_massaction([GTP],[GMP,PPi], k_forward = rxn_k['re0000000789_k1']),
    # Reaction.from_massaction([PO4,ADP],[ATP], k_forward = rxn_k['re0000000790_k1']),
    # Reaction.from_massaction([AMP,PO4],[ADP], k_forward = rxn_k['re0000000791_k1']),
    # Reaction.from_massaction([AMP,PPi],[ATP], k_forward = rxn_k['re0000000792_k1']),

    # Reaction.from_massaction([PO4,GDP],[GTP], k_forward = rxn_k['re0000000793_k1']),
    # Reaction.from_massaction([GMP,PO4],[GDP], k_forward = rxn_k['re0000000794_k1']),
    # Reaction.from_massaction([GMP,PPi],[GTP], k_forward = rxn_k['re0000000795_k1']),
    # Reaction.from_massaction([RF1],[RF1_degraded], k_forward = rxn_k['re0000000803_k1']),
    # Reaction.from_massaction([RF2],[RF2_degraded], k_forward = rxn_k['re0000000818_k1']),
    # Reaction.from_massaction([RF3],[RF3_degraded], k_forward = rxn_k['re0000000824_k1']),
    Reaction.from_massaction([RF3_GDP],[GDP,RF3], k_forward = rxn_k['re0000000825_k1']),
    Reaction.from_massaction([RF3,GDP],[RF3_GDP], k_forward = rxn_k['re0000000826_k1']),
    Reaction.from_massaction([RF3,GTP],[RF3_GTP], k_forward = rxn_k['re0000000827_k1']),
    Reaction.from_massaction([RF3_GTP],[RF3,GTP], k_forward = rxn_k['re0000000828_k1']),

    # Reaction.from_massaction([RF3_GDP],[RF3_degraded,GDP], k_forward = rxn_k['re0000000848_k1']),
    # Reaction.from_massaction([RF3_GTP],[RF3_degraded,GTP], k_forward = rxn_k['re0000000849_k1']),
    # Reaction.from_massaction([RRF],[RRF_degraded], k_forward = rxn_k['re0000000894_k1']),
    Reaction.from_massaction([RS50S_RRF],[RS50S,RRF], k_forward= rxn_k['re0000000918_k1']),
    Reaction.from_massaction([RS50S_RRF_EFG_GDP],[RS50S_RRF,EFG_GDP], k_forward= rxn_k['re0000000922_k1']),
    Reaction.from_massaction([RS50S_RRF_EFG_GDP],[RS50S_EFG_GDP,RRF], k_forward= rxn_k['re0000000923_k1']),
    # Reaction.from_massaction([RS50S_RRF],[RS50S_degraded,RRF], k_forward= rxn_k['re0000000925_k1']),
    # Reaction.from_massaction([RS50S_RRF],[RRF_degraded,RS50S], k_forward = rxn_k['re0000000928_k1']),
    # Reaction.from_massaction([RS50S_RRF_EFG_GDP],[RRF_degraded,RS50S,EFG,GDP], k_forward= rxn_k['re0000000935_k1']),
    # Reaction.from_massaction([RS50S_RRF_EFG_GDP],[RS50S_degraded,EFG,RRF,GDP], k_forward= rxn_k['re0000000936_k1']),
    # Reaction.from_massaction([RS50S_RRF_EFG_GDP],[EFG_degraded,RS50S,RRF,GDP], k_forward= rxn_k['re0000000937_k1']),

    # Reaction.from_massaction([RRF,RS50S_EFG_GDP],[RS50S_RRF_EFG_GDP], k_forward = rxn_k['re0000000964_k1']),
    # Reaction.from_massaction([EFG_GDP,RS50S_RRF],[RS50S_RRF_EFG_GDP], k_forward= rxn_k['re0000000966_k1']),
    # Reaction.from_massaction([RRF,RS50S],[RS50S_RRF], k_forward = rxn_k['re0000000968_k1']),
    ]
```

#### Met Species and Reaction¶

In [9]:

```
#Create al the fMet and needed MetRS Species
fMettRNAfMetCAU = Species('fMettRNAfMetCAU')
fMettRNAfMetCAU_degraded = Species('fMettRNAfMetCAU_degraded')
tRNAfMetCAU = Species('tRNAfMetCAU')
tRNAfMetCAU_degraded = Species('tRNAfMetCAU_degraded')
EFTu_GTP_MettRNAfMetCAU = Species('EFTu_GTP_MettRNAfMetCAU')

Met = Species('Met')
MetAMP = Species('MetAMP')
MetRS = Species('MetRS')

MetRS_AMP = Species('MetRS_AMP')
MetRS_AMP_MettRNAfMetCAU = Species('MetRS_AMP_MettRNAfMetCAU')
MetRS_ATP = Species('MetRS_ATP')
MetRS_ATP_tRNAfMetCAU = Species('MetRS_ATP_tRNAfMetCAU')
MetRS_degraded = Species('MetRS_degraded')
MetRS_Met = Species('MetRS_Met')
MetRS_Met_ATP = Species('MetRS_Met_ATP')
MetRS_Met_ATP_tRNAfMetCAU = Species('MetRS_Met_ATP_tRNAfMetCAU')
MetRS_Met_tRNAfMetCAU = Species('MetRS_Met_tRNAfMetCAU')
MetRS_MetAMP = Species('MetRS_MetAMP')
MetRS_MetAMP_PPi = Species('MetRS_MetAMP_PPi')
MetRS_MetAMP_PPi_tRNAfMetCAU = Species('MetRS_MetAMP_PPi_tRNAfMetCAU')
MetRS_MetAMP_tRNAfMetCAU = Species('MetRS_MetAMP_tRNAfMetCAU')
MetRS_MettRNAfMetCAU=Species('MetRS_MettRNAfMetCAU')
MetRS_tRNAfMetCAU = Species('MetRS_tRNAfMetCAU')

MettRNAfMetCAU = Species('MettRNAfMetCAU')
MettRNAfMetCAU_degraded = Species('MettRNAfMetCAU_degraded')
MTF_FD_MettRNAfMetCAU = Species('MTF_FD_MettRNAfMetCAU')
MTF_fMettRNAfMetCAU = Species('MTF_fMettRNAfMetCAU')
MTF_MettRNAfMetCAU = Species('MTF_MettRNAfMetCAU')
MTF_THF_fMettRNAfMetCAU = Species('MTF_THF_fMettRNAfMetCAU')

RS30S_fMettRNAfMetCAU_mRNA = Species('RS30S_fMettRNAfMetCAU_mRNA')
RS30S_IF1_fMettRNAfMetCAU_mRNA = Species('RS30S_IF1_fMettRNAfMetCAU_mRNA')
RS30S_IF1_IF2_GTP_fMettRNAfMetCAU = Species('RS30S_IF1_IF2_GTP_fMettRNAfMetCAU')
RS30S_IF1_IF2_GTP_fMettRNAfMetCAU_mRNA = Species('RS30S_IF1_IF2_GTP_fMettRNAfMetCAU_mRNA')
RS30S_IF1_IF3_fMettRNAfMetCAU_mRNA = Species('RS30S_IF1_IF3_fMettRNAfMetCAU_mRNA')
RS30S_IF1_IF3_IF2_GTP_fMettRNAfMetCAU = Species('RS30S_IF1_IF3_IF2_GTP_fMettRNAfMetCAU')
RS30S_IF1_IF3_IF2_GTP_fMettRNAfMetCAU_mRNA = Species('RS30S_IF1_IF3_IF2_GTP_fMettRNAfMetCAU_mRNA')
RS30S_IF2_GTP_fMettRNAfMetCAU = Species('RS30S_IF2_GTP_fMettRNAfMetCAU')
RS30S_IF2_GTP_fMettRNAfMetCAU_mRNA = Species('RS30S_IF2_GTP_fMettRNAfMetCAU_mRNA')
RS30S_IF3_fMettRNAfMetCAU_mRNA = Species('RS30S_IF3_fMettRNAfMetCAU_mRNA')
RS30S_IF3_IF2_GTP_fMettRNAfMetCAU = Species('RS30S_IF3_IF2_GTP_fMettRNAfMetCAU')
RS30S_IF3_IF2_GTP_fMettRNAfMetCAU_mRNA = Species('RS30S_IF3_IF2_GTP_fMettRNAfMetCAU_mRNA')

RS70S_IF1_fMettRNAfMetCAU_mRNA = Species('RS70S_IF1_fMettRNAfMetCAU_mRNA')
RS70S_IF1_IF2_GDP_fMettRNAfMetCAU_mRNA = Species('RS70S_IF1_IF2_GDP_fMettRNAfMetCAU_mRNA')
RS70S_IF1_IF3_fMettRNAfMetCAU_mRNA = Species('RS70S_IF1_IF3_fMettRNAfMetCAU_mRNA')
RS70S_IF1_IF3_IF2_GDP_fMettRNAfMetCAU_mRNA = Species('RS70S_IF1_IF3_IF2_GDP_fMettRNAfMetCAU_mRNA')
RS70S_IF1_IF3_IF2_GDP_PO4_fMettRNAfMetCAU_mRNA = Species('RS70S_IF1_IF3_IF2_GDP_PO4_fMettRNAfMetCAU_mRNA')
RS70S_IF1_IF3_IF2_GTP_fMettRNAfMetCAU_mRNA = Species('RS70S_IF1_IF3_IF2_GTP_fMettRNAfMetCAU_mRNA')
RS70S_IF2_GDP_fMettRNAfMetCAU_mRNA = Species('RS70S_IF2_GDP_fMettRNAfMetCAU_mRNA')
RS70S_IF3_fMettRNAfMetCAU_mRNA = Species('RS70S_IF3_fMettRNAfMetCAU_mRNA')
RS70S_IF3_IF2_GDP_fMettRNAfMetCAU_mRNA = Species('RS70S_IF3_IF2_GDP_fMettRNAfMetCAU_mRNA')
RS70S_IF3_IF2_GDP_PO4_fMettRNAfMetCAU_mRNA = Species('RS70S_IF3_IF2_GDP_PO4_fMettRNAfMetCAU_mRNA')
RS70S_IF3_IF2_GTP_fMettRNAfMetCAU_mRNA = Species('RS70S_IF3_IF2_GTP_fMettRNAfMetCAU_mRNA')

#######################################################################################################################

#A list of Species in relation to fMet
list_species_fmet=[
    fMettRNAfMetCAU, fMettRNAfMetCAU_degraded, Met, MetAMP, MetRS,tRNAfMetCAU, tRNAfMetCAU_degraded, EFTu_GTP_MettRNAfMetCAU,
    
    MetRS_AMP, MetRS_AMP_MettRNAfMetCAU, MetRS_ATP, MetRS_ATP_tRNAfMetCAU, MetRS_degraded,
    MetRS_Met, MetRS_Met_ATP, MetRS_Met_ATP_tRNAfMetCAU, MetRS_Met_tRNAfMetCAU, MetRS_MetAMP,
    MetRS_MetAMP_PPi, MetRS_MetAMP_PPi_tRNAfMetCAU, MetRS_MetAMP_tRNAfMetCAU, MetRS_MettRNAfMetCAU, MetRS_tRNAfMetCAU,
    MettRNAfMetCAU, MettRNAfMetCAU_degraded, MTF_FD_MettRNAfMetCAU, MTF_fMettRNAfMetCAU, MTF_MettRNAfMetCAU, MTF_THF_fMettRNAfMetCAU,
    
    RS30S_fMettRNAfMetCAU_mRNA, RS30S_IF1_fMettRNAfMetCAU_mRNA, RS30S_IF1_IF2_GTP_fMettRNAfMetCAU, 
    RS30S_IF1_IF2_GTP_fMettRNAfMetCAU_mRNA, RS30S_IF1_IF3_fMettRNAfMetCAU_mRNA, RS30S_IF1_IF3_IF2_GTP_fMettRNAfMetCAU, 
    RS30S_IF1_IF3_IF2_GTP_fMettRNAfMetCAU_mRNA, RS30S_IF2_GTP_fMettRNAfMetCAU, RS30S_IF2_GTP_fMettRNAfMetCAU_mRNA, 
    RS30S_IF3_fMettRNAfMetCAU_mRNA, RS30S_IF3_IF2_GTP_fMettRNAfMetCAU, RS30S_IF3_IF2_GTP_fMettRNAfMetCAU_mRNA,

    RS70S_IF1_fMettRNAfMetCAU_mRNA, RS70S_IF1_IF2_GDP_fMettRNAfMetCAU_mRNA, RS70S_IF1_IF3_fMettRNAfMetCAU_mRNA,
    RS70S_IF1_IF3_IF2_GDP_fMettRNAfMetCAU_mRNA, RS70S_IF1_IF3_IF2_GDP_PO4_fMettRNAfMetCAU_mRNA, RS70S_IF1_IF3_IF2_GTP_fMettRNAfMetCAU_mRNA,
    RS70S_IF2_GDP_fMettRNAfMetCAU_mRNA, RS70S_IF3_fMettRNAfMetCAU_mRNA, RS70S_IF3_IF2_GDP_fMettRNAfMetCAU_mRNA,
    RS70S_IF3_IF2_GDP_PO4_fMettRNAfMetCAU_mRNA, RS70S_IF3_IF2_GTP_fMettRNAfMetCAU_mRNA]

#######################################################################################################################

#fMet Reactions 
list_reaction_fmet=[
    # Reaction.from_massaction([fMettRNAfMetCAU],[fMettRNAfMetCAU_degraded], k_forward= rxn_k['re0000000005_k1']),
    # Reaction.from_massaction([tRNAfMetCAU],[tRNAfMetCAU_degraded], k_forward= rxn_k['re0000000006_k1']),
    Reaction.from_massaction([MetRS,Met],[MetRS_Met], k_forward= rxn_k['re0000000151_k1']),
    Reaction.from_massaction([MetRS_MetAMP_PPi],[MetRS_MetAMP,PPi], k_forward= rxn_k['re0000000152_k1']),
    # Reaction.from_massaction([MetRS],[MetRS_degraded], k_forward= rxn_k['re0000000153_k1']),
    # Reaction.from_massaction([MetRS_Met_ATP],[MetRS_degraded,Met,ATP], k_forward= rxn_k['re0000000154_k1']),
    # Reaction.from_massaction([MetRS_MetAMP],[MetRS_degraded,MetAMP], k_forward= rxn_k['re0000000155_k1']),
    Reaction.from_massaction([MetRS_Met],[MetRS,Met], k_forward= rxn_k['re0000000156_k1']),
    Reaction.from_massaction([MetRS,ATP],[MetRS_ATP], k_forward= rxn_k['re0000000157_k1']),
    Reaction.from_massaction([MetRS_ATP],[MetRS,ATP], k_forward= rxn_k['re0000000158_k1']),

    Reaction.from_massaction([MetRS_ATP,Met],[MetRS_Met_ATP], k_forward= rxn_k['re0000000159_k1']),
    Reaction.from_massaction([MetRS_Met_ATP],[MetRS_ATP,Met], k_forward= rxn_k['re0000000160_k1']),
    Reaction.from_massaction([MetRS_Met,ATP],[MetRS_Met_ATP], k_forward= rxn_k['re0000000161_k1']),
    Reaction.from_massaction([MetRS_Met_ATP],[MetRS_Met,ATP], k_forward= rxn_k['re0000000162_k1']),
    # Reaction.from_massaction([MetRS_ATP],[MetRS_degraded,ATP], k_forward= rxn_k['re0000000163_k1']),
    # Reaction.from_massaction([MetRS_Met],[MetRS_degraded,Met], k_forward= rxn_k['re0000000164_k1']),
    Reaction.from_massaction([MetRS_Met_ATP],[MetRS_MetAMP_PPi], k_forward= rxn_k['re0000000165_k1']),
    Reaction.from_massaction([MetRS_MetAMP_PPi],[MetRS_Met_ATP], k_forward= rxn_k['re0000000166_k1']),
    # Reaction.from_massaction([MetRS_MetAMP_PPi],[MetRS_degraded,MetAMP,PPi], k_forward= rxn_k['re0000000167_k1']),
    # Reaction.from_massaction([MetAMP],[Met,AMP], k_forward= rxn_k['re0000000168_k1']),

    # Reaction.from_massaction([MetRS_AMP],[MetRS_degraded,AMP], k_forward= rxn_k['re0000000169_k1']),
    Reaction.from_massaction([MetRS_AMP],[MetRS,AMP], k_forward= rxn_k['re0000000170_k1']),
    # Reaction.from_massaction([MetRS,AMP],[MetRS_AMP], k_forward= rxn_k['re0000000171_k1']),
    Reaction.from_massaction([MetRS_MetAMP],[MetRS,MetAMP], k_forward= rxn_k['re0000000172_k1']),
    Reaction.from_massaction([MetRS,MetAMP],[MetRS_MetAMP], k_forward= rxn_k['re0000000173_k1']),
    # Reaction.from_massaction([Met,AMP],[MetAMP], k_forward= rxn_k['re0000000174_k1']),
    # Reaction.from_massaction([MetRS_MetAMP,PPi],[MetRS_MetAMP_PPi], k_forward= rxn_k['re0000000175_k1']),
    # Reaction.from_massaction([MettRNAfMetCAU],[tRNAfMetCAU,Met], k_forward= rxn_k['re0000000218_k1']),
    # Reaction.from_massaction([MetRS_MetAMP_tRNAfMetCAU],[MetRS_degraded,MetAMP,tRNAfMetCAU], k_forward= rxn_k['re0000000219_k1']),
    Reaction.from_massaction([MetRS_MetAMP_tRNAfMetCAU],[MetRS_AMP_MettRNAfMetCAU], k_forward= rxn_k['re0000000220_k1']),

    # Reaction.from_massaction([MetRS_AMP_MettRNAfMetCAU],[MetRS_MetAMP_tRNAfMetCAU], k_forward= rxn_k['re0000000221_k1']),
    Reaction.from_massaction([MetRS_AMP_MettRNAfMetCAU],[MetRS_MettRNAfMetCAU,AMP], k_forward= rxn_k['re0000000222_k1']),
    # Reaction.from_massaction([MetRS_MettRNAfMetCAU,AMP],[MetRS_AMP_MettRNAfMetCAU], k_forward= rxn_k['re0000000223_k1']),
    Reaction.from_massaction([MetRS_AMP_MettRNAfMetCAU],[MettRNAfMetCAU,MetRS_AMP], k_forward= rxn_k['re0000000224_k1']),
    Reaction.from_massaction([MetRS_AMP,MettRNAfMetCAU],[MetRS_AMP_MettRNAfMetCAU], k_forward= rxn_k['re0000000225_k1']),
    Reaction.from_massaction([MetRS_MettRNAfMetCAU],[MetRS,MettRNAfMetCAU], k_forward= rxn_k['re0000000226_k1']),
    Reaction.from_massaction([MetRS,MettRNAfMetCAU],[MetRS_MettRNAfMetCAU], k_forward= rxn_k['re0000000227_k1']),
    # Reaction.from_massaction([MetRS_AMP_MettRNAfMetCAU],[MetRS_degraded,AMP,MettRNAfMetCAU], k_forward= rxn_k['re0000000228_k1']),
    # Reaction.from_massaction([MetRS_MettRNAfMetCAU],[MetRS_degraded,MettRNAfMetCAU], k_forward= rxn_k['re0000000229_k1']),
    Reaction.from_massaction([MetRS_tRNAfMetCAU,Met],[MetRS_Met_tRNAfMetCAU], k_forward= rxn_k['re0000000230_k1']),

    Reaction.from_massaction([MetRS_MetAMP_PPi_tRNAfMetCAU],[MetRS_MetAMP_tRNAfMetCAU,PPi], k_forward= rxn_k['re0000000231_k1']),
    Reaction.from_massaction([MetRS_Met_tRNAfMetCAU],[MetRS_tRNAfMetCAU,Met], k_forward= rxn_k['re0000000232_k1']),
    Reaction.from_massaction([MetRS_tRNAfMetCAU,ATP],[MetRS_ATP_tRNAfMetCAU], k_forward= rxn_k['re0000000233_k1']),
    Reaction.from_massaction([MetRS_ATP_tRNAfMetCAU],[MetRS_tRNAfMetCAU,ATP], k_forward= rxn_k['re0000000234_k1']),
    Reaction.from_massaction([MetRS_ATP_tRNAfMetCAU,Met],[MetRS_Met_ATP_tRNAfMetCAU], k_forward= rxn_k['re0000000235_k1']),
    Reaction.from_massaction([MetRS_Met_ATP_tRNAfMetCAU],[MetRS_ATP_tRNAfMetCAU,Met], k_forward= rxn_k['re0000000236_k1']),
    Reaction.from_massaction([MetRS_Met_tRNAfMetCAU,ATP],[MetRS_Met_ATP_tRNAfMetCAU], k_forward= rxn_k['re0000000237_k1']),
    Reaction.from_massaction([MetRS_Met_ATP_tRNAfMetCAU],[MetRS_Met_tRNAfMetCAU,ATP], k_forward= rxn_k['re0000000238_k1']),
    Reaction.from_massaction([MetRS_Met_ATP_tRNAfMetCAU],[MetRS_MetAMP_PPi_tRNAfMetCAU], k_forward= rxn_k['re0000000239_k1']),
    # Reaction.from_massaction([MetRS_MetAMP_PPi_tRNAfMetCAU],[MetRS_Met_ATP_tRNAfMetCAU], k_forward= rxn_k['re0000000240_k1']),

    Reaction.from_massaction([MetRS,tRNAfMetCAU],[MetRS_tRNAfMetCAU], k_forward= rxn_k['re0000000241_k1']),
    Reaction.from_massaction([MetRS_Met,tRNAfMetCAU],[MetRS_Met_tRNAfMetCAU], k_forward= rxn_k['re0000000242_k1']),
    Reaction.from_massaction([MetRS_tRNAfMetCAU],[MetRS,tRNAfMetCAU], k_forward= rxn_k['re0000000243_k1']),
    Reaction.from_massaction([MetRS_Met_tRNAfMetCAU],[MetRS_Met,tRNAfMetCAU], k_forward= rxn_k['re0000000244_k1']),
    Reaction.from_massaction([MetRS_ATP,tRNAfMetCAU],[MetRS_ATP_tRNAfMetCAU], k_forward= rxn_k['re0000000245_k1']),
    Reaction.from_massaction([MetRS_ATP_tRNAfMetCAU],[MetRS_ATP,tRNAfMetCAU], k_forward= rxn_k['re0000000246_k1']),
    Reaction.from_massaction([MetRS_Met_ATP,tRNAfMetCAU],[MetRS_Met_ATP_tRNAfMetCAU], k_forward= rxn_k['re0000000247_k1']),
    Reaction.from_massaction([MetRS_Met_ATP_tRNAfMetCAU],[MetRS_Met_ATP,tRNAfMetCAU], k_forward= rxn_k['re0000000248_k1']),
    Reaction.from_massaction([MetRS_MetAMP_PPi,tRNAfMetCAU],[MetRS_MetAMP_PPi_tRNAfMetCAU], k_forward= rxn_k['re0000000249_k1']),
    Reaction.from_massaction([MetRS_MetAMP_PPi_tRNAfMetCAU],[MetRS_MetAMP_PPi,tRNAfMetCAU], k_forward= rxn_k['re0000000250_k1']),

    Reaction.from_massaction([MetRS_MetAMP,tRNAfMetCAU],[MetRS_MetAMP_tRNAfMetCAU], k_forward= rxn_k['re0000000251_k1']),
    Reaction.from_massaction([MetRS_MetAMP_tRNAfMetCAU],[MetRS_MetAMP,tRNAfMetCAU], k_forward= rxn_k['re0000000252_k1']),
    # Reaction.from_massaction([MetRS_tRNAfMetCAU],[MetRS_degraded,tRNAfMetCAU], k_forward= rxn_k['re0000000253_k1']),
    # Reaction.from_massaction([MetRS_ATP_tRNAfMetCAU],[MetRS_degraded,tRNAfMetCAU,ATP], k_forward= rxn_k['re0000000254_k1']),
    # Reaction.from_massaction([MetRS_Met_tRNAfMetCAU],[MetRS_degraded,tRNAfMetCAU,Met], k_forward= rxn_k['re0000000255_k1']),
    # Reaction.from_massaction([MetRS_Met_ATP_tRNAfMetCAU],[MetRS_degraded,tRNAfMetCAU,ATP,Met], k_forward= rxn_k['re0000000256_k1']),
    # Reaction.from_massaction([MetRS_MetAMP_PPi_tRNAfMetCAU],[MetRS_degraded,tRNAfMetCAU,MetAMP,PPi], k_forward= rxn_k['re0000000257_k1']),
    # Reaction.from_massaction([MettRNAfMetCAU],[MettRNAfMetCAU_degraded], k_forward= rxn_k['re0000000258_k1']),
    # Reaction.from_massaction([tRNAfMetCAU,Met],[MettRNAfMetCAU], k_forward= rxn_k['re0000000259_k1']),
    # Reaction.from_massaction([MetRS_MetAMP_tRNAfMetCAU,PPi],[MetRS_MetAMP_PPi_tRNAfMetCAU], k_forward= rxn_k['re0000000260_k1']),

    Reaction.from_massaction([EFTu_GTP,MettRNAfMetCAU],[EFTu_GTP_MettRNAfMetCAU], k_forward= rxn_k['re0000000288_k1']),
    Reaction.from_massaction([EFTu_GTP_MettRNAfMetCAU],[EFTu_GTP,MettRNAfMetCAU], k_forward= rxn_k['re0000000289_k1']),
    # Reaction.from_massaction([EFTu_GTP_MettRNAfMetCAU],[EFTu_degraded,MettRNAfMetCAU,GTP], k_forward= rxn_k['re0000000290_k1']),
    # Reaction.from_massaction([EFTu_GTP_MettRNAfMetCAU],[MettRNAfMetCAU_degraded,EFTu,GTP], k_forward= rxn_k['re0000000291_k1']),
    # Reaction.from_massaction([fMettRNAfMetCAU],[tRNAfMetCAU,fMet], k_forward= rxn_k['re0000000417_k1']),
    Reaction.from_massaction([MTF,MettRNAfMetCAU],[MTF_MettRNAfMetCAU], k_forward= rxn_k['re0000000420_k1']),
    Reaction.from_massaction([MTF_MettRNAfMetCAU],[MTF,MettRNAfMetCAU], k_forward= rxn_k['re0000000421_k1']),
    Reaction.from_massaction([MTF_FD,MettRNAfMetCAU],[MTF_FD_MettRNAfMetCAU], k_forward= rxn_k['re0000000422_k1']),
    Reaction.from_massaction([MTF_FD_MettRNAfMetCAU],[MettRNAfMetCAU,MTF_FD], k_forward= rxn_k['re0000000423_k1']),
    Reaction.from_massaction([MTF_MettRNAfMetCAU,FD],[MTF_FD_MettRNAfMetCAU], k_forward= rxn_k['re0000000424_k1']),

    Reaction.from_massaction([MTF_FD_MettRNAfMetCAU],[MTF_MettRNAfMetCAU,FD], k_forward= rxn_k['re0000000425_k1']),
    Reaction.from_massaction([MTF_FD_MettRNAfMetCAU],[MTF_THF_fMettRNAfMetCAU], k_forward= rxn_k['re0000000426_k1']),
    # Reaction.from_massaction([MTF_THF_fMettRNAfMetCAU],[MTF_FD_MettRNAfMetCAU], k_forward= rxn_k['re0000000427_k1']),
    Reaction.from_massaction([MTF_THF_fMettRNAfMetCAU],[MTF_THF,fMettRNAfMetCAU], k_forward= rxn_k['re0000000428_k1']),
    # Reaction.from_massaction([MTF_THF,fMettRNAfMetCAU],[MTF_THF_fMettRNAfMetCAU], k_forward= rxn_k['re0000000429_k1']),
    Reaction.from_massaction([MTF_THF_fMettRNAfMetCAU],[MTF_fMettRNAfMetCAU,THF], k_forward= rxn_k['re0000000430_k1']),
    # Reaction.from_massaction([MTF_fMettRNAfMetCAU,THF],[MTF_THF_fMettRNAfMetCAU], k_forward= rxn_k['re0000000431_k1']),
    Reaction.from_massaction([MTF_fMettRNAfMetCAU],[MTF,fMettRNAfMetCAU], k_forward= rxn_k['re0000000432_k1']),
    # Reaction.from_massaction([MTF,fMettRNAfMetCAU],[MTF_fMettRNAfMetCAU], k_forward= rxn_k['re0000000433_k1']),
    # Reaction.from_massaction([MTF_MettRNAfMetCAU],[MTF_degraded,MettRNAfMetCAU], k_forward= rxn_k['re0000000437_k1']),

    # Reaction.from_massaction([MTF_FD_MettRNAfMetCAU],[MTF_degraded,FD,MettRNAfMetCAU], k_forward= rxn_k['re0000000438_k1']),
    # Reaction.from_massaction([MTF_THF_fMettRNAfMetCAU],[MTF_degraded,THF,fMettRNAfMetCAU], k_forward= rxn_k['re0000000439_k1']),
    # Reaction.from_massaction([MTF_fMettRNAfMetCAU],[MTF_degraded,fMettRNAfMetCAU], k_forward= rxn_k['re0000000440_k1']),
    # Reaction.from_massaction([tRNAfMetCAU,fMet],[fMettRNAfMetCAU], k_forward= rxn_k['re0000000443_k1']),
    Reaction.from_massaction([IF2_GTP,fMettRNAfMetCAU],[IF2_GTP_fMettRNAfMetCAU], k_forward= rxn_k['re0000000449_k1']),
    Reaction.from_massaction([IF2_GTP_fMettRNAfMetCAU],[IF2_GTP,fMettRNAfMetCAU], k_forward= rxn_k['re0000000450_k1']),
    # Reaction.from_massaction([IF2_GTP_fMettRNAfMetCAU],[IF2_degraded,fMettRNAfMetCAU,GTP], k_forward= rxn_k['re0000000454_k1']),
    Reaction.from_massaction([RS30S_IF3,IF2_GTP_fMettRNAfMetCAU],[RS30S_IF3_IF2_GTP_fMettRNAfMetCAU], k_forward= rxn_k['re0000000465_k1']),
    Reaction.from_massaction([RS30S_IF3_IF2_GTP_fMettRNAfMetCAU],[RS30S_IF3,IF2_GTP_fMettRNAfMetCAU], k_forward= rxn_k['re0000000466_k1']),
    Reaction.from_massaction([RS30S_IF3_IF2_GTP,fMettRNAfMetCAU],[RS30S_IF3_IF2_GTP_fMettRNAfMetCAU], k_forward= rxn_k['re0000000467_k1']),

    Reaction.from_massaction([RS30S_IF3_IF2_GTP_fMettRNAfMetCAU],[RS30S_IF3_IF2_GTP,fMettRNAfMetCAU], k_forward= rxn_k['re0000000468_k1']),
    Reaction.from_massaction([RS30S_IF3_mRNA,fMettRNAfMetCAU],[RS30S_IF3_fMettRNAfMetCAU_mRNA], k_forward= rxn_k['re0000000473_k1']),
    Reaction.from_massaction([RS30S_IF3_fMettRNAfMetCAU_mRNA],[RS30S_IF3_mRNA,fMettRNAfMetCAU], k_forward= rxn_k['re0000000474_k1']),
    Reaction.from_massaction([RS30S_IF3_mRNA,IF2_GTP_fMettRNAfMetCAU],[RS30S_IF3_IF2_GTP_fMettRNAfMetCAU_mRNA], k_forward= rxn_k['re0000000475_k1']),
    Reaction.from_massaction([RS30S_IF3_IF2_GTP_fMettRNAfMetCAU_mRNA],[RS30S_IF3_mRNA,IF2_GTP_fMettRNAfMetCAU], k_forward= rxn_k['re0000000476_k1']),
    Reaction.from_massaction([RS30S_IF3_fMettRNAfMetCAU_mRNA,IF2_GTP],[RS30S_IF3_IF2_GTP_fMettRNAfMetCAU_mRNA], k_forward= rxn_k['re0000000477_k1']),
    Reaction.from_massaction([RS30S_IF3_IF2_GTP_fMettRNAfMetCAU_mRNA],[RS30S_IF3_fMettRNAfMetCAU_mRNA,IF2_GTP], k_forward= rxn_k['re0000000478_k1']),
    Reaction.from_massaction([RS30S_IF3_IF2_GTP_mRNA,fMettRNAfMetCAU],[RS30S_IF3_IF2_GTP_fMettRNAfMetCAU_mRNA], k_forward= rxn_k['re0000000479_k1']),
    Reaction.from_massaction([RS30S_IF3_IF2_GTP_fMettRNAfMetCAU_mRNA],[RS30S_IF3_IF2_GTP_mRNA,fMettRNAfMetCAU], k_forward= rxn_k['re0000000480_k1']),
    Reaction.from_massaction([RS30S_IF3_IF2_GTP_fMettRNAfMetCAU,mRNA],[RS30S_IF3_IF2_GTP_fMettRNAfMetCAU_mRNA], k_forward= rxn_k['re0000000483_k1']),

    Reaction.from_massaction([RS30S_IF3_IF2_GTP_fMettRNAfMetCAU_mRNA],[RS30S_IF3_IF2_GTP_fMettRNAfMetCAU,mRNA], k_forward= rxn_k['re0000000484_k1']),
    Reaction.from_massaction([RS30S_IF3_IF2_GTP_fMettRNAfMetCAU_mRNA,RS50S],[RS70S_IF3_IF2_GTP_fMettRNAfMetCAU_mRNA], k_forward= rxn_k['re0000000485_k1']),
    Reaction.from_massaction([RS70S_IF3_IF2_GTP_fMettRNAfMetCAU_mRNA],[RS30S_IF3_IF2_GTP_fMettRNAfMetCAU_mRNA,RS50S], k_forward= rxn_k['re0000000486_k1']),
    Reaction.from_massaction([RS30S_IF1_IF3,IF2_GTP_fMettRNAfMetCAU],[RS30S_IF1_IF3_IF2_GTP_fMettRNAfMetCAU], k_forward= rxn_k['re0000000497_k1']),
    Reaction.from_massaction([RS30S_IF1_IF3_IF2_GTP_fMettRNAfMetCAU],[RS30S_IF1_IF3,IF2_GTP_fMettRNAfMetCAU], k_forward= rxn_k['re0000000498_k1']),
    Reaction.from_massaction([RS30S_IF1_IF3_IF2_GTP,fMettRNAfMetCAU],[RS30S_IF1_IF3_IF2_GTP_fMettRNAfMetCAU], k_forward= rxn_k['re0000000499_k1']),
    Reaction.from_massaction([RS30S_IF1_IF3_IF2_GTP_fMettRNAfMetCAU],[RS30S_IF1_IF3_IF2_GTP,fMettRNAfMetCAU], k_forward= rxn_k['re0000000500_k1']),
    Reaction.from_massaction([RS30S_IF3_IF2_GTP_fMettRNAfMetCAU,IF1],[RS30S_IF1_IF3_IF2_GTP_fMettRNAfMetCAU], k_forward= rxn_k['re0000000511_k1']),
    Reaction.from_massaction([RS30S_IF1_IF3_IF2_GTP_fMettRNAfMetCAU],[RS30S_IF3_IF2_GTP_fMettRNAfMetCAU,IF1], k_forward= rxn_k['re0000000512_k1']),
    Reaction.from_massaction([RS30S_IF1_IF3_mRNA,fMettRNAfMetCAU],[RS30S_IF1_IF3_fMettRNAfMetCAU_mRNA], k_forward= rxn_k['re0000000517_k1']),

    Reaction.from_massaction([RS30S_IF1_IF3_fMettRNAfMetCAU_mRNA],[RS30S_IF1_IF3_mRNA,fMettRNAfMetCAU], k_forward= rxn_k['re0000000518_k1']),
    Reaction.from_massaction([RS30S_IF1_IF3_mRNA,IF2_GTP_fMettRNAfMetCAU],[RS30S_IF1_IF3_IF2_GTP_fMettRNAfMetCAU_mRNA], k_forward= rxn_k['re0000000519_k1']),
    Reaction.from_massaction([RS30S_IF1_IF3_IF2_GTP_fMettRNAfMetCAU_mRNA],[RS30S_IF1_IF3_mRNA,IF2_GTP_fMettRNAfMetCAU], k_forward= rxn_k['re0000000520_k1']),
    Reaction.from_massaction([RS30S_IF1_IF3_fMettRNAfMetCAU_mRNA,IF2_GTP],[RS30S_IF1_IF3_IF2_GTP_fMettRNAfMetCAU_mRNA], k_forward= rxn_k['re0000000521_k1']),
    Reaction.from_massaction([RS30S_IF1_IF3_IF2_GTP_fMettRNAfMetCAU_mRNA],[RS30S_IF1_IF3_fMettRNAfMetCAU_mRNA,IF2_GTP], k_forward= rxn_k['re0000000522_k1']),
    Reaction.from_massaction([RS30S_IF1_IF3_IF2_GTP_mRNA,fMettRNAfMetCAU],[RS30S_IF1_IF3_IF2_GTP_fMettRNAfMetCAU_mRNA], k_forward= rxn_k['re0000000523_k1']),
    Reaction.from_massaction([RS30S_IF1_IF3_IF2_GTP_fMettRNAfMetCAU_mRNA],[RS30S_IF1_IF3_IF2_GTP_mRNA,fMettRNAfMetCAU], k_forward= rxn_k['re0000000524_k1']),
    Reaction.from_massaction([RS30S_IF1_IF3_IF2_GTP_fMettRNAfMetCAU,mRNA],[RS30S_IF1_IF3_IF2_GTP_fMettRNAfMetCAU_mRNA], k_forward= rxn_k['re0000000527_k1']),
    Reaction.from_massaction([RS30S_IF1_IF3_IF2_GTP_fMettRNAfMetCAU_mRNA],[RS30S_IF1_IF3_IF2_GTP_fMettRNAfMetCAU,mRNA], k_forward= rxn_k['re0000000528_k1']),

    Reaction.from_massaction([RS30S_IF1_IF3_IF2_GTP_fMettRNAfMetCAU_mRNA,RS50S],[RS70S_IF1_IF3_IF2_GTP_fMettRNAfMetCAU_mRNA], k_forward= rxn_k['re0000000529_k1']),
    Reaction.from_massaction([RS70S_IF1_IF3_IF2_GTP_fMettRNAfMetCAU_mRNA],[RS30S_IF1_IF3_IF2_GTP_fMettRNAfMetCAU_mRNA,RS50S], k_forward= rxn_k['re0000000530_k1']),
    Reaction.from_massaction([RS30S_IF3_fMettRNAfMetCAU_mRNA,IF1],[RS30S_IF1_IF3_fMettRNAfMetCAU_mRNA], k_forward= rxn_k['re0000000533_k1']),
    Reaction.from_massaction([RS30S_IF1_IF3_fMettRNAfMetCAU_mRNA],[RS30S_IF3_fMettRNAfMetCAU_mRNA,IF1], k_forward= rxn_k['re0000000534_k1']),
    Reaction.from_massaction([RS30S_IF3_IF2_GTP_fMettRNAfMetCAU_mRNA,IF1],[RS30S_IF1_IF3_IF2_GTP_fMettRNAfMetCAU_mRNA], k_forward= rxn_k['re0000000537_k1']),
    Reaction.from_massaction([RS30S_IF1_IF3_IF2_GTP_fMettRNAfMetCAU_mRNA],[RS30S_IF3_IF2_GTP_fMettRNAfMetCAU_mRNA,IF1], k_forward= rxn_k['re0000000538_k1']),
    Reaction.from_massaction([RS70S_IF3_IF2_GTP_fMettRNAfMetCAU_mRNA,IF1],[RS70S_IF1_IF3_IF2_GTP_fMettRNAfMetCAU_mRNA], k_forward= rxn_k['re0000000539_k1']),
    Reaction.from_massaction([RS70S_IF1_IF3_IF2_GTP_fMettRNAfMetCAU_mRNA],[RS70S_IF3_IF2_GTP_fMettRNAfMetCAU_mRNA,IF1], k_forward= rxn_k['re0000000540_k1']),
    # Reaction.from_massaction([RS30S_IF3_fMettRNAfMetCAU_mRNA],[IF3_degraded,mRNA,fMettRNAfMetCAU,RS30S], k_forward= rxn_k['re0000000551_k1']),
    # Reaction.from_massaction([RS30S_IF3_fMettRNAfMetCAU_mRNA],[RS30S_degraded,mRNA,IF3,fMettRNAfMetCAU], k_forward= rxn_k['re0000000552_k1']),

    # Reaction.from_massaction([RS30S_IF3_IF2_GTP_fMettRNAfMetCAU],[RS30S_degraded,fMettRNAfMetCAU,GTP,IF3,IF2], k_forward= rxn_k['re0000000565_k1']),
    # Reaction.from_massaction([RS30S_IF3_IF2_GTP_fMettRNAfMetCAU],[IF2_degraded,GTP,fMettRNAfMetCAU,RS30S,IF3], k_forward= rxn_k['re0000000566_k1']),
    # Reaction.from_massaction([RS30S_IF3_IF2_GTP_fMettRNAfMetCAU],[IF3_degraded,GTP,fMettRNAfMetCAU,IF2,RS30S], k_forward= rxn_k['re0000000567_k1']),
    # Reaction.from_massaction([RS30S_IF1_IF3_fMettRNAfMetCAU_mRNA],[IF3_degraded,mRNA,fMettRNAfMetCAU,RS30S,IF1], k_forward= rxn_k['re0000000574_k1']),
    # Reaction.from_massaction([RS30S_IF1_IF3_fMettRNAfMetCAU_mRNA],[RS30S_degraded,mRNA,fMettRNAfMetCAU,IF3,IF1], k_forward= rxn_k['re0000000575_k1']),
    # Reaction.from_massaction([RS30S_IF1_IF3_fMettRNAfMetCAU_mRNA],[IF1_degraded,fMettRNAfMetCAU,RS30S,mRNA,IF3], k_forward= rxn_k['re0000000576_k1']),
    # Reaction.from_massaction([RS30S_IF3_IF2_GTP_fMettRNAfMetCAU_mRNA],[RS30S_degraded,mRNA,fMettRNAfMetCAU,IF3,IF2,GTP], k_forward= rxn_k['re0000000577_k1']),
    # Reaction.from_massaction([RS30S_IF3_IF2_GTP_fMettRNAfMetCAU_mRNA],[IF2_degraded,mRNA,fMettRNAfMetCAU,GTP,RS30S,IF3], k_forward= rxn_k['re0000000578_k1']),
    # Reaction.from_massaction([RS30S_IF3_IF2_GTP_fMettRNAfMetCAU_mRNA],[IF3_degraded,GTP,IF2,fMettRNAfMetCAU,mRNA,RS30S], k_forward= rxn_k['re0000000579_k1']),
    # Reaction.from_massaction([RS30S_IF1_IF3_IF2_GTP_fMettRNAfMetCAU],[IF3_degraded,GTP,fMettRNAfMetCAU,IF2,IF1,RS30S], k_forward= rxn_k['re0000000588_k1']),

    # Reaction.from_massaction([RS30S_IF1_IF3_IF2_GTP_fMettRNAfMetCAU],[IF2_degraded,RS30S,fMettRNAfMetCAU,GTP,IF3,IF1], k_forward= rxn_k['re0000000589_k1']),
    # Reaction.from_massaction([RS30S_IF1_IF3_IF2_GTP_fMettRNAfMetCAU],[IF1_degraded,GTP,IF2,RS30S,IF3,fMettRNAfMetCAU], k_forward= rxn_k['re0000000590_k1']),
    # Reaction.from_massaction([RS30S_IF1_IF3_IF2_GTP_fMettRNAfMetCAU],[RS30S_degraded,GTP,fMettRNAfMetCAU,IF2,IF3,IF1], k_forward= rxn_k['re0000000591_k1']),
    # Reaction.from_massaction([RS30S_IF1_IF3_IF2_GTP_fMettRNAfMetCAU_mRNA],[IF3_degraded,mRNA,fMettRNAfMetCAU,IF2,GTP,IF1,RS30S], k_forward= rxn_k['re0000000596_k1']),
    # Reaction.from_massaction([RS30S_IF1_IF3_IF2_GTP_fMettRNAfMetCAU_mRNA],[RS30S_degraded,mRNA,GTP,fMettRNAfMetCAU,IF2,IF3,IF1], k_forward= rxn_k['re0000000597_k1']),
    # Reaction.from_massaction([RS30S_IF1_IF3_IF2_GTP_fMettRNAfMetCAU_mRNA],[IF2_degraded,RS30S,mRNA,fMettRNAfMetCAU,GTP,IF3,IF1], k_forward= rxn_k['re0000000598_k1']),
    # Reaction.from_massaction([RS30S_IF1_IF3_IF2_GTP_fMettRNAfMetCAU_mRNA],[IF1_degraded,fMettRNAfMetCAU,GTP,mRNA,IF2,RS30S,IF3], k_forward= rxn_k['re0000000599_k1']),
    # Reaction.from_massaction([RS70S_IF3_IF2_GTP_fMettRNAfMetCAU_mRNA],[IF2_degraded,mRNA,GTP,fMettRNAfMetCAU,RS50S,RS30S,IF3], k_forward= rxn_k['re0000000600_k1']),
    # Reaction.from_massaction([RS70S_IF3_IF2_GTP_fMettRNAfMetCAU_mRNA],[RS50S_degraded,IF2,mRNA,fMettRNAfMetCAU,GTP,RS30S,IF3], k_forward= rxn_k['re0000000601_k1']),
    # Reaction.from_massaction([RS70S_IF3_IF2_GTP_fMettRNAfMetCAU_mRNA],[IF3_degraded,GTP,fMettRNAfMetCAU,mRNA,RS30S,RS50S,IF2], k_forward= rxn_k['re0000000602_k1']),

    # Reaction.from_massaction([RS70S_IF3_IF2_GTP_fMettRNAfMetCAU_mRNA],[RS30S_degraded,RS50S,mRNA,fMettRNAfMetCAU,IF2,IF3,GTP], k_forward= rxn_k['re0000000603_k1']),
    # Reaction.from_massaction([RS70S_IF1_IF3_IF2_GTP_fMettRNAfMetCAU_mRNA],[RS30S_degraded,RS50S,GTP,mRNA,fMettRNAfMetCAU,IF2,IF3,IF1], k_forward= rxn_k['re0000000604_k1']),
    # Reaction.from_massaction([RS70S_IF1_IF3_IF2_GTP_fMettRNAfMetCAU_mRNA],[IF2_degraded,mRNA,GTP,fMettRNAfMetCAU,RS50S,RS30S,IF3,IF1], k_forward= rxn_k['re0000000605_k1']),
    # Reaction.from_massaction([RS70S_IF1_IF3_IF2_GTP_fMettRNAfMetCAU_mRNA],[RS50S_degraded,mRNA,IF2,GTP,fMettRNAfMetCAU,RS30S,IF3,IF1], k_forward= rxn_k['re0000000606_k1']),
    # Reaction.from_massaction([RS70S_IF1_IF3_IF2_GTP_fMettRNAfMetCAU_mRNA],[IF1_degraded,mRNA,GTP,fMettRNAfMetCAU,IF2,RS30S,IF3,RS50S], k_forward= rxn_k['re0000000607_k1']),
    # Reaction.from_massaction([RS70S_IF1_IF3_IF2_GTP_fMettRNAfMetCAU_mRNA],[IF3_degraded,RS50S,RS30S,IF1,mRNA,IF2,fMettRNAfMetCAU,GTP], k_forward= rxn_k['re0000000608_k1']),
    Reaction.from_massaction([RS30S,IF2_GTP_fMettRNAfMetCAU],[RS30S_IF2_GTP_fMettRNAfMetCAU], k_forward= rxn_k['re0000000611_k1']),
    Reaction.from_massaction([RS30S_IF2_GTP_fMettRNAfMetCAU],[IF2_GTP_fMettRNAfMetCAU,RS30S], k_forward= rxn_k['re0000000612_k1']),
    Reaction.from_massaction([RS30S_IF2_GTP,fMettRNAfMetCAU],[RS30S_IF2_GTP_fMettRNAfMetCAU], k_forward= rxn_k['re0000000613_k1']),
    Reaction.from_massaction([RS30S_IF2_GTP_fMettRNAfMetCAU],[fMettRNAfMetCAU,RS30S_IF2_GTP], k_forward= rxn_k['re0000000614_k1']),

    Reaction.from_massaction([RS30S_IF2_GTP_fMettRNAfMetCAU,IF3],[RS30S_IF3_IF2_GTP_fMettRNAfMetCAU], k_forward= rxn_k['re0000000617_k1']),
    Reaction.from_massaction([RS30S_IF3_IF2_GTP_fMettRNAfMetCAU],[IF3,RS30S_IF2_GTP_fMettRNAfMetCAU], k_forward= rxn_k['re0000000618_k1']),
    Reaction.from_massaction([RS30S_mRNA,fMettRNAfMetCAU],[RS30S_fMettRNAfMetCAU_mRNA], k_forward= rxn_k['re0000000621_k1']),
    Reaction.from_massaction([RS30S_fMettRNAfMetCAU_mRNA],[fMettRNAfMetCAU,RS30S_mRNA], k_forward= rxn_k['re0000000622_k1']),
    Reaction.from_massaction([RS30S_mRNA,IF2_GTP_fMettRNAfMetCAU],[RS30S_IF2_GTP_fMettRNAfMetCAU_mRNA], k_forward= rxn_k['re0000000623_k1']),
    Reaction.from_massaction([RS30S_IF2_GTP_fMettRNAfMetCAU_mRNA],[IF2_GTP_fMettRNAfMetCAU,RS30S_mRNA], k_forward= rxn_k['re0000000624_k1']),
    Reaction.from_massaction([RS30S_fMettRNAfMetCAU_mRNA,IF2_GTP],[RS30S_IF2_GTP_fMettRNAfMetCAU_mRNA], k_forward= rxn_k['re0000000627_k1']),
    Reaction.from_massaction([RS30S_IF2_GTP_fMettRNAfMetCAU_mRNA],[IF2_GTP,RS30S_fMettRNAfMetCAU_mRNA], k_forward= rxn_k['re0000000628_k1']),
    Reaction.from_massaction([RS30S_IF2_GTP_mRNA,fMettRNAfMetCAU],[RS30S_IF2_GTP_fMettRNAfMetCAU_mRNA], k_forward= rxn_k['re0000000629_k1']),
    Reaction.from_massaction([RS30S_IF2_GTP_fMettRNAfMetCAU_mRNA],[fMettRNAfMetCAU,RS30S_IF2_GTP_mRNA], k_forward= rxn_k['re0000000630_k1']),

    Reaction.from_massaction([mRNA,RS30S_IF2_GTP_fMettRNAfMetCAU],[RS30S_IF2_GTP_fMettRNAfMetCAU_mRNA], k_forward= rxn_k['re0000000633_k1']),
    Reaction.from_massaction([RS30S_IF2_GTP_fMettRNAfMetCAU_mRNA],[RS30S_IF2_GTP_fMettRNAfMetCAU,mRNA], k_forward= rxn_k['re0000000634_k1']),
    Reaction.from_massaction([RS30S_fMettRNAfMetCAU_mRNA,IF3],[RS30S_IF3_fMettRNAfMetCAU_mRNA], k_forward= rxn_k['re0000000637_k1']),
    Reaction.from_massaction([RS30S_IF3_fMettRNAfMetCAU_mRNA],[IF3,RS30S_fMettRNAfMetCAU_mRNA], k_forward= rxn_k['re0000000638_k1']),
    Reaction.from_massaction([RS30S_IF2_GTP_fMettRNAfMetCAU_mRNA,IF3],[RS30S_IF3_IF2_GTP_fMettRNAfMetCAU_mRNA], k_forward= rxn_k['re0000000641_k1']),
    Reaction.from_massaction([RS30S_IF3_IF2_GTP_fMettRNAfMetCAU_mRNA],[IF3,RS30S_IF2_GTP_fMettRNAfMetCAU_mRNA], k_forward= rxn_k['re0000000642_k1']),
    Reaction.from_massaction([RS30S_IF1,IF2_GTP_fMettRNAfMetCAU],[RS30S_IF1_IF2_GTP_fMettRNAfMetCAU], k_forward= rxn_k['re0000000645_k1']),
    Reaction.from_massaction([RS30S_IF1_IF2_GTP_fMettRNAfMetCAU],[IF2_GTP_fMettRNAfMetCAU,RS30S_IF1], k_forward= rxn_k['re0000000646_k1']),
    Reaction.from_massaction([RS30S_IF1_IF2_GTP,fMettRNAfMetCAU],[RS30S_IF1_IF2_GTP_fMettRNAfMetCAU], k_forward= rxn_k['re0000000647_k1']),
    Reaction.from_massaction([RS30S_IF1_IF2_GTP_fMettRNAfMetCAU],[fMettRNAfMetCAU,RS30S_IF1_IF2_GTP], k_forward= rxn_k['re0000000648_k1']),

    Reaction.from_massaction([RS30S_IF2_GTP_fMettRNAfMetCAU,IF1],[RS30S_IF1_IF2_GTP_fMettRNAfMetCAU], k_forward= rxn_k['re0000000651_k1']),
    Reaction.from_massaction([RS30S_IF1_IF2_GTP_fMettRNAfMetCAU],[IF1,RS30S_IF2_GTP_fMettRNAfMetCAU], k_forward= rxn_k['re0000000652_k1']),
    Reaction.from_massaction([RS30S_IF1_IF2_GTP_fMettRNAfMetCAU,IF3],[RS30S_IF1_IF3_IF2_GTP_fMettRNAfMetCAU], k_forward= rxn_k['re0000000655_k1']),
    Reaction.from_massaction([RS30S_IF1_IF3_IF2_GTP_fMettRNAfMetCAU],[IF3,RS30S_IF1_IF2_GTP_fMettRNAfMetCAU], k_forward= rxn_k['re0000000656_k1']),
    Reaction.from_massaction([RS30S_IF1_mRNA,fMettRNAfMetCAU],[RS30S_IF1_fMettRNAfMetCAU_mRNA], k_forward= rxn_k['re0000000659_k1']),
    Reaction.from_massaction([RS30S_IF1_fMettRNAfMetCAU_mRNA],[fMettRNAfMetCAU,RS30S_IF1_mRNA], k_forward= rxn_k['re0000000660_k1']),
    Reaction.from_massaction([RS30S_IF1_IF2_GTP_mRNA,fMettRNAfMetCAU],[RS30S_IF1_IF2_GTP_fMettRNAfMetCAU_mRNA], k_forward= rxn_k['re0000000661_k1']),
    Reaction.from_massaction([RS30S_IF1_IF2_GTP_fMettRNAfMetCAU_mRNA],[fMettRNAfMetCAU,RS30S_IF1_IF2_GTP_mRNA], k_forward= rxn_k['re0000000662_k1']),
    Reaction.from_massaction([RS30S_IF1_fMettRNAfMetCAU_mRNA,IF2_GTP],[RS30S_IF1_IF2_GTP_fMettRNAfMetCAU_mRNA], k_forward= rxn_k['re0000000663_k1']),
    Reaction.from_massaction([RS30S_IF1_IF2_GTP_fMettRNAfMetCAU_mRNA],[IF2_GTP,RS30S_IF1_fMettRNAfMetCAU_mRNA], k_forward= rxn_k['re0000000664_k1']),

    Reaction.from_massaction([RS30S_IF1_mRNA,IF2_GTP_fMettRNAfMetCAU],[RS30S_IF1_IF2_GTP_fMettRNAfMetCAU_mRNA], k_forward= rxn_k['re0000000665_k1']),
    Reaction.from_massaction([RS30S_IF1_IF2_GTP_fMettRNAfMetCAU_mRNA],[IF2_GTP_fMettRNAfMetCAU,RS30S_IF1_mRNA], k_forward= rxn_k['re0000000666_k1']),
    Reaction.from_massaction([RS30S_IF1_fMettRNAfMetCAU_mRNA,IF3],[RS30S_IF1_IF3_fMettRNAfMetCAU_mRNA], k_forward= rxn_k['re0000000669_k1']),
    Reaction.from_massaction([RS30S_IF1_IF3_fMettRNAfMetCAU_mRNA],[IF3,RS30S_IF1_fMettRNAfMetCAU_mRNA], k_forward= rxn_k['re0000000670_k1']),
    Reaction.from_massaction([RS30S_IF1_IF2_GTP_fMettRNAfMetCAU_mRNA,IF3],[RS30S_IF1_IF3_IF2_GTP_fMettRNAfMetCAU_mRNA], k_forward= rxn_k['re0000000673_k1']),
    Reaction.from_massaction([RS30S_IF1_IF3_IF2_GTP_fMettRNAfMetCAU_mRNA],[IF3,RS30S_IF1_IF2_GTP_fMettRNAfMetCAU_mRNA], k_forward= rxn_k['re0000000674_k1']),
    Reaction.from_massaction([RS30S_fMettRNAfMetCAU_mRNA,IF1],[RS30S_IF1_fMettRNAfMetCAU_mRNA], k_forward= rxn_k['re0000000683_k1']),
    Reaction.from_massaction([RS30S_IF1_fMettRNAfMetCAU_mRNA],[RS30S_fMettRNAfMetCAU_mRNA,IF1], k_forward= rxn_k['re0000000684_k1']),
    Reaction.from_massaction([IF1,RS30S_IF2_GTP_fMettRNAfMetCAU_mRNA],[RS30S_IF1_IF2_GTP_fMettRNAfMetCAU_mRNA], k_forward= rxn_k['re0000000685_k1']),
    Reaction.from_massaction([RS30S_IF1_IF2_GTP_fMettRNAfMetCAU_mRNA],[RS30S_IF2_GTP_fMettRNAfMetCAU_mRNA,IF1], k_forward= rxn_k['re0000000686_k1']),

    Reaction.from_massaction([RS30S_IF1_IF2_GTP_fMettRNAfMetCAU,mRNA],[RS30S_IF1_IF2_GTP_fMettRNAfMetCAU_mRNA], k_forward= rxn_k['re0000000687_k1']),
    Reaction.from_massaction([RS30S_IF1_IF2_GTP_fMettRNAfMetCAU_mRNA],[RS30S_IF1_IF2_GTP_fMettRNAfMetCAU,mRNA], k_forward= rxn_k['re0000000688_k1']),
    # Reaction.from_massaction([RS30S_IF2_GTP_fMettRNAfMetCAU],[RS30S_degraded,IF2,GTP,fMettRNAfMetCAU], k_forward= rxn_k['re0000000691_k1']),
    # Reaction.from_massaction([RS30S_IF2_GTP_fMettRNAfMetCAU],[IF2_degraded,fMettRNAfMetCAU,RS30S,GTP], k_forward= rxn_k['re0000000692_k1']),
    # Reaction.from_massaction([RS30S_fMettRNAfMetCAU_mRNA],[RS30S_degraded,fMettRNAfMetCAU,mRNA], k_forward= rxn_k['re0000000694_k1']),
    # Reaction.from_massaction([RS30S_IF2_GTP_fMettRNAfMetCAU_mRNA],[RS30S_degraded,IF2,GTP,fMettRNAfMetCAU,mRNA], k_forward= rxn_k['re0000000697_k1']),
    # Reaction.from_massaction([RS30S_IF2_GTP_fMettRNAfMetCAU_mRNA],[IF2_degraded,fMettRNAfMetCAU,mRNA,RS30S,GTP], k_forward= rxn_k['re0000000698_k1']),
    # Reaction.from_massaction([RS30S_IF1_IF2_GTP_fMettRNAfMetCAU],[RS30S_degraded,IF2,GTP,IF1,fMettRNAfMetCAU], k_forward= rxn_k['re0000000702_k1']),
    # Reaction.from_massaction([RS30S_IF1_IF2_GTP_fMettRNAfMetCAU],[IF2_degraded,GTP,RS30S,IF1,fMettRNAfMetCAU], k_forward= rxn_k['re0000000703_k1']),
    # Reaction.from_massaction([RS30S_IF1_IF2_GTP_fMettRNAfMetCAU],[IF1_degraded,RS30S,fMettRNAfMetCAU,IF2,GTP], k_forward= rxn_k['re0000000704_k1']),

    # Reaction.from_massaction([RS30S_IF1_fMettRNAfMetCAU_mRNA],[RS30S_degraded,IF1,fMettRNAfMetCAU,mRNA], k_forward= rxn_k['re0000000707_k1']),
    # Reaction.from_massaction([RS30S_IF1_fMettRNAfMetCAU_mRNA],[IF1_degraded,mRNA,fMettRNAfMetCAU,RS30S], k_forward= rxn_k['re0000000708_k1']),
    # Reaction.from_massaction([RS30S_IF1_IF2_GTP_fMettRNAfMetCAU_mRNA],[RS30S_degraded,mRNA,fMettRNAfMetCAU,IF2,GTP,IF1], k_forward= rxn_k['re0000000712_k1']),
    # Reaction.from_massaction([RS30S_IF1_IF2_GTP_fMettRNAfMetCAU_mRNA],[IF2_degraded,mRNA,fMettRNAfMetCAU,RS30S,IF1,GTP], k_forward= rxn_k['re0000000713_k1']),
    # Reaction.from_massaction([RS30S_IF1_IF2_GTP_fMettRNAfMetCAU_mRNA],[IF1_degraded,GTP,IF2,RS30S,fMettRNAfMetCAU,mRNA], k_forward= rxn_k['re0000000714_k1']),
    Reaction.from_massaction([RS70S_IF3_IF2_GTP_fMettRNAfMetCAU_mRNA],[RS70S_IF3_IF2_GDP_PO4_fMettRNAfMetCAU_mRNA], k_forward= rxn_k['re0000000715_k1']),
    Reaction.from_massaction([RS70S_IF3_IF2_GDP_PO4_fMettRNAfMetCAU_mRNA],[RS70S_IF3_IF2_GTP_fMettRNAfMetCAU_mRNA], k_forward= rxn_k['re0000000716_k1']),
    Reaction.from_massaction([RS70S_IF1_IF3_IF2_GTP_fMettRNAfMetCAU_mRNA],[RS70S_IF1_IF3_IF2_GDP_PO4_fMettRNAfMetCAU_mRNA], k_forward= rxn_k['re0000000717_k1']),
    Reaction.from_massaction([RS70S_IF1_IF3_IF2_GDP_PO4_fMettRNAfMetCAU_mRNA],[RS70S_IF1_IF3_IF2_GTP_fMettRNAfMetCAU_mRNA], k_forward= rxn_k['re0000000718_k1']),
    Reaction.from_massaction([RS70S_IF3_IF2_GDP_PO4_fMettRNAfMetCAU_mRNA,IF1],[RS70S_IF1_IF3_IF2_GDP_PO4_fMettRNAfMetCAU_mRNA], k_forward= rxn_k['re0000000719_k1']),

    Reaction.from_massaction([RS70S_IF1_IF3_IF2_GDP_PO4_fMettRNAfMetCAU_mRNA],[RS70S_IF3_IF2_GDP_PO4_fMettRNAfMetCAU_mRNA,IF1], k_forward= rxn_k['re0000000720_k1']),
    Reaction.from_massaction([RS70S_IF3_IF2_GDP_PO4_fMettRNAfMetCAU_mRNA],[RS70S_IF3_IF2_GDP_fMettRNAfMetCAU_mRNA,PO4], k_forward= rxn_k['re0000000721_k1']),
    Reaction.from_massaction([RS70S_IF1_IF3_IF2_GDP_PO4_fMettRNAfMetCAU_mRNA],[RS70S_IF1_IF3_IF2_GDP_fMettRNAfMetCAU_mRNA,PO4], k_forward= rxn_k['re0000000722_k1']),
    Reaction.from_massaction([RS70S_IF3_IF2_GDP_fMettRNAfMetCAU_mRNA,IF1],[RS70S_IF1_IF3_IF2_GDP_fMettRNAfMetCAU_mRNA], k_forward= rxn_k['re0000000723_k1']),
    Reaction.from_massaction([RS70S_IF1_IF3_IF2_GDP_fMettRNAfMetCAU_mRNA],[RS70S_IF3_IF2_GDP_fMettRNAfMetCAU_mRNA,IF1], k_forward= rxn_k['re0000000724_k1']),
    Reaction.from_massaction([RS70S_IF3_IF2_GDP_fMettRNAfMetCAU_mRNA],[RS70S_IF3_fMettRNAfMetCAU_mRNA,IF2_GDP], k_forward= rxn_k['re0000000725_k1']),
    # Reaction.from_massaction([RS70S_IF3_IF2_GDP_fMettRNAfMetCAU_mRNA],[IF2_degraded,mRNA,GDP,fMettRNAfMetCAU,RS50S,RS30S,IF3], k_forward= rxn_k['re0000000727_k1']),
    # Reaction.from_massaction([RS70S_IF3_IF2_GDP_fMettRNAfMetCAU_mRNA],[RS50S_degraded,IF2,mRNA,fMettRNAfMetCAU,GDP,RS30S,IF3], k_forward= rxn_k['re0000000728_k1']),
    # Reaction.from_massaction([RS70S_IF3_IF2_GDP_fMettRNAfMetCAU_mRNA],[IF3_degraded,GDP,fMettRNAfMetCAU,mRNA,RS30S,RS50S,IF2], k_forward= rxn_k['re0000000729_k1']),
    # Reaction.from_massaction([RS70S_IF3_IF2_GDP_fMettRNAfMetCAU_mRNA],[RS30S_degraded,RS50S,mRNA,fMettRNAfMetCAU,IF2,IF3,GDP], k_forward= rxn_k['re0000000730_k1']),

    # Reaction.from_massaction([RS70S_IF1_IF3_IF2_GDP_fMettRNAfMetCAU_mRNA],[RS30S_degraded,RS50S,GDP,mRNA,fMettRNAfMetCAU,IF2,IF3,IF1], k_forward= rxn_k['re0000000731_k1']),
    # Reaction.from_massaction([RS70S_IF1_IF3_IF2_GDP_fMettRNAfMetCAU_mRNA],[IF2_degraded,mRNA,GDP,fMettRNAfMetCAU,RS50S,RS30S,IF3,IF1], k_forward= rxn_k['re0000000732_k1']),
    # Reaction.from_massaction([RS70S_IF1_IF3_IF2_GDP_fMettRNAfMetCAU_mRNA],[RS50S_degraded,mRNA,IF2,GDP,fMettRNAfMetCAU,RS30S,IF3,IF1], k_forward= rxn_k['re0000000733_k1']),
    # Reaction.from_massaction([RS70S_IF1_IF3_IF2_GDP_fMettRNAfMetCAU_mRNA],[IF1_degraded,mRNA,GDP,fMettRNAfMetCAU,IF2,RS30S,IF3,RS50S], k_forward= rxn_k['re0000000734_k1']),
    # Reaction.from_massaction([RS70S_IF1_IF3_IF2_GDP_fMettRNAfMetCAU_mRNA],[IF3_degraded,RS50S,RS30S,IF1,mRNA,IF2,fMettRNAfMetCAU,GDP], k_forward= rxn_k['re0000000735_k1']),
    # Reaction.from_massaction([RS70S_IF3_IF2_GDP_PO4_fMettRNAfMetCAU_mRNA],[IF2_degraded,mRNA,GDP,fMettRNAfMetCAU,RS50S,RS30S,IF3,PO4], k_forward= rxn_k['re0000000736_k1']),
    # Reaction.from_massaction([RS70S_IF3_IF2_GDP_PO4_fMettRNAfMetCAU_mRNA],[RS50S_degraded,IF2,mRNA,fMettRNAfMetCAU,GDP,RS30S,IF3,PO4], k_forward= rxn_k['re0000000737_k1']),
    # Reaction.from_massaction([RS70S_IF3_IF2_GDP_PO4_fMettRNAfMetCAU_mRNA],[IF3_degraded,GDP,fMettRNAfMetCAU,mRNA,RS30S,RS50S,IF2,PO4], k_forward= rxn_k['re0000000738_k1']),
    # Reaction.from_massaction([RS70S_IF3_IF2_GDP_PO4_fMettRNAfMetCAU_mRNA],[RS30S_degraded,RS50S,mRNA,fMettRNAfMetCAU,IF2,IF3,GDP,PO4], k_forward= rxn_k['re0000000739_k1']),
    # Reaction.from_massaction([RS70S_IF1_IF3_IF2_GDP_PO4_fMettRNAfMetCAU_mRNA],[RS30S_degraded,RS50S,GDP,mRNA,fMettRNAfMetCAU,IF2,IF3,IF1,PO4], k_forward= rxn_k['re0000000740_k1']),

    # Reaction.from_massaction([RS70S_IF1_IF3_IF2_GDP_PO4_fMettRNAfMetCAU_mRNA],[IF2_degraded,mRNA,GDP,fMettRNAfMetCAU,RS50S,RS30S,IF3,IF1,PO4], k_forward= rxn_k['re0000000741_k1']),
    # Reaction.from_massaction([RS70S_IF1_IF3_IF2_GDP_PO4_fMettRNAfMetCAU_mRNA],[RS50S_degraded,mRNA,IF2,GDP,fMettRNAfMetCAU,RS30S,IF3,IF1,PO4], k_forward= rxn_k['re0000000742_k1']),
    # Reaction.from_massaction([RS70S_IF1_IF3_IF2_GDP_PO4_fMettRNAfMetCAU_mRNA],[IF1_degraded,mRNA,GDP,fMettRNAfMetCAU,IF2,RS30S,IF3,RS50S,PO4], k_forward= rxn_k['re0000000743_k1']),
    # Reaction.from_massaction([RS70S_IF1_IF3_IF2_GDP_PO4_fMettRNAfMetCAU_mRNA],[IF3_degraded,RS50S,RS30S,IF1,mRNA,IF2,fMettRNAfMetCAU,GDP,PO4], k_forward= rxn_k['re0000000744_k1']),
    # Reaction.from_massaction([RS70S_IF3_IF2_GDP_fMettRNAfMetCAU_mRNA,PO4],[RS70S_IF3_IF2_GDP_PO4_fMettRNAfMetCAU_mRNA], k_forward= rxn_k['re0000000745_k1']),
    # Reaction.from_massaction([RS70S_IF1_IF3_IF2_GDP_fMettRNAfMetCAU_mRNA,PO4],[RS70S_IF1_IF3_IF2_GDP_PO4_fMettRNAfMetCAU_mRNA], k_forward= rxn_k['re0000000746_k1']),
    Reaction.from_massaction([RS70S_IF1_IF3_IF2_GDP_fMettRNAfMetCAU_mRNA],[RS70S_IF1_IF2_GDP_fMettRNAfMetCAU_mRNA,IF3], k_forward= rxn_k['re0000000747_k1']),
    # Reaction.from_massaction([RS70S_IF1_IF2_GDP_fMettRNAfMetCAU_mRNA,IF3],[RS70S_IF1_IF3_IF2_GDP_fMettRNAfMetCAU_mRNA], k_forward= rxn_k['re0000000748_k1']),
    Reaction.from_massaction([RS70S_IF1_IF3_IF2_GDP_fMettRNAfMetCAU_mRNA],[RS70S_IF1_IF3_fMettRNAfMetCAU_mRNA,IF2_GDP], k_forward= rxn_k['re0000000749_k1']),
    Reaction.from_massaction([RS70S_IF1_IF3_fMettRNAfMetCAU_mRNA,IF2_GDP],[RS70S_IF1_IF3_IF2_GDP_fMettRNAfMetCAU_mRNA], k_forward= rxn_k['re0000000750_k1']),

    Reaction.from_massaction([RS70S_IF3_fMettRNAfMetCAU_mRNA,IF2_GDP],[RS70S_IF3_IF2_GDP_fMettRNAfMetCAU_mRNA], k_forward= rxn_k['re0000000751_k1']),
    Reaction.from_massaction([RS70S_IF1_IF3_fMettRNAfMetCAU_mRNA],[RS70S_IF3_fMettRNAfMetCAU_mRNA,IF1], k_forward= rxn_k['re0000000753_k1']),
    # Reaction.from_massaction([RS70S_IF3_fMettRNAfMetCAU_mRNA,IF1],[RS70S_IF1_IF3_fMettRNAfMetCAU_mRNA], k_forward= rxn_k['re0000000754_k1']),
    Reaction.from_massaction([RS70S_IF3_IF2_GDP_fMettRNAfMetCAU_mRNA],[RS70S_IF2_GDP_fMettRNAfMetCAU_mRNA,IF3], k_forward= rxn_k['re0000000755_k1']),
    # Reaction.from_massaction([RS70S_IF2_GDP_fMettRNAfMetCAU_mRNA,IF3],[RS70S_IF3_IF2_GDP_fMettRNAfMetCAU_mRNA], k_forward= rxn_k['re0000000756_k1']),
    Reaction.from_massaction([RS70S_IF1_IF2_GDP_fMettRNAfMetCAU_mRNA],[RS70S_IF2_GDP_fMettRNAfMetCAU_mRNA,IF1], k_forward= rxn_k['re0000000757_k1']),
    # Reaction.from_massaction([RS70S_IF2_GDP_fMettRNAfMetCAU_mRNA,IF1],[RS70S_IF1_IF2_GDP_fMettRNAfMetCAU_mRNA], k_forward= rxn_k['re0000000758_k1']),
    Reaction.from_massaction([RS70S_IF1_IF2_GDP_fMettRNAfMetCAU_mRNA],[RS70S_IF1_fMettRNAfMetCAU_mRNA,IF2_GDP], k_forward= rxn_k['re0000000759_k1']),
    Reaction.from_massaction([RS70S_IF1_fMettRNAfMetCAU_mRNA,IF2_GDP],[RS70S_IF1_IF2_GDP_fMettRNAfMetCAU_mRNA], k_forward= rxn_k['re0000000760_k1']),
    Reaction.from_massaction([RS70S_IF1_IF3_fMettRNAfMetCAU_mRNA],[RS70S_IF1_fMettRNAfMetCAU_mRNA,IF3], k_forward= rxn_k['re0000000761_k1']),

    # Reaction.from_massaction([RS70S_IF1_fMettRNAfMetCAU_mRNA,IF3],[RS70S_IF1_IF3_fMettRNAfMetCAU_mRNA], k_forward= rxn_k['re0000000762_k1']),
    # Reaction.from_massaction([RS70S_IF2_GDP_fMettRNAfMetCAU_mRNA],[RS30S_degraded,RS50S,GDP,mRNA,fMettRNAfMetCAU,IF2], k_forward= rxn_k['re0000000767_k1']),
    # Reaction.from_massaction([RS70S_IF2_GDP_fMettRNAfMetCAU_mRNA],[IF2_degraded,mRNA,GDP,fMettRNAfMetCAU,RS50S,RS30S], k_forward= rxn_k['re0000000768_k1']),
    # Reaction.from_massaction([RS70S_IF2_GDP_fMettRNAfMetCAU_mRNA],[RS50S_degraded,mRNA,IF2,GDP,fMettRNAfMetCAU,RS30S], k_forward= rxn_k['re0000000769_k1']),
    # Reaction.from_massaction([RS70S_IF3_fMettRNAfMetCAU_mRNA],[RS30S_degraded,RS50S,mRNA,fMettRNAfMetCAU,IF3], k_forward= rxn_k['re0000000770_k1']),
    # Reaction.from_massaction([RS70S_IF3_fMettRNAfMetCAU_mRNA],[RS50S_degraded,mRNA,fMettRNAfMetCAU,RS30S,IF3], k_forward= rxn_k['re0000000771_k1']),
    # Reaction.from_massaction([RS70S_IF3_fMettRNAfMetCAU_mRNA],[IF3_degraded,RS50S,RS30S,mRNA,fMettRNAfMetCAU], k_forward= rxn_k['re0000000772_k1']),
    # Reaction.from_massaction([RS70S_IF1_IF3_fMettRNAfMetCAU_mRNA],[RS30S_degraded,RS50S,mRNA,fMettRNAfMetCAU,IF3,IF1], k_forward= rxn_k['re0000000773_k1']),
    # Reaction.from_massaction([RS70S_IF1_IF3_fMettRNAfMetCAU_mRNA],[RS50S_degraded,mRNA,fMettRNAfMetCAU,RS30S,IF3,IF1], k_forward= rxn_k['re0000000774_k1']),
    # Reaction.from_massaction([RS70S_IF1_IF3_fMettRNAfMetCAU_mRNA],[IF1_degraded,mRNA,fMettRNAfMetCAU,RS30S,IF3,RS50S], k_forward= rxn_k['re0000000775_k1']),

    # Reaction.from_massaction([RS70S_IF1_IF3_fMettRNAfMetCAU_mRNA],[IF3_degraded,RS50S,RS30S,IF1,mRNA,fMettRNAfMetCAU], k_forward= rxn_k['re0000000776_k1']),
    # Reaction.from_massaction([RS70S_IF1_fMettRNAfMetCAU_mRNA],[RS30S_degraded,RS50S,mRNA,fMettRNAfMetCAU,IF1], k_forward= rxn_k['re0000000777_k1']),
    # Reaction.from_massaction([RS70S_IF1_fMettRNAfMetCAU_mRNA],[RS50S_degraded,mRNA,fMettRNAfMetCAU,RS30S,IF1], k_forward= rxn_k['re0000000778_k1']),
    # Reaction.from_massaction([RS70S_IF1_fMettRNAfMetCAU_mRNA],[IF1_degraded,mRNA,fMettRNAfMetCAU,RS30S,RS50S], k_forward= rxn_k['re0000000779_k1']),
    # Reaction.from_massaction([RS70S_IF1_IF2_GDP_fMettRNAfMetCAU_mRNA],[RS30S_degraded,RS50S,GDP,mRNA,fMettRNAfMetCAU,IF2,IF1], k_forward= rxn_k['re0000000780_k1']),
    # Reaction.from_massaction([RS70S_IF1_IF2_GDP_fMettRNAfMetCAU_mRNA],[IF2_degraded,mRNA,GDP,fMettRNAfMetCAU,RS50S,RS30S,IF1], k_forward= rxn_k['re0000000781_k1']),
    # Reaction.from_massaction([RS70S_IF1_IF2_GDP_fMettRNAfMetCAU_mRNA],[RS50S_degraded,mRNA,IF2,GDP,fMettRNAfMetCAU,RS30S,IF1], k_forward= rxn_k['re0000000782_k1']),
    # Reaction.from_massaction([RS70S_IF1_IF2_GDP_fMettRNAfMetCAU_mRNA],[IF1_degraded,mRNA,GDP,fMettRNAfMetCAU,IF2,RS30S,RS50S], k_forward= rxn_k['re0000000783_k1']),
    ]
```

#### Addition of AA individual reactions required for desire protein production¶

In [10]:

```
#######################################################################################################################
#Creates a list of all the species associated with the aa in the chain, except if aa=Met
#Iterates over all the aa in input aa chain
for aa in AA:
    xyz= list_condon[list_AA.index(aa)] #matches the codon for each of the AA
    
    if aa != 'Met': #does not include Met since equations already exists for it.
        asub=aa
        asub = Species(asub)
        aaAMP = Species(aa+'AMP')
        aaRS = Species(aa+'RS')
        aaRS_AMP = Species(aa+'RS_AMP')
        aaRS_ATP = Species(aa+'RS_ATP')
        aaRS_degraded = Species(aa+'RS_degraded')
        aaRS_aa= Species(aa+'RS_'+aa)
        aaRS_aa_ATP = Species(aa+'RS_'+aa+'_ATP')
        aaRS_aaAMP = Species(aa+'RS_'+aa+'AMP')
        aaRS_aaAMP_PPi = Species(aa+'RS_'+aa+'AMP_PPi')
        
        
        aaRS_AMP_aatRNAaaxyz = Species(aa+'RS_AMP_'+aa+'tRNA'+aa+xyz)
        aaRS_ATP_tRNAaaxyz = Species(aa+'RS_ATP_tRNA'+aa+xyz)
        aaRS_aa_ATP_tRNAaaxyz = Species(aa+'RS_'+aa+'_ATP_tRNA'+aa+xyz)
        aaRS_aa_tRNAaaxyz = Species(aa+'RS_'+aa+'_tRNA'+aa+xyz)
        aaRS_aaAMP_PPi_tRNAaaxyz = Species(aa+'RS_'+aa+'AMP_PPi_tRNA'+aa+xyz)
        aaRS_aaAMP_tRNAaaxyz = Species(aa+'RS_'+aa+'AMP_tRNA'+aa+xyz)
        aaRS_aatRNAaaxyz = Species(aa+'RS_'+aa+'tRNA'+aa+xyz)
        aaRS_tRNAaaxyz = Species(aa+'RS_tRNA'+aa+xyz)
        aatRNAaaxyz = Species(aa+'tRNA'+aa+xyz)
        aatRNAaaxyz_degraded = Species(aa+'tRNA'+aa+xyz+'_degraded')
        tRNAaaxyz = Species('tRNA'+aa+xyz)
        tRNAaaxyz_degraded = Species('tRNA'+aa+xyz+'_degraded')
        
        RS50S_tRNAaaxyz= Species('RS50S_tRNA'+aa+xyz)
        RS50S_tRNAaaxyz_EFG_GDP = Species('RS50S_tRNA'+aa+xyz+'_EFG_GDP')
        RS50S_tRNAaaxyz_RRF = Species('RS50S_tRNA'+aa+xyz+'_RRF')
        RS50S_tRNAaaxyz_RRF_EFG_GDP = Species('RS50S_tRNA'+aa+xyz+'_RRF_EFG_GDP')
        EFTu_GTP_aatRNAaaxyz = Species('EFTu_GTP_'+aa+'tRNA'+aa+xyz)
        
        aa = Species(aa)
        #######################################################################################################################

        #A list of Species
        species_aa= [
            asub, aaAMP, aaRS, aaRS_AMP, aaRS_ATP,
            aaRS_degraded, aaRS_aa, aaRS_aa_ATP, aaRS_aaAMP, 
            aaRS_aaAMP_PPi, aaRS_AMP_aatRNAaaxyz, aaRS_ATP_tRNAaaxyz, aaRS_aa_ATP_tRNAaaxyz,
            aaRS_aa_tRNAaaxyz, aaRS_aaAMP_PPi_tRNAaaxyz, aaRS_aaAMP_tRNAaaxyz, aaRS_aatRNAaaxyz, 
            aaRS_tRNAaaxyz, aatRNAaaxyz, aatRNAaaxyz_degraded,tRNAaaxyz, tRNAaaxyz_degraded,
            
            RS50S_tRNAaaxyz, RS50S_tRNAaaxyz_EFG_GDP, RS50S_tRNAaaxyz_RRF, RS50S_tRNAaaxyz_RRF_EFG_GDP, EFTu_GTP_aatRNAaaxyz]        
        
        #######################################################################################################################
        
        #Reactions
        rxn=[Reaction.from_massaction([aaRS,aa],[aaRS_aa], k_forward = rxn_k['re0000000126_k1']),
            Reaction.from_massaction([aaRS_aaAMP_PPi],[aaRS_aaAMP,PPi], k_forward = rxn_k['re0000000127_k1']),
            # Reaction.from_massaction([aaRS],[aaRS_degraded], k_forward = rxn_k['re0000000128_k1']),
            # Reaction.from_massaction([aaRS_aa_ATP],[aaRS_degraded,aa,ATP], k_forward = rxn_k['re0000000129_k1']),
            # Reaction.from_massaction([aaRS_aaAMP],[aaRS_degraded,aaAMP], k_forward = rxn_k['re0000000130_k1']),
            Reaction.from_massaction([aaRS_aa],[aaRS,aa], k_forward = rxn_k['re0000000131_k1']),
            Reaction.from_massaction([aaRS,ATP],[aaRS_ATP], k_forward = rxn_k['re0000000132_k1']),
            Reaction.from_massaction([aaRS_ATP],[aaRS,ATP], k_forward = rxn_k['re0000000133_k1']),
            Reaction.from_massaction([aaRS_ATP,aa],[aaRS_aa_ATP], k_forward = rxn_k['re0000000134_k1']),
            Reaction.from_massaction([aaRS_aa_ATP],[aaRS_ATP,aa], k_forward = rxn_k['re0000000135_k1']),
            Reaction.from_massaction([aaRS_aa,ATP],[aaRS_aa_ATP], k_forward = rxn_k['re0000000136_k1']),
            Reaction.from_massaction([aaRS_aa_ATP],[aaRS_aa,ATP], k_forward = rxn_k['re0000000137_k1']),
            # Reaction.from_massaction([aaRS_ATP],[aaRS_degraded,ATP], k_forward = rxn_k['re0000000138_k1']),
            # Reaction.from_massaction([aaRS_aa],[aaRS_degraded,aa], k_forward = rxn_k['re0000000139_k1']),
            Reaction.from_massaction([aaRS_aa_ATP],[aaRS_aaAMP_PPi], k_forward = rxn_k['re0000000140_k1']),
            Reaction.from_massaction([aaRS_aaAMP_PPi],[aaRS_aa_ATP], k_forward = rxn_k['re0000000141_k1']),
            # Reaction.from_massaction([aaRS_aaAMP_PPi],[aaRS_degraded,aaAMP,PPi], k_forward = rxn_k['re0000000142_k1']),
            # Reaction.from_massaction([aaAMP],[aa,AMP], k_forward = rxn_k['re0000000143_k1']),
            # Reaction.from_massaction([aaRS_AMP],[aaRS_degraded,AMP], k_forward = rxn_k['re0000000144_k1']),
            Reaction.from_massaction([aaRS_AMP],[aaRS,AMP], k_forward = rxn_k['re0000000145_k1']),
            # Reaction.from_massaction([aaRS,AMP],[aaRS_AMP], k_forward = rxn_k['re0000000146_k1']),
            Reaction.from_massaction([aaRS_aaAMP],[aaRS,aaAMP], k_forward = rxn_k['re0000000147_k1']),
            Reaction.from_massaction([aaRS,aaAMP],[aaRS_aaAMP], k_forward = rxn_k['re0000000148_k1']),
            # Reaction.from_massaction([aa,AMP],[aaAMP], k_forward = rxn_k['re0000000149_k1']),
            # Reaction.from_massaction([aaRS_aaAMP,PPi],[aaRS_aaAMP_PPi], k_forward = rxn_k['re0000000150_k1']),
             
            # Reaction.from_massaction([aatRNAaaxyz],[tRNAaaxyz,aa], k_forward = rxn_k['re0000000176_k1']),
            # Reaction.from_massaction([aaRS_aaAMP_tRNAaaxyz],[aaRS_degraded,aaAMP,tRNAaaxyz], k_forward = rxn_k['re0000000177_k1']),
            Reaction.from_massaction([aaRS_aaAMP_tRNAaaxyz],[aaRS_AMP_aatRNAaaxyz], k_forward = rxn_k['re0000000178_k1']),
            # Reaction.from_massaction([aaRS_AMP_aatRNAaaxyz],[aaRS_aaAMP_tRNAaaxyz], k_forward = rxn_k['re0000000179_k1']),
            Reaction.from_massaction([aaRS_AMP_aatRNAaaxyz],[aaRS_aatRNAaaxyz,AMP], k_forward = rxn_k['re0000000180_k1']),
            # Reaction.from_massaction([aaRS_aatRNAaaxyz,AMP],[aaRS_AMP_aatRNAaaxyz], k_forward = rxn_k['re0000000181_k1']),
            Reaction.from_massaction([aaRS_AMP_aatRNAaaxyz],[aatRNAaaxyz,aaRS_AMP], k_forward = rxn_k['re0000000182_k1']),
            Reaction.from_massaction([aaRS_AMP,aatRNAaaxyz],[aaRS_AMP_aatRNAaaxyz], k_forward = rxn_k['re0000000183_k1']),
            Reaction.from_massaction([aaRS_aatRNAaaxyz],[aaRS,aatRNAaaxyz], k_forward = rxn_k['re0000000184_k1']),
            Reaction.from_massaction([aaRS,aatRNAaaxyz],[aaRS_aatRNAaaxyz], k_forward = rxn_k['re0000000185_k1']),
            # Reaction.from_massaction([aaRS_AMP_aatRNAaaxyz],[aaRS_degraded,AMP,aatRNAaaxyz], k_forward = rxn_k['re0000000186_k1']),
            # Reaction.from_massaction([aaRS_aatRNAaaxyz],[aaRS_degraded,aatRNAaaxyz], k_forward = rxn_k['re0000000187_k1']),
            Reaction.from_massaction([aaRS_tRNAaaxyz,aa],[aaRS_aa_tRNAaaxyz], k_forward = rxn_k['re0000000188_k1']),
            Reaction.from_massaction([aaRS_aaAMP_PPi_tRNAaaxyz],[aaRS_aaAMP_tRNAaaxyz,PPi], k_forward = rxn_k['re0000000189_k1']),
            Reaction.from_massaction([aaRS_aa_tRNAaaxyz],[aaRS_tRNAaaxyz,aa], k_forward = rxn_k['re0000000190_k1']),
            Reaction.from_massaction([aaRS_tRNAaaxyz,ATP],[aaRS_ATP_tRNAaaxyz], k_forward = rxn_k['re0000000191_k1']),
            Reaction.from_massaction([aaRS_ATP_tRNAaaxyz],[aaRS_tRNAaaxyz,ATP], k_forward = rxn_k['re0000000192_k1']),
            Reaction.from_massaction([aaRS_ATP_tRNAaaxyz,aa],[aaRS_aa_ATP_tRNAaaxyz], k_forward = rxn_k['re0000000193_k1']),
            Reaction.from_massaction([aaRS_aa_ATP_tRNAaaxyz],[aaRS_ATP_tRNAaaxyz,aa], k_forward = rxn_k['re0000000194_k1']),
            Reaction.from_massaction([aaRS_aa_tRNAaaxyz,ATP],[aaRS_aa_ATP_tRNAaaxyz], k_forward = rxn_k['re0000000195_k1']),
            Reaction.from_massaction([aaRS_aa_ATP_tRNAaaxyz],[aaRS_aa_tRNAaaxyz,ATP], k_forward = rxn_k['re0000000196_k1']),
            Reaction.from_massaction([aaRS_aa_ATP_tRNAaaxyz],[aaRS_aaAMP_PPi_tRNAaaxyz], k_forward = rxn_k['re0000000197_k1']),
            # Reaction.from_massaction([aaRS_aaAMP_PPi_tRNAaaxyz],[aaRS_aa_ATP_tRNAaaxyz], k_forward = rxn_k['re0000000198_k1']),
            Reaction.from_massaction([aaRS,tRNAaaxyz],[aaRS_tRNAaaxyz], k_forward = rxn_k['re0000000199_k1']),
            Reaction.from_massaction([aaRS_aa,tRNAaaxyz],[aaRS_aa_tRNAaaxyz], k_forward = rxn_k['re0000000200_k1']),
            Reaction.from_massaction([aaRS_tRNAaaxyz],[aaRS,tRNAaaxyz], k_forward = rxn_k['re0000000201_k1']),
            Reaction.from_massaction([aaRS_aa_tRNAaaxyz],[aaRS_aa,tRNAaaxyz], k_forward = rxn_k['re0000000202_k1']),
            Reaction.from_massaction([aaRS_ATP,tRNAaaxyz],[aaRS_ATP_tRNAaaxyz], k_forward = rxn_k['re0000000203_k1']),
            Reaction.from_massaction([aaRS_ATP_tRNAaaxyz],[aaRS_ATP,tRNAaaxyz], k_forward = rxn_k['re0000000204_k1']),
            Reaction.from_massaction([aaRS_aa_ATP,tRNAaaxyz],[aaRS_aa_ATP_tRNAaaxyz], k_forward = rxn_k['re0000000205_k1']),
            Reaction.from_massaction([aaRS_aa_ATP_tRNAaaxyz],[aaRS_aa_ATP,tRNAaaxyz], k_forward = rxn_k['re0000000206_k1']),
            Reaction.from_massaction([aaRS_aaAMP_PPi,tRNAaaxyz],[aaRS_aaAMP_PPi_tRNAaaxyz], k_forward = rxn_k['re0000000207_k1']),
            Reaction.from_massaction([aaRS_aaAMP_PPi_tRNAaaxyz],[aaRS_aaAMP_PPi,tRNAaaxyz], k_forward = rxn_k['re0000000208_k1']),
            Reaction.from_massaction([aaRS_aaAMP,tRNAaaxyz],[aaRS_aaAMP_tRNAaaxyz], k_forward = rxn_k['re0000000209_k1']),
            Reaction.from_massaction([aaRS_aaAMP_tRNAaaxyz],[aaRS_aaAMP,tRNAaaxyz], k_forward = rxn_k['re0000000210_k1']),
            # Reaction.from_massaction([aaRS_tRNAaaxyz],[aaRS_degraded,tRNAaaxyz], k_forward = rxn_k['re0000000211_k1']),
            # Reaction.from_massaction([aaRS_ATP_tRNAaaxyz],[aaRS_degraded,tRNAaaxyz,ATP], k_forward = rxn_k['re0000000212_k1']),
            # Reaction.from_massaction([aaRS_aa_tRNAaaxyz],[aaRS_degraded,tRNAaaxyz,aa], k_forward = rxn_k['re0000000213_k1']),
            # Reaction.from_massaction([aaRS_aa_ATP_tRNAaaxyz],[aaRS_degraded,tRNAaaxyz,ATP,aa], k_forward = rxn_k['re0000000214_k1']),
            # Reaction.from_massaction([aaRS_aaAMP_PPi_tRNAaaxyz],[aaRS_degraded,tRNAaaxyz,aaAMP,PPi], k_forward = rxn_k['re0000000215_k1']),
            # Reaction.from_massaction([tRNAaaxyz,aa],[aatRNAaaxyz], k_forward = rxn_k['re0000000216_k1']),
            # Reaction.from_massaction([aaRS_aaAMP_tRNAaaxyz,PPi],[aaRS_aaAMP_PPi_tRNAaaxyz], k_forward = rxn_k['re0000000217_k1']),
             
            Reaction.from_massaction([EFTu_GTP,aatRNAaaxyz],[EFTu_GTP_aatRNAaaxyz], k_forward = rxn_k['re0000000275_k1']),
            Reaction.from_massaction([EFTu_GTP_aatRNAaaxyz],[EFTu_GTP,aatRNAaaxyz], k_forward = rxn_k['re0000000276_k1']),
            # Reaction.from_massaction([EFTu_GTP_aatRNAaaxyz],[EFTu_degraded,aatRNAaaxyz,GTP], k_forward = rxn_k['re0000000286_k1']),
            # Reaction.from_massaction([EFTu_GTP_aatRNAaaxyz],[aatRNAaaxyz_degraded,EFTu,GTP], k_forward = rxn_k['re0000000287_k1']),
            
            # Reaction.from_massaction([aatRNAaaxyz],[aatRNAaaxyz_degraded], k_forward = rxn_k['re0000000031_k1']),
            # Reaction.from_massaction([tRNAaaxyz],[tRNAaaxyz_degraded], k_forward = rxn_k['re0000000070_k1']),
            # Reaction.from_massaction([RRF,RS50S_tRNAaaxyz_EFG_GDP],[RS50S_tRNAaaxyz_RRF_EFG_GDP], k_forward = rxn_k['re0000000958_k1']), 
            # Reaction.from_massaction([EFG_GDP,RS50S_tRNAaaxyz_RRF],[RS50S_tRNAaaxyz_RRF_EFG_GDP], k_forward = rxn_k['re0000000959_k1']),
            # Reaction.from_massaction([EFG_GDP,RS50S_tRNAaaxyz],[RS50S_tRNAaaxyz_EFG_GDP], k_forward = rxn_k['re0000000961_k1']),
            # Reaction.from_massaction([RRF,RS50S_tRNAaaxyz],[RS50S_tRNAaaxyz_RRF], k_forward = rxn_k['re0000000962_k1']),

            Reaction.from_massaction([RS50S_tRNAaaxyz_RRF_EFG_GDP],[RS50S_tRNAaaxyz_RRF,EFG_GDP], k_forward = rxn_k['re0000000913_k1']),
            Reaction.from_massaction([RS50S_tRNAaaxyz_RRF_EFG_GDP],[RS50S_tRNAaaxyz_EFG_GDP,RRF], k_forward = rxn_k['re0000000914_k1']),
            Reaction.from_massaction([RS50S_tRNAaaxyz_RRF_EFG_GDP],[RS50S_RRF_EFG_GDP,tRNAaaxyz], k_forward = rxn_k['re0000000915_k1']),
            Reaction.from_massaction([RS50S_tRNAaaxyz_RRF],[RS50S_RRF,tRNAaaxyz], k_forward = rxn_k['re0000000916_k1']),
            Reaction.from_massaction([RS50S_tRNAaaxyz_RRF],[RS50S_tRNAaaxyz,RRF], k_forward = rxn_k['re0000000917_k1']),

            Reaction.from_massaction([RS50S_tRNAaaxyz],[tRNAaaxyz,RS50S], k_forward = rxn_k['re0000000919_k1']),
            Reaction.from_massaction([RS50S_tRNAaaxyz_EFG_GDP],[EFG_GDP,RS50S_tRNAaaxyz], k_forward = rxn_k['re0000000920_k1']),
            Reaction.from_massaction([RS50S_tRNAaaxyz_EFG_GDP],[RS50S_EFG_GDP,tRNAaaxyz], k_forward = rxn_k['re0000000921_k1']),
            # Reaction.from_massaction([RS50S_tRNAaaxyz],[RS50S_degraded,tRNAaaxyz], k_forward = rxn_k['re0000000924_k1']),
            # Reaction.from_massaction([RS50S_tRNAaaxyz],[tRNAaaxyz_degraded,RS50S], k_forward = rxn_k['re0000000927_k1']),

            # Reaction.from_massaction([RS50S_tRNAaaxyz_RRF],[tRNAaaxyz_degraded,RS50S,RRF], k_forward = rxn_k['re0000000929_k1']),
            # Reaction.from_massaction([RS50S_tRNAaaxyz_RRF],[RS50S_degraded,RRF,tRNAaaxyz], k_forward = rxn_k['re0000000930_k1']),
            # Reaction.from_massaction([RS50S_tRNAaaxyz_RRF],[RRF_degraded,RS50S,tRNAaaxyz], k_forward = rxn_k['re0000000931_k1']),
            # Reaction.from_massaction([RS50S_tRNAaaxyz_EFG_GDP],[tRNAaaxyz_degraded,RS50S,EFG,GDP], k_forward = rxn_k['re0000000932_k1']),
            # Reaction.from_massaction([RS50S_tRNAaaxyz_EFG_GDP],[RS50S_degraded,EFG,tRNAaaxyz,GDP], k_forward = rxn_k['re0000000933_k1']),
            # Reaction.from_massaction([RS50S_tRNAaaxyz_EFG_GDP],[EFG_degraded,RS50S,tRNAaaxyz,GDP], k_forward = rxn_k['re0000000934_k1']),

            # Reaction.from_massaction([RS50S_tRNAaaxyz_RRF_EFG_GDP],[tRNAaaxyz_degraded,RS50S,EFG,GDP,RRF], k_forward = rxn_k['re0000000952_k1']),
            # Reaction.from_massaction([RS50S_tRNAaaxyz_RRF_EFG_GDP],[RS50S_degraded,EFG,tRNAaaxyz,GDP,RRF], k_forward = rxn_k['re0000000953_k1']),
            # Reaction.from_massaction([RS50S_tRNAaaxyz_RRF_EFG_GDP],[EFG_degraded,RS50S,tRNAaaxyz,GDP,RRF], k_forward = rxn_k['re0000000954_k1']),
            # Reaction.from_massaction([RS50S_tRNAaaxyz_RRF_EFG_GDP],[RRF_degraded,tRNAaaxyz,EFG,GDP,RS50S], k_forward = rxn_k['re0000000955_k1']),


            # Reaction.from_massaction([tRNAaaxyz,RS50S_EFG_GDP],[RS50S_tRNAaaxyz_EFG_GDP], k_forward = rxn_k['re0000000963_k1']),
            # Reaction.from_massaction([tRNAaaxyz,RS50S_RRF_EFG_GDP],[RS50S_tRNAaaxyz_RRF_EFG_GDP], k_forward = rxn_k['re0000000960_k1']),
            # Reaction.from_massaction([tRNAaaxyz,RS50S_RRF],[RS50S_tRNAaaxyz_RRF], k_forward = rxn_k['re0000000965_k1']),
            # Reaction.from_massaction([tRNAaaxyz,RS50S],[RS50S_tRNAaaxyz], k_forward = rxn_k['re0000000967_k1']),
            ]

        list_of_reactions.append(rxn)
        list_species_aa.append(species_aa)
    
    else:
            #Adding MetRS
            #This is different than fMet species and reactions for fMet

            MetRS_AMP_MettRNAMetCAU = Species('MetRS_AMP_tRNAMetCAU')
            MetRS_ATP_tRNAMetCAU = Species('MetRS_ATP_tRNAMetCAU')
            MetRS_Met_ATP_tRNAMetCAU = Species('MetRS_Met_ATP_tRNAMetCAU')
            MetRS_Met_tRNAMetCAU = Species('MetRS_Met_tRNAMetCAU')
            MetRS_MetAMP_PPi_tRNAMetCAU = Species('MetRS_MetAMP_PPi_tRNAMetCAU')
            MetRS_MetAMP_tRNAMetCAU = Species('MetRS_MetAMP_tRNAMetCAU')
            MetRS_MettRNAMetCAU = Species('MetRS_MettRNAMetCAU')
            MetRS_tRNAMetCAU = Species('MetRS_tRNAMetCAU')
            MettRNAMetCAU = Species('MettRNAMetCAU')
            MettRNAMetCAU_degraded = Species('MettRNAMetCAU_degraded')
            tRNAMetCAU = Species('tRNAMetCAU')
            tRNAMetCAU_degraded = Species('tRNAMetCAU_degraded')

            RS50S_tRNAMetCAU= Species('RS50S_tRNAMetCAU')
            RS50S_tRNAMetCAU_EFG_GDP = Species('RS50S_tRNAMetCAU_EFG_GDP')
            RS50S_tRNAMetCAU_RRF = Species('RS50S_tRNAMetCAU_RRF')
            RS50S_tRNAMetCAU_RRF_EFG_GDP = Species('RS50S_tRNAMetCAU_RRF_EFG_GDP')
            EFTu_GTP_MettRNAMetCAU = Species('EFTu_GTP_MettRNAMetCAU')

            #######################################################################################################################
            species_Met= [MetRS_AMP_MettRNAMetCAU, MetRS_ATP_tRNAMetCAU, MetRS_Met_ATP_tRNAMetCAU,
                            MetRS_Met_tRNAMetCAU, MetRS_MetAMP_PPi_tRNAMetCAU, MetRS_MetAMP_tRNAMetCAU, MetRS_MettRNAMetCAU, 
                            MetRS_tRNAMetCAU, MettRNAMetCAU, MettRNAMetCAU_degraded,tRNAMetCAU, tRNAMetCAU_degraded,

                            RS50S_tRNAMetCAU, RS50S_tRNAMetCAU_EFG_GDP, RS50S_tRNAMetCAU_RRF, RS50S_tRNAMetCAU_RRF_EFG_GDP, EFTu_GTP_MettRNAMetCAU]  

            #######################################################################################################################
            reactions_Met= [
                        Reaction.from_massaction([MetRS_MetAMP_tRNAMetCAU],[MetRS_AMP_MettRNAMetCAU], k_forward = rxn_k['re0000000178_k1']),
                        Reaction.from_massaction([MetRS_AMP_MettRNAMetCAU],[MetRS_MettRNAMetCAU,AMP], k_forward = rxn_k['re0000000180_k1']),

                        Reaction.from_massaction([MetRS_AMP_MettRNAMetCAU],[MettRNAMetCAU,MetRS_AMP], k_forward = rxn_k['re0000000182_k1']),
                        Reaction.from_massaction([MetRS_AMP,MettRNAMetCAU],[MetRS_AMP_MettRNAMetCAU], k_forward = rxn_k['re0000000183_k1']),
                        Reaction.from_massaction([MetRS_MettRNAMetCAU],[MetRS,MettRNAMetCAU], k_forward = rxn_k['re0000000184_k1']),
                        Reaction.from_massaction([MetRS,MettRNAMetCAU],[MetRS_MettRNAMetCAU], k_forward = rxn_k['re0000000185_k1']),

                        Reaction.from_massaction([MetRS_tRNAMetCAU,Met],[MetRS_Met_tRNAMetCAU], k_forward = rxn_k['re0000000188_k1']),
                        Reaction.from_massaction([MetRS_MetAMP_PPi_tRNAMetCAU],[MetRS_MetAMP_tRNAMetCAU,PPi], k_forward = rxn_k['re0000000189_k1']),
                        Reaction.from_massaction([MetRS_Met_tRNAMetCAU],[MetRS_tRNAMetCAU,Met], k_forward = rxn_k['re0000000190_k1']),
                        Reaction.from_massaction([MetRS_tRNAMetCAU,ATP],[MetRS_ATP_tRNAMetCAU], k_forward = rxn_k['re0000000191_k1']),
                        Reaction.from_massaction([MetRS_ATP_tRNAMetCAU],[MetRS_tRNAMetCAU,ATP], k_forward = rxn_k['re0000000192_k1']),
                        Reaction.from_massaction([MetRS_ATP_tRNAMetCAU,Met],[MetRS_Met_ATP_tRNAMetCAU], k_forward = rxn_k['re0000000193_k1']),
                        Reaction.from_massaction([MetRS_Met_ATP_tRNAMetCAU],[MetRS_ATP_tRNAMetCAU,Met], k_forward = rxn_k['re0000000194_k1']),
                        Reaction.from_massaction([MetRS_Met_tRNAMetCAU,ATP],[MetRS_Met_ATP_tRNAMetCAU], k_forward = rxn_k['re0000000195_k1']),
                        Reaction.from_massaction([MetRS_Met_ATP_tRNAMetCAU],[MetRS_Met_tRNAMetCAU,ATP], k_forward = rxn_k['re0000000196_k1']),
                        Reaction.from_massaction([MetRS_Met_ATP_tRNAMetCAU],[MetRS_MetAMP_PPi_tRNAMetCAU], k_forward = rxn_k['re0000000197_k1']),

                        Reaction.from_massaction([MetRS,tRNAMetCAU],[MetRS_tRNAMetCAU], k_forward = rxn_k['re0000000199_k1']),
                        Reaction.from_massaction([MetRS_Met,tRNAMetCAU],[MetRS_Met_tRNAMetCAU], k_forward = rxn_k['re0000000200_k1']),
                        Reaction.from_massaction([MetRS_tRNAMetCAU],[MetRS,tRNAMetCAU], k_forward = rxn_k['re0000000201_k1']),
                        Reaction.from_massaction([MetRS_Met_tRNAMetCAU],[MetRS_Met,tRNAMetCAU], k_forward = rxn_k['re0000000202_k1']),
                        Reaction.from_massaction([MetRS_ATP,tRNAMetCAU],[MetRS_ATP_tRNAMetCAU], k_forward = rxn_k['re0000000203_k1']),
                        Reaction.from_massaction([MetRS_ATP_tRNAMetCAU],[MetRS_ATP,tRNAMetCAU], k_forward = rxn_k['re0000000204_k1']),
                        Reaction.from_massaction([MetRS_Met_ATP,tRNAMetCAU],[MetRS_Met_ATP_tRNAMetCAU], k_forward = rxn_k['re0000000205_k1']),
                        Reaction.from_massaction([MetRS_Met_ATP_tRNAMetCAU],[MetRS_Met_ATP,tRNAMetCAU], k_forward = rxn_k['re0000000206_k1']),
                        Reaction.from_massaction([MetRS_MetAMP_PPi,tRNAMetCAU],[MetRS_MetAMP_PPi_tRNAMetCAU], k_forward = rxn_k['re0000000207_k1']),
                        Reaction.from_massaction([MetRS_MetAMP_PPi_tRNAMetCAU],[MetRS_MetAMP_PPi,tRNAMetCAU], k_forward = rxn_k['re0000000208_k1']),
                        Reaction.from_massaction([MetRS_MetAMP,tRNAMetCAU],[MetRS_MetAMP_tRNAMetCAU], k_forward = rxn_k['re0000000209_k1']),
                        Reaction.from_massaction([MetRS_MetAMP_tRNAMetCAU],[MetRS_MetAMP,tRNAMetCAU], k_forward = rxn_k['re0000000210_k1']),

                        Reaction.from_massaction([EFTu_GTP,MettRNAMetCAU],[EFTu_GTP_MettRNAMetCAU], k_forward = rxn_k['re0000000275_k1']),
                        Reaction.from_massaction([EFTu_GTP_MettRNAMetCAU],[EFTu_GTP,MettRNAMetCAU], k_forward = rxn_k['re0000000276_k1']),

                        Reaction.from_massaction([RS50S_tRNAMetCAU_RRF_EFG_GDP],[RS50S_tRNAMetCAU_RRF,EFG_GDP], k_forward = rxn_k['re0000000913_k1']),
                        Reaction.from_massaction([RS50S_tRNAMetCAU_RRF_EFG_GDP],[RS50S_tRNAMetCAU_EFG_GDP,RRF], k_forward = rxn_k['re0000000914_k1']),
                        Reaction.from_massaction([RS50S_tRNAMetCAU_RRF_EFG_GDP],[RS50S_RRF_EFG_GDP,tRNAMetCAU], k_forward = rxn_k['re0000000915_k1']),
                        Reaction.from_massaction([RS50S_tRNAMetCAU_RRF],[RS50S_RRF,tRNAMetCAU], k_forward = rxn_k['re0000000916_k1']),
                        Reaction.from_massaction([RS50S_tRNAMetCAU_RRF],[RS50S_tRNAMetCAU,RRF], k_forward = rxn_k['re0000000917_k1']),

                        Reaction.from_massaction([RS50S_tRNAMetCAU],[tRNAMetCAU,RS50S], k_forward = rxn_k['re0000000919_k1']),
                        Reaction.from_massaction([RS50S_tRNAMetCAU_EFG_GDP],[EFG_GDP,RS50S_tRNAMetCAU], k_forward = rxn_k['re0000000920_k1']),
                        Reaction.from_massaction([RS50S_tRNAMetCAU_EFG_GDP],[RS50S_EFG_GDP,tRNAMetCAU], k_forward = rxn_k['re0000000921_k1']),
                        ]
            list_of_reactions.append(reactions_Met)
            list_species_aa.append(species_Met)
list_of_reactions=flatten(list_of_reactions)
```

#### Elongations¶

In [11]:

```
protein_lenth = '0000'
list_of_reaction_pept3 = []
list_species_el3p3=[]
for L in [l for l in range(len(x)) if l !=0]:
    #Identification of the previous amino acid
    oA=aa
    oX=xyz
    
    #Identification of the second amino acid
    aa=x[L]
    xyz= list_condon[list_AA.index(aa)]
    Spept= protein_lenth[:-len(str(L))]+str(L)
    Gpept= protein_lenth[:-len(str(L+1))]+str((L+1))
    
    #The initial step of reading the 2nd amino acid
    if L==1:
        elRS70SAxyz0002_fMet = Species('elRS70SA'+xyz+'0002_fMet')
        elRS70SAxyz0002_fMet_EFTu_GDP = Species('elRS70SA'+xyz+'0002_fMet_EFTu_GDP')
        elRS70SAxyz0002_fMettRNAfMetCAU = Species('elRS70SA'+xyz+'0002_fMettRNAfMetCAU')

        list_species_elfmet= [elRS70SAxyz0002_fMet, elRS70SAxyz0002_fMet_EFTu_GDP, elRS70SAxyz0002_fMettRNAfMetCAU]
        
        list_reactions_elfmet= [
            Reaction.from_massaction([elRS70SAxyz0002_fMettRNAfMetCAU],[elRS70SAxyz0002_fMet,tRNAfMetCAU], k_forward=rxn_k['re0000000001_k1']),
            # Reaction.from_massaction([elRS70SAxyz0002_fMet,tRNAfMetCAU],[elRS70SAxyz0002_fMettRNAfMetCAU], k_forward= rxn_k['re0000000002_k1']),
            # Reaction.from_massaction([elRS70SAxyz0002_fMettRNAfMetCAU],[RS50S_degraded,mRNA,RS30S,fMettRNAfMetCAU], k_forward= rxn_k['re0000000009_k1']),
            # Reaction.from_massaction([elRS70SAxyz0002_fMettRNAfMetCAU],[RS30S_degraded,RS50S,mRNA,fMettRNAfMetCAU], k_forward= rxn_k['re0000000010_k1']),
            # Reaction.from_massaction([elRS70SAxyz0002_fMet],[RS50S_degraded,mRNA,RS30S,fMet], k_forward= rxn_k['re0000000011_k1']),
            # Reaction.from_massaction([elRS70SAxyz0002_fMet],[RS30S_degraded,RS50S,mRNA,fMet], k_forward= rxn_k['re0000000012_k1']),
            Reaction.from_massaction([elRS70SAxyz0002_fMet_EFTu_GDP],[elRS70SAxyz0002_fMet,EFTu_GDP], k_forward= rxn_k['re0000000028_k1']),
            # Reaction.from_massaction([elRS70SAxyz0002_fMet_EFTu_GDP],[RS50S_degraded,mRNA,RS30S,fMet,EFTu,GDP], k_forward= rxn_k['re0000000042_k1']),
            # Reaction.from_massaction([elRS70SAxyz0002_fMet_EFTu_GDP],[RS30S_degraded,RS50S,mRNA,fMet,EFTu,GDP], k_forward= rxn_k['re0000000043_k1']),
            # Reaction.from_massaction([elRS70SAxyz0002_fMet_EFTu_GDP],[EFTu_degraded,fMet,mRNA,RS30S,RS50S,GDP], k_forward= rxn_k['re0000000044_k1']),
            # Reaction.from_massaction([elRS70SAxyz0002_fMet,EFTu_GDP],[elRS70SAxyz0002_fMet_EFTu_GDP], k_forward= rxn_k['re0000000065_k1']),
            Reaction.from_massaction([RS70S_IF2_GDP_fMettRNAfMetCAU_mRNA],[elRS70SAxyz0002_fMettRNAfMetCAU,IF2_GDP], k_forward= rxn_k['re0000000726_k1']),
            Reaction.from_massaction([elRS70SAxyz0002_fMettRNAfMetCAU,IF2_GDP],[RS70S_IF2_GDP_fMettRNAfMetCAU_mRNA], k_forward= rxn_k['re0000000752_k1']),
            Reaction.from_massaction([RS70S_IF3_fMettRNAfMetCAU_mRNA],[elRS70SAxyz0002_fMettRNAfMetCAU,IF3], k_forward= rxn_k['re0000000763_k1']),
            # Reaction.from_massaction([elRS70SAxyz0002_fMettRNAfMetCAU,IF3],[RS70S_IF3_fMettRNAfMetCAU_mRNA], k_forward= rxn_k['re0000000764_k1']),
            Reaction.from_massaction([RS70S_IF1_fMettRNAfMetCAU_mRNA],[elRS70SAxyz0002_fMettRNAfMetCAU,IF1], k_forward= rxn_k['re0000000765_k1']),
            # Reaction.from_massaction([elRS70SAxyz0002_fMettRNAfMetCAU,IF1],[RS70S_IF1_fMettRNAfMetCAU_mRNA], k_forward= rxn_k['re0000000766_k1']),
            ]
    
        # Incorporation of the 2nd amino acid
        ## A2_fmet
        elRS70SAxyz0002_fMet_EFTu_GDP_aatRNAaaxyz = Species('elRS70SA'+xyz+'0002_fMet_EFTu_GDP_'+aa+'tRNA'+aa+xyz)
        elRS70SAxyz0002_fMet_EFTu_GDP_PO4_aatRNAaaxyz = Species('elRS70SA'+xyz+'0002_fMet_EFTu_GDP_PO4_'+aa+'tRNA'+aa+xyz)
        elRS70SAxyz0002_fMet_EFTu_GTP_aatRNAaaxyz = Species('elRS70SA'+xyz+'0002_fMet_EFTu_GTP_'+aa+'tRNA'+aa+xyz)
        elRS70SAxyz0002_fMet_aatRNAaaxyz = Species('elRS70SA'+xyz+'0002_fMet_'+aa+'tRNA'+aa+xyz)

        ## B2_pept2
        elRS70SBxyz0002_Pept0002tRNAaaxyz = Species('elRS70SB'+xyz+'0002_Pept0002tRNA'+aa+xyz)
        elRS70SBxyz0002_Pept0002tRNAaaxyz_EFG_GDP_PO4 = Species('elRS70SB'+xyz+'0002_Pept0002tRNA'+aa+xyz+'_EFG_GDP_PO4')
        elRS70SBxyz0002_Pept0002tRNAaaxyz_EFG_GTP = Species('elRS70SB'+xyz+'0002_Pept0002tRNA'+aa+xyz+'_EFG_GTP')     

        Pept0002 = Species('Pept0002')
        Pept0002_degraded = Species('Pept0002_degraded')
        Pept0002tRNAaaxyz = Species('Pept0002tRNA'+aa+xyz)
        Pept0002tRNAaaxyz_degraded = Species('Pept0002tRNA'+aa+xyz+'_degraded')

        list_species_el2p2= [
            elRS70SAxyz0002_fMet_EFTu_GDP_aatRNAaaxyz, elRS70SAxyz0002_fMet_EFTu_GDP_PO4_aatRNAaaxyz, 
            elRS70SAxyz0002_fMet_EFTu_GTP_aatRNAaaxyz, elRS70SAxyz0002_fMet_aatRNAaaxyz,

            elRS70SBxyz0002_Pept0002tRNAaaxyz, elRS70SBxyz0002_Pept0002tRNAaaxyz_EFG_GDP_PO4,
            elRS70SBxyz0002_Pept0002tRNAaaxyz_EFG_GTP,  

            Pept0002, Pept0002_degraded, Pept0002tRNAaaxyz, Pept0002tRNAaaxyz_degraded]

        ## Reactions
        list_of_reaction_pept2=[
            Reaction.from_massaction([Species('EFTu_GTP_'+aa+'tRNA'+aa+xyz),elRS70SAxyz0002_fMet],[elRS70SAxyz0002_fMet_EFTu_GTP_aatRNAaaxyz], k_forward= rxn_k['re0000000013_k1']),
            Reaction.from_massaction([elRS70SAxyz0002_fMet_EFTu_GTP_aatRNAaaxyz],[elRS70SAxyz0002_fMet_EFTu_GDP_PO4_aatRNAaaxyz], k_forward= rxn_k['re0000000014_k1']),
            # Reaction.from_massaction([elRS70SAxyz0002_fMet_EFTu_GDP_PO4_aatRNAaaxyz],[elRS70SAxyz0002_fMet_EFTu_GTP_aatRNAaaxyz], k_forward= rxn_k['re0000000015_k1']),
            Reaction.from_massaction([elRS70SAxyz0002_fMet_EFTu_GDP_PO4_aatRNAaaxyz],[PO4,elRS70SAxyz0002_fMet_EFTu_GDP_aatRNAaaxyz], k_forward= rxn_k['re0000000016_k1']),
            Reaction.from_massaction([elRS70SAxyz0002_fMet_EFTu_GDP_aatRNAaaxyz],[EFTu_GDP,elRS70SAxyz0002_fMet_aatRNAaaxyz], k_forward= rxn_k['re0000000017_k1']),

            Reaction.from_massaction([elRS70SAxyz0002_fMet_aatRNAaaxyz],[elRS70SBxyz0002_Pept0002tRNAaaxyz], k_forward= rxn_k['re0000000018_k1']),
            Reaction.from_massaction([elRS70SBxyz0002_Pept0002tRNAaaxyz,EFG_GTP],[elRS70SBxyz0002_Pept0002tRNAaaxyz_EFG_GTP], k_forward= rxn_k['re0000000019_k1']),
            Reaction.from_massaction([elRS70SBxyz0002_Pept0002tRNAaaxyz_EFG_GTP],[EFG_GTP,elRS70SBxyz0002_Pept0002tRNAaaxyz], k_forward= rxn_k['re0000000020_k1']),
            Reaction.from_massaction([elRS70SAxyz0002_fMet_EFTu_GTP_aatRNAaaxyz],[Species('EFTu_GTP_'+aa+'tRNA'+aa+xyz),elRS70SAxyz0002_fMet], k_forward= rxn_k['re0000000021_k1']),
            Reaction.from_massaction([elRS70SBxyz0002_Pept0002tRNAaaxyz_EFG_GTP],[elRS70SBxyz0002_Pept0002tRNAaaxyz_EFG_GDP_PO4], k_forward= rxn_k['re0000000022_k1']),
            Reaction.from_massaction([elRS70SBxyz0002_Pept0002tRNAaaxyz_EFG_GDP_PO4],[elRS70SBxyz0002_Pept0002tRNAaaxyz_EFG_GTP], k_forward= rxn_k['re0000000023_k1']),
            # Reaction.from_massaction([elRS70SAxyz0002_fMet_EFTu_GDP_aatRNAaaxyz],[elRS70SAxyz0002_fMet_EFTu_GDP,Species(aa+'tRNA'+aa+xyz)], k_forward= rxn_k['re0000000026_k1']),
            # Reaction.from_massaction([elRS70SAxyz0002_fMet_aatRNAaaxyz],[elRS70SAxyz0002_fMet,Species(aa+'tRNA'+aa+xyz)], k_forward= rxn_k['re0000000027_k1']),

            # Reaction.from_massaction([Pept0002tRNAaaxyz],[Pept0002tRNAaaxyz_degraded], k_forward= rxn_k['re0000000032_k1']),
            # Reaction.from_massaction([elRS70SAxyz0002_fMet_EFTu_GTP_aatRNAaaxyz],[RS50S_degraded,mRNA,RS30S,fMet,Species(aa+'tRNA'+aa+xyz),EFTu,GTP], k_forward= rxn_k['re0000000033_k1']),
            # Reaction.from_massaction([elRS70SAxyz0002_fMet_EFTu_GTP_aatRNAaaxyz],[RS30S_degraded,RS50S,mRNA,fMet,Species(aa+'tRNA'+aa+xyz),EFTu,GTP], k_forward= rxn_k['re0000000034_k1']),
            # Reaction.from_massaction([elRS70SAxyz0002_fMet_EFTu_GTP_aatRNAaaxyz],[EFTu_degraded,fMet,Species(aa+'tRNA'+aa+xyz),mRNA,RS30S,RS50S,GTP], k_forward= rxn_k['re0000000035_k1']),
            # Reaction.from_massaction([elRS70SAxyz0002_fMet_EFTu_GDP_PO4_aatRNAaaxyz],[RS50S_degraded,mRNA,RS30S,fMet,Species(aa+'tRNA'+aa+xyz),EFTu,GDP,PO4], k_forward= rxn_k['re0000000036_k1']),
            # Reaction.from_massaction([elRS70SAxyz0002_fMet_EFTu_GDP_PO4_aatRNAaaxyz],[RS30S_degraded,RS50S,mRNA,fMet,Species(aa+'tRNA'+aa+xyz),EFTu,GDP,PO4], k_forward= rxn_k['re0000000037_k1']),
            # Reaction.from_massaction([elRS70SAxyz0002_fMet_EFTu_GDP_PO4_aatRNAaaxyz],[EFTu_degraded,fMet,Species(aa+'tRNA'+aa+xyz),mRNA,RS30S,RS50S,GDP,PO4], k_forward= rxn_k['re0000000038_k1']),
            # Reaction.from_massaction([elRS70SAxyz0002_fMet_EFTu_GDP_aatRNAaaxyz],[RS50S_degraded,mRNA,RS30S,fMet,Species(aa+'tRNA'+aa+xyz),EFTu,GDP], k_forward= rxn_k['re0000000039_k1']),
            # Reaction.from_massaction([elRS70SAxyz0002_fMet_EFTu_GDP_aatRNAaaxyz],[RS30S_degraded,RS50S,mRNA,fMet,Species(aa+'tRNA'+aa+xyz),EFTu,GDP], k_forward= rxn_k['re0000000040_k1']),
            # Reaction.from_massaction([elRS70SAxyz0002_fMet_EFTu_GDP_aatRNAaaxyz],[EFTu_degraded,fMet,Species(aa+'tRNA'+aa+xyz),mRNA,RS30S,RS50S,GDP], k_forward= rxn_k['re0000000041_k1']),

            # Reaction.from_massaction([elRS70SAxyz0002_fMet_aatRNAaaxyz],[RS50S_degraded,mRNA,RS30S,fMet,Species(aa+'tRNA'+aa+xyz)], k_forward= rxn_k['re0000000045_k1']),
            # Reaction.from_massaction([elRS70SAxyz0002_fMet_aatRNAaaxyz],[RS30S_degraded,RS50S,mRNA,fMet,Species(aa+'tRNA'+aa+xyz)], k_forward= rxn_k['re0000000046_k1']),
            # Reaction.from_massaction([elRS70SBxyz0002_Pept0002tRNAaaxyz],[RS50S_degraded,mRNA,RS30S,Pept0002tRNAaaxyz], k_forward= rxn_k['re0000000047_k1']),
            # Reaction.from_massaction([elRS70SBxyz0002_Pept0002tRNAaaxyz],[RS30S_degraded,RS50S,mRNA,Pept0002tRNAaaxyz], k_forward= rxn_k['re0000000048_k1']),
            # Reaction.from_massaction([elRS70SBxyz0002_Pept0002tRNAaaxyz_EFG_GTP],[RS50S_degraded,mRNA,RS30S,Pept0002tRNAaaxyz,EFG,GTP], k_forward= rxn_k['re0000000049_k1']),
            # Reaction.from_massaction([elRS70SBxyz0002_Pept0002tRNAaaxyz_EFG_GTP],[RS30S_degraded,RS50S,mRNA,Pept0002tRNAaaxyz,EFG,GTP], k_forward= rxn_k['re0000000050_k1']),
            # Reaction.from_massaction([elRS70SBxyz0002_Pept0002tRNAaaxyz_EFG_GTP],[EFG_degraded,GTP,RS50S,RS30S,mRNA,Pept0002tRNAaaxyz], k_forward= rxn_k['re0000000051_k1']),

            # Reaction.from_massaction([elRS70SBxyz0002_Pept0002tRNAaaxyz_EFG_GDP_PO4],[RS50S_degraded,mRNA,RS30S,Pept0002tRNAaaxyz,EFG,GDP,PO4], k_forward= rxn_k['re0000000055_k1']),
            # Reaction.from_massaction([elRS70SBxyz0002_Pept0002tRNAaaxyz_EFG_GDP_PO4],[RS30S_degraded,RS50S,mRNA,Pept0002tRNAaaxyz,EFG,GDP,PO4], k_forward= rxn_k['re0000000056_k1']),
            # Reaction.from_massaction([elRS70SBxyz0002_Pept0002tRNAaaxyz_EFG_GDP_PO4],[EFG_degraded,GDP,RS50S,RS30S,mRNA,Pept0002tRNAaaxyz,PO4], k_forward= rxn_k['re0000000057_k1']),
            # Reaction.from_massaction([elRS70SAxyz0002_fMet_EFTu_GDP_aatRNAaaxyz,PO4],[elRS70SAxyz0002_fMet_EFTu_GDP_PO4_aatRNAaaxyz], k_forward= rxn_k['re0000000060_k1']),
            # Reaction.from_massaction([EFTu_GDP,elRS70SAxyz0002_fMet_aatRNAaaxyz],[elRS70SAxyz0002_fMet_EFTu_GDP_aatRNAaaxyz], k_forward= rxn_k['re0000000061_k1']),
            # Reaction.from_massaction([elRS70SBxyz0002_Pept0002tRNAaaxyz],[elRS70SAxyz0002_fMet_aatRNAaaxyz], k_forward= rxn_k['re0000000062_k1']),
            # Reaction.from_massaction([elRS70SAxyz0002_fMet_EFTu_GDP,Species(aa+'tRNA'+aa+xyz)],[elRS70SAxyz0002_fMet_EFTu_GDP_aatRNAaaxyz], k_forward= rxn_k['re0000000063_k1']),
            # Reaction.from_massaction([elRS70SAxyz0002_fMet,Species(aa+'tRNA'+aa+xyz)],[elRS70SAxyz0002_fMet_aatRNAaaxyz], k_forward= rxn_k['re0000000064_k1']),
            # Reaction.from_massaction([Pept0002],[Pept0002_degraded], k_forward= rxn_k['re0000000071_k1']),
            ]
    
    #The reading the 3rd+ amino acid to the second to last amino acid
    elif L<len(x):

        elRS70SCxyzGpept_PeptSpepttRNAoAoX_EFG_GDP = Species('elRS70SC'+xyz+Gpept+'_Pept'+Spept+'tRNA'+oA+oX+'_EFG_GDP')
        
        elRS70SAxyzGpept_PeptSpept = Species('elRS70SA'+xyz+Gpept+'_Pept'+Spept)
        elRS70SAxyzGpept_PeptSpept_EFTu_GDP = Species('elRS70SA'+xyz+Gpept+'_Pept'+Spept+'_EFTu_GDP')
        elRS70SAxyzGpept_PeptSpept_EFTu_GDP_aatRNAaaxyz = Species('elRS70SA'+xyz+Gpept+'_Pept'+Spept+'_EFTu_GDP_'+aa+'tRNA'+aa+xyz)
        elRS70SAxyzGpept_PeptSpept_EFTu_GDP_PO4_aatRNAaaxyz = Species('elRS70SA'+xyz+Gpept+'_Pept'+Spept+'_EFTu_GDP_PO4_'+aa+'tRNA'+aa+xyz)
        elRS70SAxyzGpept_PeptSpept_EFTu_GTP_aatRNAaaxyz = Species('elRS70SA'+xyz+Gpept+'_Pept'+Spept+'_EFTu_GTP_'+aa+'tRNA'+aa+xyz)
        elRS70SAxyzGpept_PeptSpept_aatRNAaaxyz = Species('elRS70SA'+xyz+Gpept+'_Pept'+Spept+'_'+aa+'tRNA'+aa+xyz)
        elRS70SAxyzGpept_PeptSpepttRNAoAoX= Species('elRS70SA'+xyz+Gpept+'_Pept'+Spept+'tRNA'+oA+oX)

        elRS70SBxyzGpept_PeptGpepttRNAaaxyz_EFG_GTP = Species('elRS70SB'+xyz+Gpept+'_Pept'+Gpept+'tRNA'+aa+xyz+'_EFG_GTP')
        elRS70SBxyzGpept_PeptGpepttRNAaaxyz = Species('elRS70SB'+xyz+Gpept+'_Pept'+Gpept+'tRNA'+aa+xyz)
        elRS70SBxyzGpept_PeptGpepttRNAaaxyz_EFG_GDP_PO4 = Species('elRS70SB'+xyz+Gpept+'_Pept'+Gpept+'tRNA'+aa+xyz+'_EFG_GDP_PO4')

        PeptGpept = Species('Pept'+Gpept)
        PeptGpepttRNAaaxyz = Species('Pept'+Gpept+'tRNA'+aa+xyz)
        PeptGpepttRNAaaxyz_degraded = Species('Pept'+Gpept+'tRNA'+aa+xyz+'_degraded')

        #List of Species for peptide 3+ through n-1
        species_el3p3= [
            elRS70SCxyzGpept_PeptSpepttRNAoAoX_EFG_GDP, elRS70SAxyzGpept_PeptSpept, 
            
            elRS70SAxyzGpept_PeptSpept_EFTu_GDP, elRS70SAxyzGpept_PeptSpept_EFTu_GDP_aatRNAaaxyz, 
            elRS70SAxyzGpept_PeptSpept_EFTu_GDP_PO4_aatRNAaaxyz, elRS70SAxyzGpept_PeptSpept_EFTu_GTP_aatRNAaaxyz, 
            elRS70SAxyzGpept_PeptSpept_aatRNAaaxyz, elRS70SAxyzGpept_PeptSpepttRNAoAoX,

            elRS70SBxyzGpept_PeptGpepttRNAaaxyz_EFG_GTP, elRS70SBxyzGpept_PeptGpepttRNAaaxyz,
            elRS70SBxyzGpept_PeptGpepttRNAaaxyz_EFG_GDP_PO4,

            PeptGpept, PeptGpepttRNAaaxyz, PeptGpepttRNAaaxyz_degraded,
            ]
        #List of reactions for peptide 3+ through n-1
        rxn_pept3=[
            Reaction.from_massaction([Species('elRS70SB'+oX+Spept+'_Pept'+Spept+'tRNA'+oA+oX+'_EFG_GDP_PO4')],[PO4,elRS70SCxyzGpept_PeptSpepttRNAoAoX_EFG_GDP], k_forward= rxn_k['re0000000024_k1']),
            Reaction.from_massaction([elRS70SCxyzGpept_PeptSpepttRNAoAoX_EFG_GDP],[elRS70SAxyzGpept_PeptSpepttRNAoAoX,EFG_GDP], k_forward= rxn_k['re0000000025_k1']),
            # Reaction.from_massaction([elRS70SCxyzGpept_PeptSpepttRNAoAoX_EFG_GDP],[RS50S_degraded,mRNA,RS30S,Species('Pept'+Spept+'tRNA'+oA+oX),EFG,GDP], k_forward= rxn_k['re0000000052_k1']),
            # Reaction.from_massaction([elRS70SCxyzGpept_PeptSpepttRNAoAoX_EFG_GDP],[RS30S_degraded,RS50S,mRNA,Species('Pept'+Spept+'tRNA'+oA+oX),EFG,GDP], k_forward= rxn_k['re0000000053_k1']),
            # Reaction.from_massaction([elRS70SCxyzGpept_PeptSpepttRNAoAoX_EFG_GDP],[EFG_degraded,GDP,RS50S,RS30S,mRNA,Species('Pept'+Spept+'tRNA'+oA+oX)], k_forward= rxn_k['re0000000054_k1']),
            # Reaction.from_massaction([PO4,elRS70SCxyzGpept_PeptSpepttRNAoAoX_EFG_GDP],[Species('elRS70SB'+xyz+Spept+'_Pept'+Spept+'tRNA'+oA+oX+'_EFG_GDP_PO4')], k_forward= rxn_k['re0000000066_k1']),
            
            # Reaction.from_massaction([elRS70SAxyzGpept_PeptSpepttRNAoAoX],[RS50S_degraded,mRNA,RS30S,Species('Pept'+Spept+'tRNA'+oA+oX)], k_forward= rxn_k['re0000000058_k1']),
            # Reaction.from_massaction([elRS70SAxyzGpept_PeptSpepttRNAoAoX],[RS30S_degraded,RS50S,mRNA,Species('Pept'+Spept+'tRNA'+oA+oX)], k_forward= rxn_k['re0000000059_k1']),
            # Reaction.from_massaction([elRS70SAxyzGpept_PeptSpepttRNAoAoX,EFG_GDP],[elRS70SCxyzGpept_PeptSpepttRNAoAoX_EFG_GDP], k_forward= rxn_k['re0000000067_k1']),
            Reaction.from_massaction([elRS70SAxyzGpept_PeptSpepttRNAoAoX],[elRS70SAxyzGpept_PeptSpept,Species('tRNA'+oA+oX)], k_forward= rxn_k['re0000000068_k1']),
            # Reaction.from_massaction([elRS70SAxyzGpept_PeptSpept,Species('tRNA'+oA+oX)],[elRS70SAxyzGpept_PeptSpepttRNAoAoX], k_forward= rxn_k['re0000000069_k1']),
            #Switched to the new aa
            # Reaction.from_massaction([elRS70SAxyzGpept_PeptSpept],[RS50S_degraded,mRNA,RS30S,Species('Pept'+Spept)], k_forward= rxn_k['re0000000072_k1']),
            # Reaction.from_massaction([elRS70SAxyzGpept_PeptSpept],[RS30S_degraded,RS50S,mRNA,Species('Pept'+Spept)], k_forward= rxn_k['re0000000073_k1']),
            Reaction.from_massaction([Species('EFTu_GTP_'+aa+'tRNA'+aa+xyz),elRS70SAxyzGpept_PeptSpept],[elRS70SAxyzGpept_PeptSpept_EFTu_GTP_aatRNAaaxyz], k_forward= rxn_k['re0000000074_k1']),
            
            Reaction.from_massaction([elRS70SAxyzGpept_PeptSpept_EFTu_GTP_aatRNAaaxyz],[elRS70SAxyzGpept_PeptSpept_EFTu_GDP_PO4_aatRNAaaxyz], k_forward= rxn_k['re0000000075_k1']),
            # Reaction.from_massaction([elRS70SAxyzGpept_PeptSpept_EFTu_GDP_PO4_aatRNAaaxyz],[elRS70SAxyzGpept_PeptSpept_EFTu_GTP_aatRNAaaxyz], k_forward= rxn_k['re0000000076_k1']),
            Reaction.from_massaction([elRS70SAxyzGpept_PeptSpept_EFTu_GDP_PO4_aatRNAaaxyz],[PO4,elRS70SAxyzGpept_PeptSpept_EFTu_GDP_aatRNAaaxyz], k_forward= rxn_k['re0000000077_k1']),
            Reaction.from_massaction([elRS70SAxyzGpept_PeptSpept_EFTu_GDP_aatRNAaaxyz],[EFTu_GDP,elRS70SAxyzGpept_PeptSpept_aatRNAaaxyz], k_forward= rxn_k['re0000000078_k1']),
            Reaction.from_massaction([elRS70SAxyzGpept_PeptSpept_aatRNAaaxyz],[elRS70SBxyzGpept_PeptGpepttRNAaaxyz], k_forward= rxn_k['re0000000079_k1']),

            Reaction.from_massaction([elRS70SBxyzGpept_PeptGpepttRNAaaxyz,EFG_GTP],[elRS70SBxyzGpept_PeptGpepttRNAaaxyz_EFG_GTP], k_forward= rxn_k['re0000000080_k1']),
            Reaction.from_massaction([elRS70SBxyzGpept_PeptGpepttRNAaaxyz_EFG_GTP],[EFG_GTP,elRS70SBxyzGpept_PeptGpepttRNAaaxyz], k_forward= rxn_k['re0000000081_k1']),
            Reaction.from_massaction([elRS70SAxyzGpept_PeptSpept_EFTu_GTP_aatRNAaaxyz],[Species('EFTu_GTP_'+aa+'tRNA'+aa+xyz),elRS70SAxyzGpept_PeptSpept], k_forward= rxn_k['re0000000082_k1']),
            Reaction.from_massaction([elRS70SBxyzGpept_PeptGpepttRNAaaxyz_EFG_GTP],[elRS70SBxyzGpept_PeptGpepttRNAaaxyz_EFG_GDP_PO4], k_forward= rxn_k['re0000000083_k1']),
            Reaction.from_massaction([elRS70SBxyzGpept_PeptGpepttRNAaaxyz_EFG_GDP_PO4],[elRS70SBxyzGpept_PeptGpepttRNAaaxyz_EFG_GTP], k_forward= rxn_k['re0000000084_k1']),
            # Reaction.from_massaction([elRS70SAxyzGpept_PeptSpept_EFTu_GDP_aatRNAaaxyz],[elRS70SAxyzGpept_PeptSpept_EFTu_GDP,Species(aa+'tRNA'+aa+xyz)], k_forward= rxn_k['re0000000087_k1']),
            # Reaction.from_massaction([elRS70SAxyzGpept_PeptSpept_aatRNAaaxyz],[elRS70SAxyzGpept_PeptSpept,Species(aa+'tRNA'+aa+xyz)], k_forward= rxn_k['re0000000088_k1']),
            Reaction.from_massaction([elRS70SAxyzGpept_PeptSpept_EFTu_GDP],[elRS70SAxyzGpept_PeptSpept,EFTu_GDP], k_forward= rxn_k['re0000000089_k1']),
            # Reaction.from_massaction([PeptGpepttRNAaaxyz],[PeptGpepttRNAaaxyz_degraded], k_forward= rxn_k['re0000000090_k1']),
            # Reaction.from_massaction([elRS70SAxyzGpept_PeptSpept_EFTu_GTP_aatRNAaaxyz],[RS50S_degraded,mRNA,RS30S,Species('Pept'+Spept),Species(aa+'tRNA'+aa+xyz),EFTu,GTP], k_forward= rxn_k['re0000000091_k1']),
            # Reaction.from_massaction([elRS70SAxyzGpept_PeptSpept_EFTu_GTP_aatRNAaaxyz],[RS30S_degraded,RS50S,mRNA,Species('Pept'+Spept),Species(aa+'tRNA'+aa+xyz),EFTu,GTP], k_forward= rxn_k['re0000000092_k1']),
            # Reaction.from_massaction([elRS70SAxyzGpept_PeptSpept_EFTu_GTP_aatRNAaaxyz],[EFTu_degraded,Species('Pept'+Spept),Species(aa+'tRNA'+aa+xyz),mRNA,RS30S,RS50S,GTP], k_forward= rxn_k['re0000000093_k1']),
            # Reaction.from_massaction([elRS70SAxyzGpept_PeptSpept_EFTu_GDP_PO4_aatRNAaaxyz],[RS50S_degraded,mRNA,RS30S,Species('Pept'+Spept),Species(aa+'tRNA'+aa+xyz),EFTu,GDP,PO4], k_forward= rxn_k['re0000000094_k1']),
            # Reaction.from_massaction([elRS70SAxyzGpept_PeptSpept_EFTu_GDP_PO4_aatRNAaaxyz],[RS30S_degraded,RS50S,mRNA,Species('Pept'+Spept),Species(aa+'tRNA'+aa+xyz),EFTu,GDP,PO4], k_forward= rxn_k['re0000000095_k1']),
            # Reaction.from_massaction([elRS70SAxyzGpept_PeptSpept_EFTu_GDP_PO4_aatRNAaaxyz],[EFTu_degraded,Species('Pept'+Spept),Species(aa+'tRNA'+aa+xyz),mRNA,RS30S,RS50S,GDP,PO4], k_forward= rxn_k['re0000000096_k1']),
            # Reaction.from_massaction([elRS70SAxyzGpept_PeptSpept_EFTu_GDP_aatRNAaaxyz],[RS50S_degraded,mRNA,RS30S,Species('Pept'+Spept),Species(aa+'tRNA'+aa+xyz),EFTu,GDP], k_forward= rxn_k['re0000000097_k1']),
            # Reaction.from_massaction([elRS70SAxyzGpept_PeptSpept_EFTu_GDP_aatRNAaaxyz],[RS30S_degraded,RS50S,mRNA,Species('Pept'+Spept),Species(aa+'tRNA'+aa+xyz),EFTu,GDP], k_forward= rxn_k['re0000000098_k1']),
            # Reaction.from_massaction([elRS70SAxyzGpept_PeptSpept_EFTu_GDP_aatRNAaaxyz],[EFTu_degraded,Species('Pept'+Spept),Species(aa+'tRNA'+aa+xyz),mRNA,RS30S,RS50S,GDP], k_forward= rxn_k['re0000000099_k1']),
            # Reaction.from_massaction([elRS70SAxyzGpept_PeptSpept_EFTu_GDP],[RS50S_degraded,mRNA,RS30S,Species('Pept'+Spept),EFTu,GDP], k_forward= rxn_k['re0000000100_k1']),
            # Reaction.from_massaction([elRS70SAxyzGpept_PeptSpept_EFTu_GDP],[RS30S_degraded,RS50S,mRNA,Species('Pept'+Spept),EFTu,GDP], k_forward= rxn_k['re0000000101_k1']),

            # Reaction.from_massaction([elRS70SAxyzGpept_PeptSpept_EFTu_GDP],[EFTu_degraded,Species('Pept'+Spept),mRNA,RS30S,RS50S,GDP], k_forward= rxn_k['re0000000102_k1']),
            # Reaction.from_massaction([elRS70SAxyzGpept_PeptSpept_aatRNAaaxyz],[RS50S_degraded,mRNA,RS30S,Species('Pept'+Spept),Species(aa+'tRNA'+aa+xyz)], k_forward= rxn_k['re0000000103_k1']),
            # Reaction.from_massaction([elRS70SAxyzGpept_PeptSpept_aatRNAaaxyz],[RS30S_degraded,RS50S,mRNA,Species('Pept'+Spept),Species(aa+'tRNA'+aa+xyz)], k_forward= rxn_k['re0000000104_k1']),
            # Reaction.from_massaction([elRS70SBxyzGpept_PeptGpepttRNAaaxyz],[RS50S_degraded,mRNA,RS30S,PeptGpepttRNAaaxyz], k_forward= rxn_k['re0000000105_k1']),
            # Reaction.from_massaction([elRS70SBxyzGpept_PeptGpepttRNAaaxyz],[RS30S_degraded,RS50S,mRNA,PeptGpepttRNAaaxyz], k_forward= rxn_k['re0000000106_k1']),
            # Reaction.from_massaction([elRS70SBxyzGpept_PeptGpepttRNAaaxyz_EFG_GTP],[RS50S_degraded,mRNA,RS30S,PeptGpepttRNAaaxyz,EFG,GTP], k_forward= rxn_k['re0000000107_k1']),
            # Reaction.from_massaction([elRS70SBxyzGpept_PeptGpepttRNAaaxyz_EFG_GTP],[RS30S_degraded,RS50S,mRNA,PeptGpepttRNAaaxyz,EFG,GTP], k_forward= rxn_k['re0000000108_k1']),
            # Reaction.from_massaction([elRS70SBxyzGpept_PeptGpepttRNAaaxyz_EFG_GTP],[EFG_degraded,GTP,RS50S,RS30S,mRNA,PeptGpepttRNAaaxyz], k_forward= rxn_k['re0000000109_k1']),

            # Reaction.from_massaction([elRS70SBxyzGpept_PeptGpepttRNAaaxyz_EFG_GDP_PO4],[RS50S_degraded,mRNA,RS30S,PeptGpepttRNAaaxyz,EFG,GDP,PO4], k_forward= rxn_k['re0000000113_k1']),
            # Reaction.from_massaction([elRS70SBxyzGpept_PeptGpepttRNAaaxyz_EFG_GDP_PO4],[RS30S_degraded,RS50S,mRNA,PeptGpepttRNAaaxyz,EFG,GDP,PO4], k_forward= rxn_k['re0000000114_k1']),
            # Reaction.from_massaction([elRS70SBxyzGpept_PeptGpepttRNAaaxyz_EFG_GDP_PO4],[EFG_degraded,GDP,RS50S,RS30S,mRNA,PeptGpepttRNAaaxyz,PO4], k_forward= rxn_k['re0000000115_k1']),
            # Reaction.from_massaction([elRS70SAxyzGpept_PeptSpept_EFTu_GDP_aatRNAaaxyz,PO4],[elRS70SAxyzGpept_PeptSpept_EFTu_GDP_PO4_aatRNAaaxyz], k_forward= rxn_k['re0000000118_k1']),
            # Reaction.from_massaction([EFTu_GDP,elRS70SAxyzGpept_PeptSpept_aatRNAaaxyz],[elRS70SAxyzGpept_PeptSpept_EFTu_GDP_aatRNAaaxyz], k_forward= rxn_k['re0000000119_k1']),
            # Reaction.from_massaction([elRS70SBxyzGpept_PeptGpepttRNAaaxyz],[elRS70SAxyzGpept_PeptSpept_aatRNAaaxyz], k_forward= rxn_k['re0000000120_k1']),
            # Reaction.from_massaction([elRS70SAxyzGpept_PeptSpept_EFTu_GDP,Species(aa+'tRNA'+aa+xyz)],[elRS70SAxyzGpept_PeptSpept_EFTu_GDP_aatRNAaaxyz], k_forward= rxn_k['re0000000121_k1']),
            # Reaction.from_massaction([elRS70SAxyzGpept_PeptSpept,Species(aa+'tRNA'+aa+xyz)],[elRS70SAxyzGpept_PeptSpept_aatRNAaaxyz], k_forward= rxn_k['re0000000122_k1']),
            # Reaction.from_massaction([elRS70SAxyzGpept_PeptSpept,EFTu_GDP],[elRS70SAxyzGpept_PeptSpept_EFTu_GDP], k_forward= rxn_k['re0000000123_k1']),
            ]
        list_of_reaction_pept3.append(rxn_pept3)
        list_species_el3p3.append(species_el3p3)
    else:
        print('error')
        
#Add reaction to change the aa and xyz for the next number
list_of_reaction_pept3=flatten(list_of_reaction_pept3)
list_species_el3p3=flatten(list_species_el3p3)
```

#### Termination¶

Reading of the Stop codon and initiation of separation of complete mRNA strand

In [12]:

```
# Incorporation of the 2nd to last amino acid, including TAA
Lpept= protein_lenth[:-len(str(L))]+str((L+2))
# print(Lpept)

#######################################################################################################################
#Species
elRS70SAUAALpept_PeptGpepttRNAaaxyz = Species('elRS70SAUAA'+Lpept+'_Pept'+Gpept+'tRNA'+aa+xyz)
elRS70SAUAALpept_PeptGpepttRNAaaxyz_RF1 = Species('elRS70SAUAA'+Lpept+'_Pept'+Gpept+'tRNA'+aa+xyz+'_RF1')
elRS70SAUAALpept_PeptGpepttRNAaaxyz_RF2 = Species('elRS70SAUAA'+Lpept+'_Pept'+Gpept+'tRNA'+aa+xyz+'_RF2')
elRS70SCUAALpept_PeptGpepttRNAaaxyz_EFG_GDP = Species('elRS70SCUAA'+Lpept+'_Pept'+Gpept+'tRNA'+aa+xyz+'_EFG_GDP')

#List of Species
list_species_el4p3= [
    elRS70SAUAALpept_PeptGpepttRNAaaxyz, elRS70SAUAALpept_PeptGpepttRNAaaxyz_RF1,
    elRS70SAUAALpept_PeptGpepttRNAaaxyz_RF2, elRS70SCUAALpept_PeptGpepttRNAaaxyz_EFG_GDP]

#######################################################################################################################
#Reactions
list_of_reactions_end= [
    Reaction.from_massaction([elRS70SBxyzGpept_PeptGpepttRNAaaxyz_EFG_GDP_PO4],[PO4,elRS70SCUAALpept_PeptGpepttRNAaaxyz_EFG_GDP], k_forward= rxn_k['re0000000085_k1']),
    Reaction.from_massaction([elRS70SCUAALpept_PeptGpepttRNAaaxyz_EFG_GDP],[elRS70SAUAALpept_PeptGpepttRNAaaxyz,EFG_GDP], k_forward= rxn_k['re0000000086_k1']),

    # Reaction.from_massaction([elRS70SCUAALpept_PeptGpepttRNAaaxyz_EFG_GDP],[RS50S_degraded,mRNA,RS30S,PeptGpepttRNAaaxyz,EFG,GDP], k_forward= rxn_k['re0000000110_k1']),
    # Reaction.from_massaction([elRS70SCUAALpept_PeptGpepttRNAaaxyz_EFG_GDP],[RS30S_degraded,RS50S,mRNA,PeptGpepttRNAaaxyz,EFG,GDP], k_forward= rxn_k['re0000000111_k1']),
    # Reaction.from_massaction([elRS70SCUAALpept_PeptGpepttRNAaaxyz_EFG_GDP],[EFG_degraded,GDP,RS50S,RS30S,mRNA,PeptGpepttRNAaaxyz], k_forward= rxn_k['re0000000112_k1']),

    # Reaction.from_massaction([elRS70SAUAALpept_PeptGpepttRNAaaxyz],[RS50S_degraded,mRNA,RS30S,PeptGpepttRNAaaxyz], k_forward= rxn_k['re0000000116_k1']),
    # Reaction.from_massaction([elRS70SAUAALpept_PeptGpepttRNAaaxyz],[RS30S_degraded,RS50S,mRNA,PeptGpepttRNAaaxyz], k_forward= rxn_k['re0000000117_k1']),

    # Reaction.from_massaction([PO4,elRS70SCUAALpept_PeptGpepttRNAaaxyz_EFG_GDP],[elRS70SBxyzGpept_PeptGpepttRNAaaxyz_EFG_GDP_PO4], k_forward= rxn_k['re0000000124_k1']),
    # Reaction.from_massaction([elRS70SAUAALpept_PeptGpepttRNAaaxyz,EFG_GDP],[elRS70SCUAALpept_PeptGpepttRNAaaxyz_EFG_GDP], k_forward= rxn_k['re0000000125_k1']),

    Reaction.from_massaction([elRS70SAUAALpept_PeptGpepttRNAaaxyz,RF1],[elRS70SAUAALpept_PeptGpepttRNAaaxyz_RF1], k_forward= rxn_k['re0000000796_k1']),
    Reaction.from_massaction([elRS70SAUAALpept_PeptGpepttRNAaaxyz_RF1],[elRS70SAUAALpept_PeptGpepttRNAaaxyz,RF1], k_forward= rxn_k['re0000000797_k1']),

    # Reaction.from_massaction([elRS70SAUAALpept_PeptGpepttRNAaaxyz_RF1],[RS30S_degraded,mRNA,PeptGpepttRNAaaxyz,RS50S,RF1], k_forward= rxn_k['re0000000801_k1']),
    # Reaction.from_massaction([elRS70SAUAALpept_PeptGpepttRNAaaxyz_RF1],[RS50S_degraded,mRNA,PeptGpepttRNAaaxyz,RS30S,RF1], k_forward= rxn_k['re0000000802_k1']),
    # Reaction.from_massaction([elRS70SAUAALpept_PeptGpepttRNAaaxyz_RF1],[RF1_degraded,RS50S,RS30S,mRNA,PeptGpepttRNAaaxyz], k_forward= rxn_k['re0000000804_k1']),

    Reaction.from_massaction([elRS70SAUAALpept_PeptGpepttRNAaaxyz,RF2],[elRS70SAUAALpept_PeptGpepttRNAaaxyz_RF2], k_forward= rxn_k['re0000000811_k1']),
    Reaction.from_massaction([elRS70SAUAALpept_PeptGpepttRNAaaxyz_RF2],[elRS70SAUAALpept_PeptGpepttRNAaaxyz,RF2], k_forward= rxn_k['re0000000812_k1']),

    # Reaction.from_massaction([elRS70SAUAALpept_PeptGpepttRNAaaxyz_RF2],[RS30S_degraded,mRNA,PeptGpepttRNAaaxyz,RS50S,RF2], k_forward= rxn_k['re0000000816_k1']),
    # Reaction.from_massaction([elRS70SAUAALpept_PeptGpepttRNAaaxyz_RF2],[RS50S_degraded,mRNA,PeptGpepttRNAaaxyz,RS30S,RF2], k_forward= rxn_k['re0000000817_k1']),
    # Reaction.from_massaction([elRS70SAUAALpept_PeptGpepttRNAaaxyz_RF2],[RF2_degraded,RS50S,RS30S,mRNA,PeptGpepttRNAaaxyz], k_forward= rxn_k['re0000000819_k1']),
    ]
```

#### Removal of ribosome from finished mRNA strand¶

In [13]:

```
#Species
termRS30S_mRNA = Species('termRS30S_mRNA')
termRS70SUAALpept_tRNAaaxyz = Species('termRS70SUAA'+Lpept+'_tRNA'+aa+xyz)
termRS70SUAALpept_tRNAaaxyz_EFG_GTP = Species('termRS70SUAA'+Lpept+'_tRNA'+aa+xyz+'_EFG_GTP')
termRS70SUAALpept_tRNAaaxyz_RF1 = Species('termRS70SUAA'+Lpept+'_tRNA'+aa+xyz+'_RF1')
termRS70SUAALpept_tRNAaaxyz_RF1_RF3 = Species('termRS70SUAA'+Lpept+'_tRNA'+aa+xyz+'_RF1_RF3')
termRS70SUAALpept_tRNAaaxyz_RF1_RF3_GDP = Species('termRS70SUAA'+Lpept+'_tRNA'+aa+xyz+'_RF1_RF3_GDP')
termRS70SUAALpept_tRNAaaxyz_RF1_RF3_GTP = Species('termRS70SUAA'+Lpept+'_tRNA'+aa+xyz+'_RF1_RF3_GTP')
termRS70SUAALpept_tRNAaaxyz_RF2 = Species('termRS70SUAA'+Lpept+'_tRNA'+aa+xyz+'_RF2')
termRS70SUAALpept_tRNAaaxyz_RF2_RF3 = Species('termRS70SUAA'+Lpept+'_tRNA'+aa+xyz+'_RF2_RF3')
termRS70SUAALpept_tRNAaaxyz_RF2_RF3_GDP = Species('termRS70SUAA'+Lpept+'_tRNA'+aa+xyz+'_RF2_RF3_GDP')
termRS70SUAALpept_tRNAaaxyz_RF2_RF3_GTP = Species('termRS70SUAA'+Lpept+'_tRNA'+aa+xyz+'_RF2_RF3_GTP')
termRS70SUAALpept_tRNAaaxyz_RF3_GDP = Species('termRS70SUAA'+Lpept+'_tRNA'+aa+xyz+'_RF3_GDP')
termRS70SUAALpept_tRNAaaxyz_RF3_GDP_PO4 = Species('termRS70SUAA'+Lpept+'_tRNA'+aa+xyz+'_RF3_GDP_PO4')
termRS70SUAALpept_tRNAaaxyz_RF3_GTP = Species('termRS70SUAA'+Lpept+'_tRNA'+aa+xyz+'_RF3_GTP')
termRS70SUAALpept_tRNAaaxyz_RRF = Species('termRS70SUAA'+Lpept+'_tRNA'+aa+xyz+'_RRF')
termRS70SUAALpept_tRNAaaxyz_RRF_EFG_GDP = Species('termRS70SUAA'+Lpept+'_tRNA'+aa+xyz+'_RRF_EFG_GDP')
termRS70SUAALpept_tRNAaaxyz_RRF_EFG_GDP_PO4 = Species('termRS70SUAA'+Lpept+'_tRNA'+aa+xyz+'_RRF_EFG_GDP_PO4')
termRS70SUAALpept_tRNAaaxyz_RRF_EFG_GTP = Species('termRS70SUAA'+Lpept+'_tRNA'+aa+xyz+'_RRF_EFG_GTP')

#######################################################################################################################
#List of Species
list_species_elterm= [
    termRS30S_mRNA, termRS70SUAALpept_tRNAaaxyz,termRS70SUAALpept_tRNAaaxyz_EFG_GTP,
    termRS70SUAALpept_tRNAaaxyz_RF1, termRS70SUAALpept_tRNAaaxyz_RF1_RF3, termRS70SUAALpept_tRNAaaxyz_RF1_RF3_GDP,
    termRS70SUAALpept_tRNAaaxyz_RF1_RF3_GTP, termRS70SUAALpept_tRNAaaxyz_RF2, termRS70SUAALpept_tRNAaaxyz_RF2_RF3,
    termRS70SUAALpept_tRNAaaxyz_RF2_RF3_GDP, termRS70SUAALpept_tRNAaaxyz_RF2_RF3_GTP, termRS70SUAALpept_tRNAaaxyz_RF3_GDP,
    termRS70SUAALpept_tRNAaaxyz_RF3_GDP_PO4, termRS70SUAALpept_tRNAaaxyz_RF3_GTP, termRS70SUAALpept_tRNAaaxyz_RRF,
    termRS70SUAALpept_tRNAaaxyz_RRF_EFG_GDP, termRS70SUAALpept_tRNAaaxyz_RRF_EFG_GDP_PO4, termRS70SUAALpept_tRNAaaxyz_RRF_EFG_GTP]

#######################################################################################################################
#Reactions
list_of_reaction_term= [
    Reaction.from_massaction([elRS70SAUAALpept_PeptGpepttRNAaaxyz_RF1],[termRS70SUAALpept_tRNAaaxyz_RF1,PeptGpept], k_forward= rxn_k['re0000000798_k1']),
    Reaction.from_massaction([termRS70SUAALpept_tRNAaaxyz_RF1],[termRS70SUAALpept_tRNAaaxyz,RF1], k_forward= rxn_k['re0000000799_k1']),
    Reaction.from_massaction([termRS70SUAALpept_tRNAaaxyz,RF1],[termRS70SUAALpept_tRNAaaxyz_RF1], k_forward= rxn_k['re0000000800_k1']),
    # Reaction.from_massaction([termRS70SUAALpept_tRNAaaxyz_RF1],[RS30S_degraded,mRNA,Species('tRNA'+aa+xyz),RS50S,RF1], k_forward= rxn_k['re0000000805_k1']),
    # Reaction.from_massaction([termRS70SUAALpept_tRNAaaxyz_RF1],[RS50S_degraded,mRNA,Species('tRNA'+aa+xyz),RS30S,RF1], k_forward= rxn_k['re0000000806_k1']),
    # Reaction.from_massaction([termRS70SUAALpept_tRNAaaxyz_RF1],[RF1_degraded,RS50S,RS30S,mRNA,Species('tRNA'+aa+xyz)], k_forward= rxn_k['re0000000807_k1']),
    # Reaction.from_massaction([termRS70SUAALpept_tRNAaaxyz],[RS30S_degraded,mRNA,Species('tRNA'+aa+xyz),RS50S], k_forward= rxn_k['re0000000808_k1']),
    # Reaction.from_massaction([termRS70SUAALpept_tRNAaaxyz],[RS50S_degraded,mRNA,Species('tRNA'+aa+xyz),RS30S], k_forward= rxn_k['re0000000809_k1']),
    # Reaction.from_massaction([termRS70SUAALpept_tRNAaaxyz_RF1,PeptGpept],[elRS70SAUAALpept_PeptGpepttRNAaaxyz_RF1], k_forward= rxn_k['re0000000810_k1']),
    Reaction.from_massaction([elRS70SAUAALpept_PeptGpepttRNAaaxyz_RF2],[termRS70SUAALpept_tRNAaaxyz_RF2,PeptGpept], k_forward= rxn_k['re0000000813_k1']),

    Reaction.from_massaction([termRS70SUAALpept_tRNAaaxyz_RF2],[termRS70SUAALpept_tRNAaaxyz,RF2], k_forward= rxn_k['re0000000814_k1']),
    Reaction.from_massaction([termRS70SUAALpept_tRNAaaxyz,RF2],[termRS70SUAALpept_tRNAaaxyz_RF2], k_forward= rxn_k['re0000000815_k1']),
    # Reaction.from_massaction([termRS70SUAALpept_tRNAaaxyz_RF2],[RS30S_degraded,mRNA,Species('tRNA'+aa+xyz),RS50S,RF2], k_forward= rxn_k['re0000000820_k1']),
    # Reaction.from_massaction([termRS70SUAALpept_tRNAaaxyz_RF2],[RS50S_degraded,mRNA,Species('tRNA'+aa+xyz),RS30S,RF2], k_forward= rxn_k['re0000000821_k1']),
    # Reaction.from_massaction([termRS70SUAALpept_tRNAaaxyz_RF2],[RF2_degraded,RS50S,RS30S,mRNA,Species('tRNA'+aa+xyz)], k_forward= rxn_k['re0000000822_k1']),
    # Reaction.from_massaction([termRS70SUAALpept_tRNAaaxyz_RF2,PeptGpept],[elRS70SAUAALpept_PeptGpepttRNAaaxyz_RF2], k_forward= rxn_k['re0000000823_k1']),
    Reaction.from_massaction([RF3_GDP,termRS70SUAALpept_tRNAaaxyz_RF1],[termRS70SUAALpept_tRNAaaxyz_RF1_RF3_GDP], k_forward= rxn_k['re0000000829_k1']),
    Reaction.from_massaction([termRS70SUAALpept_tRNAaaxyz_RF1_RF3_GDP],[termRS70SUAALpept_tRNAaaxyz_RF1,RF3_GDP], k_forward= rxn_k['re0000000830_k1']),
    Reaction.from_massaction([termRS70SUAALpept_tRNAaaxyz_RF1,RF3_GTP],[termRS70SUAALpept_tRNAaaxyz_RF1_RF3_GTP], k_forward= rxn_k['re0000000831_k1']),
    Reaction.from_massaction([termRS70SUAALpept_tRNAaaxyz_RF1_RF3_GTP],[termRS70SUAALpept_tRNAaaxyz_RF1,RF3_GTP], k_forward= rxn_k['re0000000832_k1']),

    Reaction.from_massaction([termRS70SUAALpept_tRNAaaxyz_RF1,RF3],[termRS70SUAALpept_tRNAaaxyz_RF1_RF3], k_forward= rxn_k['re0000000833_k1']),
    Reaction.from_massaction([termRS70SUAALpept_tRNAaaxyz_RF1_RF3],[termRS70SUAALpept_tRNAaaxyz_RF1,RF3], k_forward= rxn_k['re0000000834_k1']),
    # Reaction.from_massaction([termRS70SUAALpept_tRNAaaxyz_RF1_RF3_GTP],[RS30S_degraded,mRNA,RF1,Species('tRNA'+aa+xyz),RS50S,RF3,GTP], k_forward= rxn_k['re0000000835_k1']),
    # Reaction.from_massaction([termRS70SUAALpept_tRNAaaxyz_RF1_RF3_GTP],[RS50S_degraded,mRNA,RF1,Species('tRNA'+aa+xyz),RS30S,RF3,GTP], k_forward= rxn_k['re0000000836_k1']),
    # Reaction.from_massaction([termRS70SUAALpept_tRNAaaxyz_RF1_RF3_GTP],[RF1_degraded,RS50S,RS30S,Species('tRNA'+aa+xyz),mRNA,GTP,RF3], k_forward= rxn_k['re0000000837_k1']),
    Reaction.from_massaction([termRS70SUAALpept_tRNAaaxyz_RF1_RF3_GTP],[termRS70SUAALpept_tRNAaaxyz_RF3_GTP,RF1], k_forward= rxn_k['re0000000838_k1']),
    Reaction.from_massaction([termRS70SUAALpept_tRNAaaxyz_RF3_GTP,RF1],[termRS70SUAALpept_tRNAaaxyz_RF1_RF3_GTP], k_forward= rxn_k['re0000000839_k1']),
    Reaction.from_massaction([termRS70SUAALpept_tRNAaaxyz_RF3_GTP],[termRS70SUAALpept_tRNAaaxyz_RF3_GDP_PO4], k_forward= rxn_k['re0000000840_k1']),
    Reaction.from_massaction([termRS70SUAALpept_tRNAaaxyz_RF3_GDP_PO4],[termRS70SUAALpept_tRNAaaxyz_RF3_GTP], k_forward= rxn_k['re0000000841_k1']),
    Reaction.from_massaction([termRS70SUAALpept_tRNAaaxyz_RF3_GDP_PO4],[termRS70SUAALpept_tRNAaaxyz_RF3_GDP,PO4], k_forward= rxn_k['re0000000842_k1']),

    Reaction.from_massaction([termRS70SUAALpept_tRNAaaxyz_RF1_RF3_GDP],[termRS70SUAALpept_tRNAaaxyz_RF1_RF3,GDP], k_forward= rxn_k['re0000000843_k1']),
    Reaction.from_massaction([termRS70SUAALpept_tRNAaaxyz_RF1_RF3,GDP],[termRS70SUAALpept_tRNAaaxyz_RF1_RF3_GDP], k_forward= rxn_k['re0000000844_k1']),
    Reaction.from_massaction([termRS70SUAALpept_tRNAaaxyz_RF1_RF3_GTP],[termRS70SUAALpept_tRNAaaxyz_RF1_RF3,GTP], k_forward= rxn_k['re0000000845_k1']),
    Reaction.from_massaction([termRS70SUAALpept_tRNAaaxyz_RF1_RF3,GTP],[termRS70SUAALpept_tRNAaaxyz_RF1_RF3_GTP], k_forward= rxn_k['re0000000846_k1']),
    Reaction.from_massaction([termRS70SUAALpept_tRNAaaxyz_RF3_GDP],[termRS70SUAALpept_tRNAaaxyz,RF3_GDP], k_forward= rxn_k['re0000000847_k1']),
    # Reaction.from_massaction([termRS70SUAALpept_tRNAaaxyz_RF1_RF3_GTP],[RF3_degraded,mRNA,Species('tRNA'+aa+xyz),RF1,RS30S,RS50S,GTP], k_forward= rxn_k['re0000000850_k1']),
    # Reaction.from_massaction([termRS70SUAALpept_tRNAaaxyz_RF1_RF3_GDP],[RS30S_degraded,mRNA,RF1,Species('tRNA'+aa+xyz),RS50S,RF3,GDP], k_forward= rxn_k['re0000000851_k1']),
    # Reaction.from_massaction([termRS70SUAALpept_tRNAaaxyz_RF1_RF3_GDP],[RS50S_degraded,mRNA,RF1,Species('tRNA'+aa+xyz),RS30S,RF3,GDP], k_forward= rxn_k['re0000000852_k1']),
    # Reaction.from_massaction([termRS70SUAALpept_tRNAaaxyz_RF1_RF3_GDP],[RF1_degraded,RS50S,RS30S,Species('tRNA'+aa+xyz),mRNA,GDP,RF3], k_forward= rxn_k['re0000000853_k1']),
    # Reaction.from_massaction([termRS70SUAALpept_tRNAaaxyz_RF1_RF3_GDP],[RF3_degraded,mRNA,Species('tRNA'+aa+xyz),RF1,RS30S,RS50S,GDP], k_forward= rxn_k['re0000000854_k1']),

    # Reaction.from_massaction([termRS70SUAALpept_tRNAaaxyz_RF1_RF3],[RS30S_degraded,mRNA,RF1,Species('tRNA'+aa+xyz),RS50S,RF3], k_forward= rxn_k['re0000000855_k1']),
    # Reaction.from_massaction([termRS70SUAALpept_tRNAaaxyz_RF1_RF3],[RS50S_degraded,mRNA,RF1,Species('tRNA'+aa+xyz),RS30S,RF3], k_forward= rxn_k['re0000000856_k1']),
    # Reaction.from_massaction([termRS70SUAALpept_tRNAaaxyz_RF1_RF3],[RF1_degraded,RS50S,RS30S,Species('tRNA'+aa+xyz),mRNA,RF3], k_forward= rxn_k['re0000000857_k1']),
    # Reaction.from_massaction([termRS70SUAALpept_tRNAaaxyz_RF1_RF3],[RF3_degraded,mRNA,Species('tRNA'+aa+xyz),RF1,RS30S,RS50S], k_forward= rxn_k['re0000000858_k1']),
    # Reaction.from_massaction([termRS70SUAALpept_tRNAaaxyz_RF3_GTP],[RS30S_degraded,mRNA,Species('tRNA'+aa+xyz),RS50S,RF3,GTP], k_forward= rxn_k['re0000000859_k1']),
    # Reaction.from_massaction([termRS70SUAALpept_tRNAaaxyz_RF3_GTP],[RS50S_degraded,mRNA,Species('tRNA'+aa+xyz),RS30S,RF3,GTP], k_forward= rxn_k['re0000000860_k1']),
    # Reaction.from_massaction([termRS70SUAALpept_tRNAaaxyz_RF3_GTP],[RF3_degraded,RS50S,RS30S,Species('tRNA'+aa+xyz),mRNA,GTP], k_forward= rxn_k['re0000000861_k1']),
    # Reaction.from_massaction([termRS70SUAALpept_tRNAaaxyz_RF3_GDP_PO4],[RS30S_degraded,mRNA,Species('tRNA'+aa+xyz),RS50S,RF3,GDP,PO4], k_forward= rxn_k['re0000000862_k1']),
    # Reaction.from_massaction([termRS70SUAALpept_tRNAaaxyz_RF3_GDP_PO4],[RS50S_degraded,mRNA,Species('tRNA'+aa+xyz),RS30S,RF3,GDP,PO4], k_forward= rxn_k['re0000000863_k1']),
    # Reaction.from_massaction([termRS70SUAALpept_tRNAaaxyz_RF3_GDP_PO4],[RF3_degraded,RS50S,RS30S,Species('tRNA'+aa+xyz),mRNA,GDP,PO4], k_forward= rxn_k['re0000000864_k1']),

    # Reaction.from_massaction([termRS70SUAALpept_tRNAaaxyz_RF3_GDP],[RS30S_degraded,mRNA,Species('tRNA'+aa+xyz),RS50S,RF3,GDP], k_forward= rxn_k['re0000000865_k1']),
    # Reaction.from_massaction([termRS70SUAALpept_tRNAaaxyz_RF3_GDP],[RS50S_degraded,mRNA,Species('tRNA'+aa+xyz),RS30S,RF3,GDP], k_forward= rxn_k['re0000000866_k1']),
    # Reaction.from_massaction([termRS70SUAALpept_tRNAaaxyz_RF3_GDP],[RF3_degraded,RS50S,RS30S,Species('tRNA'+aa+xyz),mRNA,GDP], k_forward= rxn_k['re0000000867_k1']),
    # Reaction.from_massaction([termRS70SUAALpept_tRNAaaxyz_RF3_GDP,PO4],[termRS70SUAALpept_tRNAaaxyz_RF3_GDP_PO4], k_forward= rxn_k['re0000000868_k1']),
    # Reaction.from_massaction([RF3_GDP,termRS70SUAALpept_tRNAaaxyz],[termRS70SUAALpept_tRNAaaxyz_RF3_GDP], k_forward= rxn_k['re0000000869_k1']),
    Reaction.from_massaction([RF3_GDP,termRS70SUAALpept_tRNAaaxyz_RF2],[termRS70SUAALpept_tRNAaaxyz_RF2_RF3_GDP], k_forward= rxn_k['re0000000870_k1']),
    Reaction.from_massaction([termRS70SUAALpept_tRNAaaxyz_RF2_RF3_GDP],[termRS70SUAALpept_tRNAaaxyz_RF2,RF3_GDP], k_forward= rxn_k['re0000000871_k1']),
    Reaction.from_massaction([termRS70SUAALpept_tRNAaaxyz_RF2,RF3_GTP],[termRS70SUAALpept_tRNAaaxyz_RF2_RF3_GTP], k_forward= rxn_k['re0000000872_k1']),
    Reaction.from_massaction([termRS70SUAALpept_tRNAaaxyz_RF2_RF3_GTP],[termRS70SUAALpept_tRNAaaxyz_RF2,RF3_GTP], k_forward= rxn_k['re0000000873_k1']),
    Reaction.from_massaction([termRS70SUAALpept_tRNAaaxyz_RF2,RF3],[termRS70SUAALpept_tRNAaaxyz_RF2_RF3], k_forward= rxn_k['re0000000874_k1']),

    Reaction.from_massaction([termRS70SUAALpept_tRNAaaxyz_RF2_RF3],[termRS70SUAALpept_tRNAaaxyz_RF2,RF3], k_forward= rxn_k['re0000000875_k1']),
    # Reaction.from_massaction([termRS70SUAALpept_tRNAaaxyz_RF2_RF3_GTP],[RS30S_degraded,mRNA,RF2,Species('tRNA'+aa+xyz),RS50S,RF3,GTP], k_forward= rxn_k['re0000000876_k1']),
    # Reaction.from_massaction([termRS70SUAALpept_tRNAaaxyz_RF2_RF3_GTP],[RS50S_degraded,mRNA,RF2,Species('tRNA'+aa+xyz),RS30S,RF3,GTP], k_forward= rxn_k['re0000000877_k1']),
    # Reaction.from_massaction([termRS70SUAALpept_tRNAaaxyz_RF2_RF3_GTP],[RF2_degraded,RS50S,RS30S,Species('tRNA'+aa+xyz),mRNA,GTP,RF3], k_forward= rxn_k['re0000000878_k1']),
    Reaction.from_massaction([termRS70SUAALpept_tRNAaaxyz_RF2_RF3_GTP],[termRS70SUAALpept_tRNAaaxyz_RF3_GTP,RF2], k_forward= rxn_k['re0000000879_k1']),
    Reaction.from_massaction([termRS70SUAALpept_tRNAaaxyz_RF3_GTP,RF2],[termRS70SUAALpept_tRNAaaxyz_RF2_RF3_GTP], k_forward= rxn_k['re0000000880_k1']),
    Reaction.from_massaction([termRS70SUAALpept_tRNAaaxyz_RF2_RF3_GDP],[termRS70SUAALpept_tRNAaaxyz_RF2_RF3,GDP], k_forward= rxn_k['re0000000881_k1']),
    Reaction.from_massaction([termRS70SUAALpept_tRNAaaxyz_RF2_RF3,GDP],[termRS70SUAALpept_tRNAaaxyz_RF2_RF3_GDP], k_forward= rxn_k['re0000000882_k1']),
    Reaction.from_massaction([termRS70SUAALpept_tRNAaaxyz_RF2_RF3_GTP],[termRS70SUAALpept_tRNAaaxyz_RF2_RF3,GTP], k_forward= rxn_k['re0000000883_k1']),
    Reaction.from_massaction([termRS70SUAALpept_tRNAaaxyz_RF2_RF3,GTP],[termRS70SUAALpept_tRNAaaxyz_RF2_RF3_GTP], k_forward= rxn_k['re0000000884_k1']),

    # Reaction.from_massaction([termRS70SUAALpept_tRNAaaxyz_RF2_RF3_GTP],[RF3_degraded,mRNA,Species('tRNA'+aa+xyz),RF2,RS30S,RS50S,GTP], k_forward= rxn_k['re0000000885_k1']),
    # Reaction.from_massaction([termRS70SUAALpept_tRNAaaxyz_RF2_RF3_GDP],[RS30S_degraded,mRNA,RF2,Species('tRNA'+aa+xyz),RS50S,RF3,GDP], k_forward= rxn_k['re0000000886_k1']),
    # Reaction.from_massaction([termRS70SUAALpept_tRNAaaxyz_RF2_RF3_GDP],[RS50S_degraded,mRNA,RF2,Species('tRNA'+aa+xyz),RS30S,RF3,GDP], k_forward= rxn_k['re0000000887_k1']),
    # Reaction.from_massaction([termRS70SUAALpept_tRNAaaxyz_RF2_RF3_GDP],[RF2_degraded,RS50S,RS30S,Species('tRNA'+aa+xyz),mRNA,GDP,RF3], k_forward= rxn_k['re0000000888_k1']),
    # Reaction.from_massaction([termRS70SUAALpept_tRNAaaxyz_RF2_RF3_GDP],[RF3_degraded,mRNA,Species('tRNA'+aa+xyz),RF2,RS30S,RS50S,GDP], k_forward= rxn_k['re0000000889_k1']),
    # Reaction.from_massaction([termRS70SUAALpept_tRNAaaxyz_RF2_RF3],[RS30S_degraded,mRNA,RF2,Species('tRNA'+aa+xyz),RS50S,RF3], k_forward= rxn_k['re0000000890_k1']),
    # Reaction.from_massaction([termRS70SUAALpept_tRNAaaxyz_RF2_RF3],[RS50S_degraded,mRNA,RF2,Species('tRNA'+aa+xyz),RS30S,RF3], k_forward= rxn_k['re0000000891_k1']),
    # Reaction.from_massaction([termRS70SUAALpept_tRNAaaxyz_RF2_RF3],[RF2_degraded,RS50S,RS30S,Species('tRNA'+aa+xyz),mRNA,RF3], k_forward= rxn_k['re0000000892_k1']),
    # Reaction.from_massaction([termRS70SUAALpept_tRNAaaxyz_RF2_RF3],[RF3_degraded,mRNA,Species('tRNA'+aa+xyz),RF2,RS30S,RS50S], k_forward= rxn_k['re0000000893_k1']),
    Reaction.from_massaction([termRS70SUAALpept_tRNAaaxyz_RRF,EFG_GTP],[termRS70SUAALpept_tRNAaaxyz_RRF_EFG_GTP], k_forward= rxn_k['re0000000895_k1']),

    Reaction.from_massaction([termRS70SUAALpept_tRNAaaxyz_RRF_EFG_GTP],[termRS70SUAALpept_tRNAaaxyz_RRF,EFG_GTP], k_forward= rxn_k['re0000000896_k1']),
    # Reaction.from_massaction([termRS70SUAALpept_tRNAaaxyz_RRF_EFG_GTP],[RS30S_degraded,mRNA,RRF,Species('tRNA'+aa+xyz),RS50S,EFG,GTP], k_forward= rxn_k['re0000000897_k1']),
    # Reaction.from_massaction([termRS70SUAALpept_tRNAaaxyz_RRF_EFG_GTP],[RS50S_degraded,mRNA,RRF,Species('tRNA'+aa+xyz),RS30S,EFG,GTP], k_forward= rxn_k['re0000000898_k1']),
    # Reaction.from_massaction([termRS70SUAALpept_tRNAaaxyz_RRF_EFG_GTP],[RRF_degraded,RS50S,RS30S,Species('tRNA'+aa+xyz),mRNA,GTP,EFG], k_forward= rxn_k['re0000000899_k1']),
    Reaction.from_massaction([termRS70SUAALpept_tRNAaaxyz_RRF_EFG_GTP],[termRS70SUAALpept_tRNAaaxyz_EFG_GTP,RRF], k_forward= rxn_k['re0000000900_k1']),
    Reaction.from_massaction([termRS70SUAALpept_tRNAaaxyz_EFG_GTP,RRF],[termRS70SUAALpept_tRNAaaxyz_RRF_EFG_GTP], k_forward= rxn_k['re0000000901_k1']),
    Reaction.from_massaction([termRS70SUAALpept_tRNAaaxyz_RRF_EFG_GDP_PO4],[termRS70SUAALpept_tRNAaaxyz_RRF_EFG_GDP,PO4], k_forward= rxn_k['re0000000902_k1']),
    # Reaction.from_massaction([termRS70SUAALpept_tRNAaaxyz_RRF_EFG_GTP],[EFG_degraded,mRNA,Species('tRNA'+aa+xyz),RRF,RS30S,RS50S,GTP], k_forward= rxn_k['re0000000903_k1']),
    Reaction.from_massaction([termRS70SUAALpept_tRNAaaxyz,RRF],[termRS70SUAALpept_tRNAaaxyz_RRF], k_forward= rxn_k['re0000000904_k1']),
    Reaction.from_massaction([termRS70SUAALpept_tRNAaaxyz_RRF],[termRS70SUAALpept_tRNAaaxyz,RRF], k_forward= rxn_k['re0000000905_k1']),

    Reaction.from_massaction([termRS70SUAALpept_tRNAaaxyz,EFG_GTP],[termRS70SUAALpept_tRNAaaxyz_EFG_GTP], k_forward= rxn_k['re0000000906_k1']),
    Reaction.from_massaction([termRS70SUAALpept_tRNAaaxyz_EFG_GTP],[termRS70SUAALpept_tRNAaaxyz,EFG_GTP], k_forward= rxn_k['re0000000907_k1']),
    Reaction.from_massaction([termRS70SUAALpept_tRNAaaxyz_RRF_EFG_GTP],[termRS70SUAALpept_tRNAaaxyz_RRF_EFG_GDP_PO4], k_forward= rxn_k['re0000000908_k1']),
    Reaction.from_massaction([termRS70SUAALpept_tRNAaaxyz_RRF_EFG_GDP_PO4],[termRS70SUAALpept_tRNAaaxyz_RRF_EFG_GTP], k_forward= rxn_k['re0000000909_k1']),
    Reaction.from_massaction([termRS70SUAALpept_tRNAaaxyz_RRF_EFG_GDP],[Species('RS50S_tRNA'+aa+xyz+'_RRF_EFG_GDP'),termRS30S_mRNA], k_forward= rxn_k['re0000000910_k1']),
    Reaction.from_massaction([termRS30S_mRNA],[RS30S,mRNA], k_forward= rxn_k['re0000000911_k1']),
    # Reaction.from_massaction([RS30S,mRNA],[termRS30S_mRNA], k_forward= rxn_k['re0000000912_k1']),
    # Reaction.from_massaction([termRS30S_mRNA],[RS30S_degraded,mRNA], k_forward= rxn_k['re0000000926_k1']),
    # Reaction.from_massaction([termRS70SUAALpept_tRNAaaxyz_RRF],[RS30S_degraded,mRNA,RRF,Species('tRNA'+aa+xyz),RS50S], k_forward= rxn_k['re0000000938_k1']),
    # Reaction.from_massaction([termRS70SUAALpept_tRNAaaxyz_RRF],[RS50S_degraded,mRNA,RRF,Species('tRNA'+aa+xyz),RS30S], k_forward= rxn_k['re0000000939_k1']),

    # Reaction.from_massaction([termRS70SUAALpept_tRNAaaxyz_RRF],[RRF_degraded,RS50S,RS30S,Species('tRNA'+aa+xyz),mRNA], k_forward= rxn_k['re0000000940_k1']),
    # Reaction.from_massaction([termRS70SUAALpept_tRNAaaxyz_EFG_GTP],[RS30S_degraded,mRNA,Species('tRNA'+aa+xyz),RS50S,EFG,GTP], k_forward= rxn_k['re0000000941_k1']),
    # Reaction.from_massaction([termRS70SUAALpept_tRNAaaxyz_EFG_GTP],[RS50S_degraded,mRNA,Species('tRNA'+aa+xyz),RS30S,EFG,GTP], k_forward= rxn_k['re0000000942_k1']),
    # Reaction.from_massaction([termRS70SUAALpept_tRNAaaxyz_EFG_GTP],[EFG_degraded,RS50S,RS30S,Species('tRNA'+aa+xyz),mRNA,GTP], k_forward= rxn_k['re0000000943_k1']),
    # Reaction.from_massaction([termRS70SUAALpept_tRNAaaxyz_RRF_EFG_GDP],[RS30S_degraded,mRNA,RRF,Species('tRNA'+aa+xyz),RS50S,EFG,GDP], k_forward= rxn_k['re0000000944_k1']),
    #Reaction.from_massaction([termRS70SUAALpept_tRNAaaxyz_RRF_EFG_GDP],[RS50S_degraded,mRNA,RRF,Species('tRNA'+aa+xyz),RS30S,EFG,GDP], k_forward= rxn_k['re0000000945_k1']),
    # Reaction.from_massaction([termRS70SUAALpept_tRNAaaxyz_RRF_EFG_GDP],[RRF_degraded,RS50S,RS30S,Species('tRNA'+aa+xyz),mRNA,GDP,EFG], k_forward= rxn_k['re0000000946_k1']),
    # Reaction.from_massaction([termRS70SUAALpept_tRNAaaxyz_RRF_EFG_GDP],[EFG_degraded,mRNA,Species('tRNA'+aa+xyz),RRF,RS30S,RS50S,GDP], k_forward= rxn_k['re0000000947_k1']),
    # Reaction.from_massaction([termRS70SUAALpept_tRNAaaxyz_RRF_EFG_GDP_PO4],[RS30S_degraded,mRNA,RRF,Species('tRNA'+aa+xyz),RS50S,EFG,GDP,PO4], k_forward= rxn_k['re0000000948_k1']),
    # Reaction.from_massaction([termRS70SUAALpept_tRNAaaxyz_RRF_EFG_GDP_PO4],[RS50S_degraded,mRNA,RRF,Species('tRNA'+aa+xyz),RS30S,EFG,GDP,PO4], k_forward= rxn_k['re0000000949_k1']),

    # Reaction.from_massaction([termRS70SUAALpept_tRNAaaxyz_RRF_EFG_GDP_PO4],[RRF_degraded,RS50S,RS30S,Species('tRNA'+aa+xyz),mRNA,GDP,EFG,PO4], k_forward= rxn_k['re0000000950_k1']),
    # Reaction.from_massaction([termRS70SUAALpept_tRNAaaxyz_RRF_EFG_GDP_PO4],[EFG_degraded,mRNA,Species('tRNA'+aa+xyz),RRF,RS30S,RS50S,GDP,PO4], k_forward= rxn_k['re0000000951_k1']),
    # Reaction.from_massaction([PO4,termRS70SUAALpept_tRNAaaxyz_RRF_EFG_GDP],[termRS70SUAALpept_tRNAaaxyz_RRF_EFG_GDP_PO4], k_forward= rxn_k['re0000000956_k1']),
    # Reaction.from_massaction([Species('RS50S_tRNA'+aa+xyz+'_RRF_EFG_GDP'),termRS30S_mRNA],[termRS70SUAALpept_tRNAaaxyz_RRF_EFG_GDP], k_forward= rxn_k['re0000000957_k1']),
    ]
```

### Model¶

In [14]:

```
#Compile all reactions and species
All_species = flatten([list_species_gen, list_species_elfmet, list_species_fmet,
                       flatten(list_species_aa), list_species_el2p2, list_species_el3p3,
                       list_species_el4p3, list_species_elterm])

All_rxn = flatten([list_of_reaction_gen, list_reaction_fmet, 
                   list_of_reactions, list_reactions_elfmet,
                   list_of_reaction_pept2, list_of_reaction_pept3,
                   list_of_reactions_end, list_of_reaction_term,])

CRN_TL = ChemicalReactionNetwork(species = All_species, reactions = All_rxn)
# CRN_TL.write_sbml_file(directory+"/CRN_PURE_TLonly/CRN_PURE_TLonlywZeros_deGFP.xml")
```

In [15]:

```
#Get intial conditons and run simulation
folder= '/CRN_PURE_TLonly/'
filename_ic = 'fMGG_synthesis_initial_values_CRNv2.csv'
with open(directory + folder+ filename_ic, mode='r') as infile:
    reader = csv.reader(infile)
    initial_con= {rows[0]:float(rows[1]) for rows in reader}

t0=time.time()
timepoints0 = np.linspace(10**-5, 10**4, 500)

PURE_TLonly = CRN_TL.simulate_with_bioscrape_via_sbml(timepoints = timepoints0, initial_condition_dict = initial_con)
# PURE_TLonly.to_csv(directory +'/CRN_PURE_TLonly_MGaptdeGFP_results.csv')

t1=time.time()

ct = datetime.datetime.now()
print((t1-t0)/60, "current time:-", ct)
```

```
C:\Users\zoila\anaconda3\envs\python38\lib\site-packages\scipy\integrate\_odepack_py.py:248: ODEintWarning: Excess work done on this call (perhaps wrong Dfun type). Run with full_output = 1 to get quantitative information.
  warnings.warn(warning_msg, ODEintWarning)
```

```
13.508141819636027 current time:- 2023-08-14 13:32:37.686598
```

### Plot results along with Experimental data¶

In [16]:

```
p1 = bokeh.plotting.figure(toolbar_location='right',
    outline_line_color= None,
    min_border_right=10,
    height=500,
    width=500,)

p1.title.text = "B) TX only simulation results of deGFP"
p1.xaxis.axis_label = 'Time (hours)'
p1.yaxis.axis_label = 'deGFP (μM)'
p1.y_range=Range1d(0, 8)
p1.x_range=Range1d(0, 4)
p1.outline_line_color=None

# p1.yaxis
p1.ygrid.visible = False
p1.yaxis.axis_label_text_font_size='15pt' 
p1.yaxis.major_label_text_font_size = '15pt'
p1.yaxis.major_label_text_font='Work Sans'
p1.yaxis.axis_label_standoff=15
p1.yaxis.axis_label_text_font_style='normal' 

# p1.xaxis
p1.xgrid.visible = False
p1.xaxis.axis_label_text_font_size='15pt' 
p1.xaxis.major_label_text_font_size = '15pt'
p1.xaxis.major_label_text_font='Work Sans'
p1.xaxis.axis_label_standoff=15
p1.xaxis.axis_label_text_font_style='normal' 

# p1.title
p1.title.text_font_size= '18pt' 
p1.title.align= 'left'

p1.title.offset=-65.0
```

In [17]:

```
#Experimental data set of one repeat
folder_exp='/MGapt_deGFP_Exp_PURE_Data'
filename='/2022.08.18_PURE_deGFP.csv'
deGFP_Data = pd.read_csv(directory + folder_exp+filename)
deGFP_Data['deGFP_uM']=(deGFP_Data['deGFP']+391.44)/1624.27-0.24

#Experimental data set 1
p1.scatter(deGFP_Data['timepoints']/3600, deGFP_Data['deGFP_uM'], legend_label = "T7-deGFP.repeat1", radius=0.01, fill_alpha=0.2, color="navy") 

#Simulation results
p1.line(PURE_TLonly['time']/3600, PURE_TLonly[str(PeptGpept)],legend_label = "k1*Model", line_color="magenta", line_width=5) 

p1.legend.location='bottom_right'
p1.x_range=Range1d(0, 2.5)
bokeh.io.show(p1)
```

### Computing environment¶

In [18]:

```
%load_ext watermark
%watermark -v -p bioscrape,bokeh,panel,jupyterlab,biocrnpyler
```

```
Python implementation: CPython
Python version       : 3.8.17
IPython version      : 8.12.2

bioscrape  : 1.2.1
bokeh      : 2.4.0
panel      : 0.13.1
jupyterlab : 3.6.5
biocrnpyler: 1.1.1
```
