## Supporting scripts, data, and simulation results. for "A chemical reaction network model of PURE": CRN_PURE_TXonly_Final.html


### PURE TX-only Model in BioCRNpyler¶

August 8, 2023

[1] Division of Engineering and Applied Science, California Institute of Technology, Pasadena, CA \
[2] Division of Biology and Biological Engineering, California Institute of Technology, Pasadena, CA \

In this notebook, we built a chemical reaction network transcription model for PURE protein synthesis. This model is based on the TX-TL from Tuza *et al.* [1] originally built for cell lysate protein expression. This section of code requires a DNA sequence input from the user. This TX-only model was used in the parameter inferencing and sensitivity analysis described in the BioRxiv paper. The steps of the TX are split into 3 subsections: species and reactions based on nucleotide needed (1), generates complement sequence (2), and species and reactions of growing mRNA strand (3).

1. Non-nucleic acid Species and Reactions
2. RNA sequence
3. Addition of nucleotide and individual reactions for the desired DNA

##### Citation:¶

[1] Zoltán A. Tuza, Dan Siegal-Gaskins, Jongmin Kim, and Gábor Szederkényi. Analysis-based parameter estimation of an *in vitro* transcription-translation system. In Eur. Control Conf. 2015, page 1554–1560, 2015. (PNAS, 2015)

#### Importing required packages and definitions¶

In [1]:

```
import biocrnpyler
from biocrnpyler import *
from biocrnpyler.component import Component
from biocrnpyler.chemical_reaction_network import Species, Reaction, ChemicalReactionNetwork
from biocrnpyler.mechanism import Mechanism
from biocrnpyler.reaction import Reaction
from biocrnpyler.species import Complex, Species, WeightedSpecies

import bioscrape

from typing import List, Union
import numpy as np
import pandas as pd
import pylab as plt
import csv
import numpy as np
import pandas as pd
import math
import seaborn as sns; sns.set()
import matplotlib.pyplot as plt
import random 
import time
import datetime
from itertools import groupby
import libsbml
import itertools as it
import itertools as it

#Get directory
import os
directory = os.getcwd()

def flatten(t):
    return [item for sublist in t for item in sublist]

t0=time.time()
```

##### Bokeh¶

In [2]:

```
%matplotlib inline
import bokeh.io
import bokeh.plotting
bokeh.io.output_notebook()
from bokeh.themes import Theme

# Modules needed from Bokeh.
from bokeh.io import output_file, show
from bokeh.plotting import figure
from bokeh.models import LinearAxis, Range1d

colors = bokeh.palettes.Colorblind[8]

try:
    import dnaplotlib as dpl
    dpl_enabled = True
except (ModuleNotFoundError,ImportError) as e:
    dpl_enabled = False
    
theme = Theme(json={'attrs': {
# apply defaults to Figure properties
'Figure': {
    'toolbar_location': 'right',
    'outline_line_color': None,
    'min_border_right': 10,
#     'sizing_mode': 'stretch_width',
    'height':600,
    'width':800,
},'Grid': {
    'grid_line_color': None,
},
'Title': {
    'text_font_size': '20pt',
    'align': 'center'
},
    
    
# apply defaults to Axis properties
'Axis': {
#     'minor_tick_out': None,
#     'minor_tick_in': None,
    'major_label_text_font_size': '15pt',
    'axis_label_text_font_size': '15pt',
#     'axis_label_text_font': 'Work Sans',
    'axis_label_text_font_style': 'normal',
    'axis_label_standoff':15
},


# apply defaults to Legend properties
'Legend': {
    'background_fill_alpha': 0.8,
    'location': 'top_right',
    "label_text_font_size": '15pt'
}}})

bokeh.io.curdoc().theme = theme
from bokeh.io import export_png
```

Loading BokehJS ...

#### Getting experimental data¶

In [3]:

```
# DNA= MGapt_Data (no RBS)
folder_data='/MGapt_deGFP_Exp_PURE_Data/'
filename = '2022.08.19_OnePot_PURE_mGapt_uM.csv'
mGapt_Data =  pd.read_csv(directory+folder_data+filename, delimiter = '\,', names = ['time','mRNA0', 'mRNA1','mRNA2', 'mRNA3'], skiprows = 1)
```

```
C:\Users\zoila\AppData\Local\Temp\ipykernel_8336\210321641.py:4: ParserWarning: Falling back to the 'python' engine because the 'c' engine does not support regex separators (separators > 1 char and different from '\s+' are interpreted as regex); you can avoid this warning by specifying engine='python'.
  mGapt_Data =  pd.read_csv(directory+folder_data+filename, delimiter = '\,', names = ['time','mRNA0', 'mRNA1','mRNA2', 'mRNA3'], skiprows = 1)
```

In [4]:

```
# DNA= MGapt-UTR1-deGFP_Data
filename1 = '2022.11.14_OnePotPURE_deGFPuM.csv'
MGaptdeGFP_gfp =  pd.read_csv(directory+folder_data+filename1, delimiter = '\,', names = ['timepoints','deGFP_1','deGFP_2','deGFP_3'], skiprows = 1)

filename2 = '2022.11.14_OnePotPURE_MGaptuM.csv'
MGaptdeGFP_rna =  pd.read_csv(directory+folder_data+filename2, delimiter = '\,', names = ['timepoints','MGapt_1','MGapt_2','MGapt_3'], skiprows = 1)
```

```
C:\Users\zoila\AppData\Local\Temp\ipykernel_8336\1803188381.py:3: ParserWarning: Falling back to the 'python' engine because the 'c' engine does not support regex separators (separators > 1 char and different from '\s+' are interpreted as regex); you can avoid this warning by specifying engine='python'.
  MGaptdeGFP_gfp =  pd.read_csv(directory+folder_data+filename1, delimiter = '\,', names = ['timepoints','deGFP_1','deGFP_2','deGFP_3'], skiprows = 1)
C:\Users\zoila\AppData\Local\Temp\ipykernel_8336\1803188381.py:6: ParserWarning: Falling back to the 'python' engine because the 'c' engine does not support regex separators (separators > 1 char and different from '\s+' are interpreted as regex); you can avoid this warning by specifying engine='python'.
  MGaptdeGFP_rna =  pd.read_csv(directory+folder_data+filename2, delimiter = '\,', names = ['timepoints','MGapt_1','MGapt_2','MGapt_3'], skiprows = 1)
```

### User defined transcribed unit¶

In [5]:

```
#User will copy the sequence bwteen end of promoter and start of terminator of DNA strain
#mGapt sequence
dna_seq='GGATCCCGACTGGCGAGAGCCAGGTAACGAATGGATCTCGAGCCTTAGGAGATCCGGCTGCTAACAAAGCCCGAAAGGAAGCTGAGTTG'

#mGapt_UTR1_deGFP (uncomment below if modeling indicated plasmid)
# dna_seq= 'GGGATCCCGACTGGCGAGAGCCAGGTAACGAATGGATCCAATAATTTTGTTTAACTTTAAGAAGGAGATATACCATGGAGCTTTTCACTGGCGTTGTTCCCATCCTGGTCGAGCTGGACGGCGACGTAAACGGCCACAAGTTCAGCGTGTCCGGCGAGGGCGAGGGCGATGCCACCTACGGCAAGCTGACCCTGAAGTTCATCTGCACCACCGGCAAGCTGCCCGTGCCCTGGCCCACCCTCGTGACCACCCTGACCTACGGCGTGCAGTGCTTCAGCCGCTACCCCGACCACATGAAGCAGCACGACTTCTTCAAGTCCGCCATGCCCGAAGGCTACGTCCAGGAGCGCACCATCTTCTTCAAGGACGACGGCAACTACAAGACCCGCGCCGAGGTGAAGTTCGAGGGCGACACCCTGGTGAACCGCATCGAGCTGAAGGGCATCGACTTCAAGGAGGACGGCAACATCCTGGGGCACAAGCTGGAGTACAACTACAACAGCCACAACGTCTATATCATGGCCGACAAGCAGAAGAACGGCATCAAGGTGAACTTCAAGATCCGCCACAACATCGAGGACGGCAGCGTGCAGCTCGCCGACCACTACCAGCAGAACACCCCCATCGGCGACGGCCCCGTGCTGCTGCCCGACAACCACTACCTGAGCACCCAGTCCGCCCTGAGCAAAGACCCCAACGAGAAGCGCGATCACATGGTCCTGCTGGAGTTCGTGACCGCCGCCGGGATCTAACTCGAGCCTTAGGAGATCCGGCTGCTAACAAAGCCCGAAAGGAAGCTGAGTTG'
```

### Transcription (TX)¶

#### Non-nucleic acid Species and Reactions¶

Creates all the species and reaction around the nucleotide sequences in the given transcribed region, common protein and small molecules.

In [6]:

```
## Species
#Define species needed for TX 
RNAPa = Species('RNAPa') #ActivePolymerase
DNA = Species("DNA") 
mRNA=Species('mRNA')
RNAPa_bound = Species("RNAPa_bound") 
RNAPa_bound_GTP=Species('RNAPa_bound_GTP') #Attached RNAP to DNA
RNAPa_bound_GDP_PO4= Species('RNAPa_bound_GDP_PO4')

#Nucleotide bases that are "active"
ATP=Species('ATP')
GTP=Species('GTP')
CTP=Species('CTP')
UTP=Species('UTP')

#Other small molecules
GDP=Species('GDP')
PPi=Species('PPi')
PO4=Species('PO4')

##Rates
#Degradation rate
NTP_deg= 1.75*10**(-4) 
k_rnapbF = 2.9*10**(-1)
k_rnapbF2 = 6.3*10**(-2)
k_rnapbF3 = 4.2*10**(-1)


k_trap= NTP_deg 
NTP_charge= 223.6 


#Define placing all species in one array
Species_general= [RNAPa, DNA, mRNA, RNAPa_bound_GTP, RNAPa_bound, 
                  ATP, GTP, CTP, UTP,
                  GDP, PPi, PO4,
                  RNAPa_bound_GDP_PO4, RNAPa_bound_GTP,]

#Define Reactions needed for TX regardless of the sequence defined
RXN_general= [
    #NTP degradation
    Reaction.from_massaction([ATP], [], k_forward=NTP_deg), 
    Reaction.from_massaction([GTP], [], k_forward=NTP_deg), 
    Reaction.from_massaction([CTP], [], k_forward=NTP_deg), 
    Reaction.from_massaction([UTP], [], k_forward=NTP_deg), 
    #Binding of RNAP and beginning of transcription
    Reaction.from_massaction([RNAPa, DNA, GTP], [RNAPa_bound_GTP], k_forward=k_rnapbF),
    Reaction.from_massaction([RNAPa_bound_GTP], [RNAPa_bound_GDP_PO4], k_forward=k_rnapbF2),
    Reaction.from_massaction([RNAPa_bound_GDP_PO4], [RNAPa_bound, GDP, PO4,], k_forward=k_rnapbF3)]
```

#### Transcribing DNA to mRNA nucleotides¶

In [7]:

```
# Transcribed given DNA sequence into mRNA sequence
rna_seq = []
for bp in dna_seq:
    if bp == 'A':
        tx = UTP
    elif bp == 'T':
        tx =ATP
    elif bp == 'G':
        tx =CTP
    elif bp == 'C':
        tx =GTP
    else:
        print('Non-standard nucleotides')
    rna_seq.append(tx)
```

#### Addition of each nucleotide and individual reactions needed¶

In [8]:

```
nt_len = len(rna_seq)

k_mrna0=8.2*10**(-2)
k_ntpf = 6.2*10**(-1)
k_addnt = (8.9)*10**(-0) 
k_ppi = 1000 
k_term= 1.8 

#Start iternations
mrna_length='0000'
rxn_list=[]
species_list=[]

for L in [l for l in range(len(rna_seq))]:
    ntp = rna_seq[L]
    mrna0 = mrna_length[:-len(str(L))]+str(L) #starting mRNA
    mrnaG = mrna_length[:-len(str(L+1))]+str(L+1) #Growing mRNA

    if L == 0:        
        RNAPa_bound_mRNAmrna0 = Species('RNAPa_bound_mRNA'+mrna0)
        RNAPa_bound_mRNAmrna0_ntp = Species('RNAPa_bound_mRNA'+mrna0+'_'+ str(ntp))
        RNAPa_bound_mRNAmrnaG_nmp_PPi = Species('RNAPa_bound_mRNA'+mrnaG+'_PPi')
        RNAPa_bound_mRNAmrnaG = Species('RNAPa_bound_mRNA'+mrnaG)
        
        species= [RNAPa_bound_mRNAmrna0, RNAPa_bound_mRNAmrna0_ntp,
                  RNAPa_bound_mRNAmrnaG_nmp_PPi, RNAPa_bound_mRNAmrnaG]
        
        rxns=[Reaction.from_massaction([RNAPa_bound],[Species('RNAPa_bound_mRNA'+mrna0)],k_forward=k_mrna0), 
              Reaction.from_massaction([Species('RNAPa_bound_mRNA'+mrna0), ntp], [Species('RNAPa_bound_mRNA'+mrna0+'_'+ str(ntp))], k_forward=k_ntpf),
              Reaction.from_massaction([Species('RNAPa_bound_mRNA'+mrna0+'_'+ str(ntp))],[Species('RNAPa_bound_mRNA'+mrnaG+'_PPi')],k_forward=k_addnt),
            Reaction.from_massaction([Species('RNAPa_bound_mRNA'+mrnaG+'_PPi')],[Species('RNAPa_bound_mRNA'+mrnaG), PPi,],k_forward=k_ppi),] 
    elif L<nt_len-1:
        
        RNAPa_bound_mRNAmrna0_ntp = Species('RNAPa_bound_mRNA'+mrna0+'_'+ str(ntp))
        RNAPa_bound_mRNAmrnaG_nmp_PPi = Species('RNAPa_bound_mRNA'+mrnaG+'_PPi')
        RNAPa_bound_mRNAmrnaG = Species('RNAPa_bound_mRNA'+mrnaG)
        
        species= [RNAPa_bound_mRNAmrna0_ntp,
                  RNAPa_bound_mRNAmrnaG_nmp_PPi, RNAPa_bound_mRNAmrnaG]
        
        rxns=[Reaction.from_massaction([Species('RNAPa_bound_mRNA'+mrna0), ntp], [Species('RNAPa_bound_mRNA'+mrna0+'_'+ str(ntp))], k_forward=k_ntpf),

              Reaction.from_massaction([Species('RNAPa_bound_mRNA'+mrna0+'_'+ str(ntp))],[Species('RNAPa_bound_mRNA'+mrnaG+'_PPi')],k_forward=k_addnt),
              Reaction.from_massaction([Species('RNAPa_bound_mRNA'+mrnaG+'_PPi')],[Species('RNAPa_bound_mRNA'+mrnaG), PPi,],k_forward=k_ppi),]  
#         
    else:
        RNAPa_bound_mRNAmrna0_ntp = Species('RNAPa_bound_mRNA'+mrna0+'_'+ str(ntp))
        RNAPa_bound_mRNAmrnaG_nmp_PPi = Species('RNAPa_bound_mRNA'+mrnaG+'_PPi')
        RNAPa_bound_mRNAmrnaG = Species('RNAPa_bound_mRNA'+mrnaG)
        
        species= [RNAPa_bound_mRNAmrna0_ntp,
                  RNAPa_bound_mRNAmrnaG_nmp_PPi, RNAPa_bound_mRNAmrnaG,]
                   
        rxns=[Reaction.from_massaction([Species('RNAPa_bound_mRNA'+mrna0), ntp], [Species('RNAPa_bound_mRNA'+mrna0+'_'+ str(ntp))], k_forward=k_ntpf),
              Reaction.from_massaction([Species('RNAPa_bound_mRNA'+mrna0+'_'+ str(ntp))],[Species('RNAPa_bound_mRNA'+mrnaG+'_PPi')],k_forward=k_addnt),  
              Reaction.from_massaction([Species('RNAPa_bound_mRNA'+mrnaG+'_PPi')],[Species('RNAPa_bound_mRNA'+mrnaG), PPi],k_forward=k_ppi),
              Reaction.from_massaction([Species('RNAPa_bound_mRNA'+mrnaG),],[RNAPa, DNA, mRNA],k_forward=k_term,)]  
        
    rxn_list.append(rxns)
    species_list.append(species)

species_list = flatten(species_list)
rxn_list= flatten(rxn_list)
```

#### Simulate mRNA¶

In [9]:

```
All_species = flatten([Species_general, species_list])
All_rxn = flatten([RXN_general, rxn_list])

CRN_TX = ChemicalReactionNetwork(species = All_species, reactions = All_rxn)

#IC for MGapt
initial_con={'RNAPa':(0.1), 'DNA':(.005), 'ATP':(3000), 'GTP':(3000), 'CTP':(1000), 'UTP':(1000),} #in uM

# #IC for MGapt-deGFP
# initial_con={'RNAPa':(0.1), 'DNA':(.001), 'ATP':(3000), 'GTP':(3000), 'CTP':(1000), 'UTP':(1000),} #in uM


timepoints = np.linspace(10**-5, 6*10**4, 500)
CRN_TX_only = CRN_TX.simulate_with_bioscrape_via_sbml(timepoints = timepoints, initial_condition_dict = initial_con )

t1=time.time()
total = t1-t0
print('total time (s)= ')
print(total)
```

```
total time (s)= 
1.93406343460083
```

#### Plotting for MGapt\_mRNA only¶

In [10]:

```
p1 = bokeh.plotting.figure()
p1.title.text = "mRNA Concentration Over Time"
p1.xaxis.axis_label = 'Time (hours)'
p1.yaxis.axis_label = 'mRNA (uM)'

p1.line(timepoints/3600, CRN_TX_only[str(mRNA)], legend_label = "Model_TXonly", line_color="magenta", line_width=5) #mRNA

#Experimental data set 1
p1.scatter(mGapt_Data['time']/3600, mGapt_Data['mRNA1'], legend_label = "T7-mGapt.repeat1", radius=0.05, fill_alpha=0.5, color="navy") #mRNA
p1.scatter(mGapt_Data['time']/3600, mGapt_Data['mRNA2'], legend_label = "T7-mGapt.repeat2", radius=0.05, fill_alpha=0.5, color="blue")#mRNA
p1.scatter(mGapt_Data['time']/3600, mGapt_Data['mRNA3'], legend_label = "T7-mGapt.repeat3", radius=0.05, fill_alpha=0.5, color= "steelblue") #mRNA
#Experimental data set 2 lightsteelblue, powderblue, lightblue, skyblue, lightskyblue, deepskyblue, dodgerblue, cornflowerblue, steelblue, royalblue, blue, mediumblue, darkblue, navy, midnightblue
p1.scatter(mGapt_Data['time']/3600, mGapt_Data['mRNA0'], legend_label = "T7-mGapt.repeat4", radius=0.05, fill_alpha=0.5, color= "royalblue") #mRNAp1.legend.location='bottom_right'

p1.legend.location='bottom_right'
p1.x_range=Range1d(0, 8)
bokeh.io.show(p1)
```

#### Plotting for MGapt-UTR1-deGFP\_mRNA only¶

In [11]:

```
# p1 = bokeh.plotting.figure()
# p1.title.text = "mRNA Concentration Over Time"
# p1.xaxis.axis_label = 'Time (hours)'
# p1.yaxis.axis_label = 'mRNA (uM)'

# p1.line(timepoints/3600, CRN_TX_only[str(mRNA)], legend_label = "Model_TXonly", line_color="magenta", line_width=5) #mRNA

# #Experimental data set 1
# p1.scatter(MGaptdeGFP_rna['timepoints']/60, MGaptdeGFP_rna['MGapt_1'], legend_label = "T7-mGapt_deGFP.R1", radius=0.01, fill_alpha=0.5, color="navy") #mRNA
# p1.scatter(MGaptdeGFP_rna['timepoints']/60, MGaptdeGFP_rna['MGapt_2'], legend_label = "T7-mGapt_deGFP.R2", radius=0.01, fill_alpha=0.5, color="navy") #mRNA
# p1.scatter(MGaptdeGFP_rna['timepoints']/60, MGaptdeGFP_rna['MGapt_3'], legend_label = "T7-mGapt_deGFP.R3", radius=0.01, fill_alpha=0.5, color="navy") #mRNA
# p1.legend.location='top_left'
# p1.x_range=Range1d(0, 6)
# bokeh.io.show(p1)
```

### Computing environment¶

In [12]:

```
%load_ext watermark
%watermark -v -p bioscrape,bokeh,panel,jupyterlab,biocrnpyler
```

```
Python implementation: CPython
Python version       : 3.8.17
IPython version      : 8.12.2

bioscrape  : 1.2.1
bokeh      : 2.4.0
panel      : 0.13.1
jupyterlab : 3.6.5
biocrnpyler: 1.1.1
```
