## Supporting scripts, data, and simulation results. for "A chemical reaction network model of PURE": CRN_PURE_TXTL_MGaptdeGFP_FINAL.html

August 8, 2023

[1] Division of Engineering and Applied Science, California Institute of Technology, Pasadena, CA \
[2] Division of Biology and Biological Engineering, California Institute of Technology, Pasadena, CA \

In this notebook, we built a chemical reaction network transcription model for PURE protein synthesis. The transcription and translation models are separately described and generated. All species and reactions in the transcription and translation model are then combined together with additional reactions and species. The combined PURE model can be used to model the expression of a protein in PURE. We used this notebook to model the transcription and translation of a plasmid containing the construct: pT7-MGapt-UTR1-deGFP-tT7.

A full description of the model can be found in the BioRvix.

### Importing required packages and definitions¶

In [1]:

```
import biocrnpyler
from biocrnpyler import *
from biocrnpyler.component import Component
from biocrnpyler.chemical_reaction_network import Species, Reaction, ChemicalReactionNetwork
from biocrnpyler.mechanism import Mechanism
from biocrnpyler.reaction import Reaction
from biocrnpyler.species import Complex, Species, WeightedSpecies

import bioscrape

import numpy as np
import pandas as pd
import pylab as plt
from scipy import stats

import libsbml
import time
from typing import List, Union
import csv
import math
import matplotlib.pyplot as plt
import random 
import datetime

#Get directory
import os
directory = os.getcwd()
```

#### Bokeh¶

In [2]:

```
%matplotlib inline
import bokeh.io
import bokeh.plotting
bokeh.io.output_notebook()
from bokeh.themes import Theme

# Modules needed from Bokeh.
from bokeh.io import output_file, show
from bokeh.plotting import gridplot,figure
from bokeh.models import LinearAxis, Range1d

colors2 = bokeh.palettes.Colorblind[8]

try:
    import dnaplotlib as dpl
    dpl_enabled = True
except (ModuleNotFoundError,ImportError) as e:
    dpl_enabled = False
    
from bokeh.io import export_png
```

Loading BokehJS ...

#### Definitions¶

In [3]:

```
#Definition to un-nest lists that will be created by models
def flatten(t):
    return [item for sublist in t for item in sublist]
```

### User defined transcribed unit¶

In [4]:

```
#User will copy the sequence bwteen end of promoter and start of terminator of DNA strain
#mGapt_UTR1_deGFP 
dna_seq= 'GGGATCCCGACTGGCGAGAGCCAGGTAACGAATGGATCCAATAATTTTGTTTAACTTTAAGAAGGAGATATACCATGGAGCTTTTCACTGGCGTTGTTCCCATCCTGGTCGAGCTGGACGGCGACGTAAACGGCCACAAGTTCAGCGTGTCCGGCGAGGGCGAGGGCGATGCCACCTACGGCAAGCTGACCCTGAAGTTCATCTGCACCACCGGCAAGCTGCCCGTGCCCTGGCCCACCCTCGTGACCACCCTGACCTACGGCGTGCAGTGCTTCAGCCGCTACCCCGACCACATGAAGCAGCACGACTTCTTCAAGTCCGCCATGCCCGAAGGCTACGTCCAGGAGCGCACCATCTTCTTCAAGGACGACGGCAACTACAAGACCCGCGCCGAGGTGAAGTTCGAGGGCGACACCCTGGTGAACCGCATCGAGCTGAAGGGCATCGACTTCAAGGAGGACGGCAACATCCTGGGGCACAAGCTGGAGTACAACTACAACAGCCACAACGTCTATATCATGGCCGACAAGCAGAAGAACGGCATCAAGGTGAACTTCAAGATCCGCCACAACATCGAGGACGGCAGCGTGCAGCTCGCCGACCACTACCAGCAGAACACCCCCATCGGCGACGGCCCCGTGCTGCTGCCCGACAACCACTACCTGAGCACCCAGTCCGCCCTGAGCAAAGACCCCAACGAGAAGCGCGATCACATGGTCCTGCTGGAGTTCGTGACCGCCGCCGGGATCTAACTCGAGCCTTAGGAGATCCGGCTGCTAACAAAGCCCGAAAGGAAGCTGAGTTG'
```

### PURE TX Model in BioCRNpyler¶

This model is based on the TX-TL from Tuza *et al.* [1] originally built for cell lysate protein expression. This section of code requires a DNA sequence input from the user. If transcription is not desired, an amino acid list can also serve as the input and this section can be omitted. The steps of the TX are split into 3 subsections: species and reactions based on nucleotide needed (1), generates complement sequence (2), and species and reactions of growing mRNA strand (3).

1. Non-nucleic acid Species and Reactions
2. RNA sequence
3. Addition of nucleotide and individual reactions for the desired DNA

##### Citation:¶

[1] Zoltán A. Tuza, Dan Siegal-Gaskins, Jongmin Kim, and Gábor Szederkényi. Analysis-based parameter estimation of an *in vitro* transcription-translation system. In Eur. Control Conf. 2015, page 1554–1560, 2015. (PNAS, 2015)

#### Non-nucleic acid Species and Reactions¶

Creates all the species and reaction around the nucleotide sequences in the given transcribed regions, common protein and small molecules.

In [5]:

```
## Species
#Define species needed for TX 
T7RNAP = Species('T7RNAP') #ActivePolymerase
DNA = Species("DNA") 
mRNA_i=Species('mRNA_i') # mRNA intermediate species
mRNA_t=Species('mRNA_t') # mRNA species used to track total mRNA production
T7RNAP_bound = Species("T7RNAP_bound") 
T7RNAP_bound_GTP=Species('T7RNAP_bound_GTP')
T7RNAP_bound_GDP_PO4= Species('T7RNAP_bound_GDP_PO4')

#Nucleotide bases that are "active"
ATP=Species('ATP')
GTP=Species('GTP')
CTP=Species('CTP')
UTP=Species('UTP')

CDP=Species('CDP')
UDP=Species('UDP')

#Other small molecules
GDP=Species('GDP')
PPi=Species('PPi')
PO4=Species('PO4')

##Rates
#Degradation rate
NTP_deg=  8.75*10**(-5)
k_rnapbF1 = 0.145 
k_rnapbF2 = 0.0529 
k_rnapbF3 = 0.21

#Define placing all species in one array 
Species_general= [T7RNAP, DNA, T7RNAP_bound_GTP, T7RNAP_bound, 
                  ATP, GTP, CTP, UTP, CDP, UDP,
                  PPi, PO4, mRNA_t, mRNA_i,
                  T7RNAP_bound_GDP_PO4, T7RNAP_bound_GTP,]

#Define Reactions needed for TX regardless of the sequence defined
RXN_general= [
    #NTP degradation
    # Reaction.from_massaction([ATP], [ADP, PO4], k_forward=NTP_deg), #Removed due to redundacy in TL model below
    # Reaction.from_massaction([GTP], [GDP, PO4], k_forward=NTP_deg), #Removed due to redundacy in TL model below
    Reaction.from_massaction([CTP], [CDP, PO4], k_forward=NTP_deg), 
    Reaction.from_massaction([UTP], [UDP, PO4], k_forward=NTP_deg), 
    
    #Binding of RNAP and beginning of transcription
    Reaction.from_massaction([T7RNAP, DNA, GTP], [T7RNAP_bound_GTP], k_forward=k_rnapbF1),
    Reaction.from_massaction([T7RNAP_bound_GTP], [T7RNAP_bound_GDP_PO4], k_forward=k_rnapbF2),
    Reaction.from_massaction([T7RNAP_bound_GDP_PO4], [T7RNAP_bound,GDP, PO4], k_forward=k_rnapbF3)]
```

#### Transcribing DNA sequence to mRNA sequence¶

Transcribed the DNA input into RNA nucleotides. Output is a list of the nucleotides used to make the mRNA.

In [6]:

```
# Transcribed given DNA sequence into mRNA sequence
def getTranscript(dna):
    transcript = []
    for bp in dna:
        if bp == 'A':
            tx = UTP
        elif bp == 'T':
            tx =ATP
        elif bp == 'G':
            tx =CTP
        elif bp == 'C':
            tx =GTP
        else:
            print('Non-standard nucleotides')
            
        transcript.append(tx)

    return transcript
```

In [7]:

```
#Translating RNA sequence from DNA sequence
rna_seq = getTranscript(dna_seq)
# print(rna_seq)
```

#### Addition of nucleotide and individual reactions for the desired DNA¶

Individually adds each nucleotide of the growing mRNA strand.

In [8]:

```
#Reaction rates for transcription
k_start= 0.1039 
k_ntpbound = 0.365 
k_ntpadd = 8.88 
k_ntpdis =  557.64 
k_term= 1.3 

#Start iternations to loop over mRNA sequence
nt_len = len(rna_seq)
mrna_length='0000'
rxn_list=[]
species_list=[]

for L in [l for l in range(len(rna_seq))]:
    ntp = rna_seq[L]
    mrna0 = mrna_length[:-len(str(L))]+str(L) #starting mRNA
    mrnaG = mrna_length[:-len(str(L+1))]+str(L+1) #Growing mRNA
    
    #Initiation of mRNA strand 
    if L == 0:       
        T7RNAP_bound_mRNAmrna0 = Species('T7RNAP_bound_mRNA'+mrna0)
        T7RNAP_bound_mRNAmrna0_ntp = Species('T7RNAP_bound_mRNA'+mrna0+'_'+ str(ntp))
        T7RNAP_bound_mRNAmrnaG_nmp_PPi = Species('T7RNAP_bound_mRNA'+mrnaG+'_PPi')
        T7RNAP_bound_mRNAmrnaG = Species('T7RNAP_bound_mRNA'+mrnaG)
        
        species= [T7RNAP_bound_mRNAmrna0, T7RNAP_bound_mRNAmrna0_ntp, T7RNAP_bound_mRNAmrnaG_nmp_PPi, T7RNAP_bound_mRNAmrnaG]
        
        rxns=[Reaction.from_massaction([T7RNAP_bound],[Species('T7RNAP_bound_mRNA'+mrna0)],k_forward=k_start), #can add DNA, DNA
              Reaction.from_massaction([Species('T7RNAP_bound_mRNA'+mrna0), ntp], [Species('T7RNAP_bound_mRNA'+mrna0+'_'+ str(ntp))], k_forward=k_ntpbound),
              Reaction.from_massaction([Species('T7RNAP_bound_mRNA'+mrna0+'_'+ str(ntp))],[Species('T7RNAP_bound_mRNA'+mrnaG+'_PPi')],k_forward=k_ntpadd),
              Reaction.from_massaction([Species('T7RNAP_bound_mRNA'+mrnaG+'_PPi')],[Species('T7RNAP_bound_mRNA'+mrnaG), PPi],k_forward=k_ntpdis),] 
     
    #Elongation of mRNA strand  
    elif L<nt_len-1:
        T7RNAP_bound_mRNAmrna0_ntp = Species('T7RNAP_bound_mRNA'+mrna0+'_'+ str(ntp))
        T7RNAP_bound_mRNAmrnaG_nmp_PPi = Species('T7RNAP_bound_mRNA'+mrnaG+'_PPi')
        T7RNAP_bound_mRNAmrnaG = Species('T7RNAP_bound_mRNA'+mrnaG)
        
        species= [T7RNAP_bound_mRNAmrna0_ntp, T7RNAP_bound_mRNAmrnaG_nmp_PPi, T7RNAP_bound_mRNAmrnaG]
        
        rxns=[Reaction.from_massaction([Species('T7RNAP_bound_mRNA'+mrna0), ntp], [Species('T7RNAP_bound_mRNA'+mrna0+'_'+ str(ntp))], k_forward=k_ntpbound),
              Reaction.from_massaction([Species('T7RNAP_bound_mRNA'+mrna0+'_'+ str(ntp))],[Species('T7RNAP_bound_mRNA'+mrnaG+'_PPi')],k_forward=k_ntpadd),
              Reaction.from_massaction([Species('T7RNAP_bound_mRNA'+mrnaG+'_PPi')],[Species('T7RNAP_bound_mRNA'+mrnaG), PPi,],k_forward=k_ntpdis),] 
        
    #Termination of mRNA strand      
    else:
        T7RNAP_bound_mRNAmrna0_ntp = Species('T7RNAP_bound_mRNA'+mrna0+'_'+ str(ntp))
        T7RNAP_bound_mRNAmrnaG_nmp_PPi = Species('T7RNAP_bound_mRNA'+mrnaG+'_PPi')
        T7RNAP_bound_mRNAmrnaG = Species('T7RNAP_bound_mRNA'+mrnaG)
        
        species= [T7RNAP_bound_mRNAmrna0_ntp, T7RNAP_bound_mRNAmrnaG_nmp_PPi, T7RNAP_bound_mRNAmrnaG,]
                   
        rxns=[Reaction.from_massaction([Species('T7RNAP_bound_mRNA'+mrna0), ntp], [Species('T7RNAP_bound_mRNA'+mrna0+'_'+ str(ntp))], k_forward=k_ntpbound),
              Reaction.from_massaction([Species('T7RNAP_bound_mRNA'+mrna0+'_'+ str(ntp))],[Species('T7RNAP_bound_mRNA'+mrnaG+'_PPi')],k_forward=k_ntpadd),  
              Reaction.from_massaction([Species('T7RNAP_bound_mRNA'+mrnaG+'_PPi')],[Species('T7RNAP_bound_mRNA'+mrnaG), PPi],k_forward=k_ntpdis),
              Reaction.from_massaction([Species('T7RNAP_bound_mRNA'+mrnaG),],[T7RNAP,mRNA_i, DNA, mRNA_t],k_forward=k_term)]
        
    #Appending each reaction and species to the one before       
    rxn_list.append(rxns)
    species_list.append(species)

#Flattening to th nested lists 
species_list = flatten(species_list)
rxn_list= flatten(rxn_list)
```

#### Compiling TX CRN (CRN\_TX)¶

In [9]:

```
#List of all species and reactions
All_species_TX = flatten([Species_general, species_list])
All_rxn_TX = flatten([RXN_general, rxn_list])

#IC for Transcription only reactions
initial_con={'T7RNAP':(1), 'DNA':(.005), 'ATP':(3750), 'GTP':(2500), 'CTP':(1250), 'UTP':(1250),} #in uM

#Buliding the CRN_TX and saving as a SBML file
CRN_TX = ChemicalReactionNetwork(species = All_species_TX, reactions = All_rxn_TX)
# CRN_TX.write_sbml_file("PURE_TX_Model_degfp.xml")
```

#### Simulations of TX¶

In [10]:

```
# Run simulation
t0_tx=time.time()

timepoints = np.linspace(10**-5, 3*10**4, 500)
Model_TX = CRN_TX.simulate_with_bioscrape_via_sbml(timepoints = timepoints, initial_condition_dict = initial_con)

t1_tx=time.time()
print(f'TX Simulation total time= {t1_tx-t0_tx} s')
```

```
TX Simulation total time= 108.61792969703674 s
```

##### Plot¶

In [11]:

```
pTXonly = bokeh.plotting.figure()
pTXonly.title.text = "mRNA Concentration Over Time"
pTXonly.xaxis.axis_label = 'Time (hours)'
pTXonly.yaxis.axis_label = 'mRNA (uM)'

pTXonly.line(timepoints/3600, Model_TX[str(mRNA_t)], legend_label = "mRNA_t", line_color="magenta", line_width=5, line_alpha=.5) #mRNA transcribed 
pTXonly.line(timepoints/3600, Model_TX[str(mRNA_i)], legend_label = "mRNA_i", line_color="orange", line_width=5,line_alpha=.5) #mRNA transcribed 

pTXonly.legend.click_policy = "hide"
pTXonly.legend.location='top_left'
pTXonly.x_range=Range1d(0, 6)
bokeh.io.show(pTXonly)
```

### PURE TL Model in BioCRNpyler¶

This model is based on the PURE TL models from Matsuura *et al.* [2,3] originally modeled in MATLAB. This section of code uses the amino acids list derived from the DNA sequence input to generate all species and reactions associated with desired protein. If transcription is not wanted, an amino acid list can also serve as the input. The PURE TL does not account for multiple tRNAs coding for the same amino acid.

For order and simplicity, the reactions were broken up into 6 subsections: species and reactions based on amino acids needed (1-2) and species and reactions of growing peptide (3-6).

1. Non-nucleic acid Species and Reactions
2. Addition of AA individual reactions
3. Met Species and Reaction (Start codon)
4. Elongation
5. Termination (End codon)
6. Removal of Ribosome

##### Citation:¶

[2] Matsuura, T., Tanimura, N., Hosoda, K., and Shimizu, Y. Reaction dynamics analysis of a reconstituted *Escherichia coli* protein translation system by computational modeling. PNAS. 2017, 114, 8. (PNAS, 2017) \
[3] Matsuura, T., Hosoda, K., and Shimizu, Y. Robustness of a reconstituted *Escherichia coli* protein translation system analyzed by computational modeling. ACS Synth. Biol. 2018, 7, 8, 1964–1972. (ACS Synth. Biol., 2018)

#### Define the directory where parameters and initial conditions are saved as a CSV.¶

In [12]:

```
#Directory and file for the reaction rates
filename_parameters = '/fMGG_synthesis_parameters_CRN.csv'

with open(directory + filename_parameters, mode='r') as infile:
    reader = csv.reader(infile)
    rxn_k= {rows[0]:float(rows[1]) for rows in reader}
```

#### Translating DNA sequence to peptide amino acid list¶

Converting DNA sequence into amino acid list for translation model

In [13]:

```
#DNA code chart for coding domain
def translate(seq): 
    table = {
        'ATA':'Ile', 'ATC':'Ile', 'ATT':'Ile', 'ATG':'Met',
        'ACA':'Thr', 'ACC':'Thr', 'ACG':'Thr', 'ACT':'Thr',
        'AAC':'Asn', 'AAT':'Asn', 'AAA':'Lys', 'AAG':'Lys',
        'AGC':'Ser', 'AGT':'Ser', 'AGA':'Arg', 'AGG':'Arg',                 
        'CTA':'Leu', 'CTC':'Leu', 'CTG':'Leu', 'CTT':'Leu',
        'CCA':'Pro', 'CCC':'Pro', 'CCG':'Pro', 'CCT':'Pro',
        'CAC':'His', 'CAT':'His', 'CAA':'Gln', 'CAG':'Gln',
        'CGA':'Arg', 'CGC':'Arg', 'CGG':'Arg', 'CGT':'Arg',
        'GTA':'Val', 'GTC':'Val', 'GTG':'Val', 'GTT':'Val',
        'GCA':'Ala', 'GCC':'Ala', 'GCG':'Ala', 'GCT':'Ala',
        'GAC':'Asp', 'GAT':'Asp', 'GAA':'Glu', 'GAG':'Glu',
        'GGA':'Gly', 'GGC':'Gly', 'GGG':'Gly', 'GGT':'Gly',
        'TCA':'Ser', 'TCC':'Ser', 'TCG':'Ser', 'TCT':'Ser',
        'TTC':'Phe', 'TTT':'Phe', 'TTA':'Leu', 'TTG':'Leu',
        'TAC':'Tyr', 'TAT':'Tyr', 'TAA':'_', 'TAG':'_',
        'TGC':'Cys', 'TGT':'Cys', 'TGA':'_', 'TGG':'Trp', }
    
    protein =[]
    if len(seq)%3 == 0:
        for i in range(0, len(seq), 3):
            codon = seq[i:i + 3]
            if table[codon] != '_':
                protein+= [table[codon]]
            else:
                break
    return protein

#Translation of CDS
def coding_protein(dna, start=0):
    for bp in range(start,len(dna)-2):
        bp_aa = dna_seq[bp:bp+3]
        
        if bp_aa=='ATG':
            start = bp
            remainder= len(dna[bp:]) % 3
            
            if remainder ==0:
                CDS= dna
            else:
                CDS= dna[bp:-remainder]
            
            translation= translate(CDS)
            
            if len(translation)>0:
                return translation
                break
            
        if bp == len(dna)-2:
            print('No start codon found')
```

In [14]:

```
#Translating protein sequence from DNA sequence
protein=coding_protein(dna_seq)
# print(protein,len(protein))
```

#### Non-nucleic acid Species and Reactions¶

Creates all the species and reaction around the amino acids in the given chain, common protein and small molecules.

In [15]:

```
# Makes a list of the aa needed in the given amino acid chain, removing any duplicates.
AA= list(set(protein[1:],)) #removes any duplicated amino acids

#Makes empty array for species and reactions
list_of_reactions = []
list_species_aa=[]

#Reference lists for aa and respective codon.
list_AA =['Ala', 'Arg', 'Asn', 'Asp', 'Cys', 
          'Gln', 'Glu', 'Gly', 'His', 'Ile', 
          'Leu', 'Lys', 'Met', 'Phe', 'Pro',
          'Ser', 'Thr', 'Trp', 'Tyr', 'Val']
list_condon =['AGU', 'CCG', 'AUU','AUC','ACA',
             'UUG','UUC','GCC','AUG','AAU',
             'CAG','CUU','CAU','GAA','AGG',
             'GCU','AGU','CCA','AUA','CAC',]
```

##### General proteins and small molecules¶

In [16]:

```
#Make species for general proteins and small molecules
mRNA=Species('mRNA')
ADP=Species('ADP')
GDP=Species('GDP')
AMP=Species('AMP')
GMP=Species('GMP')
CMP=Species('CMP')
UMP=Species('UMP')

CK = Species('CK')
CK_ADP = Species('CK_ADP')
CK_ATP = Species('CK_ATP')
CK_CP = Species('CK_CP')
CK_CP_ADP = Species('CK_CP_ADP')
CK_Cr = Species('CK_Cr')
CK_Cr_ATP = Species('CK_Cr_ATP')

CK_degraded = Species('CK_degraded')
CP = Species('CP')
Cr = Species('Cr')
EFG = Species('EFG')
EFG_degraded = Species('EFG_degraded')
EFG_GDP = Species('EFG_GDP')
EFG_GTP = Species('EFG_GTP')
EFTs = Species('EFTs')
EFTs_degraded = Species('EFTs_degraded')
EFTu = Species('EFTu')

EFTu_degraded = Species('EFTu_degraded')
EFTu_EFTs = Species('EFTu_EFTs')
EFTu_GDP = Species('EFTu_GDP')
EFTu_GDP_EFTs = Species('EFTu_GDP_EFTs')
EFTu_GTP = Species('EFTu_GTP')
EFTu_GTP_EFTs = Species('EFTu_GTP_EFTs')
FD = Species('FD')

IF1 = Species('IF1')
IF1_degraded = Species('IF1_degraded')
IF2 = Species('IF2')
IF2_degraded = Species('IF2_degraded')
IF2_GDP = Species('IF2_GDP')
IF2_GTP = Species('IF2_GTP')
IF2_GTP_fMettRNAfMetCAU = Species('IF2_GTP_fMettRNAfMetCAU')
IF3 = Species('IF3')
IF3_degraded = Species('IF3_degraded')
MK = Species('MK')

MK_ADP_1 = Species('MK_ADP_1')
MK_ADP_2 = Species('MK_ADP_2')
MK_ADP_ADP = Species('MK_ADP_ADP')
MK_AMP = Species('MK_AMP')
MK_ATP = Species('MK_ATP')
MK_ATP_AMP = Species('MK_ATP_AMP')
MK_degraded = Species('MK_degraded')
mRNA_degraded = Species('mRNA_degraded')
MTF = Species('MTF')

MTF_degraded = Species('MTF_degraded')
MTF_FD = Species('MTF_FD')
MTF_THF = Species('MTF_THF')
NDK = Species('NDK')
NDK_ADP = Species('NDK_ADP')
NDK_ATP = Species('NDK_ATP')
NDK_degraded = Species('NDK_degraded')
NDK_GDP = Species('NDK_GDP')
NDK_GDP_ATP = Species('NDK_GDP_ATP')
NDK_GTP = Species('NDK_GTP')

NDK_GTP_ADP = Species('NDK_GTP_ADP')
PPiase = Species('PPiase')
PPiase_degraded = Species('PPiase_degraded')
PPiase_PO4 = Species('PPiase_PO4')
PPiase_PO4_PO4 = Species('PPiase_PO4_PO4')
PPiase_PPi = Species('PPiase_PPi')
RF1 = Species('RF1')
RF1_degraded = Species('RF1_degraded')

RF2 = Species('RF2')
RF2_degraded = Species('RF2_degraded')
RF3 = Species('RF3')
RF3_degraded = Species('RF3_degraded')
RF3_GDP = Species('RF3_GDP')
RF3_GTP = Species('RF3_GTP')
RRF = Species('RRF')
RRF_degraded = Species('RRF_degraded')

RS30S = Species('RS30S')
RS30S_degraded = Species('RS30S_degraded')
RS30S_IF1_IF3_IF2_GTP_mRNA = Species('RS30S_IF1_IF3_IF2_GTP_mRNA')
RS30S_IF1_IF3_mRNA = Species('RS30S_IF1_IF3_mRNA')
RS30S_IF1_mRNA = Species('RS30S_IF1_mRNA')
RS30S_IF1 = Species('RS30S_IF1')
RS30S_IF1_IF2_GTP = Species('RS30S_IF1_IF2_GTP')
RS30S_IF1_IF2_GTP_mRNA = Species('RS30S_IF1_IF2_GTP_mRNA')
RS30S_IF1_IF3 = Species('RS30S_IF1_IF3')
RS30S_IF1_IF3_IF2_GTP = Species('RS30S_IF1_IF3_IF2_GTP')
RS30S_IF3_IF2_GTP = Species('RS30S_IF3_IF2_GTP')

RS30S_IF3_IF2_GTP_mRNA = Species('RS30S_IF3_IF2_GTP_mRNA')      
RS30S_IF2_GTP = Species('RS30S_IF2_GTP')
RS30S_IF2_GTP_mRNA = Species('RS30S_IF2_GTP_mRNA')
RS30S_IF3 = Species('RS30S_IF3')
RS30S_IF3_mRNA = Species('RS30S_IF3_mRNA')
RS30S_mRNA = Species('RS30S_mRNA')
RS50S = Species('RS50S')
RS50S_degraded = Species('RS50S_degraded')
RS50S_EFG_GDP = Species('RS50S_EFG_GDP')
RS50S_EFG_GDP_PO4 = Species('RS50S_EFG_GDP_PO4')

RS50S_EFG_GTP = Species('RS50S_EFG_GTP')
RS50S_RRF = Species('RS50S_RRF')
RS50S_RRF_EFG_GDP = Species('RS50S_RRF_EFG_GDP')
RS70S = Species('RS70S')
RS70S_EFG_GDP = Species('RS70S_EFG_GDP')
RS70S_EFG_GDP_PO4 = Species('RS70S_EFG_GDP_PO4')
RS70S_EFG_GTP = Species('RS70S_EFG_GTP')
RS70S_IF1 = Species('RS70S_IF1')
RS70S_IF3 = Species('RS70S_IF3')
RS70S_IF1_IF3 = Species('RS70S_IF1_IF3')         

THF = Species('THF')
fMet = Species('fMet')
fMet_degraded = Species('fMet_degraded')

#######################################################################################################################

#Creates a list of all the general species
list_species_gen=[ADP, AMP, ATP, CK, CK_ADP, CK_ATP, 
                  CK_CP, CK_CP_ADP, CK_Cr, CK_Cr_ATP, CK_degraded,
                  CP, Cr, EFG, EFG_degraded, EFG_GDP,
                  EFG_GTP, EFTs, EFTs_degraded, EFTu, EFTu_degraded,
                  EFTu_EFTs, EFTu_GDP, EFTu_GDP_EFTs, EFTu_GTP, EFTu_GTP_EFTs,
                  FD, GDP, GMP, GTP, IF1,
                  IF1_degraded, IF2, IF2_degraded, IF2_GDP, IF2_GTP,
                  IF2_GTP_fMettRNAfMetCAU, IF3, IF3_degraded, MK, MK_ADP_1, MK_ADP_2,
                  MK_ADP_ADP, MK_AMP, MK_ATP, MK_ATP_AMP, MK_degraded,
                  mRNA, mRNA_degraded, MTF, MTF_degraded, MTF_FD, MTF_THF,
                  NDK, NDK_ADP, NDK_ATP, NDK_degraded, NDK_GDP, NDK_GDP_ATP,
                  NDK_GTP, NDK_GTP_ADP, PO4, PPi, PPiase, PPiase_degraded,
                  PPiase_PO4, PPiase_PO4_PO4, PPiase_PPi, RF1, RF1_degraded,
                  RF2, RF2_degraded, RF3, RF3_degraded, RF3_GDP,
                  RF3_GTP, RRF, RRF_degraded, RS30S, RS30S_degraded,
                  RS30S_IF1_IF3_IF2_GTP_mRNA, RS30S_IF1_IF3_mRNA, RS30S_IF1_mRNA, RS30S_IF1, RS30S_IF1_IF2_GTP,
                  RS30S_IF1_IF2_GTP_mRNA, RS30S_IF1_IF3, RS30S_IF1_IF3_IF2_GTP, RS30S_IF3_IF2_GTP_mRNA,   
                  RS30S_IF2_GTP, RS30S_IF2_GTP_mRNA, RS30S_IF3, RS30S_IF3_mRNA, RS30S_IF3_IF2_GTP,
                  RS30S_mRNA, RS50S, RS50S_degraded, RS50S_EFG_GDP, RS50S_EFG_GDP_PO4,
                  RS50S_EFG_GTP, RS50S_RRF, RS50S_RRF_EFG_GDP, RS70S, RS70S_EFG_GDP,
                  RS70S_EFG_GDP_PO4, RS70S_EFG_GTP, RS70S_IF1, RS70S_IF3, RS70S_IF1_IF3,        
                  THF, fMet, fMet_degraded,]

#######################################################################################################################

#Creates a list of all the reactions only involving the general species. The reaction rates are read from the rxn_k dictionary created initially.
list_of_reaction_gen= [    
    Reaction.from_massaction([EFTu,EFTs],[EFTu_EFTs], k_forward = rxn_k['re0000000261_k1']),
    Reaction.from_massaction([EFTu_EFTs],[EFTu,EFTs], k_forward = rxn_k['re0000000262_k1']),
    Reaction.from_massaction([EFTu_EFTs,GDP],[EFTu_GDP_EFTs], k_forward = rxn_k['re0000000263_k1']),
    Reaction.from_massaction([EFTu_GDP_EFTs],[EFTu_EFTs,GDP], k_forward = rxn_k['re0000000264_k1']),
    Reaction.from_massaction([EFTu_GDP_EFTs],[EFTu_GDP,EFTs], k_forward = rxn_k['re0000000265_k1']),
    Reaction.from_massaction([EFTu_GDP,EFTs],[EFTu_GDP_EFTs], k_forward = rxn_k['re0000000266_k1']),
    Reaction.from_massaction([EFTu_GDP],[EFTu,GDP], k_forward = rxn_k['re0000000267_k1']),
    Reaction.from_massaction([EFTu,GDP],[EFTu_GDP], k_forward = rxn_k['re0000000268_k1']),
    Reaction.from_massaction([EFTu_EFTs,GTP],[EFTu_GTP_EFTs], k_forward = rxn_k['re0000000269_k1']),
    Reaction.from_massaction([EFTu_GTP_EFTs],[EFTu_EFTs,GTP], k_forward = rxn_k['re0000000270_k1']),
    Reaction.from_massaction([EFTu_GTP_EFTs],[EFTu_GTP,EFTs], k_forward = rxn_k['re0000000271_k1']),
    Reaction.from_massaction([EFTu_GTP,EFTs],[EFTu_GTP_EFTs], k_forward = rxn_k['re0000000272_k1']),
    Reaction.from_massaction([EFTu,GTP],[EFTu_GTP], k_forward = rxn_k['re0000000273_k1']),
    Reaction.from_massaction([EFTu_GTP],[EFTu,GTP], k_forward = rxn_k['re0000000274_k1']),

    Reaction.from_massaction([EFG_GDP],[EFG,GDP], k_forward = rxn_k['re0000000292_k1']),
    Reaction.from_massaction([EFG,GDP],[EFG_GDP], k_forward = rxn_k['re0000000293_k1']),
    Reaction.from_massaction([EFG,GTP],[EFG_GTP], k_forward = rxn_k['re0000000294_k1']),
    Reaction.from_massaction([EFG_GTP],[EFG,GTP], k_forward = rxn_k['re0000000295_k1']),

    Reaction.from_massaction([RS50S,EFG_GTP],[RS50S_EFG_GTP], k_forward = rxn_k['re0000000298_k1']),
    Reaction.from_massaction([RS50S_EFG_GTP],[RS50S,EFG_GTP], k_forward = rxn_k['re0000000299_k1']),
    Reaction.from_massaction([RS70S,EFG_GTP],[RS70S_EFG_GTP], k_forward = rxn_k['re0000000300_k1']),
    Reaction.from_massaction([RS70S_EFG_GTP],[RS70S,EFG_GTP], k_forward = rxn_k['re0000000301_k1']),

    Reaction.from_massaction([RS50S_EFG_GTP],[RS50S_EFG_GDP_PO4], k_forward = rxn_k['re0000000302_k1']),
    Reaction.from_massaction([RS50S_EFG_GDP_PO4],[RS50S_EFG_GTP], k_forward = rxn_k['re0000000303_k1']),
    Reaction.from_massaction([RS70S_EFG_GTP],[RS70S_EFG_GDP_PO4], k_forward = rxn_k['re0000000304_k1']),
    Reaction.from_massaction([RS70S_EFG_GDP_PO4],[RS70S_EFG_GTP], k_forward = rxn_k['re0000000305_k1']),
    Reaction.from_massaction([RS50S_EFG_GDP_PO4],[RS50S_EFG_GDP,PO4,], k_forward = rxn_k['re0000000306_k1']),
    Reaction.from_massaction([RS70S_EFG_GDP_PO4],[RS70S_EFG_GDP,PO4,], k_forward = rxn_k['re0000000307_k1']),
    Reaction.from_massaction([RS50S_EFG_GDP],[RS50S,EFG_GDP], k_forward = rxn_k['re0000000308_k1']),
    Reaction.from_massaction([RS70S_EFG_GDP],[RS70S,EFG_GDP], k_forward = rxn_k['re0000000309_k1']),

    Reaction.from_massaction([CK,ADP],[CK_ADP], k_forward = rxn_k['re0000000330_k1']),
    Reaction.from_massaction([CK_ADP],[CK,ADP], k_forward = rxn_k['re0000000331_k1']),

    Reaction.from_massaction([CK,CP],[CK_CP], k_forward = rxn_k['re0000000332_k1']),
    Reaction.from_massaction([CK_CP],[CK,CP], k_forward = rxn_k['re0000000333_k1']),
    Reaction.from_massaction([CK_CP,ADP],[CK_CP_ADP], k_forward = rxn_k['re0000000334_k1']),
    Reaction.from_massaction([CK_CP_ADP],[CK_CP,ADP], k_forward = rxn_k['re0000000335_k1']),
    Reaction.from_massaction([CK_ADP,CP],[CK_CP_ADP], k_forward = rxn_k['re0000000336_k1']),
    Reaction.from_massaction([CK_CP_ADP],[CP,CK_ADP], k_forward = rxn_k['re0000000337_k1']),
    Reaction.from_massaction([CK_CP_ADP],[CK_Cr_ATP], k_forward = rxn_k['re0000000338_k1']),
    Reaction.from_massaction([CK_Cr_ATP],[CK_CP_ADP], k_forward = rxn_k['re0000000339_k1']),
    Reaction.from_massaction([CK_Cr_ATP],[CK_Cr,ATP], k_forward = rxn_k['re0000000340_k1']),
    Reaction.from_massaction([CK_Cr,ATP],[CK_Cr_ATP], k_forward = rxn_k['re0000000341_k1']),

    Reaction.from_massaction([CK_Cr_ATP],[CK_ATP,Cr], k_forward = rxn_k['re0000000342_k1']),
    Reaction.from_massaction([CK_ATP,Cr],[CK_Cr_ATP], k_forward = rxn_k['re0000000343_k1']),
    Reaction.from_massaction([CK_ATP],[CK,ATP], k_forward = rxn_k['re0000000344_k1']),
    Reaction.from_massaction([CK,ATP],[CK_ATP], k_forward = rxn_k['re0000000345_k1']),
    Reaction.from_massaction([CK_Cr],[CK,Cr], k_forward = rxn_k['re0000000346_k1']),
    Reaction.from_massaction([CK,Cr],[CK_Cr], k_forward = rxn_k['re0000000347_k1']),

    Reaction.from_massaction([NDK,ATP],[NDK_ATP], k_forward = rxn_k['re0000000355_k1']),
    Reaction.from_massaction([NDK_ATP],[NDK,ATP], k_forward = rxn_k['re0000000356_k1']),
    Reaction.from_massaction([NDK,GDP],[NDK_GDP], k_forward = rxn_k['re0000000357_k1']),
    Reaction.from_massaction([NDK_GDP],[NDK,GDP], k_forward = rxn_k['re0000000358_k1']),
    Reaction.from_massaction([NDK_GDP,ATP],[NDK_GDP_ATP], k_forward = rxn_k['re0000000359_k1']),
    Reaction.from_massaction([NDK_GDP_ATP],[NDK_GDP,ATP], k_forward = rxn_k['re0000000360_k1']),
    Reaction.from_massaction([NDK_ATP,GDP],[NDK_GDP_ATP], k_forward = rxn_k['re0000000361_k1']),

    Reaction.from_massaction([NDK_GDP_ATP],[NDK_ATP,GDP], k_forward = rxn_k['re0000000362_k1']),
    Reaction.from_massaction([NDK_GDP_ATP],[NDK_GTP_ADP], k_forward = rxn_k['re0000000363_k1']),

    Reaction.from_massaction([NDK_GTP_ADP],[NDK_ADP,GTP], k_forward = rxn_k['re0000000365_k1']),
    Reaction.from_massaction([NDK_GTP_ADP],[NDK_GTP,ADP], k_forward = rxn_k['re0000000366_k1']),
    Reaction.from_massaction([NDK_ADP],[NDK,ADP], k_forward = rxn_k['re0000000367_k1']),
    Reaction.from_massaction([NDK_GTP],[NDK,GTP], k_forward = rxn_k['re0000000368_k1']),

    Reaction.from_massaction([NDK_GTP,ADP],[NDK_GTP_ADP], k_forward = rxn_k['re0000000375_k1']),
    Reaction.from_massaction([NDK_ADP,GTP],[NDK_GTP_ADP], k_forward = rxn_k['re0000000376_k1']),
    Reaction.from_massaction([NDK,GTP],[NDK_GTP], k_forward = rxn_k['re0000000377_k1']),
    Reaction.from_massaction([NDK,ADP],[NDK_ADP], k_forward = rxn_k['re0000000378_k1']),

    Reaction.from_massaction([MK,ATP],[MK_ATP], k_forward = rxn_k['re0000000380_k1']),
    Reaction.from_massaction([MK_ATP],[ATP,MK], k_forward = rxn_k['re0000000381_k1']),

    Reaction.from_massaction([MK,AMP],[MK_AMP], k_forward = rxn_k['re0000000382_k1']),
    Reaction.from_massaction([MK_AMP],[MK,AMP], k_forward = rxn_k['re0000000383_k1']),
    Reaction.from_massaction([MK_AMP,ATP],[MK_ATP_AMP], k_forward = rxn_k['re0000000384_k1']),
    Reaction.from_massaction([MK_ATP_AMP],[ATP,MK_AMP], k_forward = rxn_k['re0000000385_k1']),
    Reaction.from_massaction([MK_ATP,AMP],[MK_ATP_AMP], k_forward = rxn_k['re0000000386_k1']),
    Reaction.from_massaction([MK_ATP_AMP],[MK_ATP,AMP], k_forward = rxn_k['re0000000387_k1']),
    Reaction.from_massaction([MK_ATP_AMP],[MK_ADP_ADP], k_forward = rxn_k['re0000000388_k1']),
    Reaction.from_massaction([MK_ADP_ADP],[MK_ATP_AMP], k_forward = rxn_k['re0000000389_k1']),
    Reaction.from_massaction([MK_ADP_ADP],[MK_ADP_1,ADP], k_forward = rxn_k['re0000000390_k1']),
    Reaction.from_massaction([MK_ADP_1,ADP],[MK_ADP_ADP], k_forward = rxn_k['re0000000391_k1']),

    Reaction.from_massaction([MK_ADP_ADP],[MK_ADP_2,ADP], k_forward = rxn_k['re0000000392_k1']),
    Reaction.from_massaction([MK_ADP_2,ADP],[MK_ADP_ADP], k_forward = rxn_k['re0000000393_k1']),
    Reaction.from_massaction([MK_ADP_1],[MK,ADP], k_forward = rxn_k['re0000000394_k1']),
    Reaction.from_massaction([MK,ADP],[MK_ADP_1], k_forward = rxn_k['re0000000395_k1']),
    Reaction.from_massaction([MK_ADP_2],[ADP,MK], k_forward = rxn_k['re0000000396_k1']),
    Reaction.from_massaction([MK,ADP],[MK_ADP_2], k_forward = rxn_k['re0000000397_k1']),

    Reaction.from_massaction([PPiase,PPi],[PPiase_PPi], k_forward = rxn_k['re0000000405_k1']),
    Reaction.from_massaction([PPiase_PPi],[PPiase,PPi], k_forward = rxn_k['re0000000406_k1']),
    Reaction.from_massaction([PPiase_PPi],[PPiase_PO4_PO4], k_forward = rxn_k['re0000000407_k1']),
    Reaction.from_massaction([PPiase_PO4_PO4],[PPiase_PPi], k_forward = rxn_k['re0000000408_k1']),
    Reaction.from_massaction([PPiase_PO4_PO4],[PPiase_PO4,PO4], k_forward = rxn_k['re0000000409_k1']),
    Reaction.from_massaction([PPiase_PO4,PO4],[PPiase_PO4_PO4], k_forward = rxn_k['re0000000410_k1']),
    Reaction.from_massaction([PPiase_PO4],[PPiase,PO4], k_forward = rxn_k['re0000000411_k1']),

    Reaction.from_massaction([PPiase,PO4],[PPiase_PO4], k_forward = rxn_k['re0000000412_k1']),

    Reaction.from_massaction([MTF,FD],[MTF_FD], k_forward = rxn_k['re0000000418_k1']),
    Reaction.from_massaction([MTF_FD],[MTF,FD], k_forward = rxn_k['re0000000419_k1']),
    Reaction.from_massaction([MTF_THF],[MTF,THF], k_forward = rxn_k['re0000000434_k1']),

    Reaction.from_massaction([IF2,GTP],[IF2_GTP], k_forward = rxn_k['re0000000445_k1']),
    Reaction.from_massaction([IF2_GTP],[GTP,IF2], k_forward = rxn_k['re0000000446_k1']),
    Reaction.from_massaction([IF2,GDP],[IF2_GDP], k_forward = rxn_k['re0000000447_k1']),
    Reaction.from_massaction([IF2_GDP],[IF2,GDP], k_forward = rxn_k['re0000000448_k1']),

    Reaction.from_massaction([RS70S],[RS30S,RS50S], k_forward = rxn_k['re0000000455_k1']),
    Reaction.from_massaction([RS30S,RS50S],[RS70S], k_forward = rxn_k['re0000000456_k1']),
    Reaction.from_massaction([RS70S,IF3],[RS70S_IF3], k_forward = rxn_k['re0000000457_k1']),
    Reaction.from_massaction([RS70S_IF3],[RS70S,IF3], k_forward = rxn_k['re0000000458_k1']),
    Reaction.from_massaction([RS30S,IF3],[RS30S_IF3], k_forward = rxn_k['re0000000459_k1']),
    Reaction.from_massaction([RS30S_IF3],[RS30S,IF3], k_forward = rxn_k['re0000000460_k1']),
    Reaction.from_massaction([RS70S_IF3],[RS30S_IF3,RS50S], k_forward = rxn_k['re0000000461_k1']),
    Reaction.from_massaction([RS30S_IF3,RS50S],[RS70S_IF3], k_forward = rxn_k['re0000000462_k1']),
    Reaction.from_massaction([RS30S_IF3,IF2_GTP],[RS30S_IF3_IF2_GTP], k_forward = rxn_k['re0000000463_k1']),
    Reaction.from_massaction([RS30S_IF3_IF2_GTP],[RS30S_IF3,IF2_GTP], k_forward = rxn_k['re0000000464_k1']),

    Reaction.from_massaction([RS30S_IF3,mRNA],[RS30S_IF3_mRNA], k_forward= rxn_k['re0000000469_k1']),
    Reaction.from_massaction([RS30S_IF3_mRNA],[RS30S_IF3,mRNA], k_forward= rxn_k['re0000000470_k1']),
    Reaction.from_massaction([RS30S_IF3_mRNA,IF2_GTP],[RS30S_IF3_IF2_GTP_mRNA], k_forward= rxn_k['re0000000471_k1']),
    Reaction.from_massaction([RS30S_IF3_IF2_GTP_mRNA],[RS30S_IF3_mRNA,IF2_GTP], k_forward= rxn_k['re0000000472_k1']),
    Reaction.from_massaction([RS30S_IF3_IF2_GTP,mRNA],[RS30S_IF3_IF2_GTP_mRNA], k_forward= rxn_k['re0000000481_k1']),
    Reaction.from_massaction([RS30S_IF3_IF2_GTP_mRNA],[RS30S_IF3_IF2_GTP,mRNA], k_forward= rxn_k['re0000000482_k1']),
    Reaction.from_massaction([RS70S_IF1],[RS30S_IF1,RS50S], k_forward = rxn_k['re0000000487_k1']),
    Reaction.from_massaction([RS30S_IF1,RS50S],[RS70S_IF1], k_forward = rxn_k['re0000000488_k1']),
    Reaction.from_massaction([RS70S_IF1,IF3],[RS70S_IF1_IF3], k_forward = rxn_k['re0000000489_k1']),
    Reaction.from_massaction([RS70S_IF1_IF3],[RS70S_IF1,IF3], k_forward = rxn_k['re0000000490_k1']),

    Reaction.from_massaction([RS30S_IF1,IF3],[RS30S_IF1_IF3], k_forward = rxn_k['re0000000491_k1']),
    Reaction.from_massaction([RS30S_IF1_IF3],[RS30S_IF1,IF3], k_forward = rxn_k['re0000000492_k1']),
    Reaction.from_massaction([RS70S_IF1_IF3],[RS30S_IF1_IF3,RS50S], k_forward = rxn_k['re0000000493_k1']),
    Reaction.from_massaction([RS30S_IF1_IF3,RS50S],[RS70S_IF1_IF3], k_forward = rxn_k['re0000000494_k1']),
    Reaction.from_massaction([RS30S_IF1_IF3,IF2_GTP],[RS30S_IF1_IF3_IF2_GTP], k_forward = rxn_k['re0000000495_k1']),
    Reaction.from_massaction([RS30S_IF1_IF3_IF2_GTP],[RS30S_IF1_IF3,IF2_GTP], k_forward = rxn_k['re0000000496_k1']),
    Reaction.from_massaction([RS70S,IF1],[RS70S_IF1], k_forward= rxn_k['re0000000501_k1']),
    Reaction.from_massaction([RS70S_IF1],[RS70S,IF1], k_forward = rxn_k['re0000000502_k1']),
    Reaction.from_massaction([RS30S,IF1],[RS30S_IF1], k_forward = rxn_k['re0000000503_k1']),
    Reaction.from_massaction([RS30S_IF1],[RS30S,IF1], k_forward = rxn_k['re0000000504_k1']),

    Reaction.from_massaction([RS70S_IF3,IF1],[RS70S_IF1_IF3], k_forward = rxn_k['re0000000505_k1']),
    Reaction.from_massaction([RS70S_IF1_IF3],[RS70S_IF3,IF1], k_forward = rxn_k['re0000000506_k1']),
    Reaction.from_massaction([RS30S_IF3,IF1],[RS30S_IF1_IF3], k_forward= rxn_k['re0000000507_k1']),
    Reaction.from_massaction([RS30S_IF1_IF3],[RS30S_IF3,IF1], k_forward = rxn_k['re0000000508_k1']),
    Reaction.from_massaction([RS30S_IF3_IF2_GTP,IF1],[RS30S_IF1_IF3_IF2_GTP], k_forward= rxn_k['re0000000509_k1']),
    Reaction.from_massaction([RS30S_IF1_IF3_IF2_GTP],[RS30S_IF3_IF2_GTP,IF1], k_forward = rxn_k['re0000000510_k1']),
    Reaction.from_massaction([RS30S_IF1_IF3,mRNA],[RS30S_IF1_IF3_mRNA], k_forward = rxn_k['re0000000513_k1']),
    Reaction.from_massaction([RS30S_IF1_IF3_mRNA],[RS30S_IF1_IF3,mRNA], k_forward = rxn_k['re0000000514_k1']),
    Reaction.from_massaction([RS30S_IF1_IF3_mRNA,IF2_GTP],[RS30S_IF1_IF3_IF2_GTP_mRNA], k_forward = rxn_k['re0000000515_k1']),
    Reaction.from_massaction([RS30S_IF1_IF3_IF2_GTP_mRNA],[RS30S_IF1_IF3_mRNA,IF2_GTP], k_forward = rxn_k['re0000000516_k1']),
    Reaction.from_massaction([RS30S_IF1_IF3_IF2_GTP,mRNA],[RS30S_IF1_IF3_IF2_GTP_mRNA], k_forward = rxn_k['re0000000525_k1']),

    Reaction.from_massaction([RS30S_IF1_IF3_IF2_GTP_mRNA],[RS30S_IF1_IF3_IF2_GTP,mRNA], k_forward = rxn_k['re0000000526_k1']),
    Reaction.from_massaction([RS30S_IF3_mRNA,IF1],[RS30S_IF1_IF3_mRNA], k_forward = rxn_k['re0000000531_k1']),
    Reaction.from_massaction([RS30S_IF1_IF3_mRNA],[RS30S_IF3_mRNA,IF1], k_forward = rxn_k['re0000000532_k1']),
    Reaction.from_massaction([RS30S_IF3_IF2_GTP_mRNA,IF1],[RS30S_IF1_IF3_IF2_GTP_mRNA], k_forward= rxn_k['re0000000535_k1']),
    Reaction.from_massaction([RS30S_IF1_IF3_IF2_GTP_mRNA],[RS30S_IF3_IF2_GTP_mRNA,IF1], k_forward = rxn_k['re0000000536_k1']),

    Reaction.from_massaction([RS30S_IF2_GTP],[IF2_GTP,RS30S], k_forward= rxn_k['re0000000610_k1']),
    Reaction.from_massaction([RS30S_IF2_GTP,IF3],[RS30S_IF3_IF2_GTP], k_forward= rxn_k['re0000000615_k1']),
    Reaction.from_massaction([RS30S_IF3_IF2_GTP],[IF3,RS30S_IF2_GTP], k_forward= rxn_k['re0000000616_k1']),
    Reaction.from_massaction([mRNA,RS30S],[RS30S_mRNA], k_forward = rxn_k['re0000000619_k1']),
    Reaction.from_massaction([RS30S_mRNA],[RS30S,mRNA], k_forward= rxn_k['re0000000620_k1']),

    Reaction.from_massaction([RS30S_mRNA,IF2_GTP],[RS30S_IF2_GTP_mRNA], k_forward= rxn_k['re0000000625_k1']),
    Reaction.from_massaction([RS30S_IF2_GTP_mRNA],[IF2_GTP,RS30S_mRNA], k_forward= rxn_k['re0000000626_k1']),
    Reaction.from_massaction([mRNA,RS30S_IF2_GTP],[RS30S_IF2_GTP_mRNA], k_forward = rxn_k['re0000000631_k1']),
    Reaction.from_massaction([RS30S_IF2_GTP_mRNA],[RS30S_IF2_GTP,mRNA], k_forward= rxn_k['re0000000632_k1']),
    Reaction.from_massaction([RS30S_mRNA,IF3],[RS30S_IF3_mRNA], k_forward= rxn_k['re0000000635_k1']),
    Reaction.from_massaction([RS30S_IF3_mRNA],[IF3,RS30S_mRNA], k_forward= rxn_k['re0000000636_k1']),
    Reaction.from_massaction([RS30S_IF2_GTP_mRNA,IF3],[RS30S_IF3_IF2_GTP_mRNA], k_forward= rxn_k['re0000000639_k1']),
    Reaction.from_massaction([RS30S_IF3_IF2_GTP_mRNA],[IF3,RS30S_IF2_GTP_mRNA], k_forward= rxn_k['re0000000640_k1']),
    Reaction.from_massaction([RS30S_IF1,IF2_GTP],[RS30S_IF1_IF2_GTP], k_forward = rxn_k['re0000000643_k1']),
    Reaction.from_massaction([RS30S_IF1_IF2_GTP],[IF2_GTP,RS30S_IF1], k_forward = rxn_k['re0000000644_k1']),

    Reaction.from_massaction([IF1,RS30S_IF2_GTP],[RS30S_IF1_IF2_GTP], k_forward = rxn_k['re0000000649_k1']),
    Reaction.from_massaction([RS30S_IF1_IF2_GTP],[IF1,RS30S_IF2_GTP], k_forward = rxn_k['re0000000650_k1']),
    Reaction.from_massaction([RS30S_IF1_IF2_GTP,IF3],[RS30S_IF1_IF3_IF2_GTP], k_forward = rxn_k['re0000000653_k1']),
    Reaction.from_massaction([RS30S_IF1_IF3_IF2_GTP],[IF3,RS30S_IF1_IF2_GTP], k_forward = rxn_k['re0000000654_k1']),
    Reaction.from_massaction([RS30S_IF1_mRNA,IF2_GTP],[RS30S_IF1_IF2_GTP_mRNA], k_forward= rxn_k['re0000000657_k1']),
    Reaction.from_massaction([RS30S_IF1_IF2_GTP_mRNA],[RS30S_IF1_mRNA,IF2_GTP], k_forward = rxn_k['re0000000658_k1']),
    Reaction.from_massaction([RS30S_IF1_mRNA,IF3],[RS30S_IF1_IF3_mRNA], k_forward= rxn_k['re0000000667_k1']),
    Reaction.from_massaction([RS30S_IF1_IF3_mRNA],[IF3,RS30S_IF1_mRNA], k_forward = rxn_k['re0000000668_k1']),
    Reaction.from_massaction([RS30S_IF1_IF2_GTP_mRNA,IF3],[RS30S_IF1_IF3_IF2_GTP_mRNA], k_forward = rxn_k['re0000000671_k1']),
    Reaction.from_massaction([RS30S_IF1_IF3_IF2_GTP_mRNA],[IF3,RS30S_IF1_IF2_GTP_mRNA], k_forward = rxn_k['re0000000672_k1']),

    Reaction.from_massaction([RS30S_IF1,mRNA],[RS30S_IF1_mRNA], k_forward = rxn_k['re0000000675_k1']),
    Reaction.from_massaction([RS30S_IF1_mRNA],[RS30S_IF1,mRNA], k_forward= rxn_k['re0000000676_k1']),
    Reaction.from_massaction([RS30S_mRNA,IF1],[RS30S_IF1_mRNA], k_forward= rxn_k['re0000000677_k1']),
    Reaction.from_massaction([RS30S_IF1_mRNA],[RS30S_mRNA,IF1], k_forward= rxn_k['re0000000678_k1']),
    Reaction.from_massaction([RS30S_IF2_GTP_mRNA,IF1],[RS30S_IF1_IF2_GTP_mRNA], k_forward= rxn_k['re0000000679_k1']),
    Reaction.from_massaction([RS30S_IF1_IF2_GTP_mRNA],[RS30S_IF2_GTP_mRNA,IF1], k_forward = rxn_k['re0000000680_k1']),
    Reaction.from_massaction([RS30S_IF1_IF2_GTP,mRNA],[RS30S_IF1_IF2_GTP_mRNA], k_forward = rxn_k['re0000000681_k1']),
    Reaction.from_massaction([RS30S_IF1_IF2_GTP_mRNA],[RS30S_IF1_IF2_GTP,mRNA], k_forward = rxn_k['re0000000682_k1']),

    Reaction.from_massaction([RF3_GDP],[GDP,RF3], k_forward = rxn_k['re0000000825_k1']),
    Reaction.from_massaction([RF3,GDP],[RF3_GDP], k_forward = rxn_k['re0000000826_k1']),
    Reaction.from_massaction([RF3,GTP],[RF3_GTP], k_forward = rxn_k['re0000000827_k1']),
    Reaction.from_massaction([RF3_GTP],[RF3,GTP], k_forward = rxn_k['re0000000828_k1']),

    Reaction.from_massaction([RS50S_RRF],[RS50S,RRF], k_forward= rxn_k['re0000000918_k1']),
    Reaction.from_massaction([RS50S_RRF_EFG_GDP],[RS50S_RRF,EFG_GDP], k_forward= rxn_k['re0000000922_k1']),
    Reaction.from_massaction([RS50S_RRF_EFG_GDP],[RS50S_EFG_GDP,RRF], k_forward= rxn_k['re0000000923_k1']),
    ]
```

##### Met Species and Reaction¶

In [17]:

```
#Create al the fMet and needed MetRS Species
fMettRNAfMetCAU = Species('fMettRNAfMetCAU')
fMettRNAfMetCAU_degraded = Species('fMettRNAfMetCAU_degraded')
tRNAfMetCAU = Species('tRNAfMetCAU')
tRNAfMetCAU_degraded = Species('tRNAfMetCAU_degraded')
EFTu_GTP_MettRNAfMetCAU = Species('EFTu_GTP_MettRNAfMetCAU')

Met = Species('Met')
MetAMP = Species('MetAMP')
MetRS = Species('MetRS')

MetRS_AMP = Species('MetRS_AMP')
MetRS_AMP_MettRNAfMetCAU = Species('MetRS_AMP_MettRNAfMetCAU')
MetRS_ATP = Species('MetRS_ATP')
MetRS_ATP_tRNAfMetCAU = Species('MetRS_ATP_tRNAfMetCAU')
MetRS_degraded = Species('MetRS_degraded')
MetRS_Met = Species('MetRS_Met')
MetRS_Met_ATP = Species('MetRS_Met_ATP')
MetRS_Met_ATP_tRNAfMetCAU = Species('MetRS_Met_ATP_tRNAfMetCAU')
MetRS_Met_tRNAfMetCAU = Species('MetRS_Met_tRNAfMetCAU')
MetRS_MetAMP = Species('MetRS_MetAMP')
MetRS_MetAMP_PPi = Species('MetRS_MetAMP_PPi')
MetRS_MetAMP_PPi_tRNAfMetCAU = Species('MetRS_MetAMP_PPi_tRNAfMetCAU')
MetRS_MetAMP_tRNAfMetCAU = Species('MetRS_MetAMP_tRNAfMetCAU')
MetRS_MettRNAfMetCAU=Species('MetRS_MettRNAfMetCAU')
MetRS_tRNAfMetCAU = Species('MetRS_tRNAfMetCAU')

MettRNAfMetCAU = Species('MettRNAfMetCAU')
MettRNAfMetCAU_degraded = Species('MettRNAfMetCAU_degraded')
MTF_FD_MettRNAfMetCAU = Species('MTF_FD_MettRNAfMetCAU')
MTF_fMettRNAfMetCAU = Species('MTF_fMettRNAfMetCAU')
MTF_MettRNAfMetCAU = Species('MTF_MettRNAfMetCAU')
MTF_THF_fMettRNAfMetCAU = Species('MTF_THF_fMettRNAfMetCAU')

RS30S_fMettRNAfMetCAU_mRNA = Species('RS30S_fMettRNAfMetCAU_mRNA')
RS30S_IF1_fMettRNAfMetCAU_mRNA = Species('RS30S_IF1_fMettRNAfMetCAU_mRNA')
RS30S_IF1_IF2_GTP_fMettRNAfMetCAU = Species('RS30S_IF1_IF2_GTP_fMettRNAfMetCAU')
RS30S_IF1_IF2_GTP_fMettRNAfMetCAU_mRNA = Species('RS30S_IF1_IF2_GTP_fMettRNAfMetCAU_mRNA')
RS30S_IF1_IF3_fMettRNAfMetCAU_mRNA = Species('RS30S_IF1_IF3_fMettRNAfMetCAU_mRNA')
RS30S_IF1_IF3_IF2_GTP_fMettRNAfMetCAU = Species('RS30S_IF1_IF3_IF2_GTP_fMettRNAfMetCAU')
RS30S_IF1_IF3_IF2_GTP_fMettRNAfMetCAU_mRNA = Species('RS30S_IF1_IF3_IF2_GTP_fMettRNAfMetCAU_mRNA')
RS30S_IF2_GTP_fMettRNAfMetCAU = Species('RS30S_IF2_GTP_fMettRNAfMetCAU')
RS30S_IF2_GTP_fMettRNAfMetCAU_mRNA = Species('RS30S_IF2_GTP_fMettRNAfMetCAU_mRNA')
RS30S_IF3_fMettRNAfMetCAU_mRNA = Species('RS30S_IF3_fMettRNAfMetCAU_mRNA')
RS30S_IF3_IF2_GTP_fMettRNAfMetCAU = Species('RS30S_IF3_IF2_GTP_fMettRNAfMetCAU')
RS30S_IF3_IF2_GTP_fMettRNAfMetCAU_mRNA = Species('RS30S_IF3_IF2_GTP_fMettRNAfMetCAU_mRNA')

RS70S_IF1_fMettRNAfMetCAU_mRNA = Species('RS70S_IF1_fMettRNAfMetCAU_mRNA')
RS70S_IF1_IF2_GDP_fMettRNAfMetCAU_mRNA = Species('RS70S_IF1_IF2_GDP_fMettRNAfMetCAU_mRNA')
RS70S_IF1_IF3_fMettRNAfMetCAU_mRNA = Species('RS70S_IF1_IF3_fMettRNAfMetCAU_mRNA')
RS70S_IF1_IF3_IF2_GDP_fMettRNAfMetCAU_mRNA = Species('RS70S_IF1_IF3_IF2_GDP_fMettRNAfMetCAU_mRNA')
RS70S_IF1_IF3_IF2_GDP_PO4_fMettRNAfMetCAU_mRNA = Species('RS70S_IF1_IF3_IF2_GDP_PO4_fMettRNAfMetCAU_mRNA')
RS70S_IF1_IF3_IF2_GTP_fMettRNAfMetCAU_mRNA = Species('RS70S_IF1_IF3_IF2_GTP_fMettRNAfMetCAU_mRNA')
RS70S_IF2_GDP_fMettRNAfMetCAU_mRNA = Species('RS70S_IF2_GDP_fMettRNAfMetCAU_mRNA')
RS70S_IF3_fMettRNAfMetCAU_mRNA = Species('RS70S_IF3_fMettRNAfMetCAU_mRNA')
RS70S_IF3_IF2_GDP_fMettRNAfMetCAU_mRNA = Species('RS70S_IF3_IF2_GDP_fMettRNAfMetCAU_mRNA')
RS70S_IF3_IF2_GDP_PO4_fMettRNAfMetCAU_mRNA = Species('RS70S_IF3_IF2_GDP_PO4_fMettRNAfMetCAU_mRNA')
RS70S_IF3_IF2_GTP_fMettRNAfMetCAU_mRNA = Species('RS70S_IF3_IF2_GTP_fMettRNAfMetCAU_mRNA')

#######################################################################################################################

#A list of Species in relation to fMet
list_species_fmet=[
    fMettRNAfMetCAU, fMettRNAfMetCAU_degraded, Met, MetAMP, MetRS,tRNAfMetCAU, tRNAfMetCAU_degraded, EFTu_GTP_MettRNAfMetCAU,
    
    MetRS_AMP, MetRS_AMP_MettRNAfMetCAU, MetRS_ATP, MetRS_ATP_tRNAfMetCAU, MetRS_degraded,
    MetRS_Met, MetRS_Met_ATP, MetRS_Met_ATP_tRNAfMetCAU, MetRS_Met_tRNAfMetCAU, MetRS_MetAMP,
    MetRS_MetAMP_PPi, MetRS_MetAMP_PPi_tRNAfMetCAU, MetRS_MetAMP_tRNAfMetCAU, MetRS_MettRNAfMetCAU, MetRS_tRNAfMetCAU,
    MettRNAfMetCAU, MettRNAfMetCAU_degraded, MTF_FD_MettRNAfMetCAU, MTF_fMettRNAfMetCAU, MTF_MettRNAfMetCAU, MTF_THF_fMettRNAfMetCAU,
    
    RS30S_fMettRNAfMetCAU_mRNA, RS30S_IF1_fMettRNAfMetCAU_mRNA, RS30S_IF1_IF2_GTP_fMettRNAfMetCAU, 
    RS30S_IF1_IF2_GTP_fMettRNAfMetCAU_mRNA, RS30S_IF1_IF3_fMettRNAfMetCAU_mRNA, RS30S_IF1_IF3_IF2_GTP_fMettRNAfMetCAU, 
    RS30S_IF1_IF3_IF2_GTP_fMettRNAfMetCAU_mRNA, RS30S_IF2_GTP_fMettRNAfMetCAU, RS30S_IF2_GTP_fMettRNAfMetCAU_mRNA, 
    RS30S_IF3_fMettRNAfMetCAU_mRNA, RS30S_IF3_IF2_GTP_fMettRNAfMetCAU, RS30S_IF3_IF2_GTP_fMettRNAfMetCAU_mRNA,

    RS70S_IF1_fMettRNAfMetCAU_mRNA, RS70S_IF1_IF2_GDP_fMettRNAfMetCAU_mRNA, RS70S_IF1_IF3_fMettRNAfMetCAU_mRNA,
    RS70S_IF1_IF3_IF2_GDP_fMettRNAfMetCAU_mRNA, RS70S_IF1_IF3_IF2_GDP_PO4_fMettRNAfMetCAU_mRNA, RS70S_IF1_IF3_IF2_GTP_fMettRNAfMetCAU_mRNA,
    RS70S_IF2_GDP_fMettRNAfMetCAU_mRNA, RS70S_IF3_fMettRNAfMetCAU_mRNA, RS70S_IF3_IF2_GDP_fMettRNAfMetCAU_mRNA,
    RS70S_IF3_IF2_GDP_PO4_fMettRNAfMetCAU_mRNA, RS70S_IF3_IF2_GTP_fMettRNAfMetCAU_mRNA]

#######################################################################################################################

#fMet Reactions 
list_reaction_fmet=[
    Reaction.from_massaction([MetRS,Met],[MetRS_Met], k_forward= rxn_k['re0000000151_k1']),
    Reaction.from_massaction([MetRS_MetAMP_PPi],[MetRS_MetAMP,PPi,], k_forward= rxn_k['re0000000152_k1']),#Mg

    Reaction.from_massaction([MetRS_Met],[MetRS,Met], k_forward= rxn_k['re0000000156_k1']),
    Reaction.from_massaction([MetRS,ATP],[MetRS_ATP], k_forward= rxn_k['re0000000157_k1']),
    Reaction.from_massaction([MetRS_ATP],[MetRS,ATP], k_forward= rxn_k['re0000000158_k1']),

    Reaction.from_massaction([MetRS_ATP,Met],[MetRS_Met_ATP], k_forward= rxn_k['re0000000159_k1']),
    Reaction.from_massaction([MetRS_Met_ATP],[MetRS_ATP,Met], k_forward= rxn_k['re0000000160_k1']),
    Reaction.from_massaction([MetRS_Met,ATP],[MetRS_Met_ATP], k_forward= rxn_k['re0000000161_k1']),
    Reaction.from_massaction([MetRS_Met_ATP],[MetRS_Met,ATP], k_forward= rxn_k['re0000000162_k1']),

    Reaction.from_massaction([MetRS_Met_ATP],[MetRS_MetAMP_PPi], k_forward= rxn_k['re0000000165_k1']),
    Reaction.from_massaction([MetRS_MetAMP_PPi],[MetRS_Met_ATP], k_forward= rxn_k['re0000000166_k1']),

    Reaction.from_massaction([MetRS_AMP],[MetRS,AMP], k_forward= rxn_k['re0000000170_k1']),

    Reaction.from_massaction([MetRS_MetAMP],[MetRS,MetAMP], k_forward= rxn_k['re0000000172_k1']),
    Reaction.from_massaction([MetRS,MetAMP],[MetRS_MetAMP], k_forward= rxn_k['re0000000173_k1']),

    Reaction.from_massaction([MetRS_MetAMP_tRNAfMetCAU],[MetRS_AMP_MettRNAfMetCAU], k_forward= rxn_k['re0000000220_k1']),

    Reaction.from_massaction([MetRS_AMP_MettRNAfMetCAU],[MetRS_MettRNAfMetCAU,AMP], k_forward= rxn_k['re0000000222_k1']),
    
    Reaction.from_massaction([MetRS_AMP_MettRNAfMetCAU],[MettRNAfMetCAU,MetRS_AMP], k_forward= rxn_k['re0000000224_k1']),
    Reaction.from_massaction([MetRS_AMP,MettRNAfMetCAU],[MetRS_AMP_MettRNAfMetCAU], k_forward= rxn_k['re0000000225_k1']),
    Reaction.from_massaction([MetRS_MettRNAfMetCAU],[MetRS,MettRNAfMetCAU], k_forward= rxn_k['re0000000226_k1']),
    Reaction.from_massaction([MetRS,MettRNAfMetCAU],[MetRS_MettRNAfMetCAU], k_forward= rxn_k['re0000000227_k1']),

    Reaction.from_massaction([MetRS_tRNAfMetCAU,Met],[MetRS_Met_tRNAfMetCAU], k_forward= rxn_k['re0000000230_k1']),

    Reaction.from_massaction([MetRS_MetAMP_PPi_tRNAfMetCAU],[MetRS_MetAMP_tRNAfMetCAU,PPi], k_forward= rxn_k['re0000000231_k1']),
    Reaction.from_massaction([MetRS_Met_tRNAfMetCAU],[MetRS_tRNAfMetCAU,Met], k_forward= rxn_k['re0000000232_k1']),
    Reaction.from_massaction([MetRS_tRNAfMetCAU,ATP],[MetRS_ATP_tRNAfMetCAU], k_forward= rxn_k['re0000000233_k1']),
    Reaction.from_massaction([MetRS_ATP_tRNAfMetCAU],[MetRS_tRNAfMetCAU,ATP], k_forward= rxn_k['re0000000234_k1']),
    Reaction.from_massaction([MetRS_ATP_tRNAfMetCAU,Met],[MetRS_Met_ATP_tRNAfMetCAU], k_forward= rxn_k['re0000000235_k1']),
    Reaction.from_massaction([MetRS_Met_ATP_tRNAfMetCAU],[MetRS_ATP_tRNAfMetCAU,Met], k_forward= rxn_k['re0000000236_k1']),
    Reaction.from_massaction([MetRS_Met_tRNAfMetCAU,ATP],[MetRS_Met_ATP_tRNAfMetCAU], k_forward= rxn_k['re0000000237_k1']),
    Reaction.from_massaction([MetRS_Met_ATP_tRNAfMetCAU],[MetRS_Met_tRNAfMetCAU,ATP], k_forward= rxn_k['re0000000238_k1']),
    Reaction.from_massaction([MetRS_Met_ATP_tRNAfMetCAU],[MetRS_MetAMP_PPi_tRNAfMetCAU], k_forward= rxn_k['re0000000239_k1']),

    Reaction.from_massaction([MetRS,tRNAfMetCAU],[MetRS_tRNAfMetCAU], k_forward= rxn_k['re0000000241_k1']),
    Reaction.from_massaction([MetRS_Met,tRNAfMetCAU],[MetRS_Met_tRNAfMetCAU], k_forward= rxn_k['re0000000242_k1']),
    Reaction.from_massaction([MetRS_tRNAfMetCAU],[MetRS,tRNAfMetCAU], k_forward= rxn_k['re0000000243_k1']),
    Reaction.from_massaction([MetRS_Met_tRNAfMetCAU],[MetRS_Met,tRNAfMetCAU], k_forward= rxn_k['re0000000244_k1']),
    Reaction.from_massaction([MetRS_ATP,tRNAfMetCAU],[MetRS_ATP_tRNAfMetCAU], k_forward= rxn_k['re0000000245_k1']),
    Reaction.from_massaction([MetRS_ATP_tRNAfMetCAU],[MetRS_ATP,tRNAfMetCAU], k_forward= rxn_k['re0000000246_k1']),
    Reaction.from_massaction([MetRS_Met_ATP,tRNAfMetCAU],[MetRS_Met_ATP_tRNAfMetCAU], k_forward= rxn_k['re0000000247_k1']),
    Reaction.from_massaction([MetRS_Met_ATP_tRNAfMetCAU],[MetRS_Met_ATP,tRNAfMetCAU], k_forward= rxn_k['re0000000248_k1']),
    Reaction.from_massaction([MetRS_MetAMP_PPi,tRNAfMetCAU],[MetRS_MetAMP_PPi_tRNAfMetCAU], k_forward= rxn_k['re0000000249_k1']),
    Reaction.from_massaction([MetRS_MetAMP_PPi_tRNAfMetCAU],[MetRS_MetAMP_PPi,tRNAfMetCAU], k_forward= rxn_k['re0000000250_k1']),

    Reaction.from_massaction([MetRS_MetAMP,tRNAfMetCAU],[MetRS_MetAMP_tRNAfMetCAU], k_forward= rxn_k['re0000000251_k1']),
    Reaction.from_massaction([MetRS_MetAMP_tRNAfMetCAU],[MetRS_MetAMP,tRNAfMetCAU], k_forward= rxn_k['re0000000252_k1']),

    Reaction.from_massaction([EFTu_GTP,MettRNAfMetCAU],[EFTu_GTP_MettRNAfMetCAU], k_forward= rxn_k['re0000000288_k1']),
    Reaction.from_massaction([EFTu_GTP_MettRNAfMetCAU],[EFTu_GTP,MettRNAfMetCAU], k_forward= rxn_k['re0000000289_k1']),

    Reaction.from_massaction([MTF,MettRNAfMetCAU],[MTF_MettRNAfMetCAU], k_forward= rxn_k['re0000000420_k1']),
    Reaction.from_massaction([MTF_MettRNAfMetCAU],[MTF,MettRNAfMetCAU], k_forward= rxn_k['re0000000421_k1']),
    Reaction.from_massaction([MTF_FD,MettRNAfMetCAU],[MTF_FD_MettRNAfMetCAU], k_forward= rxn_k['re0000000422_k1']),
    Reaction.from_massaction([MTF_FD_MettRNAfMetCAU],[MettRNAfMetCAU,MTF_FD], k_forward= rxn_k['re0000000423_k1']),
    Reaction.from_massaction([MTF_MettRNAfMetCAU,FD],[MTF_FD_MettRNAfMetCAU], k_forward= rxn_k['re0000000424_k1']),

    Reaction.from_massaction([MTF_FD_MettRNAfMetCAU],[MTF_MettRNAfMetCAU,FD], k_forward= rxn_k['re0000000425_k1']),
    Reaction.from_massaction([MTF_FD_MettRNAfMetCAU],[MTF_THF_fMettRNAfMetCAU], k_forward= rxn_k['re0000000426_k1']),

    Reaction.from_massaction([MTF_THF_fMettRNAfMetCAU],[MTF_THF,fMettRNAfMetCAU], k_forward= rxn_k['re0000000428_k1']),

    Reaction.from_massaction([MTF_THF_fMettRNAfMetCAU],[MTF_fMettRNAfMetCAU,THF], k_forward= rxn_k['re0000000430_k1']),

    Reaction.from_massaction([MTF_fMettRNAfMetCAU],[MTF,fMettRNAfMetCAU], k_forward= rxn_k['re0000000432_k1']),

    Reaction.from_massaction([IF2_GTP,fMettRNAfMetCAU],[IF2_GTP_fMettRNAfMetCAU], k_forward= rxn_k['re0000000449_k1']),
    Reaction.from_massaction([IF2_GTP_fMettRNAfMetCAU],[IF2_GTP,fMettRNAfMetCAU], k_forward= rxn_k['re0000000450_k1']),

    Reaction.from_massaction([RS30S_IF3,IF2_GTP_fMettRNAfMetCAU],[RS30S_IF3_IF2_GTP_fMettRNAfMetCAU], k_forward= rxn_k['re0000000465_k1']),
    Reaction.from_massaction([RS30S_IF3_IF2_GTP_fMettRNAfMetCAU],[RS30S_IF3,IF2_GTP_fMettRNAfMetCAU], k_forward= rxn_k['re0000000466_k1']),
    Reaction.from_massaction([RS30S_IF3_IF2_GTP,fMettRNAfMetCAU],[RS30S_IF3_IF2_GTP_fMettRNAfMetCAU], k_forward= rxn_k['re0000000467_k1']),

    Reaction.from_massaction([RS30S_IF3_IF2_GTP_fMettRNAfMetCAU],[RS30S_IF3_IF2_GTP,fMettRNAfMetCAU], k_forward= rxn_k['re0000000468_k1']),
    Reaction.from_massaction([RS30S_IF3_mRNA,fMettRNAfMetCAU],[RS30S_IF3_fMettRNAfMetCAU_mRNA], k_forward= rxn_k['re0000000473_k1']),
    Reaction.from_massaction([RS30S_IF3_fMettRNAfMetCAU_mRNA],[RS30S_IF3_mRNA,fMettRNAfMetCAU], k_forward= rxn_k['re0000000474_k1']),
    Reaction.from_massaction([RS30S_IF3_mRNA,IF2_GTP_fMettRNAfMetCAU],[RS30S_IF3_IF2_GTP_fMettRNAfMetCAU_mRNA], k_forward= rxn_k['re0000000475_k1']),
    Reaction.from_massaction([RS30S_IF3_IF2_GTP_fMettRNAfMetCAU_mRNA],[RS30S_IF3_mRNA,IF2_GTP_fMettRNAfMetCAU], k_forward= rxn_k['re0000000476_k1']),
    Reaction.from_massaction([RS30S_IF3_fMettRNAfMetCAU_mRNA,IF2_GTP],[RS30S_IF3_IF2_GTP_fMettRNAfMetCAU_mRNA], k_forward= rxn_k['re0000000477_k1']),
    Reaction.from_massaction([RS30S_IF3_IF2_GTP_fMettRNAfMetCAU_mRNA],[RS30S_IF3_fMettRNAfMetCAU_mRNA,IF2_GTP], k_forward= rxn_k['re0000000478_k1']),
    Reaction.from_massaction([RS30S_IF3_IF2_GTP_mRNA,fMettRNAfMetCAU],[RS30S_IF3_IF2_GTP_fMettRNAfMetCAU_mRNA], k_forward= rxn_k['re0000000479_k1']),
    Reaction.from_massaction([RS30S_IF3_IF2_GTP_fMettRNAfMetCAU_mRNA],[RS30S_IF3_IF2_GTP_mRNA,fMettRNAfMetCAU], k_forward= rxn_k['re0000000480_k1']),
    Reaction.from_massaction([RS30S_IF3_IF2_GTP_fMettRNAfMetCAU,mRNA],[RS30S_IF3_IF2_GTP_fMettRNAfMetCAU_mRNA], k_forward= rxn_k['re0000000483_k1']),

    Reaction.from_massaction([RS30S_IF3_IF2_GTP_fMettRNAfMetCAU_mRNA],[RS30S_IF3_IF2_GTP_fMettRNAfMetCAU,mRNA], k_forward= rxn_k['re0000000484_k1']),
    Reaction.from_massaction([RS30S_IF3_IF2_GTP_fMettRNAfMetCAU_mRNA,RS50S],[RS70S_IF3_IF2_GTP_fMettRNAfMetCAU_mRNA], k_forward= rxn_k['re0000000485_k1']),
    Reaction.from_massaction([RS70S_IF3_IF2_GTP_fMettRNAfMetCAU_mRNA],[RS30S_IF3_IF2_GTP_fMettRNAfMetCAU_mRNA,RS50S], k_forward= rxn_k['re0000000486_k1']),
    Reaction.from_massaction([RS30S_IF1_IF3,IF2_GTP_fMettRNAfMetCAU],[RS30S_IF1_IF3_IF2_GTP_fMettRNAfMetCAU], k_forward= rxn_k['re0000000497_k1']),
    Reaction.from_massaction([RS30S_IF1_IF3_IF2_GTP_fMettRNAfMetCAU],[RS30S_IF1_IF3,IF2_GTP_fMettRNAfMetCAU], k_forward= rxn_k['re0000000498_k1']),
    Reaction.from_massaction([RS30S_IF1_IF3_IF2_GTP,fMettRNAfMetCAU],[RS30S_IF1_IF3_IF2_GTP_fMettRNAfMetCAU], k_forward= rxn_k['re0000000499_k1']),
    Reaction.from_massaction([RS30S_IF1_IF3_IF2_GTP_fMettRNAfMetCAU],[RS30S_IF1_IF3_IF2_GTP,fMettRNAfMetCAU], k_forward= rxn_k['re0000000500_k1']),
    Reaction.from_massaction([RS30S_IF3_IF2_GTP_fMettRNAfMetCAU,IF1],[RS30S_IF1_IF3_IF2_GTP_fMettRNAfMetCAU], k_forward= rxn_k['re0000000511_k1']),
    Reaction.from_massaction([RS30S_IF1_IF3_IF2_GTP_fMettRNAfMetCAU],[RS30S_IF3_IF2_GTP_fMettRNAfMetCAU,IF1], k_forward= rxn_k['re0000000512_k1']),
    Reaction.from_massaction([RS30S_IF1_IF3_mRNA,fMettRNAfMetCAU],[RS30S_IF1_IF3_fMettRNAfMetCAU_mRNA], k_forward= rxn_k['re0000000517_k1']),

    Reaction.from_massaction([RS30S_IF1_IF3_fMettRNAfMetCAU_mRNA],[RS30S_IF1_IF3_mRNA,fMettRNAfMetCAU], k_forward= rxn_k['re0000000518_k1']),
    Reaction.from_massaction([RS30S_IF1_IF3_mRNA,IF2_GTP_fMettRNAfMetCAU],[RS30S_IF1_IF3_IF2_GTP_fMettRNAfMetCAU_mRNA], k_forward= rxn_k['re0000000519_k1']),
    Reaction.from_massaction([RS30S_IF1_IF3_IF2_GTP_fMettRNAfMetCAU_mRNA],[RS30S_IF1_IF3_mRNA,IF2_GTP_fMettRNAfMetCAU], k_forward= rxn_k['re0000000520_k1']),
    Reaction.from_massaction([RS30S_IF1_IF3_fMettRNAfMetCAU_mRNA,IF2_GTP],[RS30S_IF1_IF3_IF2_GTP_fMettRNAfMetCAU_mRNA], k_forward= rxn_k['re0000000521_k1']),
    Reaction.from_massaction([RS30S_IF1_IF3_IF2_GTP_fMettRNAfMetCAU_mRNA],[RS30S_IF1_IF3_fMettRNAfMetCAU_mRNA,IF2_GTP], k_forward= rxn_k['re0000000522_k1']),
    Reaction.from_massaction([RS30S_IF1_IF3_IF2_GTP_mRNA,fMettRNAfMetCAU],[RS30S_IF1_IF3_IF2_GTP_fMettRNAfMetCAU_mRNA], k_forward= rxn_k['re0000000523_k1']),
    Reaction.from_massaction([RS30S_IF1_IF3_IF2_GTP_fMettRNAfMetCAU_mRNA],[RS30S_IF1_IF3_IF2_GTP_mRNA,fMettRNAfMetCAU], k_forward= rxn_k['re0000000524_k1']),
    Reaction.from_massaction([RS30S_IF1_IF3_IF2_GTP_fMettRNAfMetCAU,mRNA],[RS30S_IF1_IF3_IF2_GTP_fMettRNAfMetCAU_mRNA], k_forward= rxn_k['re0000000527_k1']),
    Reaction.from_massaction([RS30S_IF1_IF3_IF2_GTP_fMettRNAfMetCAU_mRNA],[RS30S_IF1_IF3_IF2_GTP_fMettRNAfMetCAU,mRNA], k_forward= rxn_k['re0000000528_k1']),

    Reaction.from_massaction([RS30S_IF1_IF3_IF2_GTP_fMettRNAfMetCAU_mRNA,RS50S],[RS70S_IF1_IF3_IF2_GTP_fMettRNAfMetCAU_mRNA], k_forward= rxn_k['re0000000529_k1']),
    Reaction.from_massaction([RS70S_IF1_IF3_IF2_GTP_fMettRNAfMetCAU_mRNA],[RS30S_IF1_IF3_IF2_GTP_fMettRNAfMetCAU_mRNA,RS50S], k_forward= rxn_k['re0000000530_k1']),
    Reaction.from_massaction([RS30S_IF3_fMettRNAfMetCAU_mRNA,IF1],[RS30S_IF1_IF3_fMettRNAfMetCAU_mRNA], k_forward= rxn_k['re0000000533_k1']),
    Reaction.from_massaction([RS30S_IF1_IF3_fMettRNAfMetCAU_mRNA],[RS30S_IF3_fMettRNAfMetCAU_mRNA,IF1], k_forward= rxn_k['re0000000534_k1']),
    Reaction.from_massaction([RS30S_IF3_IF2_GTP_fMettRNAfMetCAU_mRNA,IF1],[RS30S_IF1_IF3_IF2_GTP_fMettRNAfMetCAU_mRNA], k_forward= rxn_k['re0000000537_k1']),
    Reaction.from_massaction([RS30S_IF1_IF3_IF2_GTP_fMettRNAfMetCAU_mRNA],[RS30S_IF3_IF2_GTP_fMettRNAfMetCAU_mRNA,IF1], k_forward= rxn_k['re0000000538_k1']),
    Reaction.from_massaction([RS70S_IF3_IF2_GTP_fMettRNAfMetCAU_mRNA,IF1],[RS70S_IF1_IF3_IF2_GTP_fMettRNAfMetCAU_mRNA], k_forward= rxn_k['re0000000539_k1']),
    Reaction.from_massaction([RS70S_IF1_IF3_IF2_GTP_fMettRNAfMetCAU_mRNA],[RS70S_IF3_IF2_GTP_fMettRNAfMetCAU_mRNA,IF1], k_forward= rxn_k['re0000000540_k1']),

    Reaction.from_massaction([RS30S,IF2_GTP_fMettRNAfMetCAU],[RS30S_IF2_GTP_fMettRNAfMetCAU], k_forward= rxn_k['re0000000611_k1']),
    Reaction.from_massaction([RS30S_IF2_GTP_fMettRNAfMetCAU],[IF2_GTP_fMettRNAfMetCAU,RS30S], k_forward= rxn_k['re0000000612_k1']),
    Reaction.from_massaction([RS30S_IF2_GTP,fMettRNAfMetCAU],[RS30S_IF2_GTP_fMettRNAfMetCAU], k_forward= rxn_k['re0000000613_k1']),
    Reaction.from_massaction([RS30S_IF2_GTP_fMettRNAfMetCAU],[fMettRNAfMetCAU,RS30S_IF2_GTP], k_forward= rxn_k['re0000000614_k1']),

    Reaction.from_massaction([RS30S_IF2_GTP_fMettRNAfMetCAU,IF3],[RS30S_IF3_IF2_GTP_fMettRNAfMetCAU], k_forward= rxn_k['re0000000617_k1']),
    Reaction.from_massaction([RS30S_IF3_IF2_GTP_fMettRNAfMetCAU],[IF3,RS30S_IF2_GTP_fMettRNAfMetCAU], k_forward= rxn_k['re0000000618_k1']),
    Reaction.from_massaction([RS30S_mRNA,fMettRNAfMetCAU],[RS30S_fMettRNAfMetCAU_mRNA], k_forward= rxn_k['re0000000621_k1']),
    Reaction.from_massaction([RS30S_fMettRNAfMetCAU_mRNA],[fMettRNAfMetCAU,RS30S_mRNA], k_forward= rxn_k['re0000000622_k1']),
    Reaction.from_massaction([RS30S_mRNA,IF2_GTP_fMettRNAfMetCAU],[RS30S_IF2_GTP_fMettRNAfMetCAU_mRNA], k_forward= rxn_k['re0000000623_k1']),
    Reaction.from_massaction([RS30S_IF2_GTP_fMettRNAfMetCAU_mRNA],[IF2_GTP_fMettRNAfMetCAU,RS30S_mRNA], k_forward= rxn_k['re0000000624_k1']),
    Reaction.from_massaction([RS30S_fMettRNAfMetCAU_mRNA,IF2_GTP],[RS30S_IF2_GTP_fMettRNAfMetCAU_mRNA], k_forward= rxn_k['re0000000627_k1']),
    Reaction.from_massaction([RS30S_IF2_GTP_fMettRNAfMetCAU_mRNA],[IF2_GTP,RS30S_fMettRNAfMetCAU_mRNA], k_forward= rxn_k['re0000000628_k1']),
    Reaction.from_massaction([RS30S_IF2_GTP_mRNA,fMettRNAfMetCAU],[RS30S_IF2_GTP_fMettRNAfMetCAU_mRNA], k_forward= rxn_k['re0000000629_k1']),
    Reaction.from_massaction([RS30S_IF2_GTP_fMettRNAfMetCAU_mRNA],[fMettRNAfMetCAU,RS30S_IF2_GTP_mRNA], k_forward= rxn_k['re0000000630_k1']),

    Reaction.from_massaction([mRNA,RS30S_IF2_GTP_fMettRNAfMetCAU],[RS30S_IF2_GTP_fMettRNAfMetCAU_mRNA], k_forward= rxn_k['re0000000633_k1']),
    Reaction.from_massaction([RS30S_IF2_GTP_fMettRNAfMetCAU_mRNA],[RS30S_IF2_GTP_fMettRNAfMetCAU,mRNA], k_forward= rxn_k['re0000000634_k1']),
    Reaction.from_massaction([RS30S_fMettRNAfMetCAU_mRNA,IF3],[RS30S_IF3_fMettRNAfMetCAU_mRNA], k_forward= rxn_k['re0000000637_k1']),
    Reaction.from_massaction([RS30S_IF3_fMettRNAfMetCAU_mRNA],[IF3,RS30S_fMettRNAfMetCAU_mRNA], k_forward= rxn_k['re0000000638_k1']),
    Reaction.from_massaction([RS30S_IF2_GTP_fMettRNAfMetCAU_mRNA,IF3],[RS30S_IF3_IF2_GTP_fMettRNAfMetCAU_mRNA], k_forward= rxn_k['re0000000641_k1']),
    Reaction.from_massaction([RS30S_IF3_IF2_GTP_fMettRNAfMetCAU_mRNA],[IF3,RS30S_IF2_GTP_fMettRNAfMetCAU_mRNA], k_forward= rxn_k['re0000000642_k1']),
    Reaction.from_massaction([RS30S_IF1,IF2_GTP_fMettRNAfMetCAU],[RS30S_IF1_IF2_GTP_fMettRNAfMetCAU], k_forward= rxn_k['re0000000645_k1']),
    Reaction.from_massaction([RS30S_IF1_IF2_GTP_fMettRNAfMetCAU],[IF2_GTP_fMettRNAfMetCAU,RS30S_IF1], k_forward= rxn_k['re0000000646_k1']),
    Reaction.from_massaction([RS30S_IF1_IF2_GTP,fMettRNAfMetCAU],[RS30S_IF1_IF2_GTP_fMettRNAfMetCAU], k_forward= rxn_k['re0000000647_k1']),
    Reaction.from_massaction([RS30S_IF1_IF2_GTP_fMettRNAfMetCAU],[fMettRNAfMetCAU,RS30S_IF1_IF2_GTP], k_forward= rxn_k['re0000000648_k1']),

    Reaction.from_massaction([RS30S_IF2_GTP_fMettRNAfMetCAU,IF1],[RS30S_IF1_IF2_GTP_fMettRNAfMetCAU], k_forward= rxn_k['re0000000651_k1']),
    Reaction.from_massaction([RS30S_IF1_IF2_GTP_fMettRNAfMetCAU],[IF1,RS30S_IF2_GTP_fMettRNAfMetCAU], k_forward= rxn_k['re0000000652_k1']),
    Reaction.from_massaction([RS30S_IF1_IF2_GTP_fMettRNAfMetCAU,IF3],[RS30S_IF1_IF3_IF2_GTP_fMettRNAfMetCAU], k_forward= rxn_k['re0000000655_k1']),
    Reaction.from_massaction([RS30S_IF1_IF3_IF2_GTP_fMettRNAfMetCAU],[IF3,RS30S_IF1_IF2_GTP_fMettRNAfMetCAU], k_forward= rxn_k['re0000000656_k1']),
    Reaction.from_massaction([RS30S_IF1_mRNA,fMettRNAfMetCAU],[RS30S_IF1_fMettRNAfMetCAU_mRNA], k_forward= rxn_k['re0000000659_k1']),
    Reaction.from_massaction([RS30S_IF1_fMettRNAfMetCAU_mRNA],[fMettRNAfMetCAU,RS30S_IF1_mRNA], k_forward= rxn_k['re0000000660_k1']),
    Reaction.from_massaction([RS30S_IF1_IF2_GTP_mRNA,fMettRNAfMetCAU],[RS30S_IF1_IF2_GTP_fMettRNAfMetCAU_mRNA], k_forward= rxn_k['re0000000661_k1']),
    Reaction.from_massaction([RS30S_IF1_IF2_GTP_fMettRNAfMetCAU_mRNA],[fMettRNAfMetCAU,RS30S_IF1_IF2_GTP_mRNA], k_forward= rxn_k['re0000000662_k1']),
    Reaction.from_massaction([RS30S_IF1_fMettRNAfMetCAU_mRNA,IF2_GTP],[RS30S_IF1_IF2_GTP_fMettRNAfMetCAU_mRNA], k_forward= rxn_k['re0000000663_k1']),
    Reaction.from_massaction([RS30S_IF1_IF2_GTP_fMettRNAfMetCAU_mRNA],[IF2_GTP,RS30S_IF1_fMettRNAfMetCAU_mRNA], k_forward= rxn_k['re0000000664_k1']),

    Reaction.from_massaction([RS30S_IF1_mRNA,IF2_GTP_fMettRNAfMetCAU],[RS30S_IF1_IF2_GTP_fMettRNAfMetCAU_mRNA], k_forward= rxn_k['re0000000665_k1']),
    Reaction.from_massaction([RS30S_IF1_IF2_GTP_fMettRNAfMetCAU_mRNA],[IF2_GTP_fMettRNAfMetCAU,RS30S_IF1_mRNA], k_forward= rxn_k['re0000000666_k1']),
    Reaction.from_massaction([RS30S_IF1_fMettRNAfMetCAU_mRNA,IF3],[RS30S_IF1_IF3_fMettRNAfMetCAU_mRNA], k_forward= rxn_k['re0000000669_k1']),
    Reaction.from_massaction([RS30S_IF1_IF3_fMettRNAfMetCAU_mRNA],[IF3,RS30S_IF1_fMettRNAfMetCAU_mRNA], k_forward= rxn_k['re0000000670_k1']),
    Reaction.from_massaction([RS30S_IF1_IF2_GTP_fMettRNAfMetCAU_mRNA,IF3],[RS30S_IF1_IF3_IF2_GTP_fMettRNAfMetCAU_mRNA], k_forward= rxn_k['re0000000673_k1']),
    Reaction.from_massaction([RS30S_IF1_IF3_IF2_GTP_fMettRNAfMetCAU_mRNA],[IF3,RS30S_IF1_IF2_GTP_fMettRNAfMetCAU_mRNA], k_forward= rxn_k['re0000000674_k1']),
    Reaction.from_massaction([RS30S_fMettRNAfMetCAU_mRNA,IF1],[RS30S_IF1_fMettRNAfMetCAU_mRNA], k_forward= rxn_k['re0000000683_k1']),
    Reaction.from_massaction([RS30S_IF1_fMettRNAfMetCAU_mRNA],[RS30S_fMettRNAfMetCAU_mRNA,IF1], k_forward= rxn_k['re0000000684_k1']),
    Reaction.from_massaction([IF1,RS30S_IF2_GTP_fMettRNAfMetCAU_mRNA],[RS30S_IF1_IF2_GTP_fMettRNAfMetCAU_mRNA], k_forward= rxn_k['re0000000685_k1']),
    Reaction.from_massaction([RS30S_IF1_IF2_GTP_fMettRNAfMetCAU_mRNA],[RS30S_IF2_GTP_fMettRNAfMetCAU_mRNA,IF1], k_forward= rxn_k['re0000000686_k1']),

    Reaction.from_massaction([RS30S_IF1_IF2_GTP_fMettRNAfMetCAU,mRNA],[RS30S_IF1_IF2_GTP_fMettRNAfMetCAU_mRNA], k_forward= rxn_k['re0000000687_k1']),
    Reaction.from_massaction([RS30S_IF1_IF2_GTP_fMettRNAfMetCAU_mRNA],[RS30S_IF1_IF2_GTP_fMettRNAfMetCAU,mRNA], k_forward= rxn_k['re0000000688_k1']),

    Reaction.from_massaction([RS70S_IF3_IF2_GTP_fMettRNAfMetCAU_mRNA],[RS70S_IF3_IF2_GDP_PO4_fMettRNAfMetCAU_mRNA], k_forward= rxn_k['re0000000715_k1']),
    Reaction.from_massaction([RS70S_IF3_IF2_GDP_PO4_fMettRNAfMetCAU_mRNA],[RS70S_IF3_IF2_GTP_fMettRNAfMetCAU_mRNA], k_forward= rxn_k['re0000000716_k1']),
    Reaction.from_massaction([RS70S_IF1_IF3_IF2_GTP_fMettRNAfMetCAU_mRNA],[RS70S_IF1_IF3_IF2_GDP_PO4_fMettRNAfMetCAU_mRNA], k_forward= rxn_k['re0000000717_k1']),
    Reaction.from_massaction([RS70S_IF1_IF3_IF2_GDP_PO4_fMettRNAfMetCAU_mRNA],[RS70S_IF1_IF3_IF2_GTP_fMettRNAfMetCAU_mRNA], k_forward= rxn_k['re0000000718_k1']),
    Reaction.from_massaction([RS70S_IF3_IF2_GDP_PO4_fMettRNAfMetCAU_mRNA,IF1],[RS70S_IF1_IF3_IF2_GDP_PO4_fMettRNAfMetCAU_mRNA], k_forward= rxn_k['re0000000719_k1']),

    Reaction.from_massaction([RS70S_IF1_IF3_IF2_GDP_PO4_fMettRNAfMetCAU_mRNA],[RS70S_IF3_IF2_GDP_PO4_fMettRNAfMetCAU_mRNA,IF1], k_forward= rxn_k['re0000000720_k1']),
    Reaction.from_massaction([RS70S_IF3_IF2_GDP_PO4_fMettRNAfMetCAU_mRNA],[RS70S_IF3_IF2_GDP_fMettRNAfMetCAU_mRNA,PO4,], k_forward= rxn_k['re0000000721_k1']), 
    Reaction.from_massaction([RS70S_IF1_IF3_IF2_GDP_PO4_fMettRNAfMetCAU_mRNA],[RS70S_IF1_IF3_IF2_GDP_fMettRNAfMetCAU_mRNA,PO4,], k_forward= rxn_k['re0000000722_k1']),
    Reaction.from_massaction([RS70S_IF3_IF2_GDP_fMettRNAfMetCAU_mRNA,IF1],[RS70S_IF1_IF3_IF2_GDP_fMettRNAfMetCAU_mRNA], k_forward= rxn_k['re0000000723_k1']),
    Reaction.from_massaction([RS70S_IF1_IF3_IF2_GDP_fMettRNAfMetCAU_mRNA],[RS70S_IF3_IF2_GDP_fMettRNAfMetCAU_mRNA,IF1], k_forward= rxn_k['re0000000724_k1']),
    Reaction.from_massaction([RS70S_IF3_IF2_GDP_fMettRNAfMetCAU_mRNA],[RS70S_IF3_fMettRNAfMetCAU_mRNA,IF2_GDP], k_forward= rxn_k['re0000000725_k1']),

    Reaction.from_massaction([RS70S_IF1_IF3_IF2_GDP_fMettRNAfMetCAU_mRNA],[RS70S_IF1_IF2_GDP_fMettRNAfMetCAU_mRNA,IF3], k_forward= rxn_k['re0000000747_k1']),

    Reaction.from_massaction([RS70S_IF1_IF3_IF2_GDP_fMettRNAfMetCAU_mRNA],[RS70S_IF1_IF3_fMettRNAfMetCAU_mRNA,IF2_GDP], k_forward= rxn_k['re0000000749_k1']),
    Reaction.from_massaction([RS70S_IF1_IF3_fMettRNAfMetCAU_mRNA,IF2_GDP],[RS70S_IF1_IF3_IF2_GDP_fMettRNAfMetCAU_mRNA], k_forward= rxn_k['re0000000750_k1']),

    Reaction.from_massaction([RS70S_IF3_fMettRNAfMetCAU_mRNA,IF2_GDP],[RS70S_IF3_IF2_GDP_fMettRNAfMetCAU_mRNA], k_forward= rxn_k['re0000000751_k1']),
    Reaction.from_massaction([RS70S_IF1_IF3_fMettRNAfMetCAU_mRNA],[RS70S_IF3_fMettRNAfMetCAU_mRNA,IF1], k_forward= rxn_k['re0000000753_k1']),

    Reaction.from_massaction([RS70S_IF3_IF2_GDP_fMettRNAfMetCAU_mRNA],[RS70S_IF2_GDP_fMettRNAfMetCAU_mRNA,IF3], k_forward= rxn_k['re0000000755_k1']),

    Reaction.from_massaction([RS70S_IF1_IF2_GDP_fMettRNAfMetCAU_mRNA],[RS70S_IF2_GDP_fMettRNAfMetCAU_mRNA,IF1], k_forward= rxn_k['re0000000757_k1']),

    Reaction.from_massaction([RS70S_IF1_IF2_GDP_fMettRNAfMetCAU_mRNA],[RS70S_IF1_fMettRNAfMetCAU_mRNA,IF2_GDP], k_forward= rxn_k['re0000000759_k1']),
    Reaction.from_massaction([RS70S_IF1_fMettRNAfMetCAU_mRNA,IF2_GDP],[RS70S_IF1_IF2_GDP_fMettRNAfMetCAU_mRNA], k_forward= rxn_k['re0000000760_k1']),
    Reaction.from_massaction([RS70S_IF1_IF3_fMettRNAfMetCAU_mRNA],[RS70S_IF1_fMettRNAfMetCAU_mRNA,IF3], k_forward= rxn_k['re0000000761_k1']),
    ]
```

#### Elongations¶

Individually adds each amino acid of the growing peptide strand.

In [19]:

```
protein_lenth = '0000'
list_of_reaction_pept3 = []
list_species_el3p3=[]
for L in [l for l in range(len(protein)) if l !=0]:
    #Identification of the previous amino acid
    oA=aa
    oX=xyz
    
    #Identification of the second amino acid
    aa=protein[L]
    xyz= list_condon[list_AA.index(aa)]
    Spept= protein_lenth[:-len(str(L))]+str(L)
    Gpept= protein_lenth[:-len(str(L+1))]+str((L+1))
    #######################################################################################################################

    #The initial step of reading the 2nd amino acid
    if L==1:
        elRS70SAxyz0002_fMet = Species('elRS70SA'+xyz+'0002_fMet')
        elRS70SAxyz0002_fMet_EFTu_GDP = Species('elRS70SA'+xyz+'0002_fMet_EFTu_GDP')
        elRS70SAxyz0002_fMettRNAfMetCAU = Species('elRS70SA'+xyz+'0002_fMettRNAfMetCAU')

        list_species_elfmet= [elRS70SAxyz0002_fMet, elRS70SAxyz0002_fMet_EFTu_GDP, elRS70SAxyz0002_fMettRNAfMetCAU]
        
        list_reactions_elfmet= [
            Reaction.from_massaction([elRS70SAxyz0002_fMettRNAfMetCAU],[elRS70SAxyz0002_fMet,tRNAfMetCAU], k_forward=rxn_k['re0000000001_k1']),

            Reaction.from_massaction([elRS70SAxyz0002_fMet_EFTu_GDP],[elRS70SAxyz0002_fMet,EFTu_GDP], k_forward= rxn_k['re0000000028_k1']),

            Reaction.from_massaction([RS70S_IF2_GDP_fMettRNAfMetCAU_mRNA],[elRS70SAxyz0002_fMettRNAfMetCAU,IF2_GDP], k_forward= rxn_k['re0000000726_k1']),
            Reaction.from_massaction([elRS70SAxyz0002_fMettRNAfMetCAU,IF2_GDP],[RS70S_IF2_GDP_fMettRNAfMetCAU_mRNA], k_forward= rxn_k['re0000000752_k1']),
            Reaction.from_massaction([RS70S_IF3_fMettRNAfMetCAU_mRNA],[elRS70SAxyz0002_fMettRNAfMetCAU,IF3], k_forward= rxn_k['re0000000763_k1']),

            Reaction.from_massaction([RS70S_IF1_fMettRNAfMetCAU_mRNA],[elRS70SAxyz0002_fMettRNAfMetCAU,IF1], k_forward= rxn_k['re0000000765_k1']),
            ]
    #######################################################################################################################

        # Incorporation of the 2nd amino acid
        ## A2_fmet
        elRS70SAxyz0002_fMet_EFTu_GDP_aatRNAaaxyz = Species('elRS70SA'+xyz+'0002_fMet_EFTu_GDP_'+aa+'tRNA'+aa+xyz)
        elRS70SAxyz0002_fMet_EFTu_GDP_PO4_aatRNAaaxyz = Species('elRS70SA'+xyz+'0002_fMet_EFTu_GDP_PO4_'+aa+'tRNA'+aa+xyz)
        elRS70SAxyz0002_fMet_EFTu_GTP_aatRNAaaxyz = Species('elRS70SA'+xyz+'0002_fMet_EFTu_GTP_'+aa+'tRNA'+aa+xyz)
        elRS70SAxyz0002_fMet_aatRNAaaxyz = Species('elRS70SA'+xyz+'0002_fMet_'+aa+'tRNA'+aa+xyz)

        ## B2_pept2
        elRS70SBxyz0002_Pept0002tRNAaaxyz = Species('elRS70SB'+xyz+'0002_Pept0002tRNA'+aa+xyz)
        elRS70SBxyz0002_Pept0002tRNAaaxyz_EFG_GDP_PO4 = Species('elRS70SB'+xyz+'0002_Pept0002tRNA'+aa+xyz+'_EFG_GDP_PO4')
        elRS70SBxyz0002_Pept0002tRNAaaxyz_EFG_GTP = Species('elRS70SB'+xyz+'0002_Pept0002tRNA'+aa+xyz+'_EFG_GTP')     

        Pept0002 = Species('Pept0002')
        Pept0002_degraded = Species('Pept0002_degraded')
        Pept0002tRNAaaxyz = Species('Pept0002tRNA'+aa+xyz)
        Pept0002tRNAaaxyz_degraded = Species('Pept0002tRNA'+aa+xyz+'_degraded')

        list_species_el2p2= [
            elRS70SAxyz0002_fMet_EFTu_GDP_aatRNAaaxyz, elRS70SAxyz0002_fMet_EFTu_GDP_PO4_aatRNAaaxyz, 
            elRS70SAxyz0002_fMet_EFTu_GTP_aatRNAaaxyz, elRS70SAxyz0002_fMet_aatRNAaaxyz,

            elRS70SBxyz0002_Pept0002tRNAaaxyz, elRS70SBxyz0002_Pept0002tRNAaaxyz_EFG_GDP_PO4,
            elRS70SBxyz0002_Pept0002tRNAaaxyz_EFG_GTP,  

            Pept0002, Pept0002_degraded, Pept0002tRNAaaxyz, Pept0002tRNAaaxyz_degraded]

        ## Reactions
        list_of_reaction_pept2=[
            Reaction.from_massaction([Species('EFTu_GTP_'+aa+'tRNA'+aa+xyz),elRS70SAxyz0002_fMet],[elRS70SAxyz0002_fMet_EFTu_GTP_aatRNAaaxyz], k_forward= rxn_k['re0000000013_k1']),
            Reaction.from_massaction([elRS70SAxyz0002_fMet_EFTu_GTP_aatRNAaaxyz],[elRS70SAxyz0002_fMet_EFTu_GDP_PO4_aatRNAaaxyz], k_forward= rxn_k['re0000000014_k1']),
 
            Reaction.from_massaction([elRS70SAxyz0002_fMet_EFTu_GDP_PO4_aatRNAaaxyz],[PO4,elRS70SAxyz0002_fMet_EFTu_GDP_aatRNAaaxyz], k_forward= rxn_k['re0000000016_k1']),
            Reaction.from_massaction([elRS70SAxyz0002_fMet_EFTu_GDP_aatRNAaaxyz],[EFTu_GDP,elRS70SAxyz0002_fMet_aatRNAaaxyz], k_forward= rxn_k['re0000000017_k1']),

            Reaction.from_massaction([elRS70SAxyz0002_fMet_aatRNAaaxyz],[elRS70SBxyz0002_Pept0002tRNAaaxyz], k_forward= rxn_k['re0000000018_k1']),
            Reaction.from_massaction([elRS70SBxyz0002_Pept0002tRNAaaxyz,EFG_GTP],[elRS70SBxyz0002_Pept0002tRNAaaxyz_EFG_GTP], k_forward= rxn_k['re0000000019_k1']),
            Reaction.from_massaction([elRS70SBxyz0002_Pept0002tRNAaaxyz_EFG_GTP],[EFG_GTP,elRS70SBxyz0002_Pept0002tRNAaaxyz], k_forward= rxn_k['re0000000020_k1']),
            Reaction.from_massaction([elRS70SAxyz0002_fMet_EFTu_GTP_aatRNAaaxyz],[Species('EFTu_GTP_'+aa+'tRNA'+aa+xyz),elRS70SAxyz0002_fMet], k_forward= rxn_k['re0000000021_k1']),
            Reaction.from_massaction([elRS70SBxyz0002_Pept0002tRNAaaxyz_EFG_GTP],[elRS70SBxyz0002_Pept0002tRNAaaxyz_EFG_GDP_PO4], k_forward= rxn_k['re0000000022_k1']),
            Reaction.from_massaction([elRS70SBxyz0002_Pept0002tRNAaaxyz_EFG_GDP_PO4],[elRS70SBxyz0002_Pept0002tRNAaaxyz_EFG_GTP], k_forward= rxn_k['re0000000023_k1']),
            ]
    #######################################################################################################################
    #The reading the 3rd+ amino acid to the second to last amino acid
    elif L<len(protein):

        elRS70SCxyzGpept_PeptSpepttRNAoAoX_EFG_GDP = Species('elRS70SC'+xyz+Gpept+'_Pept'+Spept+'tRNA'+oA+oX+'_EFG_GDP')
        
        elRS70SAxyzGpept_PeptSpept = Species('elRS70SA'+xyz+Gpept+'_Pept'+Spept)
        elRS70SAxyzGpept_PeptSpept_EFTu_GDP = Species('elRS70SA'+xyz+Gpept+'_Pept'+Spept+'_EFTu_GDP')
        elRS70SAxyzGpept_PeptSpept_EFTu_GDP_aatRNAaaxyz = Species('elRS70SA'+xyz+Gpept+'_Pept'+Spept+'_EFTu_GDP_'+aa+'tRNA'+aa+xyz)
        elRS70SAxyzGpept_PeptSpept_EFTu_GDP_PO4_aatRNAaaxyz = Species('elRS70SA'+xyz+Gpept+'_Pept'+Spept+'_EFTu_GDP_PO4_'+aa+'tRNA'+aa+xyz)
        elRS70SAxyzGpept_PeptSpept_EFTu_GTP_aatRNAaaxyz = Species('elRS70SA'+xyz+Gpept+'_Pept'+Spept+'_EFTu_GTP_'+aa+'tRNA'+aa+xyz)
        elRS70SAxyzGpept_PeptSpept_aatRNAaaxyz = Species('elRS70SA'+xyz+Gpept+'_Pept'+Spept+'_'+aa+'tRNA'+aa+xyz)
        elRS70SAxyzGpept_PeptSpepttRNAoAoX= Species('elRS70SA'+xyz+Gpept+'_Pept'+Spept+'tRNA'+oA+oX)

        elRS70SBxyzGpept_PeptGpepttRNAaaxyz_EFG_GTP = Species('elRS70SB'+xyz+Gpept+'_Pept'+Gpept+'tRNA'+aa+xyz+'_EFG_GTP')
        elRS70SBxyzGpept_PeptGpepttRNAaaxyz = Species('elRS70SB'+xyz+Gpept+'_Pept'+Gpept+'tRNA'+aa+xyz)
        elRS70SBxyzGpept_PeptGpepttRNAaaxyz_EFG_GDP_PO4 = Species('elRS70SB'+xyz+Gpept+'_Pept'+Gpept+'tRNA'+aa+xyz+'_EFG_GDP_PO4')

        PeptGpept = Species('Pept'+Gpept)
        PeptGpepttRNAaaxyz = Species('Pept'+Gpept+'tRNA'+aa+xyz)
        PeptGpepttRNAaaxyz_degraded = Species('Pept'+Gpept+'tRNA'+aa+xyz+'_degraded')

        #List of Species for peptide 3+ through n-1
        species_el3p3= [
            elRS70SCxyzGpept_PeptSpepttRNAoAoX_EFG_GDP, elRS70SAxyzGpept_PeptSpept, 
            
            elRS70SAxyzGpept_PeptSpept_EFTu_GDP, elRS70SAxyzGpept_PeptSpept_EFTu_GDP_aatRNAaaxyz, 
            elRS70SAxyzGpept_PeptSpept_EFTu_GDP_PO4_aatRNAaaxyz, elRS70SAxyzGpept_PeptSpept_EFTu_GTP_aatRNAaaxyz, 
            elRS70SAxyzGpept_PeptSpept_aatRNAaaxyz, elRS70SAxyzGpept_PeptSpepttRNAoAoX,

            elRS70SBxyzGpept_PeptGpepttRNAaaxyz_EFG_GTP, elRS70SBxyzGpept_PeptGpepttRNAaaxyz,
            elRS70SBxyzGpept_PeptGpepttRNAaaxyz_EFG_GDP_PO4,

            PeptGpept, PeptGpepttRNAaaxyz, PeptGpepttRNAaaxyz_degraded,
            ]
        
        #List of reactions for peptide 3+ through n-1
        rxn_pept3=[
            Reaction.from_massaction([Species('elRS70SB'+oX+Spept+'_Pept'+Spept+'tRNA'+oA+oX+'_EFG_GDP_PO4')],[PO4,elRS70SCxyzGpept_PeptSpepttRNAoAoX_EFG_GDP], k_forward= rxn_k['re0000000024_k1']), 
            Reaction.from_massaction([elRS70SCxyzGpept_PeptSpepttRNAoAoX_EFG_GDP],[elRS70SAxyzGpept_PeptSpepttRNAoAoX,EFG_GDP], k_forward= rxn_k['re0000000025_k1']), 

            Reaction.from_massaction([elRS70SAxyzGpept_PeptSpepttRNAoAoX],[elRS70SAxyzGpept_PeptSpept,Species('tRNA'+oA+oX)], k_forward= rxn_k['re0000000068_k1']),

            #Switched to the new aa
            Reaction.from_massaction([Species('EFTu_GTP_'+aa+'tRNA'+aa+xyz),elRS70SAxyzGpept_PeptSpept],[elRS70SAxyzGpept_PeptSpept_EFTu_GTP_aatRNAaaxyz], k_forward= rxn_k['re0000000074_k1']),
            
            Reaction.from_massaction([elRS70SAxyzGpept_PeptSpept_EFTu_GTP_aatRNAaaxyz],[elRS70SAxyzGpept_PeptSpept_EFTu_GDP_PO4_aatRNAaaxyz], k_forward= rxn_k['re0000000075_k1']),
            Reaction.from_massaction([elRS70SAxyzGpept_PeptSpept_EFTu_GDP_PO4_aatRNAaaxyz],[PO4,elRS70SAxyzGpept_PeptSpept_EFTu_GDP_aatRNAaaxyz], k_forward= rxn_k['re0000000077_k1']),
            Reaction.from_massaction([elRS70SAxyzGpept_PeptSpept_EFTu_GDP_aatRNAaaxyz],[EFTu_GDP,elRS70SAxyzGpept_PeptSpept_aatRNAaaxyz], k_forward= rxn_k['re0000000078_k1']),
            Reaction.from_massaction([elRS70SAxyzGpept_PeptSpept_aatRNAaaxyz],[elRS70SBxyzGpept_PeptGpepttRNAaaxyz], k_forward= rxn_k['re0000000079_k1']),

            Reaction.from_massaction([elRS70SBxyzGpept_PeptGpepttRNAaaxyz,EFG_GTP],[elRS70SBxyzGpept_PeptGpepttRNAaaxyz_EFG_GTP], k_forward= rxn_k['re0000000080_k1']),
            Reaction.from_massaction([elRS70SBxyzGpept_PeptGpepttRNAaaxyz_EFG_GTP],[EFG_GTP,elRS70SBxyzGpept_PeptGpepttRNAaaxyz], k_forward= rxn_k['re0000000081_k1']),
            Reaction.from_massaction([elRS70SAxyzGpept_PeptSpept_EFTu_GTP_aatRNAaaxyz],[Species('EFTu_GTP_'+aa+'tRNA'+aa+xyz),elRS70SAxyzGpept_PeptSpept], k_forward= rxn_k['re0000000082_k1']),
            Reaction.from_massaction([elRS70SBxyzGpept_PeptGpepttRNAaaxyz_EFG_GTP],[elRS70SBxyzGpept_PeptGpepttRNAaaxyz_EFG_GDP_PO4], k_forward= rxn_k['re0000000083_k1']),
            Reaction.from_massaction([elRS70SBxyzGpept_PeptGpepttRNAaaxyz_EFG_GDP_PO4],[elRS70SBxyzGpept_PeptGpepttRNAaaxyz_EFG_GTP], k_forward= rxn_k['re0000000084_k1']),

            Reaction.from_massaction([elRS70SAxyzGpept_PeptSpept_EFTu_GDP],[elRS70SAxyzGpept_PeptSpept,EFTu_GDP], k_forward= rxn_k['re0000000089_k1']),
            ]
        list_of_reaction_pept3.append(rxn_pept3)
        list_species_el3p3.append(species_el3p3)
    else:
        print('error')
        
#Flatten the reactions/species for aa 3+
list_of_reaction_pept3=flatten(list_of_reaction_pept3)
list_species_el3p3=flatten(list_species_el3p3)
```

#### Compiling TL CRN (CRN\_TL)¶

In [22]:

```
#Compile all reactions and species
All_species_TL= flatten([list_species_gen, list_species_elfmet, list_species_fmet,
                       flatten(list_species_aa), list_species_el2p2, list_species_el3p3,
                       list_species_el4p3, list_species_elterm])

All_rxn_TL = flatten([list_of_reaction_gen, list_reaction_fmet,
                   list_of_reactions, list_reactions_elfmet,
                   list_of_reaction_pept2, list_of_reaction_pept3,
                   list_of_reactions_end, list_of_reaction_term])
```

In [23]:

```
#Buliding the CRN_TL and saving as a SBML file
CRN_TL = ChemicalReactionNetwork(species = All_species_TL, reactions = All_rxn_TL)
# CRN_TL.write_sbml_file("PURE_TL_Model_degfp.xml")
```

### Linker reactions:¶

This section includes species and reactions to account for GFP folding and mean ribosome load.

In [24]:

```
deGFP_m = Species('deGFP_m')
gfp_species= [deGFP_m] 

Rxn_folding= [Reaction.from_massaction([PeptGpept],[deGFP_m], k_forward= .001667)] #based on bionumbers
CRN_linker= [Reaction.from_massaction([mRNA_i],[4*[mRNA]],k_forward=1000)]# to account for mean ribosome load
```

### Combining all CRNs: CRN\_TX + CRN\_TL + CRN\_linker¶

In [25]:

```
Combine_PURE=ChemicalReactionNetwork(species = flatten([CRN_TX.species, CRN_TL.species, gfp_species]),
                                     reactions = flatten([CRN_TX.reactions, CRN_TL.reactions,
                                                          Rxn_folding, CRN_linker]))
# Combine_PURE.write_sbml_file("Combine_PURE_Model_gfp.xml")
```

### Simulations of Combines Model of PURE¶

In [26]:

```
#Directory and file for the initial conditions
filename_ic = '/PURE_TXTL_initial_values_Final.csv'

with open(directory + filename_ic, mode='r') as infile:
    reader = csv.reader(infile)
    initial_con= {rows[0]:float(rows[1]) for rows in reader}
```

In [27]:

```
t0=time.time()
print(f"current time: {datetime.datetime.now()}")
timepoints = np.linspace(10**-5, 4*60*60, 500) 

PURE_Model_Final = Combine_PURE.simulate_with_bioscrape_via_sbml(timepoints = timepoints, initial_condition_dict = initial_con)
# PURE_Model_Final.to_csv(directory+'combined_mGapt-deGFP_5nM.csv')  

t1=time.time() 
print(f"PURE Simulation= {(t1-t0)/60} min")
```

```
current time: 2023-08-14 13:57:30.723072
PURE Simulation= 32.46270789305369 min
```

```
C:\Users\zoila\anaconda3\envs\python38\lib\site-packages\scipy\integrate\_odepack_py.py:248: ODEintWarning: Excess work done on this call (perhaps wrong Dfun type). Run with full_output = 1 to get quantitative information.
  warnings.warn(warning_msg, ODEintWarning)
```

#### Plotting simulations of combined PURE model¶

##### mRNA¶

In [28]:

```
p1 = bokeh.plotting.figure(toolbar_location='right',
    outline_line_color= None,
    min_border_right=10,
    height=400,
    width=600,)

p1.title.text = "A) Simulation Results of mRNA"
p1.xaxis.axis_label = 'Time (hours)'
p1.yaxis.axis_label = 'mRNA (μM)'
p1.outline_line_color=None

# p1.yaxis
p1.ygrid.visible = False
p1.yaxis.axis_label_text_font_size='15pt' 
p1.yaxis.major_label_text_font_size = '15pt'
p1.yaxis.major_label_text_font='Work Sans'
p1.yaxis.axis_label_standoff=15
p1.yaxis.axis_label_text_font_style='normal' 

# p1.xaxis
p1.xgrid.visible = False
p1.xaxis.axis_label_text_font_size='15pt' 
p1.xaxis.major_label_text_font_size = '15pt'
p1.xaxis.major_label_text_font='Work Sans'
p1.xaxis.axis_label_standoff=15
p1.xaxis.axis_label_text_font_style='normal' 

# p1.title
p1.title.text_font_size= '18pt' 
p1.title.align= 'left'
p1.title.offset=-65.0

# Model results
p1.line(PURE_Model_Final['time']/3600, PURE_Model_Final['mRNA_t'],legend_label = "BioCRNpyler Model", line_color="magenta", line_width=3, line_alpha=.5) #mRNA

p1.x_range=Range1d(0, 4)
p1.y_range=Range1d(0, 1)

# p1.legend
p1.legend.location='top_left'
p1.legend.background_fill_alpha= 0
p1.legend.border_line_color= None
p1.legend.label_text_font_size= '10pt'

bokeh.io.show(p1)
```

##### deGFP¶

In [29]:

```
p2 = bokeh.plotting.figure(toolbar_location='right',
    outline_line_color= None,
    min_border_right=10,
    height=400,
    width=600,)

p2.title.text = "B) Simulation Results of deGFP"
p2.xaxis.axis_label = 'Time (hours)'
p2.yaxis.axis_label = 'deGFP (μM)'
p2.y_range=Range1d(0, 8)
p2.x_range=Range1d(0, 4)
p2.outline_line_color=None

# p2.yaxis
p2.ygrid.visible = False
p2.yaxis.axis_label_text_font_size='15pt' 
p2.yaxis.major_label_text_font_size = '15pt'
p2.yaxis.major_label_text_font='Work Sans'
p2.yaxis.axis_label_standoff=15
p2.yaxis.axis_label_text_font_style='normal' 

# p2.xaxis
p2.xgrid.visible = False
p2.xaxis.axis_label_text_font_size='15pt' 
p2.xaxis.major_label_text_font_size = '15pt'
p2.xaxis.major_label_text_font='Work Sans'
p2.xaxis.axis_label_standoff=15
p2.xaxis.axis_label_text_font_style='normal' 

# p2.title
p2.title.text_font_size= '18pt' 
p2.title.align= 'left'

p2.title.offset=-65.0

# Model data
p2.line(PURE_Model_Final['time']/3600, PURE_Model_Final['deGFP_m'],legend_label = "BioCRNpyler Model", line_color='magenta', 
        line_dash= "solid", line_width=3,line_alpha=.5) 

# p2.xaxis
p2.ygrid.visible = False
p2.yaxis.axis_label_text_font_size='15pt' 
p2.yaxis.major_label_text_font_size = '15pt'
p2.yaxis.major_label_text_font='Work Sans'
p2.yaxis.axis_label_standoff=15
p2.yaxis.axis_label_text_font_style='normal' 

# p2.legend
p2.legend.location='top_left'
p2.legend.background_fill_alpha= 0
p2.legend.border_line_color= None
p2.legend.label_text_font_size= '10pt'

bokeh.io.show(p2)
```

#### Plot as Grid¶

In [30]:

```
# using grid layout on plots
pPURE_gfp=gridplot([[p1, p2]], width=500, height=400)
show(pPURE_gfp)
```

### Simulations of Combines Model of PURE for Mean Ribosome Load¶

Combining all CRNs: CRN\_TX + CRN\_TL + CRN\_linker (n=1-10)

In [31]:

```
##Uncomment below to run all 10 additional Mean Ribosome Load simulations
# for i in range(10):
#     n=i+1
#     CRN_linker= [Reaction.from_massaction([mRNA_i],[n*[mRNA]],k_forward=1000)] 
#     Combine_PURE=ChemicalReactionNetwork(species = flatten([CRN_TX.species, CRN_TL.species, gfp_species]),
#                                          reactions = flatten([CRN_TX.reactions, CRN_TL.reactions, 
#                                                               Rxn_folding, CRN_linker]))
#     t0=time.time()
#     PURE_Model_MRL= Combine_PURE.simulate_with_bioscrape_via_sbml(timepoints = timepoints, initial_condition_dict = initial_con)
#     PURE_Model_MRL.to_csv(directory+'/MGapt-deGFP_Model_MRL_runs/mGapt-deGFP_5nM_n'+str(n)+'.csv') 
#     print(f"PURE Simulation= {(t1-t0)/60} min")
```

### Computing environment¶

In [32]:

```
%load_ext watermark
%watermark -v -p bioscrape,bokeh,panel,jupyterlab,biocrnpyler
```

```
Python implementation: CPython
Python version       : 3.8.17
IPython version      : 8.12.2

bioscrape  : 1.2.1
bokeh      : 2.4.0
panel      : 0.13.1
jupyterlab : 3.6.5
biocrnpyler: 1.1.1
```
