## Supporting scripts, data, and simulation results. for "A chemical reaction network model of PURE": inference_run.sh

```
#!/bin/bash

#Submit this script with: sbatch thefilename

#SBATCH --time=20:00:00   # walltime
#SBATCH --ntasks=4   # number of processor cores (i.e. tasks)
#SBATCH --nodes=1   # number of nodes
#SBATCH --mem-per-cpu=64G   # memory per CPU core
#SBATCH -J "PURE_run"   # job name
#SBATCH --mail-user=   # email address
#SBATCH --mail-type=END
#SBATCH --mail-type=FAIL


# LOAD MODULES, INSERT CODE, AND RUN YOUR PROGRAMS HERE
source /home/apandey/anaconda3/bin/activate python3.7-bioscrape
python PURE_inference.py
```
