## Supporting scripts, data, and simulation results. for "A chemical reaction network model of PURE": PURE_inference.py

```
"""To run this file on HPC, simply run:
`sbatch inference_run.sh`
The results are written to `mcmc_results.csv`
"""
import numpy as np
import pandas as pd

import matplotlib.pyplot as plt

from bioscrape.types import Model
from bioscrape.inference import py_inference
from bioscrape.simulator import py_simulate_model

# ## Getting experimental data
filename = 'MGapt_deGFP_Exp_PURE_Data\2022.08.19_OnePot_PURE_mGapt_uM.csv'
mGapt_Data =  pd.read_csv(filename, delimiter = '\,', names = ['time','mRNA0', 'mRNA1','mRNA2', 'mRNA3'], engine='python',skiprows = 1)
first_row = mGapt_Data.iloc[0, 1:]  # extract the first row, excluding the 'time' column
mGapt_Data.iloc[:, 1:] -= first_row  # subtract the values in the first row from all other rows, excluding the 'time' column
# Truncate at 4 hours
truncated_data = mGapt_Data[mGapt_Data['time'] <= 4*3600]
# Define timepoints
timepoints = np.array(truncated_data['time'])

# Run model to test
m = Model(sbml_filename = "CRN_PURE_TX_mGaptv4_updated.xml")
initial_con={'RNAPa':(1), 'DNA':(.005), 'ATP':(3750), 'GTP':(2500), 'CTP':(1250), 'UTP':(1250),} #in uM
m.set_species(initial_con)
#### Try to fit with parameters/2 initial values for params. 
for param, param_value in m.get_parameter_dictionary().items():
    m.set_parameter(param, param_value/2)
# Simulate model
R0 = py_simulate_model(Model = m, timepoints = timepoints)
plt.figure(figsize=(6, 8))
plt.title("mRNA Concentration Over Time")
plt.plot(R0['time']/3600, R0['mRNA'], label = "mRNA", color = 'm') #mRNA

#Experimental data set 1
plt.scatter(truncated_data['time']/3600, truncated_data['mRNA1'], label = "T7-mGapt.R1", s=5, alpha=0.5, color="navy") #mRNA
plt.scatter(truncated_data['time']/3600, truncated_data['mRNA2'], label = "T7-mGapt.R2", s=5, alpha=0.5, color="blue")#mRNA
plt.scatter(truncated_data['time']/3600, truncated_data['mRNA3'], label = "T7-mGapt.R3", s=5, alpha=0.5, color= "steelblue") #mRNA
plt.xlabel('Time (h)')
plt.ylabel('mGapt (uM)')
plt.legend()
plt.savefig('tx_only_model-vs-data_params_by_2.png')


# # Bayesian Inference 

from bioscrape.inference import py_inference

import matplotlib.pyplot as plt
import numpy as np
from bioscrape.types import Model
import time
import pandas as pd
import warnings
from scipy.integrate import odeint
import seaborn as sns
import pandas as pd


m = Model(sbml_filename = "CRN_PURE_TX_mGaptv4_updated.xml")
initial_con={'RNAPa':(1), 'DNA':(.005), 'ATP':(3750), 'GTP':(2500), 'CTP':(1250), 'UTP':(1250),} #in uM
m.set_species(initial_con)
#### Try to fit with parameters/2 initial values for params. 
for param, param_value in m.get_parameter_dictionary().items():
    m.set_parameter(param, param_value/2)
# Import data from CSV

num_trajectories = 3
exp_data = []
for i in range(num_trajectories):
    df = pd.DataFrame()
    df["mRNA"]=truncated_data['mRNA'+str(i)]
    df["timepoints"] = truncated_data["time"]
    exp_data.append(df)
#exp_data

# Choose the "true" parameters.
k_2 = 0.08/2
k_start = 0.06/2
prior = {'k_forward2' : ['gaussian', k_2, 0.1, 'positive'],
         'k_forward4' : ['gaussian', k_start, 0.1, 'positive']
         }

sampler, pid = py_inference(Model = m, exp_data = exp_data, measurements = ['mRNA'],
                            time_column = ['timepoints'], params_to_estimate = ['k_forward2', 'k_forward4'],
                            nwalkers = 20, nsteps = 2000, init_seed = 'prior', prior = prior,
                            sim_type = 'deterministic', plot_show = False,)

# # Recommended to simply use sampler object/mcmc_results.csv and generate your own custom plots
# truth_list, uncertainty_list = pid.plot_mcmc_results(sampler);
```
