## Supporting scripts, data, and simulation results. for "A chemical reaction network model of PURE": README.txt

This README file outlines the documents and subfolders in this folder.
The folder contains all Python scripts used to simulate protein expression and plot figures seen in the paper "A chemical reaction network model of PURE" by Zoila Jurado, Ayush Pandey, and Richard M. Murray. Additionally, all simulation runs and experimental results are included.

Scripts:
-CRN_PURE_Model_Figures: Script used to plot all figures.
-CRN_PURE_TLonly_Final: Expanded translation model of PURE based on MATLAB model by Matsuura, et al.
-CRN_PURE_TXonly_Final: Transcription model of PURE based on Tuza, et al.
-CRN_PURE_TXTL_MGaptdeGFP_FINAL: The proposed combined transcriptions (TX) and translation (TL) model of PURE for plasmid pT7-MGapt-UTR1-deGFP-tT7. Returns PURE_TX_Model_degfp.xml, PURE_TL_Model_degfp.xml, and Combine_PURE_Model_gfp.xml Sub-SBML files.
-PURE_inference: Parameter inference script used to identify reaction rates for the TX-only model. Uses CRN_PURE_TX_mGaptv4_updated.xml file and data in MGapt_deGFP_Exp_PURE_Data folder. 
-sensitivity_analysis: Sensitivity analysis script used to identify sensitive parameters for the TX-only model.

Folders:
-Calibrations_Biotek: Contains calibration data of MGapt and deGFP from Biotek 4.
-CRN_PURE_TLonly: Contains the first iteration of the PURE TL-only model, including initial conditions. 
-Figures_PURE_Model: Figures generated by CRN_PURE_Model_Figures.ipynb, as seen in paper.
-MGapt_deGFP_Exp_PURE_Data: Experimental results of NEB PURExpress of respective plasmids.
-MGapt-deGFP_Model_MRL_runs: Simulation results of the combined PURE model of mean ribosome load tests.
-TL_extension_arbAA: Simulation results of the PURE TL-only model of peptide test.

Etc:
-combined_mGapt-deGFP_5nM: Simulation results for expression of pT7-MGapt-UTR1-deGFP-tT7 plasmid in CRN_PURE_TXTL_MGaptdeGFP_FINAL.ipynb.
-CRN_PURE_TLonly_MGaptdeGFP_results: Simulation results for expression of pT7-MGapt-UTR1-deGFP-tT7 plasmid in CRN_PURE_TXTL_MGaptdeGFP_FINAL.ipynb.
-fMGG_MATLAB_Matsuura: MATLAB simulation results for translation of peptide fMGG (Matsuura 2017). See https://sites.google.com/view/puresimulator for more details.

-fMGG_synthesis_parameters_CRN: Parameters download from https://sites.google.com/view/puresimulator (Matsuura 2017).
-PURE_TXTL_initial_values_Final: Initial conditions used in CRN_PURE_TXTL_MGaptdeGFP_FINAL.ipynb.

The required packages to run scripts on Anaconda 3.8.16 are listed below:
•bioscrape
•biocrnpyler
•pyviz 
•holoviz
•nodejs
•black 
•blackcellmagic
•watermark 
•jupyter lab
•seaborn
•svglib
•scikit-learn
•openpyxl
•bokeh-catplot
•selenium     
•bokeh 
•jupyter_bokeh ==2.0.0         
•nbconvert
•panel==0.13.1
