## Supporting scripts, data, and simulation results. for "A chemical reaction network model of PURE": sensitivity_analysis.py

```
"""To run this file on HPC, simply run:
`sbatch sensitivity_analysis.sh`
"""
import numpy as np
import pandas as pd

from bioscrape.simulator import py_simulate_model, ModelCSimInterface, DeterministicSimulator
from bioscrape.inference import py_inference
from bioscrape.analysis import py_sensitivity_analysis

import matplotlib.pyplot as plt
from bioscrape.types import Model
import time
import pandas as pd
import warnings
import seaborn as sn

initial_con={'RNAPa':(0.1), 'DNA':(.005), 'ATP':(3000), 'GTP':(3000), 'CTP':(1000), 'UTP':(1000), 'Mg':(4000)} #in uM
# New ic from Matsuura response
initial_con={'RNAPa':(1), 'DNA':(.005), 'ATP':(3750), 'GTP':(2500), 'CTP':(1250), 'UTP':(1250),} #in uM

timepoints = np.linspace(10**-5, 1.44*10**4, 20)
# timepoints = np.linspace(10**-5, 2*10**4, 20)

m = Model(sbml_filename = "CRN_PURE_TX_17Nov_updated.xml")
m.set_species(initial_con)
ssm = py_sensitivity_analysis(model = m, timepoints = timepoints, normalize = True)
SSM=np.around(ssm[:,:,4].T, decimals=3, out=None)
R0 = py_simulate_model(Model = m, timepoints = timepoints)
import seaborn as sn
figsize = (15,8)
fig, ax = plt.subplots(figsize=figsize)
index = 1
params_names_latex = ['$k_{rev}$', '$k_{pi}$', '$k_{ntpdeg}$', '$k_{bound}$', '$k_{start}$', '$k_{ntpadd}$',
                      '$k_{ntpdis}$','$k_{ppi}$', '$k_{mrna}$']
sn_ax = sn.heatmap(SSM, ax = ax, annot=False, cmap = 'YlGnBu',
                   xticklabels = np.linspace(timepoints[0]/3600,timepoints[-1]/3600, len(timepoints),
                                             endpoint = True, dtype = 'int'))
sn_ax.figure.axes[-1].xaxis.label.set_size(20)
ax = fig.axes
_ = plt.xlabel('Time (h)', fontsize = 14)
_ = plt.ylabel('Parameters', fontsize = 14)
_ = ax[0].tick_params(axis='x', which='major', labelsize=12, bottom = False)
_ = ax[0].tick_params(axis='y', which='major', labelsize=12, left = False)
_ = ax[0].set_yticklabels(params_names_latex)
every_nth = 4
for n, label in enumerate(ax[0].xaxis.get_ticklabels()):
    if n == len(timepoints)-1:
        continue
    if n % every_nth != 0:
        label.set_visible(False)
_ = ax[1].tick_params(axis = 'x', labelsize = 18)
_ = plt.savefig('mrna-sensitivity.svg')
plt.show()
```
